# Enhanced metanephric specification to functional proximal tubule enables toxicity screening and infectious disease modelling in kidney organoids

**DOI:** 10.1101/2021.10.14.464320

**Authors:** Jessica M. Vanslambrouck, Sean B. Wilson, Ker Sin Tan, Ella Groenewegen, Rajeev Rudraraju, Jessica Neil, Kynan T. Lawlor, Sophia Mah, Michelle Scurr, Sara E. Howden, Kanta Subbarao, Melissa H. Little

## Abstract

While pluripotent stem cell-derived kidney organoids are now being used to model renal disease, the proximal nephron remains immature with limited evidence for key functional solute channels. This may reflect early mispatterning of the nephrogenic mesenchyme and/or insufficient maturation. Here we show that enhanced specification to metanephric nephron progenitors results in elongated and radially aligned proximalised nephrons with distinct S1 - S3 proximal tubule cell types. Such PT-enhanced organoids possess improved albumin and organic cation uptake, appropriate KIM-1 upregulation in response to cisplatin, and improved expression of SARS-CoV-2 entry factors resulting in increased viral replication. The striking proximo-distal orientation of nephrons resulted from localized WNT antagonism originating from the organoid stromal core. PT-enhanced organoids represent an improved model to study inherited and acquired proximal tubular disease as well as drug and viral responses.

## Introduction

The proximal tubules (PTs) of the kidney represent a highly specialised portion of the nephron performing the bulk of kidney reabsorption and secretion. This occurs via three distinct functional and anatomical segments: the convoluted (S1 and S2) and the straight (S3) segments that traverses the cortico-medullary boundary, with S1 exhibiting the highest capacity for solute, sodium, amino acid, and fluid transport (Zhuo and Li, 2013). Their unique roles and high metabolic activity render the PTs acutely vulnerable to toxins and metabolic stress (Kirita, *et al*., 2020). As such, accurately patterned and segmented PTs would represent a critical tool for drug development, toxicology research, and studies of PT dysfunction.

We and others have established robust protocols for the directed differentiation of human pluripotent stem cells to kidney (Freedman, *et al*., 2015; Morizane, *et al*., 2015; Taguchi and Nishinakamura, 2017; Takasato, *et al*., 2015; Toyohara, *et al*., 2015). While these organoids display a remarkable transcriptional similarity to the developing human kidney (Combes, *et al*., 2019; Howden, *et al*., 2021; Subramanian, *et al*., 2019; Wu, *et al*., 2018), their nephron patterning and segmentation remains immature, more closely resembling human trimester 1 fetal tissue (Takasato, *et al*., 2015). PT maturation and functional segmentation is particularly underdeveloped. Despite possessing nuclear HNF4A (responsible for driving early proximal patterning [(Marable, *et al*., 2020)]) and apical CUBILIN-MEGALIN complex expression, organoid PTs lack a range of functional solute channels that define each PT subsegment (Wu, *et al*., 2018; Wilson, *et al*., 2021). Expression levels of the principle water transporting channel, AQP1, the organic anion transporters (OATs), and the organic cation transporters (OCTs) are all low (Wilson, *et al*., 2021).

Such suboptimal PT maturation may represent inappropriate anteroposterior/mediolateral patterning, suboptimal maintenance of progenitor identity or incomplete maturation. In response to distinct temporospatial signalling, the permanent (metanephric) kidney arises during human embryogenesis as the final of three embryonic excretory organs, developing sequentially from specific rostrocaudal regions of the intermediate mesoderm (Dressler, 2009). Metanephric development, commencing during weeks 4 -5 of gestation, is preceded by the formation of two more rostral transient organs; the pronephros (human gestation week 3 – 4) and the mesonephros (human gestation week 4 – 10) (de Bakker, *et al*., 2019). While the mammalian pronephros is highly rudimentary, mesonephric nephrons also arise via MET and show similar patterning and segmentation to early metanephric nephron. However, the mesonephros possesses less definitive distal tubule segments and regresses around week 8 (Georgas, *et al*., 2011; Mugford, *et al*., 2008; Tiedemann, *et al*., 1987).

Using fluorescent reporter lines and lineage tracing in human kidney organoids, we have confirmed both the presence of a SIX2^+^ nephron progenitor population and the contribution of these cells to nephrogenesis via MET in kidney organoids (Howden, *et al*., 2019; Vanslambrouck, *et al*., 2019). However, the possibility exists that we are modelling mesonephric rather than metanephric nephrogenesis, potentially contributing to poor PT patterning and maturation (reviewed in (Little and Combes, 2019). It is also possible that suboptimal maintenance of progenitor identity during iPSC differentiation *in vitro* limits nephron maturation. Several media have been described that are able to support the maintenance of isolated nephron progenitors *in vitro* (Brown, *et al*., 2015; Li, *et al*., 2016; Tanigawa, *et al*., 2015; Tanigawa, *et al*., 2016). While each media contains low levels of canonical WNT activity and FGF2/9, distinct differences in nephron patterning result from the inclusion of a variety of TGFβ superfamily agonists (BMP4, BMP7, Activin A) and antagonists (A83-01, LDN193189), NOTCH inhibition (DAPT), and other growth factors (TGFα, IGF1/2, LIF). The inclusion of LDN193189 (inhibitor of BMP receptor-mediated SMAD1/5/8) supported tubular patterning but not formation of glomeruli (Brown, *et al*., 2015). In contrast, the addition of LIF and either dual-SMAD inhibition (LDN193189 and A83-01) or NOTCH inhibition (DAPT) resulted in the formation of nephrons with podocytes but different nephron morphologies (Li, *et al*., 2016; Tanigawa, *et al*., 2016). Finally, while proximodistal nephron patterning in mouse has previously been shown to be influenced by relative Wnt, Bmp, and Notch signalling in mouse (Lindstrom, *et al*., 2015), these data suggest that distinct nephron progenitor states may show varying competence for different nephron segments, or that distinct SIX2 populations give rise to different regions of the nephron.

In the current study, we sought to understand whether anteroposterior/mediolateral patterning, or shifts in commitment state of the nephron progenitors, could influence ultimate PT identity and maturation. Patterning to a posterior metanephric SIX2^+^ nephron progenitor population by extending the duration of mesodermal patterning, while simultaneously enhancing nephron progenitor expansion, specified progenitors with improved metanephric identity without influencing anteroposterior/mediolateral patterning. These progenitors formed strongly proximalised, elongated, and spatially aligned nephrons. The PTs within these nephrons displayed distinct segmentation into S1, S2 and S3 cell types, upregulation of key solute channels and transporters, and functional uptake of albumin and organic cations. Treatment with cisplatin elicited upregulation of Kidney Injury Marker-1 (KIM-1), while increased expression of key viral entry factors enabled improved SARS-CoV-2 infection and replication compared to standard protocols. Notably, the striking nephron alignment was shown to result from localised WNT antagonism, supporting a role for WNT gradients in human nephron proximodistal patterning. Taken together, this study suggests a requirement for optimal nephron progenitor commitment for appropriate PT identity. Such PT-enhanced kidney organoids represent a model of the human proximal nephrons with likely applications for infectious and genetic disease research, drug development, and nephrotoxicity evaluation.

## Results

### Prolonged monolayer culture and delayed nephron induction supports nephron progenitors

As noted previously, optimisation of nephron progenitor maintenance *in vitro* has been investigated by a range of studies using murine and human pluripotent stem cell-derived nephron progenitors (Brown, *et al*., 2015; Li, *et al*., 2016; Tanigawa, *et al*., 2016). While all studies reported maintenance of nephron progenitors, variations were evident with respect to the final patterning of resulting nephrons following induction. Given the clear influence that initial differentiation conditions and timing can have on nephron progenitor survival and subsequent nephron patterning, we hypothesised that expanding our nephron progenitor population whilst delaying nephron initiation may create a more metanephric population leading to organoids with improved patterning and PT maturation. We have previously shown that SIX2 expression is not detected until day 10 of pluripotent stem cell differentiation (Howden, *et al*., 2019). Hence, the initial monolayer differentiation phase was prolonged to between 12 – 14 days, along with culture in either of two previously defined nephron progenitor (NP) maintenance media, NPSR (Li, *et al*., 2016) and CDBLY (Tanigawa, *et al*., 2016) from day 7, which represents the point of intermediate mesoderm commitment (Takasato, *et al*., 2015; Takasato, *et al*., 2014) (Figure 1A). Compared to control media (TeSR-E6; E6), both NPSR and CDBLY prevented spontaneous epithelialisation of the monolayer (Figure 1B). However, very little epithelialisation and poor nephron commitment was observed after culture in NPSR (Figure 1B). In contrast, CDBLY preserved the nephron-forming capacity of the progenitor cells following their formation into a micromass and induction of nephrogenesis with a pulse of canonical WNT signalling) (Figure 1B). Nephrons of these organoids were also observed to surround a stromal core region that stained positive for markers of kidney stroma MEIS1/2/3 and SNAI2 (SLUG) (Supplementary Figure 1A) (England, *et al*., 2020). Upon prolonged organoid culture (> 14 days), portions of this core region formed patches of Alcian blue-positive cartilage (Supplementary Figure 1B).

**Figure 1:**
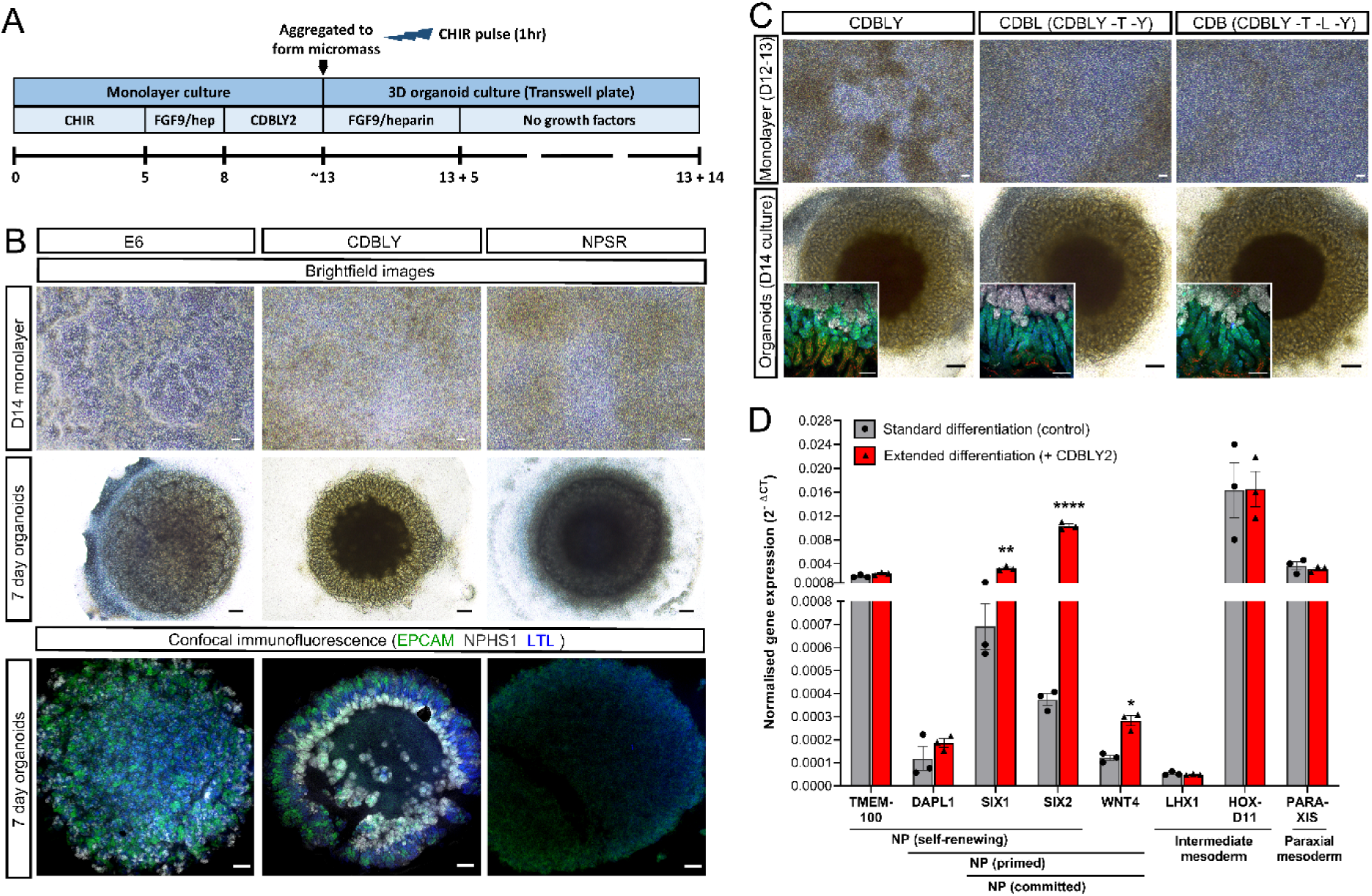
Extended monolayer culture in CDBLY supports nephron progenitors and preserves nephrogenic capacity. **A.** Schematic depicting the extended differentiation protocol in CDBLY2. **B.** Brightfield and confocal immunofluorescence images of extended monolayer differentiations in E6, CDBLY and NPSR, and resulting organoids. Immunofluorescence depicts nephrons (EPCAM; green), podocytes of glomeruli (NPHS1; grey), and proximal tubules (LTL; blue). Scale bars represent 100µm (monolayers) and 200µm (organoids). **C.** Brightfield images of extended monolayer differentiations using CDBLY variations and resulting organoids, with inset confocal immunofluorescence images highlighting organoid nephron alignment and patterning. Immunofluorescence depicts nephrons (EPCAM; green), podocytes of glomeruli (NPHS1; grey), proximal tubules (HNF4A; blue), and Loop of Henle (SLC12A1; red). Scale bars represent 100µm (monolayer brightfields and organoid immunofluorescence) and 200µm (organoid brightfields). **D.** qRT-PCR analysis of standard and extended monolayer differentiations. Error bars represent SEM from n = 3 biological replicates. Statistical significance was determined using an unpaired t test. Asterisks represent P values adjusted for multiple comparisons using the Holm-Sidak method, alpha = 0.05 (*; P ≤ 0.05, **; P ≤ 0.01, ***; P ≤ 0.001, ****; P ≤ 0.0001).

The prevention of spontaneous differentiation while preserving the nephrogenic capacity of the NP cells was found to be primarily a response to the presence of CDB (CHIR, DAPT, BMP7), with omission of LIF, Y27632, as well as the basal media component TGFα, found to produce a similar result with respect to growth, morphology and nephron segmentation compared to CDBLY (Figure 1C). The inhibition of monolayer epithelialisation with preserved nephrogenic capacity was found to be consistent at monolayer differentiation lengths tested (10, 12, 13 and 14 days) (Supplementary Fig 1C). However, a monolayer differentiation length of 12 – 13 days produced more consistent nephrogenesis between experiments, with 14 days observed to cause frequent detachment of the differentiating monolayer. Subsequent studies proceeded using prolonged culture in CDBLY noting the inclusion of an increased concentration of BMP7 (10ng/mL; CDBLY2) which improved the consistency of nephrogenesis between organoids compared to standard CDBLY (5ng/mL BMP7) (Supplementary Figure 1D). This modified differentiation protocol is detailed in Figure 1A.

Quantitative RT-PCR (qRT-PCR) of the extended monolayer differentiations in CDBLY2 confirmed an improved metanephric gene expression profile compared to standard differentiations performed in parallel (7 day protocol in E6 (Howden, *et al*., 2019; Takasato, *et al*., 2016)) (Figure 1D). Extended CDBLY2 monolayers showed a significant increase in *SIX1*/*SIX2* (self-renewing to committed NPs) and *WNT4* (primed to committed NPs), while *DAPL1* (self-renewing and primed NPs) was increased without significance and no change was observed in *TMEM100* (self-renewing NPs). This suggested that the extended protocol promotes a primed/committed, rather than self-renewing, NP population (Hochane, *et al*., 2019; Lindstrom, *et al*., 2018; Lindstrom, *et al*., 2018). Extended differentiation in CDBLY2 was not found to alter mediolateral patterning, with no change in paraxial mesodermal marker *PARAXIS* and unchanged or increased expression of intermediate mesoderm markers *HOXD11* and *LHX1* (Mugford, *et al*., 2008) (Figure 1D).

### Extended monolayer culture induces SIX2-derived proximalised nephrons

Lineage tracing studies in mouse have shown that nephrons are derived entirely from Six2+ nephron progenitors (Kobayashi, *et al*., 2008), with histological studies suggesting a similar developmental process in human (Lindstrom, *et al*., 2018; Lindstrom, *et al*., 2018) (Kobayashi, *et al*., 2008). Using a SIX2^Cre/Cre^:GAPDH^dual^ lineage tracing line, in which SIX2 expression induces a permanent GFP/mCherry switch, we have previously shown that kidney organoid nephrons contain cells derived from SIX2^+^, but also SIX2^-^, progenitor cells, resulting in a chimeric appearance (Howden, *et al*., 2019). To confirm and compare the competence of the metanephric progenitor-enriched monolayer differentiation to contribute to nephron formation, organoids were generated from both our standard protocol and the extended differentiation protocol using the SIX2^Cre/Cre^:GAPDH^dual^ lineage tracing line. Immunofluorescence re-confirmed the chimeric contribution of SIX2^+^ and SIX2^-^ progenitor-derived cells to standard organoid nephrons as shown previously (Howden, *et al*., 2019) (Figure 2A). However, confocal imaging suggested a larger contribution of SIX2+ cells to proximal nephrons in organoids derived from the extended protocol compared to the standard protocol (7 days differentiation, cultured in E6) (Howden, *et al*., 2019), including contribution to NPHS1^+^ podocytes, LTL^+^ PTs, and to a lesser extent E-CADHERIN^+^ distal tubules (Figure 2A). To quantitatively compare the contributions SIX2-derived cells to nephrons, dissociated SIX2^Cre/Cre^:GAPDH^dual^ standard and extended organoids (expressing endogenous SIX2-mCherry) were co-stained with EPCAM to mark both proximal and distal nephron epithelium, then analysed via flow cytometry (Figure 2Bi). In agreement with confocal imaging, SIX2-derived cell contribution to EPCAM^+^ nephrons was significantly higher in organoids derived from the metanephric progenitor-enriched extended monolayers compared to those derived from the standard 7 day protocol in E6 media, suggesting improved metanephric identity of prolonged monolayers exposed to CDBLY2 (Figure 2Bii).

**Figure 2:**
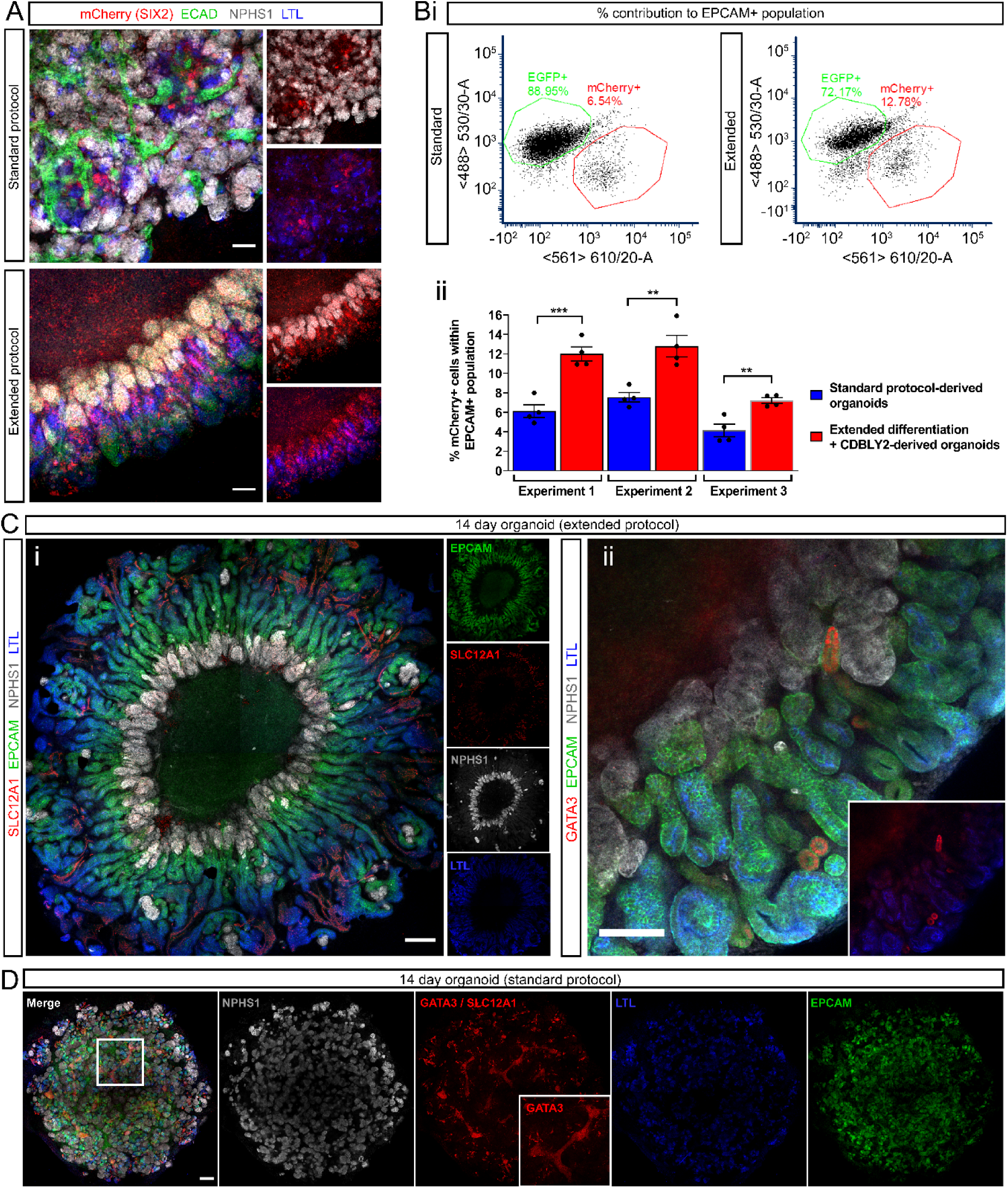
Extended monolayer culture in CDBLY2 increases SIX2+ progenitor contribution to nephrons and proximalisation. **A.** Confocal immunofluorescence of D7+14 (standard protocol) and D13+14 (extended protocol) organoids derived from the SIX2^Cre/Cre^:GAPDH^dual^ lineage tracing iPSC line. Images depict merged and separated channels showing lineage-traced SIX2+ cells (mCherry; red), distal tubules (ECAD; green), podocytes (NPHS1; grey) and proximal tubules (LTL; blue). Scale bars represent 100 µm. **B.** Flow cytometry of SIX2^Cre/Cre^:GAPDH^dual^ lineage tracing organoids derived from extended (13 day + CDBLY2) and standard (7 day + E6 media) differentiations depicting mCherry contribution to the EPCAM+ (nephron) population. Flow plots shown in (**i**) are representative of the replicates across multiple experiments. Percentage mCherry contributions from flow cytometry are depicted in bar graph (**ii**). Error bars in (**ii**) represent SEM from n = 4 biological replicates across 3 independent experiments. Statistical significance was determined using an unpaired t test. Asterisks represent P values adjusted for multiple comparisons using the Holm-Sidak method, alpha = 0.05 (*; P ≤ 0.05, **; P ≤ 0.01, ***; P ≤ 0.001, ****; P ≤ 0.0001). **C.** Confocal immunofluorescence of D13+14 organoids demonstrating (**i**) aligned nephron morphology with nephron segmentation makers (nephron epithelium [EPCAM; green], distal tubule/Loop of Henle [SLC12A1; red], proximal tubules [LTL; blue], and podocytes [NPHS1; grey]) and (**ii**) the presence of few GATA3+ connecting segment/ureteric epithelium structures (red), co-stained for nephron epithelium (EPCAM; green), podocytes (NPHS1; grey), and proximal tubules (LTL; blue). Inset in (**ii**) shows GATA3 and LTL alone. Scale bars in (**i**) and (**ii**) represent 200 µm and 100 µm, respectively. **C**. Confocal immunofluorescence of a D7+14 (standard) organoid depicting homogenous distribution of podocytes (NPHS1; grey), proximal tubules (LTL; red), and nephron epithelium (EPCAM; green), as well as the presence of extended segments of centralised GATA3+ connecting segment/ureteric epithelium (also highlighted in insets). Scale bar represents 200 µm.

The segmentation of nephrons within organoids derived from the extended protocol was examined using a range of markers for podocytes, proximal, and distal tubules, revealing distinct proximo-distal segmentation (Figure 2Ci). In contrast to the standard protocol which produced organoids with a branching GATA3+ epithelium (Figure 2D), extended protocol-derived organoids possessed few structures expressing the ureteric epithelium marker GATA3 (Figure 2Cii). The distribution of glomeruli, marked by NPHS1+ podocytes, also differed between protocols, with extended protocol-derived organoids possessing a central ring of glomeruli and elongated PTs radiating outwards that starkly opposed the more homogenous distribution of these structures in standard organoids (Figure Ci and D). This unique organoid morphology was observed in organoids derived from 6 different iPSC lines with or without gene editing and from male or female iPSC sources (3 examples evidenced in Supplementary Figure 1E).

In addition to differences in the segmentation of nephrons, organoids derived via extended differentiation in CDBLY2 appeared to possess a larger proportion of LTL- and HNF4A- positive PT compared to standard organoids (Figure 2Ci, Figure 2D, and Figure 3A). To quantify and compare the proportion of PT cells in organoids derived from these two protocols, organoids were generated using the HNF4A^YFP^ iPSC reporter line which reports the formation of PT (Vanslambrouck, *et al*., 2019) (Figure 3Bi). Flow cytometry revealed up to 6.2 times higher average proportions of HNF4A^YFP+^ PT cells in organoids derived from the extended monolayer protocol compared to the standard protocol (Figure 3Bii), confirming the use of extended monolayer differentiation combined with progenitor-supportive media, CDBLY2, as an effective method of generating proximal tubule-enhanced (PT-enhanced) kidney organoids.

**Figure 3:**
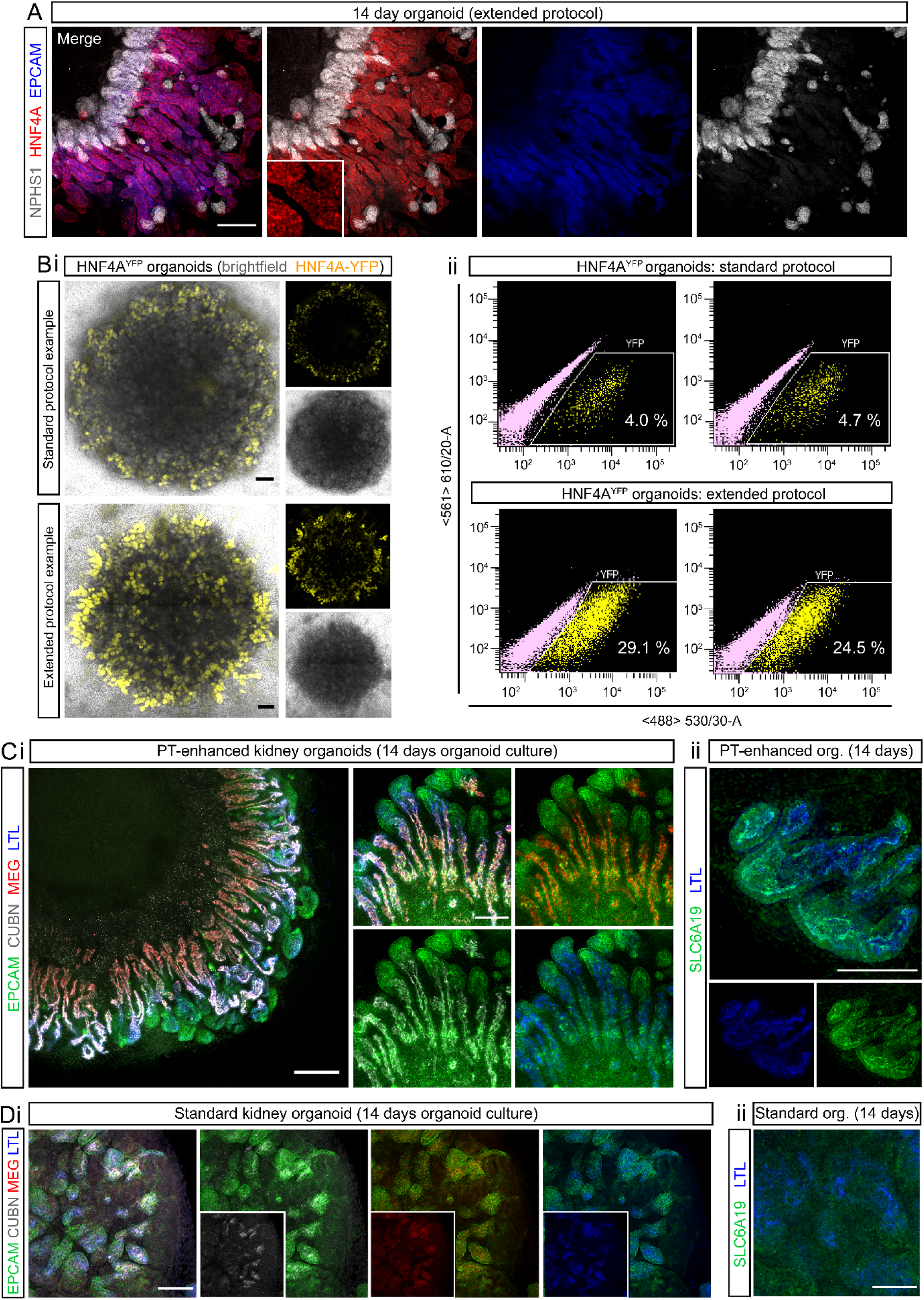
Enhanced organoids express mature and appropriately localised PT transporter proteins and transcription factors. **A.** Confocal immunofluorescence of HNF4A protein expression (red) in EPCAM-positive PTs (blue) of a day 14 organoid derived from extended differentiation of iPSCs in CDBLY2. NPHS1 (grey) marks podocytes of the glomeruli. Inset depicts higher magnification of HNF4A-expressing PT segments emphasising nuclear localisation. Scale bar represents 100 µm. **Bi.** Live confocal microscopy of 2 representative standard and extended protocol-derived organoids (also shown in [**Bii**]) generated using the HNF4A^YFP^ fluorescent iPSC reporter line (PCS-201-010/HNF4A^YFP^). YFP (yellow) marks proximal tubules. Transmitted light channel (brightfield) is shown as merged and separate images. Scale bars represent 200 µm. **Bii**. Flow cytometry plots of the 2 representative HNF4A^YFP^ organoids from experiment depicted in (**Bi**), derived from standard and extended protocols. **C-D.** Confocal immunofluorescence of PT-enhanced (**C**) and standard (D) (D13+14) organoids showing PT markers within EPCAM+ (green) nephrons, including LTL (blue; [**i**] and [**ii**]), CUBILIN (CUBN; grey [**i**]), MEGALIN (MEG; red [**i**]), and SLC6A19 (green [**ii**]). Scale bars represent 200 µm (**Ci**) and 100 µm (**Di-ii** and **Cii**).

### PT-enhanced organoids show improved proximal tubule patterning and maturation at both a protein and gene level

To establish the level of PT maturation within enhanced organoids, the expression and cellular localisation of functionally important brush border membrane proteins and markers, characteristic of mature PTs, were assessed via immunofluorescence (Figure 3C). Within LTL- positive PTs, enhanced organoids showed strong expression of the protein transport complex CUBILIN-MEGALIN (CUBN-MEG) and neutral amino acid transporter SLC6A19, with all transporters displaying a highly-specific apical brush border membrane localisation (Figure 3Ci-ii). In contrast, the PTs of standard organoids possessed weaker and diffuse staining of the CUBN-MEG complex (Figure 3Di). Furthermore, the majority of standard organoids lacked SLC6A19 expression, with staining observed in just one of three independent experiments (representative images in Figure 3Dii and Supplementary Figure 2A). Additional information regarding the maturity of PT brush border membranes was afforded by high-resolution imaging of LTL binding. LTL is a fucose-specific lectin widely used in the kidney field owing to its high-affinity binding to α-linked L fucose-containing oligosaccharides of glycoconjugates that abundantly line the brush border membrane of kidney PT cells (Hennigar, *et al*., 1985). High-resolution imaging of PTs within enhanced organoids showed LTL binding was highly restricted to the apical membrane where it co-localised with SLC6A19, a characteristic of correctly polarised, mature PT brush-border membranes (Supplementary Figure 2A). In contrast, the PTs of standard organoids possessed LTL staining that was not highly apically-restricted and instead diffuse throughout the PT, even in the instance where apical SLC6A19 was detected (Figure 2Dii and Supplementary Figure 2A). Taken together, these data suggested a more immature PT phenotype and suboptimal brush border membrane development in standard compared to enhanced organoids.

To provide a more comprehensive comparison with existing kidney organoid differentiation protocols, as well as to gain a deeper insight into the complexity and maturity of cells derived from the extended protocol, multiplexed single cell RNA sequencing (scRNAseq) with antibody-based cell barcoding was performed on both monolayer (day 13) and resulting PT- enhanced organoids (Figure 4). To account for variation, libraries were created from 4 separate differentiated monolayers representing distinct starting pools of iPSCs (CRL1502.C32) that were used to generate 4 separate batches of organoids (Figure 4A). Cells from the 4 replicates (both at day 13 [D13] monolayer stage, prior to organoid formation, and day 14 of organoid culture [D13+14]) were barcoded using hashing antibodies before being pooled. This approach produced a single library for each timepoint (sample) which could be later deconvoluted to retrieve replicate information.

**Figure 4:**
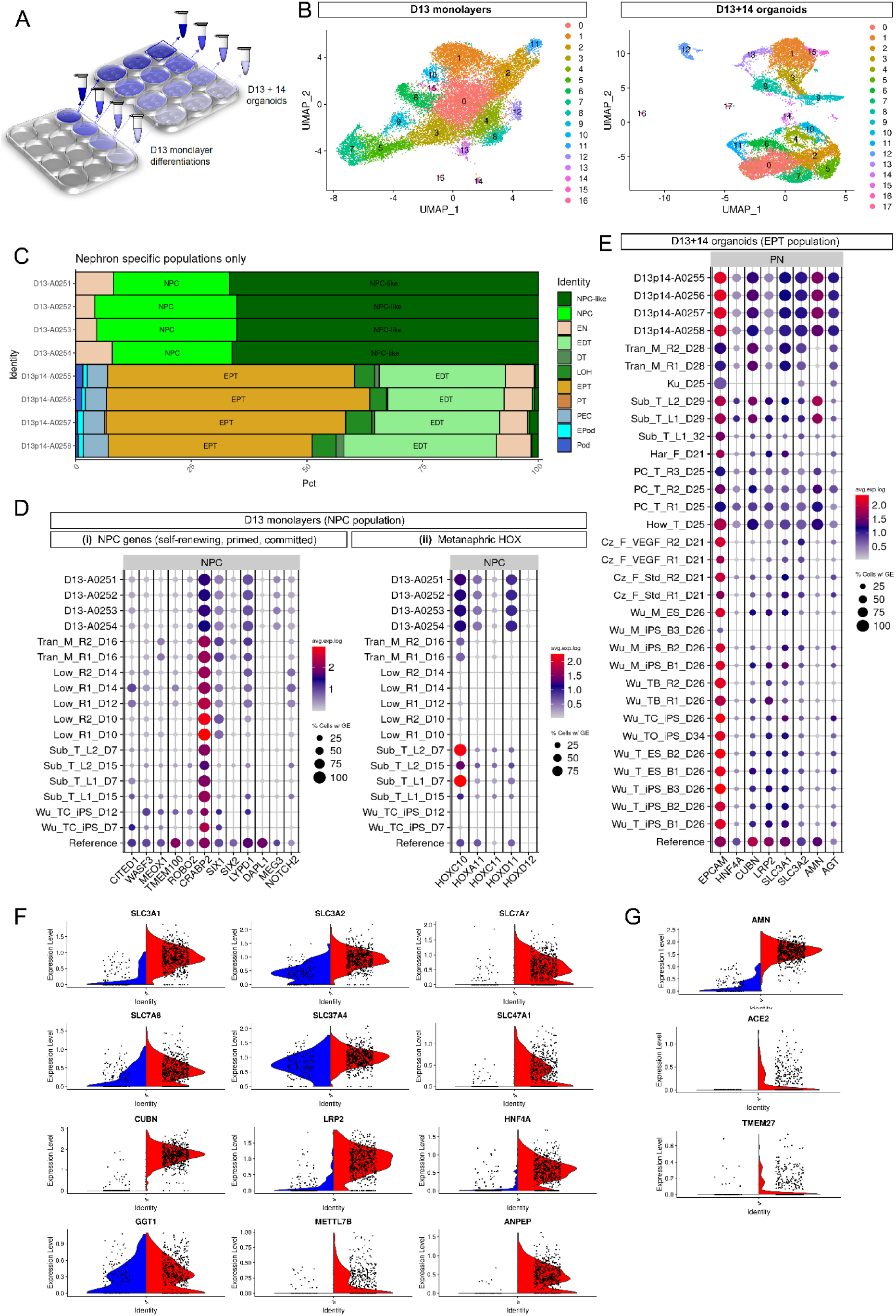
Single cell transcriptional profiling (scRNAseq) shows improved specification, patterning and maturation of proximal tubules and their progenitors. **A.** Schematic depicting experimental design and profiled samples. Multiple organoids (2.5 x 10^5^ cells per organoid) were generated from each of the 4 replicate differentiated cell monolayers at D13. The remaining portion of cells from each replicate monolayer were barcoded and pooled for generation of the D13 monolayer library. The resulting organoids were cultured for 14 days before being harvested and pooled within replicate wells, making 4 cell suspensions. These 4 suspensions were individually barcoded and pooled into a single cell suspension for generation of the D13+14 organoid library. **B.** UMAP plots of D13 and D13+14 samples (pooled replicates) identifying 16 and 17 distinct cell clusters, respectively. **C.** *ComparePlots* depicting proportions of kidney cell types (nephron-specific populations only) in D13 and D13+14 sample replicates as classified by *DevKidCC*. Population abbreviations: nephron progenitor cell (NPC), early nephron (EN), early distal tubule (EDT), DT (distal tubule), loop of Henle (LOH), early proximal tubule (EPT), proximal tubule (PT), parietal epithelial cell (PEC), early podocyte (EPod), podocyte (Pod). **D.** *DevKidCC* dot plots comparing the expression of gene signatures for (i) self-renewing (*SIX1, SIX2, CITED1, WASF3, DAPL1, MEOX1, TMEM100, ROBO2, CRABP2*), committed (*SIX1, SIX2, LYPD1*), and primed (*DAPL1, NOTCH2, MEG3*) NPC subsets, as well as (ii) metanephric HOX genes, within the D13 monolayer NPC population to that of published stem cell-derived kidney datasets and a mixed (week 11 – 18) human fetal kidney reference dataset (Hochane, *et al*., 2019; Tran, *et al*., 2019; Hollywood, *et al*., 2020). Comparisons were made to published monolayer and early nephrogenic datasets (Subramanian, *et al*., 2019; Wu, *et al*., 2018; Low, *et al*., 2019; Tran, *et al*., 2019) as outlined previously (Wilson, *et al*., 2021) . **E.** *DevKidCC* dot plot comparing the expression of proximal nephron (PN) gene signatures within the EPT population of D13+14 organoids to that of published stem cell-derived kidney organoid datasets (Czerniecki, *et al*., 2018; Harder, *et al*., 2019; Kumar, *et al*., 2019) and the mixed week 11 - 18 fetal kidney reference dataset (Hochane, *et al*., 2019; Tran, *et al*., 2019; Hollywood, *et al*., 2020) as outlined previously (Wilson, *et al*., 2021). **F-G.** Violin plots in (F) and (G) compare PT-specific gene expression of PT-enhanced organoids (red, right) with our existing standard organoid dataset of equivalent line and age (blue, left) (Howden, *et al*., 2019). Genes encoding auxiliary proteins are shown in (G).

The resulting D13 and D13+14 pooled replicate libraries resolved 19,956 and 15,852 individual cell transcriptomes per timepoint, respectively. UMAP plots showed the resolution of distinct clusters for both D13 monolayers and resulting PT-enhanced (D13+14) organoids (Figure 4B). Gene expression analyses confirmed the expression of a range of markers for mesenchymal cell states pre-kidney organogenesis in D13 monolayers, as well as markers of proximodistal patterning, stroma, and endothelium in D13+14 organoids (Supplementary Figure 2BC and Supplementary Tables 1 - 2). To enable unbiased comparisons of kidney cell types and gene expression levels between D13/D13+14 samples, published stem cell-derived, and reference reference human kidney datasets, datasets were analysed using the *DevKidCC* package (Wilson, *et al*., 2021). *DevKidCC* enables robust classification of novel developing human or stem cell-derived kidney organoid datasets without the need for integration or prior dimensional reduction or clustering. Using the *ComparePlot* function, kidney cell proportions in D13 and D13+14 samples were directly compared, confirming distinct differences in cell populations yet consistency between the 4 replicates within each sample (Figure 4C and Supplementary Figure 3A). As anticipated, over 90% of cells within the D13 monolayer differentiations were classified as nephron progenitor cells (NPC) or NPC-like, with a small contribution of cells classified as early nephron (EN) (Figure 4C). In contrast, D13+14 organoids possessed a range of proximal, distal, and renal corpuscle cell types. Early proximal tubule (EPT) formed the largest proportion of organoid nephron cell types (51% average across 4 samples), while two replicates possessed a small (<5%) fraction of maturing PT cells. By contrast, previous studies of the standard organoid protocol show on average <25% EPT and no PT (Takasato, *et al*., 2015).

*DevKidCC* was next used to compare cell type-specific markers in D13 / D13+14 samples to published stem cell-derived and reference human fetal kidney datasets (Figure 4DE). Analysis of the NPC population within D13 samples confirmed strong gene signatures for committed NPCs (*SIX1, SIX2*, and *LYPD1*) and the metanephric HOX code (*HOXC10/11, HOXA11*, and *HOXD11*) compared to relevant published monolayer and nephrogenic-stage differentiations (Subramanian, *et al*., 2019; Wu, *et al*., 2018; Low, *et al*., 2019; Tran, *et al*., 2019) that better emulated the mixed reference dataset of human fetal kidneys (weeks 11, 13, 16, 18) (Hochane, *et al*., 2019; Tran, *et al*., 2019; Holloway, *et al*., 2020). PT-enhanced organoids derived from these D13 monolayer differentiations possessed high and abundant expression of a range of proximal nephron markers in their EPT population (Figure 4E). These included genes encoding several membrane proteins critical for PT transport of proteins and amino acids (*CUBN, LRP2, SLC3A1*, and *SLC3A2*), as well as auxiliary proteins and transcription factors required for transporter regulation and functionality, such as *AMN*, *AGT*, and *HNF4A*. This gene signature showed remarkable congruence to reference human fetal kidney and improved PT identity compared to existing published kidney organoid datasets (Czerniecki, *et al*., 2018; Harder, *et al*., 2019; Kumar, *et al*., 2019) (Figure 4E).

An important anatomical feature of the mature PT is its segmentation into functionally and morphologically distinct regions defined as the S1/S2 convoluted tubule segments and the S3 straight segment. In addition to differences in proliferation characteristics and protein synthesis, the convoluted and straight segments display distinct differences in solute handling to accommodate the declining concentration of solutes as the ultrafiltrate passes through the nephron (Zhuo and Li, 2013; Avissar, *et al*., 1994). As such, early S1 – S2 convoluted segments express low-affinity/high-capacity transporters, with a gradual transition to high-affinity/low-capacity transporters in the later S3 straight segment (Palacin, *et al*., 2001; Schuh, *et al*., 2018; Verrey, *et al*., 2005). To determine whether the PTs of enhanced organoids show evidence of this segmentation, PT clusters from the 4 integrated D13+14 replicate datasets were isolated and re-clustered, resolving 4740 PT cells across 6 distinct clusters (Supplementary Figure 3B). The PT population was analysed for the expression of segment-specific PT markers with critical functional roles, including solute carriers for ions (*SLC34A1*/NPT2 (Fenollar-Ferrer, *et al*., 2015) expressed in S1>S2), glucose (*SLC2A2*/GLUT2 and *SLC5A2*/SGLT2 expressed in S1>S2; *SLC2A1*/GLUT1 and *SLC5A1*/SGLT1 expressed in S2<S3 (Hummel, *et al*., 2011; Rahmoune, *et al*., 2005; Wood and Trayhurn, 2003)), amino acids (*SLC7A9*/b(0,+)AT transporter of cystine, aspartate, and glutamate expressed in S1/S2 > S3 (Nagamori, *et al*., 2016), and cationic drugs/toxins (*SLC47A1*/MATE1 expressed in S1/S2 > S3 (Otsuka, *et al*., 2005)), as well as *AKAP12* (involved in cell cycle regulation, expressed in S2<S3 (Vogetseder, *et al*., 2008) and *GPX3* (glutathione peroxidase; secreted antioxidant synthesised in S1/S2>S3 (Avissar, *et al*., 1994)) (Supplementary Figure 3C). UMAP plots revealed the largely opposing distributions of cells expressing S1>S2 and S2>S3 gene signatures (Supplementary Figure 3C). Cells expressing S1>S2 convoluted PT markers (*SLC34A1*/MATE1, *SLC2A2*/GLUT2, and *SLC5A2*/SGLT2) were predominantly located in clusters 0, 3, and the lower portion of cluster 4, whereas cells expressing S2<S3 straight PT markers (*AKAP12*, *SLC2A1*/GLUT1, and *SLC5A1*/SGLT1) were primarily within clusters 1, 2, and the upper portion of cluster 4. When analysed for markers that exhibit a gradient of expression along the length of the nephron (S1/S2>S3), UMAP plots for each gene revealed a similar graded expression pattern, with a higher concentration of positive cells within the S1>S2 cluster (0) and decreasing in prevalence within S2<S3 clusters (0, 2) (Supplementary Figure 3C). Together this suggested that, despite the low expression of some markers indicating PT immaturity, the PTs of enhanced kidney organoids show evidence of separation into the 3 anatomically distinct PT segments.

Comparison between organoids is confounded by the inherent variability of different organoid protocols, technical variables, and individual cell line characteristics. To minimise potential bias when comparing cell maturation, PT-enhanced organoid scRNAseq data were compared to an existing standard organoid dataset derived from the same iPSC line and of equivalent organoid age (Howden, *et al*., 2019). Libraries from the PT-enhanced and standard organoid samples resolved 6737 and 1879 cells, respectively. Datasets were integrated prior to quality control measures to enable direct comparison of PT maturation and UMAP plots confirmed the resolution of distinct kidney cell clusters for both samples (Supplementary Figure 3D). Violin plots of the PT cluster alone in integrated datasets confirmed that the PT-enhanced organoid dataset possessed higher and more abundant expression of genes critical for PT functionality compared to the standard organoid (Figure 4FG). Examples included genes encoding membrane transporters CUBILIN/*CUBN* and MEGALIN/*LRP2* (important for protein uptake (Nielsen, *et al*., 2016), heavy-chain subunit solute carriers rBAT/*SLC3A1* and 4F2/*SLC3A2* (required for heteromer formation and amino acid transport by SLC7 family members (Kowalczuk, *et al*., 2008), light-chain subunit solute carriers y+LAT-1/*SLC7A7* and LAT2/*SLC7A8* (responsible for regulating intracellular amino acid pool via basolateral efflux of basic and neutral amino acids for transport systems y+L and L, respectively (Kanai, *et al*., 2000; Verrey, 2003), and solute carriers critical for PT metabolism and drug transport (G6PT1/*SLC37A4* and MATE1/*SLC47A1* (Lee, *et al*., 2015) (Figure 4F). Several auxiliary proteins essential for correct apical localisation and transporter functionality also showed higher expression in the PT-enhanced dataset, including *AMN* (Amnionless), *ACE2,* and *TMEM27* (Collectrin) (Kowalczuk, *et al*., 2008; Camargo, *et al*., 2009; Fyfe, *et al*., 2004; Ahuja, *et al*., 2008) (Figure 4G). Expression of genes encoding drug transporters *SLC22A2* (OCT2) and *SLC22A6* (OAT1) were low in both conditions but increased in PT-enhanced compared to standard organoids (Supplementary Figure 3E).

To investigate PT maturation further, an unbiased ToppFun GO Molecular Function analysis was performed on genes that were significantly differentially expressed within the PT cluster of PT-enhanced compared to standard organoids (945 input genes). This analysis revealed key differences in genes involved in cell metabolism (Supplementary Figure 3F). PT-enhanced organoid cells within the PT cluster showed increased expression of genes related to fatty acid metabolism and its regulation, such as *PPARG*, *FABP3*, *PRKAA2*, and *FAT1* (Supplementary Figure 3G). Given the known reliance of mature PT cells on fatty acid metabolism *in vivo* (reviewed in (Zhuo and Li, 2013), this gene signature was suggestive of a more mature metabolic profile in enhanced compared to standard organoid PT cells.

Together, these comprehensive scRNAseq analyses confirmed an increased abundance and relative maturation of PT within this extended protocol. Analyses of D13 monolayers suggests this higher-order PT patterning arises from improved NPC identity at the point of metanephric specification.

### Radial nephron patterning and alignment is associated with localised stroma-associated WNT antagonism

Of interest was the characteristic radial patterning observed in all PT-enhanced organoids, where tubules align with their glomeruli towards the centre of the organoid, surrounding a central core region, and distal SLC12A1+ segments towards the organoid periphery (refer to Figure 2C). This orientation was suggestive of a directional patterning cue emanating from the core region, shown earlier to express stroma marker proteins MEIS1/2/3 and SNAI2 (Supplementary Figure 1A). Previous studies have not only suggested a role of interstitial/stromal populations in nephron differentiation (England, *et al*., 2020; Das, *et al*., 2013), but have also indicated proximo-distal patterning is controlled by Wnt/β-catenin signalling along the nephron axis, with lower WNT signalling leading to improved formation and maturation of the proximal nephron (Lindstrom, *et al*., 2015). In agreement with this, WNT inhibition has been observed to promote podocyte commitment in PSC cultures (Yoshimura, *et al*., 2019). These findings suggested that the central core of PT-enhanced organoids may possess stromal populations influencing nephron patterning and/or express a localised WNT antagonist leading to directional signalling cues.

PT-enhanced scRNAseq datasets classified by *DevKidCC* were re-analysed to examine the stromal populations at greater depth. In addition to nephron-related and endothelial populations, previous classification of D13+14 organoids identified 48.2% of cells as stroma (enriched for *CRABP1*, *COL3A1*, *COL1A1*, *COL1A2* and *CXCL12*) and 23.8% of cells as unassigned but similarly enriched for collagens (e.g., *COL2A1* and *COL9A1*) (Supplementary Figure 3A). Further analyses of D13+14 populations for defined markers of stromal zones curated in mouse kidney (England, *et al*., 2020) revealed the stromal cells of PT-enhanced organoids were most like those of kidney cortex (Figure 5Ai). High expression of cortical stroma (CS) markers, including *FIBN*, *DLK1*, *MEIS1/2*, and *SNAI2*, were observed predominantly in the unassigned, stroma, and NPC-like subsets, while medullary stroma and stromal progenitor markers were largely absent (Figure 5Ai). Unassigned and stroma clusters also highly expressed the WNT antagonist Secreted Frizzled-Related Protein-2 (*SFRP2*) and developing cartilage markers (*ONG*, *MGP*, and *COL2A1*) that been previously identified in mouse kidney stromal cells (*ONG* and *MGP*) (Tanigawa, *et al*., 2022) and nephrogenic mesenchyme (*COL2A1*) (Menon, *et al*., 2018; Zhu, *et al*., 1999)(Figure 5Ai). When compared to standard organoid datasets derived from a range of relevant published protocols, D13+14 PT-enhanced organoids possessed a similar cortical stroma gene signature to several datasets, but notably higher expression of the WNT antagonist, *SFRP2*, and pre-cartilage markers, within cortical stroma and unassigned populations (Figure 5Aii).

D13 monolayers were similarly re-analysed to determine at which stage of the differentiation protocol (monolayer or 3D culture) stroma and pre-cartilage subtypes appear. Previously shown to contain just 0.9% stromal cells (Supplementary Figure 3A), analysis of the D13 sample following *DevKidCC* classification confirmed a lack of stromal progenitor (SP) and medullary stroma (MS) zone markers, while expression of cortical stroma (CS) and pre-cartilage markers were limited (Figure 5Ai). This suggested that these definitive CS and pre-cartilage populations arise during the organoid culture period, but possibly from precursor NPC-like and/or unclassified cell populations in the D13 monolayer owing to their dominance in the differentiations (83%) (Supplementary Figure 3A). Indeed, the NPC-like population in D13 monolayers showed a high similarity to the NPC population without key NPC markers (e.g. *PAX8* and *SIX2*), while Azimuth label transfer method using a human developmental reference dataset (Cao, *et al*., 2020) still classified the majority (∼75%) of D13 monolayer cells as ‘metanephric’ despite 52.3% being unclassified by *DevKidCC* (Supplementary Figure 3A, Figure 5Aiii).

The cortical stromal gene expression, notably including the WNT antagonist *SFRP2*, suggested that the central core region of PT-enhanced organoids may control WNT pathway-mediated nephron patterning, in turn driving the observed radial alignment. To functionally test this hypothesis, a WNT signalling gradient was recreated using agarose beads soaked in the tankyrase inhibitor, IWR-1 (10µM), which antagonises canonical WNT/β-catenin pathway activity (Gunaydin, *et al*., 2012)(Figure 5B). Following the 7 day (standard) differentiation protocol, iPSC-derived kidney progenitors were bioprinted and cultured to create rectangular patch organoids (Lawlor, *et al*., 2021). At 5 days of organoid culture (D7+5), by which time renal vesicle formation had occurred, IWR-1-soaked or control (PBS-soaked) beads were added to the centre of the organoids where they made contact with the early epithelial structures (Supplementary Figure 4A). After 9 days of organoid culture, organoids with IWR-1-soaked beads exhibited visible differences in the morphology of structures surrounding the beads compared to controls with PBS-soaked beads (Supplementary Figure 4B). This became more apparent when these organoids were stained via immunofluorescence (Figure 5B). In control organoids with PBS-soaked beads, beads were in contact with a mixture of proximal and distal EPCAM-positive nephron epithelium, as well as NPHS1-positive podocytes of glomeruli (Figure 5Bi). In contrast, IWR-1-soaked beads were predominantly surrounded by glomeruli, with few distal structures (LTL-negative/EPCAM-positive) visible overall (Figure 5Bii). These observations were confirmed by image quantification, showing that the percentage of NPHS1+ podocytes (glomeruli) was significantly higher in the region adjacent to IWR-1-soaked beads compared to PBS-soaked control beads (Figure 5Ci-ii and Supplementary Table 3).

**Figure 5.**
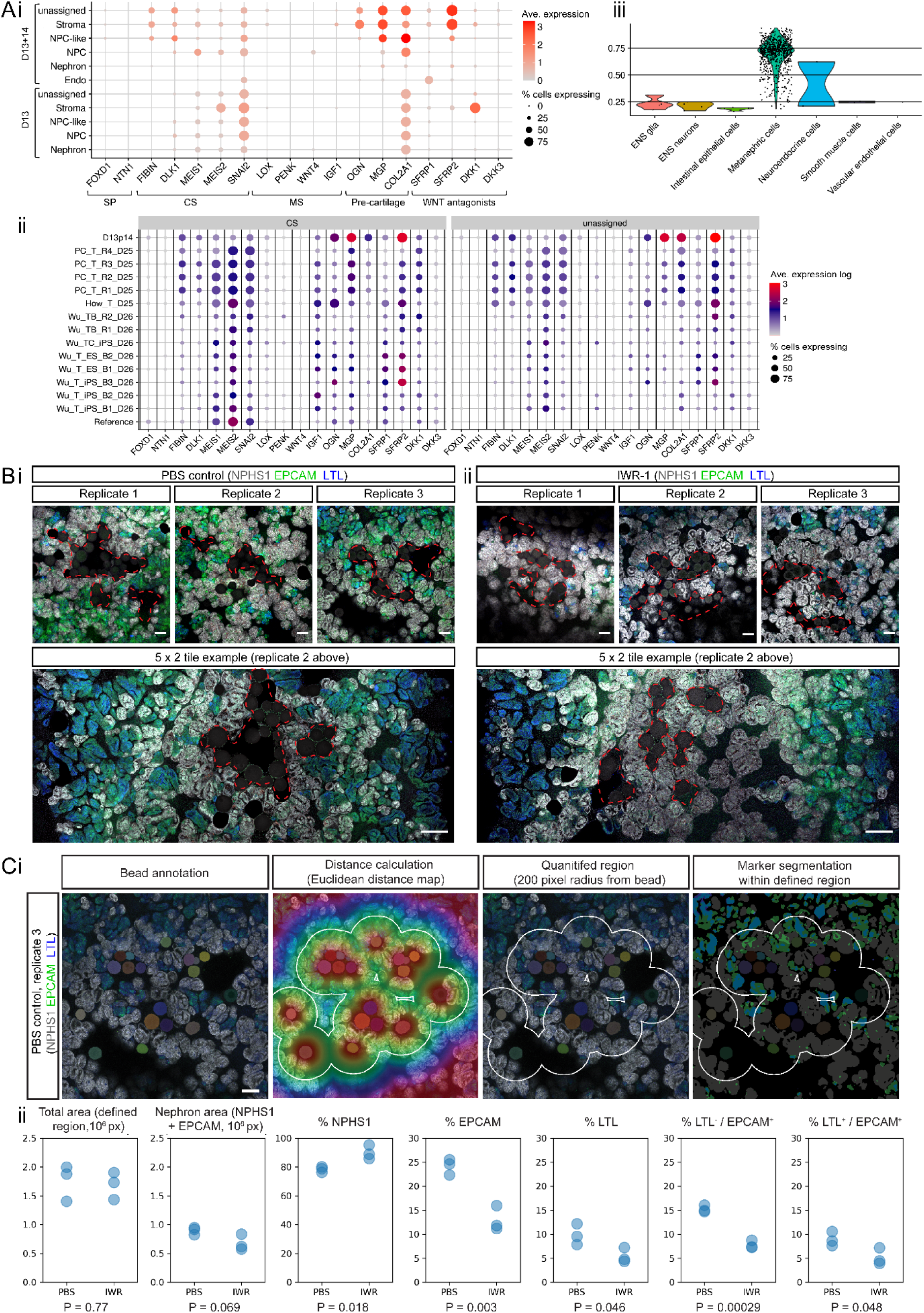
WNT signalling gradient influences nephron alignment and directionality. **A.** Analyses of scRNAseq datasets for D13+14 PT-enhanced organoid replicates (**i - iii**) and D13 monolayer replicates (**i** and **iii**) as classified by *DevKidCC*. Dot plot in (**i**) depicts expression of stroma compartment markers (England, *et al*., 2020) and WNT antagonists in D13+14 and D13 samples. *DevKidCC’s* DotPlotCompare in (**ii**) shows the comparison of D13+14 to other relevant published datasets (cortical stroma [CS] and unassigned populations only). Dot colour and size in (**i - ii**) represents unscaled gene expression and percentage of cells expressing each gene, respectively. Violin plot (**iii**) depicts the similarity scores for all unassigned cells within D13 monolayer replicates as predicted by the Azimuth label transfer method (https://azimuth.hubmapconsortium.org/) with the human developmental reference dataset (Cao, *et al*., 2020), where cells are grouped by population with the highest similarity score. Population abbreviations in (**i – iii**): nephron progenitor cell (NPC), endothelium (Endo), stroma progenitor (SP), cortical stroma (CS), medullary stroma (MS), enteric nervous system (ENS). **B.** Confocal immunofluorescence images of replicate standard organoids bioprinted in a patch conformation and in contact with either agarose beads soaked in PBS (control; **Bi**) or in 10µM IWR-1 (**Bii**). Clusters of beads are outlined with red dashed lines. Organoids are stained with markers of epithelium (EPCAM; green), proximal tubule (LTL; blue), and podocytes of the glomeruli (NPHS1; grey). Scale bars represent 100 µm (all top panels) and 200 µm (bottom tile panels). **Ci.** Example image from (**Bi**) (PBS control, replicate 3) illustrating the image annotation approach used to segment and quantify the proportion of nephron structures (NPHS1+ [grey], EPCAM+ [green], and LTL+ [blue]) within a defined region 200 pixels from any bead (white outline). Solid colours represent masks for beads and nephrons. Scale bar represents 200 µm. **Cii**. Quantification of PBS control and IWR-1 treated organoid images from (**Bi-ii**) using approach illustrated in (**Ci**), with n = 3 replicates per condition. Total area and nephron area values are shown in pixels (10^6^ px). Percentage (%) of each structure (NPHS1+, EPCAM+, LTL+) are shown as a proportion of the total nephron area. P values are indicated below each plot.

Taken together, these analyses supported establishment of a gradient arising from centralised WNT antagonism as responsible for the nephron directionality and alignment in PT-enhanced organoids.

### Mature transporter expression within PT-enhanced organoids enables nephron functionality and drug screening

The strong expression and apical cellular localisation of transporters in PT-enhanced organoids was suggestive of nephron functionality. To test this, we firstly performed multiple substrate uptake assays specific to PTs in both standard and PT-enhanced kidney organoids (Figure 6A). While standard organoids showed evidence of uptake of fluorescently labelled albumin (TRITC-albumin) into MEG-positive PTs (indicative of MEG-CUBN transport function), this uptake was visibly higher in PT-enhanced organoids, with large portions of elongated PTs displaying high-intensity TRITC-albumin fluorescence (Figure 6Ai). In addition, PTs of enhanced organoids demonstrated robust uptake of 4′,6-diamidino-2-phenylindole (DAPI), which is an effective probe for evaluation of the PT-specific SLC47 family of organic cation/H^+^ antiporters, MATE-1 (Multidrug and Toxin Extrusion Protein 1) and MATE2-K (Multidrug and Toxin Extrusion Protein 2K) (Yasujima, *et al*., 2010) (Figure 6Aii). The uptake of DAPI by PT cells was successfully inhibited via pre-treatment of organoids with the cation transporter inhibitor Cimetidine, supporting the specificity of transport activity, while the absence of DRAQ7 staining excluded the possibility of DAPI uptake in PTs due to cell death (Figure 6Aii). In contrast, standard organoids showed no uptake of DAPI, suggesting functional immaturity of these same drug transporters (Figure 6Aii).

**Figure 6:**
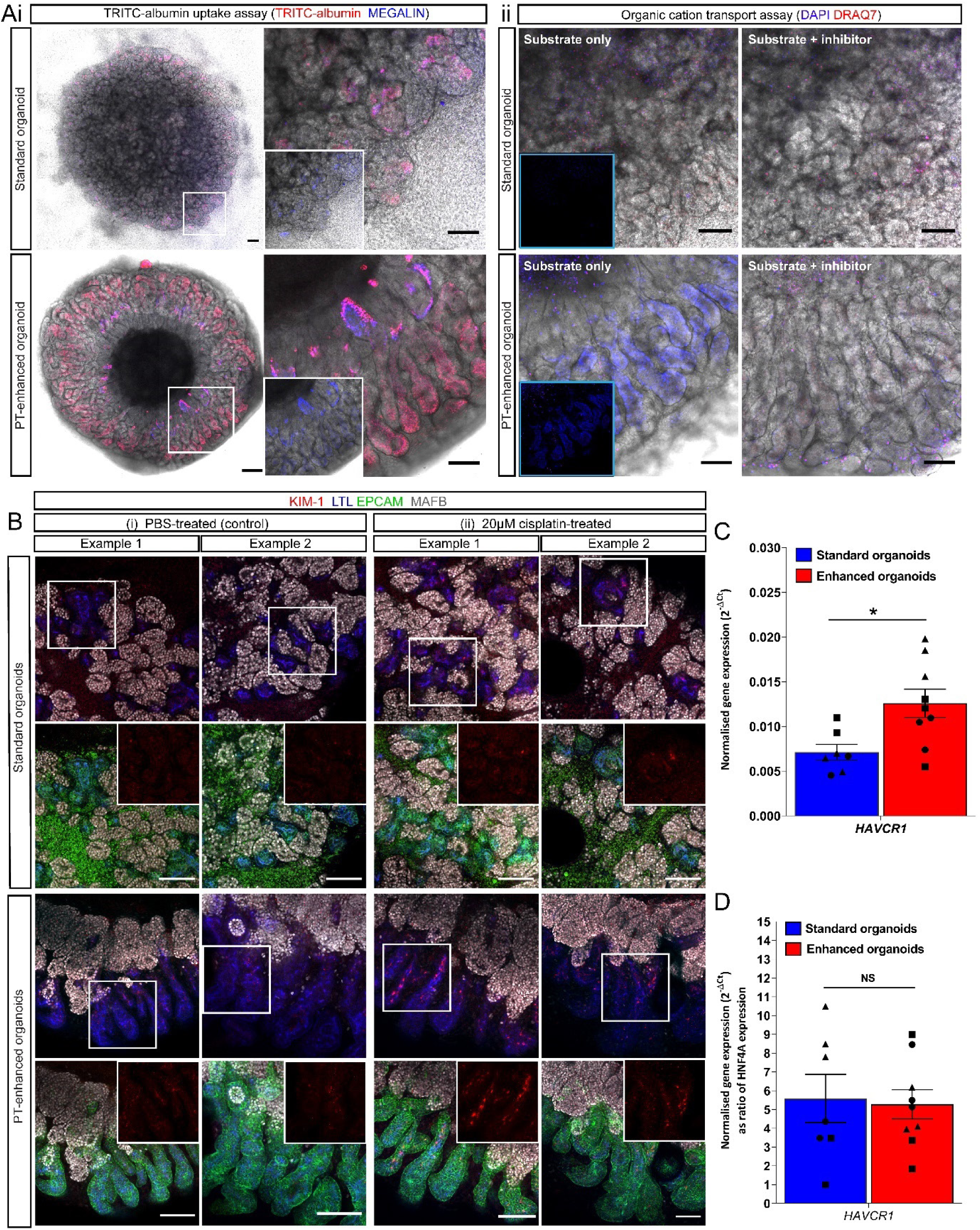
Enhanced organoids possess functional PT transporters and appropriate injury response. **A.** Live confocal images of standard and PT-enhanced organoids (transmitted light and fluorescence overlays) depicting uptake of (**i**) TRITC-albumin (red) into MEGALIN- positive PTs (blue) and (**ii**) uptake of DAPI (blue; surrogate for organic cation transport capacity). White boxed areas on left images of (**i**) are shown as higher magnification on right (insets depict MEGALIN staining alone). Organic cation uptake image set (**ii**) depicts organoids exposed to substrate alone (DAPI; blue, left images) or a combination of substrate/DAPI + inhibitor/Cimetidine (right images). Dead cells in both panels of (**ii**) are labelled with DRAQ7 (red). Insets in (**ii**) depict blue channel only without brightfield overlay. Scale bars represent 200 µm (whole organoid images in [**i**]) and 100 µm (left images in [**i**], all images in [**ii**]). **B.** Confocal immunofluorescence of representative D7+14 (standard; top panels) and D13+14 (PT-enhanced; bottom panels) line-matched organoids following 24 hours treatment with E6 media containing either (**ii**) 20 µM cisplatin or (**i**) an equivalent volume of PBS. Images depict KIM-1-expressing cells (red) in LTL+ proximal tubules (blue) with nephron epithelium co-stained with EPCAM (green). Insets of bottom row images for standard and PT-enhanced organoids show KIM-1 staining (red channel) alone from white boxed regions in top row images. Scale bars in all images represent 100 µm. **C-D.** qRT-PCR analyses depicting KIM-1 gene (*HAVCR1*) expression in standard (blue) and PT-enhanced (red) organoids from experiments shown in (**B**). *HAVCR1* gene expression values are normalised to the expression of housekeeping gene *GAPDH* and depicted both with (**D**) and without (**C**) compensation for differences in proximal tubule proportion (expressed as a ratio of *HNF4A*). Error bars represent SEM from n = 8 (control) and n = 9 (cisplatin-treated) biological replicates across 3 replicate experiments as indicated. Statistical significance was determined using an unpaired t test. Asterisk represents P value (*; P ≤ 0.05) adjusted for multiple comparisons using the Holm-Sidak method alpha = 0.05. NS = non-significant.

Having established albumin and organic cation transport capacity in PT-enhanced organoids, we next assessed their response to nephrotoxic insult (Figure 6BCD). Several recent studies have explored the suitability of kidney organoids as a human-relevant model of cisplatin-induced nephrotoxicity (Freedman, *et al*., 2015; Morizane, *et al*., 2015; Takasato, *et al*., 2015), a common complication that limits usage of this chemotherapeutic agent (Ozkok and Edelstein, 2014; Yao, *et al*., 2007). The biomarker KIM-1 is sensitive for early detection of PT injury in humans and animals (Abdelsalam, *et al*., 2018; Chiusolo, *et al*., 2010; Sasaki, *et al*., 2011; Shinke, *et al*., 2015; Vaidya, *et al*., 2010) and has been shown to increase in response to cisplatin in kidney organoids, despite conflicting reports regarding its PT-specificity (Morizane, *et al*., 2015; Takasato, *et al*., 2016; Digby, *et al*., 2020). This discrepancy may arise from immature expression of the predominant cisplatin transporters, particularly SLC22A2/OCT2 (Digby, *et al*., 2020), combined with heterogeneity in cisplatin uptake mechanisms. Re-analysis of our PT-enhanced and existing standard organoid scRNAseq datasets (Howden, *et al*., 2019) revealed higher expression of the majority of cisplatin influx and efflux transporters in enhanced compared to standard organoid PT cells (Supplementary Figure 4C), suggestive of cisplatin transport capacity. This included *SLC22A2*/OCT2, previously reported to show low expression in kidney organoids (Digby, *et al*., 2020). To confirm the functionality of these transporters and appropriate injury response by PTs, iPSC line-matched D7+14 (standard) and D13+14 (enhanced) organoids were derived from monolayer differentiations across 3 independent experiments. Organoids were exposed to 20 µM cisplatin for 24 hours prior to assessment for expression of KIM1 protein its corresponding gene, *HAVCR1*. Immunofluorescence revealed an upregulation of KIM-1 protein expression within LTL-positive PTs of both standard and enhanced organoids compared to PBS-treated controls (Figure 6Bi-ii). This was supported by a significant increase in KIM-1 gene (*HAVCR1*) expression in PT-enhanced organoids compared to standard organoids (Figure 6C). Also noteworthy was the similar *HAVCR1* expression levels in standard and PT-enhanced organoids when gene level was expressed relative to the absolute amount of PT in each organoid (marked by *HNF4A*). This suggested that the levels of *HAVCR1* upregulation may be dictated by proximal tubule proportion (Figure 6D). However, in both standard and PT-enhanced organoids, *HAVCR1* expression was significantly increased compared to control organoids (Supplementary Figure 4D).

### PT-enhanced organoids represent an improved model for SARS-CoV-2 pathogenesis research

Kidney organoids have previously proven useful to model inherited, early-onset kidney disease (Freedman, *et al*., 2015; Taguchi and Nishinakamura, 2017; Czerniecki, *et al*., 2018; Cruz, *et al*., 2017; Forbes, *et al*., 2018; Hale, *et al*., 2018; Hollywood, *et al*., 2020; Mae, *et al*., 2013; Przepiorski, *et al*., 2018; Tanigawa, *et al*., 2018). More recently, organoids have been successfully applied to understanding the pathogenesis of the infectious respiratory disease COVID-19, with SARS-CoV-2 viral infection and replication being achieved in a range of stem cell-derived tissues (Han, *et al*., 2020; Marchiano, *et al*., 2021; Mills, *et al*., 2021; Sharma, *et al*., 2020; Tiwari, *et al*., 2021). Driven by the occurrence of AKI in COVID-19 patients (Huang, *et al*., 2020; Kunutsor and Laukkanen, 2020; Yang, *et al*., 2020; Zhou, *et al*., 2020), a handful of studies have explored kidney organoids as a potential model of COVID-19 (Monteil, *et al*., 2020; Wysocki, *et al*., 2021). While it is still debated whether kidney damage results from direct viral infection or a combination of inflammatory responses and drug nephrotoxicity (reviewed in (Motavalli, *et al*., 2021), human PTs show high expression of the key SARS-CoV-2 receptor ACE2 (Kowalczuk, *et al*., 2008; Hoffmann, *et al*., 2020) and evidence of viral infection (Braun, *et al*., 2020; Farkash, *et al*., 2020; Kissling, *et al*., 2020; Puelles, *et al*., 2020; Su, *et al*., 2020; Werion, *et al*., 2020; Hanley, *et al*., 2020).

Given the high proportion of PT in enhanced organoids, we investigated their suitability as a model of SARS-CoV-2 infection and pathogenesis. Comprehensive analysis of scRNAseq data from >15,800 D13+14 organoid cells revealed expression levels and cellular localisation of a range of entry factors (receptors, proteases and binding proteins) previously implicated in SARS-CoV-2 infection (Amraei, *et al*., 2021; Singh, *et al*., 2020) (Supplementary Figure 5A). When comparing age- and line-matched organoids, all SARS-CoV-2 entry factors of the proximal and distal tubular segments showed increased expression levels and abundance in PT- enhanced organoids compared to our existing standard organoid dataset (Figure 7A). The two most frequently reported viral entry factors in literature, *ACE2*/ACE2 and *TMPRSS2*/TMPRSS2 (Hoffmann, *et al*., 2020), were confirmed to be expressed at both a gene- and protein-level in proximal and distal nephron compartments, respectively (Figure 7AB), supporting previous reports *in vivo* and in kidney organoids (Kowalczuk, *et al*., 2008; Camargo, *et al*., 2009; Han, *et al*., 2020; Monteil, *et al*., 2020; Wysocki, *et al*., 2021).

**Figure 7:**
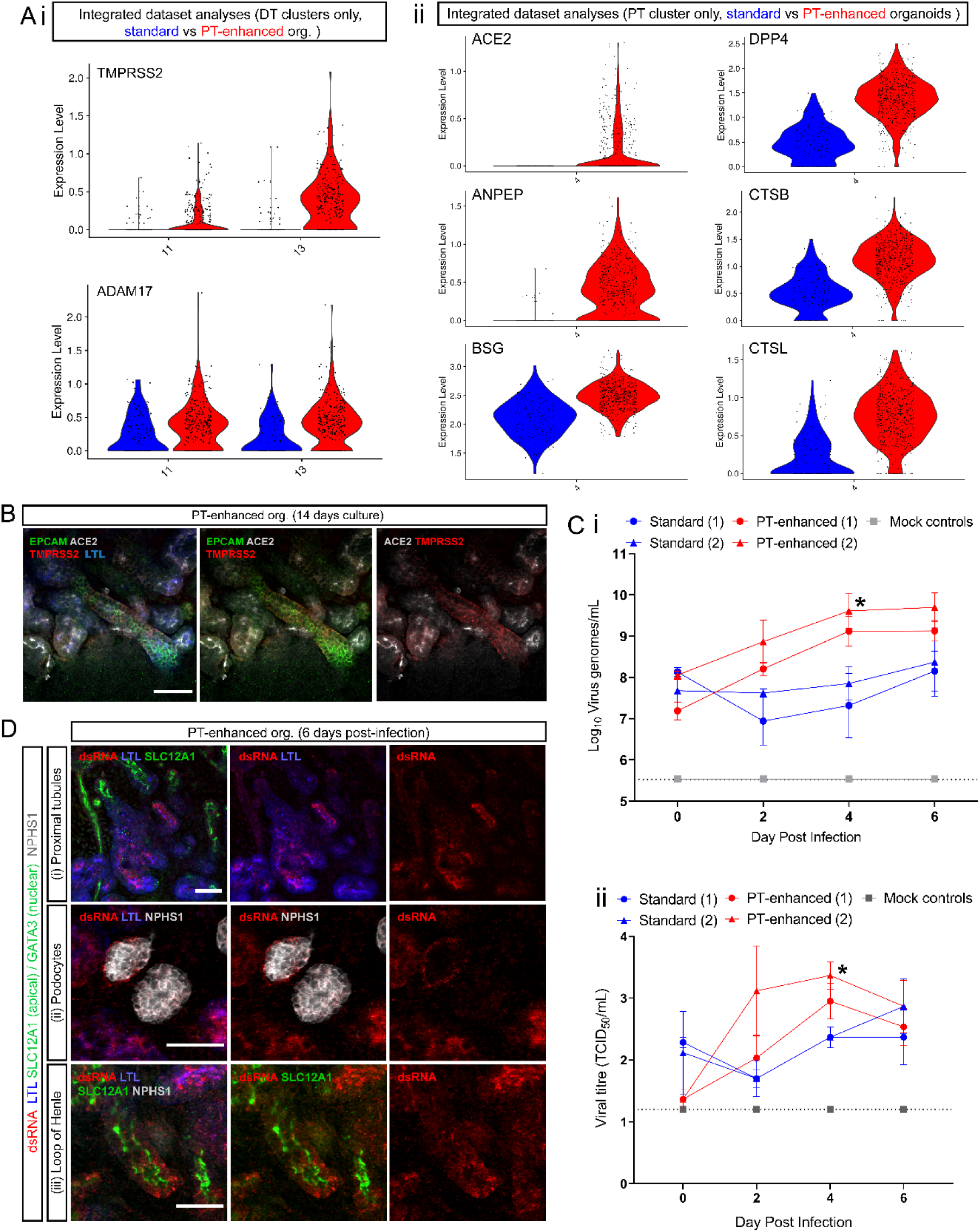
PT-enhanced organoids show improved SARS-CoV-2 entry factor expression, infectivity, and viral replication. **A.** scRNAseq analyses comparing the expression of SARS-CoV-2 entry factors in PT-enhanced organoids (red) and our existing standard organoid dataset (line- and age-matched) (Howden, *et al*., 2019). Violin plots compare expression of genes within integrated datasets from which (**i**) distal tubule (DT) and (**ii**) proximal tubule (PT) clusters have been isolated. **B.** Confocal immunofluorescence of ACE2 (green) and TMPRSS2 (red) demonstrating protein localisation in PT-enhanced kidney organoids. Nephron epithelium is stained with EPCAM (green). ACE2 and TMPRSS2 entry factors are depicted in grey and red, respectively. Scale bars represent 50µm. **Ci.** qRT-PCR for SARS-CoV-2 viral envelope (E) gene (genome copies per mL) in the same culture media samples as depicted in (**Cii**) below. Organoids from the two representative experiments are indicated (1, 2) for standard (blue), PT- enhanced (red), and mock-infected (grey line; representative of all mock results across the same 4 independent experiments). Error bars represent SEM from n = 3 individual wells of organoids (3 organoids per well). Statistical significance was determined using a one-way ANOVA with Tukey’s multiple comparisons test. Asterisk represents P value (*; P ≤ 0.05). **Cii.** Viral titre determined by Vero cell assays (Median Tissue Culture Infectious Dose; TCID_50_) of culture media sampled from SARS-CoV-2 infected standard and PT-enhanced organoids (blue and red lines, respectively), as well as mock-infected organoids (grey line; representative of all mock results across the same 4 independent experiments). Dotted line represents lower limit of detection (LOD). For both standard and PT-enhanced conditions, 2 representative independent experiments, replicated using the same iPSC line and culture conditions, are indicated as (1) and (2), with error bars representing SEM from n = 3 individual wells of organoids (3 organoids per well). Statistical significance was determined using a one-way ANOVA with Tukey’s multiple comparisons test. Asterisk represents P value (*; P ≤ 0.05). **D.** Confocal immunofluorescence of PT-enhanced organoids 6 days post-infection indicating viral dsRNA (red) localisation, co-stained for PTs (LTL; blue), Loop of Henle (SLC12A1; apical green), and podocytes (NPHS1; grey). Scale bars represent 50µm.

Apical ACE2 expression was also identified in epithelial cells lining the initial portion of Bowman’s capsule transitioning from the S1 segment of the PT (Supplementary Figure 5C). Previous studies in mice have identified these transitionary cells as cuboidal and intermediate parietal epithelial cells (cuPECs and iPECs), making up the most proximal part of the PT prior to transitioning to flat PECs that line Bowmans’s capsule (Kuppe, *et al*., 2019; Wang, 2019). Accordingly, high *ACE2* gene expression correlated with a subset of cells co-expressing general PEC markers with a cuPEC/iPEC-specific profile (*PAX8+, AKAP12+, PROM1-*) (Supplementary Figure 5D). This region also partly coincided with the *SLC34A1^Hi^/HNF4A^+^/SLC36A2^+^*population marking early (S1) PT cells (Lee, *et al*., 2015; Broer, *et al*., 2008) (Supplementary Figure 5E), which, along with LTL-positivity of the early Bowmans capsule epithelium (Supplementary Figure 5C), agreed with the known S1-PEC transitionary phenotype reported for cPECs and iPECs (Kuppe, *et al*., 2019). However, *ACE2* was absent from podocytes (cluster 12; Supplementary Figure 5ACD). These expression patterns were further supported by analyses of human fetal kidney, with expression of SARS-CoV-2 entry factors exhibiting a highly similar expression pattern to our extended kidney organoids, including low levels of ACE2 in human fetal kidney PECs (Supplementary Figure 6AB).

Having confirmed the expression of viral entry factors, PT-enhanced and standard organoids were assessed for infectivity following incubation with SARS-CoV-2. Viral infection of kidney organoids was confirmed by visualisation of GFP-expressing SARS-CoV-2 reporter virus (marking replicating virus) (Hou, *et al*., 2020) in combination with immunofluorescence staining for the spike protein (S; the transmembrane protein responsible for host cell binding and viral entry) (Supplementary Figure 5D). To confirm the presence of viral genome, culture media from standard and PT-enhanced organoids were harvested every second day post-infection for qRT-PCR of SARS-CoV-2 viral envelope gene expression (*E*; genome copies per mL) (Figure 7Ci) and virus titration in Vero cells to calculate median Tissue Culture Infectious Dose (TCID_50_) (Figure 7Cii). Infectious virus was detected earlier in PT-enhanced compared to standard organoids (at 2 days post-infection) across independent experiments replicated using the same iPSC line and organoid conditions. In both instances, infectious virus levels reached significance at 4 days post-infection (P = 0.0297 and P = 0.0457, respectively) (Figure 7Ci-ii).

To determine the kidney cell types targeted by SARS-CoV-2 in PT-enhanced organoids, infected organoids were analysed via immunofluorescence for double stranded RNA (dsRNA) and nephron-specific markers 6 days post-infection (Figure 7D). In agreement with scRNAseq analyses of ACE2 receptor expression (Supplementary Figures 5ABC and 6B), infected organoids showed dsRNA predominantly in LTL-positive PTs, as well as Bowman’s capsule surrounding NPHS1-positive podocytes (undetectable in podocytes themselves) and some detection in SLC12A1-positive Loops of Henle (Figure 7Di-iii). The specificity of this staining was confirmed by immunofluorescence of uninfected control organoids, which showed no staining for dsRNA (Supplementary Figure 6D). Despite their infection, tubular epithelium in organoids exposed to SARS-CoV-2 retained key characteristics such as apically-restricted LTL and SLC12A1, as well as membrane-bound EPCAM staining (Figure 7Di and iii, Supplementary Figure 6Ei). However, upregulation of KIM-1 was observed in infected organoids and found to be significantly higher than mock (uninfected) control organoids at a gene level, complementing results of previous publications (Supplementary Figure 6Ei-ii) (Chen, *et al*., 2021; Jansen, *et al*., 2022).

## Discussion

The utility of human PSC-derived kidney organoids as accurate models for disease research applications will rely upon their nephron maturation and functionality. To date, proximal tubules characterised within kidney organoids have lacked significant evidence of functional solute transport. In this study, we have shown that prolonged maintenance and delayed epithelialisation of the nephron progenitor population improved PT maturation and functionality compared to standard organoid protocols. Critically, this approach promoted development of distinct S1, S2, and S3 cell populations within the PT, a feature not previously identified in a kidney organoid. The application of *DevKidCC* in the current study enabled an unbiased and quantitative transcriptional comparison to previous published kidney organoid and human fetal kidney datasets, providing a reliable readout of cell identity and maturation and minimising the caveats associated with comparing restricted marker panels (Wilson, *et al*., 2021).

Treatment strategies for coronavirus infections, including SARS-CoV and MERS-CoV, are still in their infancy with progress reliant upon an improved understanding of virus biology and interaction with host factors (V’Kovski, *et al*., 2021). Despite the rapid accumulation of information on SARS-CoV-2, findings have often been conflicting or challenging to interpret, including reported heterogeneity in the expression of viral entry factors and the correlation between expression levels and disease outcome (Zlacka, *et al*., 2021; Jackson, *et al*., 2022; Muus, *et al*., 2021). PT-enhanced organoids exhibited a robust response to the nephrotoxic chemotherapeutic cisplatin and superior infectivity with SARS-CoV-2 compared to standard organoids. This enhanced patterning and functionality underscores the advantage of PT- enriched organoids for drug screening and disease modelling applications, including as a model of infectious disease in the kidney.

PT-enhanced organoids exhibited improved expression of a range of previously identified viral entry factors compared to standard organoids, including the key SARS-CoV-2 receptor (ACE2) on the apical membrane of PT cells. This translated to higher virus replication levels in PT- enhanced organoids, determined by both dsRNA quantification and infectious viral genome copies across multiple timepoints, replicates, and independent experiments. Previous kidney organoid studies have reported podocyte SARS-CoV-2 infection using stem cell-derived kidney models (Jansen, *et al*., 2022; Kalejaiye, *et al*., 2022). In contrast, we saw limited viral entry factor expression and no evidence of *ACE2*/ACE2 within podocytes of PT-enhanced organoids and human fetal kidney. It is possible that reports of podocyte infection reflected viral entry in more immature podocytes or parietal cells, given the reported variation in genuine podocyte gene expression arising from the use of different cellular models/formats (Hale, *et al*., 2018; Kalejaiye, *et al*., 2022). In addition, while previous transcriptional profiling of infected organoids claimed the presence of virus within most cell populations (Jansen, *et al*., 2022), no viral entry factor expression was observed in any cell cluster within that study. Here again we conclude that PT-enhanced organoids represent a more accurate model of the mature nephron.

It remains to be seen whether the enhanced PT development in these organoids results from improved nephron progenitor expansion or sufficient time to form a more metanephric nephron progenitor population. Transcriptional profiling of day 13 monolayers exposed to CDBLY2 showed a high proportion of nephron progenitors with a significant increase in nephron progenitor gene expression (*SIX1, LYPD1*) and metanephric HOX ortholog expression (*HOX11A/C/D*) in comparison to other relevant published scRNAseq datasets. One unique feature critical to the overall outcome of this modified protocol included the addition of nephron progenitor maintenance media that prolongs low-level canonical WNT signalling (CHIR), suppresses NOTCH signalling (DAPT), and increases BMP7 activity (BMP7) (Tanigawa, *et al*., 2016). Inclusion of these factors agreed with mouse studies which have shown a requirement for Notch to initiate nephron progenitor commitment and nephron formation, as well as demonstration that Notch2 supports proximal nephron patterning (Chung, *et al*., 2017; Surendran, *et al*., 2010). In addition, low levels of canonical Wnt activity and Bmp/BMP signalling via MAPK and PI3K pathways have been proposed to support nephron progenitor survival (Brown, *et al*., 2015; Karner, *et al*., 2011; Park, *et al*., 2007; Blank, *et al*., 2009; Lindstrom, *et al*., 2015; Muthukrishnan, *et al*., 2015). Despite containing both low CHIR and BMP7, the alternate nephron progenitor maintenance media NPSR was unable to support subsequent nephron formation in the resulting organoids, possibly due to the inclusion of BMP and TGFβ receptor inhibitors (dual inhibition of SMAD1/5/8 and SMAD2/3) (Li, *et al*., 2016), which may maintain a less competent nephron progenitor population (Tanigawa, *et al*., 2019).

The influence of timing on protocol outcome also cannot be discounted. Recent studies of the relative timing of PSC differentiation suggest that development and maturation *in vitro* is influenced by a predetermined species-specific biological clock. This has been elegantly demonstrated by Matsuda *et al* (2020), showing that the markedly different paces of differentiation exhibited by mouse and human PSCs can be attributed to biochemical rate variations that influence the segmentation clock (Matsuda, *et al*., 2020). Indeed, brain organoids require months in culture to develop specific neural subtypes, akin to human gestation (Lancaster, *et al*., 2013; Velasco, *et al*., 2019). While our PT-enhanced kidney organoid protocol already shows considerable improvements in maturation after only 3 – 4 weeks, there is likely room for additional improvements including the timing of growth factor exposure and optimisation of metabolic conditions beyond the monolayer differentiation phase.

Despite enhancing PT development, this protocol faces some limitations with respect to nephron patterning and off-target populations. While providing a powerful model of PT function, reduced patterning to distal tubular segments highlights the challenge of simultaneously generating all kidney cell types in a single protocol, as previously described in mouse ((Freedman, *et al*., 2015; Morizane, *et al*., 2015; Toyohara, *et al*., 2015; Taguchi, *et al*., 2014). In addition, the formation of pre-cartilage cells is problematic for any potential clinical application, albeit not unique to this approach. Cartilage development has been observed in organoids from several protocols following transplantation (Bantounas, *et al*., 2020; Nam, *et al*., 2019; van den Berg, *et al*., 2018). In PT-enhanced organoids, this may represent a side-effect of prolonged BMP signalling that could potentially be supressed through timed SMAD1/5/8 inhibition. The presence of central pre-cartilage within the cortical stroma population of the organoid core resulted in strong central WNT antagonism (*SFRP2*) that contributed to the striking nephron alignment observed. The establishment of a sink and source of WNT activity along the length of the tubule, driving nephron directionality, is in agreement with our current understanding of proximodistal patterning during mouse development (Lindstrom, *et al*., 2015), while the cortical stroma population likely supports and promotes the proximal nephron development (Das, *et al*., 2013). Interestingly, while standard organoids develop regions of cartilage post transplantation, they do not display this characteristic nephron spatial arrangement either before or after transplant. It is possible that this core is the result of altered biophysical parameters. We have previously shown that higher density standard organoids favour the development of a central unpatterned core, whereas a bioprinted sheet does not (Lawlor, *et al*., 2021). Such observations indicate that an interplay between cell deposition density and the patterning of the mesodermal population in the enhanced protocol facilitated the strong centralised source of WNT antagonism. Together this suggests an approach to further control the spatial organisation of bioengineered tissue through manipulation of signalling gradients.

In conclusion, we describe here a protocol that enabled improved patterning and maturation of proximal tubules within kidney organoids. These show significant advantages for modelling an appropriate damage response following drug-induced injury and SARS-CoV-2 infection, underscoring the utility of this approach as a platform to model a range of proximal tubular disease states.

## Methods

### iPSC lines and maintenance

iPSC lines used in this study include CRL1502.C32 (Takasato, *et al*., 2015; Briggs, *et al*., 2013) CRL-2429/SIX2^Cre/Cre^:GAPDH^dual^ (Howden, *et al*., 2019), PCS-201-010/HNF4A^YFP^ (Vanslambrouck, *et al*., 2019), and PB010/MCRIi010-A (Vlahos, *et al*., 2019). All iPSC lines were maintained and expanded at 37°C, 5% CO_2_ and 5% O_2_ in Essential 8 medium (Thermo Fisher Scientific, Waltham, MA) on Matrigel- (BioStrategy, Victoria, Australia) coated plates with daily media changes and passaged every 2 – 3 days with EDTA in 1X PBS as described previously (Chen, *et al*., 2011).

### Directed differentiation and kidney organoid generation

For standard organoid production, differentiation of iPSC lines and organoid culture was performed as described previously (Howden, *et al*., 2019), with minor variations in the concentration of Laminin-521 (BioLamina, Sundbyberg, Sweden) used to coat 12-well plates, initial iPSC seeding density within 12-well plates, and CHIR99021 (R&D Systems) concentration and duration of exposure according to the iPSC line used (CRL1502.C32, CRL-2429/SIX2^Cre/Cre^:GAPDH^dual^ and PB010/MCRIi010-A were seeded at 25,000 cells/well and exposed to 6µM CHIR for 5 days; PCS-201-010/HNF4A^YFP^ was seeded at 40,000 cells/well and exposed to 6µM CHIR for 4 days; CRL1502.C32, CRL-2429/SIX2^Cre/Cre^:GAPDH^dual^ were seeded with 20µL/mL Laminin-521; PB010/MCRIi010-A and PCS-201-010/HNF4A^YFP^ were seeded with 40µL/mL Laminin-521). Standard bioprinted patch organoids were generated as described previously (Lawlor, *et al*., 2021).

For PT-enhanced organoids, Matrigel concentrations and iPSC seeding density for differentiation in 12-well plates were as stated for standard organoids above. iPSCs were then subjected to prolonged monolayer differentiation in 6µM CHIR for 5 days, followed by 200ng/mL FGF9 (R&D Systems) and 1µg/mL heparin (Sigma Aldrich) until day 8, refreshing the media every second day. At day 8, the monolayer was exposed to 1mL/well nephron progenitor maintenance media, NPSR or CDBLY (Li, *et al*., 2016; Tanigawa, *et al*., 2016), refreshing these media daily. Final PT-enhanced organoid conditions utilised CDBLY2, containing 2X concentration of BMP7. Organoids were generated and cultured as described previously (Takasato, *et al*., 2016).

### Immunofluorescence and confocal microscopy

For immunofluorescence, organoids were fixed and stained as previously described (Vanslambrouck, *et al*., 2019) using the antibodies detailed in Table 1, diluted in 0.1% TX-100/PBS. Imaging was performed on the ZEISS LSM 780 confocal microscope (Carl Zeiss, Oberkochen, Germany) with acquisition and processing performed using ZEISS ZEN Black software (Zeiss Microscopy, Thornwood, NY) and Fiji ImageJ (Schindelin, *et al*., 2012).

**Table 1.** Antibodies used in immunofluorescence studies.

| Specificity | Host species | Dilution range | Manufacturer and identifier |
| --- | --- | --- | --- |
| ACE2 | Rabbit | 1:300 | Abcam (ab15348) |
| CUBILIN | Goat | 1:300 | Santa Cruz Biotechnology (sc-20607) |
| dsRNA | Mouse | 1:300 | Absolute Antibody (Ab01299-2.0) |
| ECADHERIN | Mouse | 1:300 | BD Biosciences (610181) |
| EpCAM (Alexa488 or Alexa647 conjugate) | Mouse | 1:300 | BioLegend (324210 and 324212) |
| GATA3 | Goat or rabbit | 1:300 | R&D Systems (AF2605) and Cell Signalling Technology (95852S) |
| GFP | Chicken | 1:200 - 1:300 | Sapphire Bioscience (ab13970) |
| HNF4A | Mouse | 1:300 | Life Technologies (MA1-199) |
| KIM-1 | Goat | 1:300 | R&D Systems (AF1750) |
| mCherry (RFP) | Rabbit | 1:300 – 1:400 | MBL Medical & Biological Laboratories Co. Ltd. (PM005) |
| MEGALIN | Rabbit | 1:300 | Sapphire Bioscience (NBP2-39033) |
| NEPHRIN | Sheep | 1:300 | R&D Systems (AF4269) |
| Proximal tubule brush border membrane | *Lotus tetragonobulus* lectin (LTL) | 1:300 – 1:500 | Vector Laboratories (B-1325) |
| TMPRSS2 | Mouse | 1:300 | Merck (MABF2158-25UG) |
| S2 subunit of SARS-CoV-2 spike protein (stain: Sin2774) | Rabbit | 1:300 | GeneTex/Sapphire Bioscience (GTX632604) |
| SLC6A19 | Chicken | 1:100 – 1:200 | Aves Laboratories (custom antibody) |
| SLC12A1 | Rabbit | 1:300 – 1:400 | Proteintech (18970-1-AP) |

### Flow cytometry

Flow cytometry of reporter line-derived organoids using endogenous fluorescence was performed and analysed as described previously (Vanslambrouck, *et al*., 2019). To determine the contribution of SIX2-mCherry + cells to EPCAM+ populations in organoids derived from the SIX2^Cre^ lineage tracing iPSC line, dissociated and strained cells were stained using directly conjugated anti-EPCAM Alexa Fluor-647 antibody (see Table 1) diluted 1:100 in 100 µL of FACS wash (1% fetal calf serum [FCS] in PBS) for every 5 x10^5^ cells. Following 30 minutes incubation on ice, cells were washed 3 times in 2mL FACS wash via centrifugation prior to flow cytometry.

### Histology

For Alcian Blue detection of cartilage, organoids were fixed in 4% PFA as described above and processed for routine paraffin embedding using the Excelsior AS Tissue Processor (rapid biopsy setting; Thermo Fisher Scientific). Samples were embedded in wax and 5µm sections cut using a Waterfall HM325 microtome (Thermo Fisher Scientific). Sections were dewaxed, hydrated through graded alcohols to running water, then covered with Alcian Blue Solution (1% Alcian blue in 3% acetic acid, pH 2.5). After 10 minutes, sections were washed in tap water for 2 minutes and counterstained for 7 minutes in Nuclear Fast Red stain (0.1% Nuclear Fast Red [Sigma Aldrich, St Louise, MO] and 5% ammonium potassium sulfate in water). Following staining, sections were dehydrated in graded alcohols, cleared in Safsolvent (Bacto Laboratories, NSW, Australia), and coverslipped. Images were acquired on a Zeiss Axio Imager A2 with Zeiss Zen software (Zeiss Microscopy, Thornwood, NY).

### Real-time quantitative reverse transcription PCR (qRT-PCR)

RNA extraction, cDNA synthesis and quantitative RT-PCR (qRT-PCR) were performed using the Bioline Isolate II Mini/Micro RNA Extraction Kit, SensiFAST cDNA Synthesis Kit and the SensiFAST SYBR Lo-ROX Kit (Bioline, NSW, Australia), respectively, as per manufacturer’s instructions. Each qRT-PCR reaction was performed in triplicate using the primer pairs detailed in Table 2. Data were graphed and analysed in Prism 8 (GraphPad).

**Table 2.**
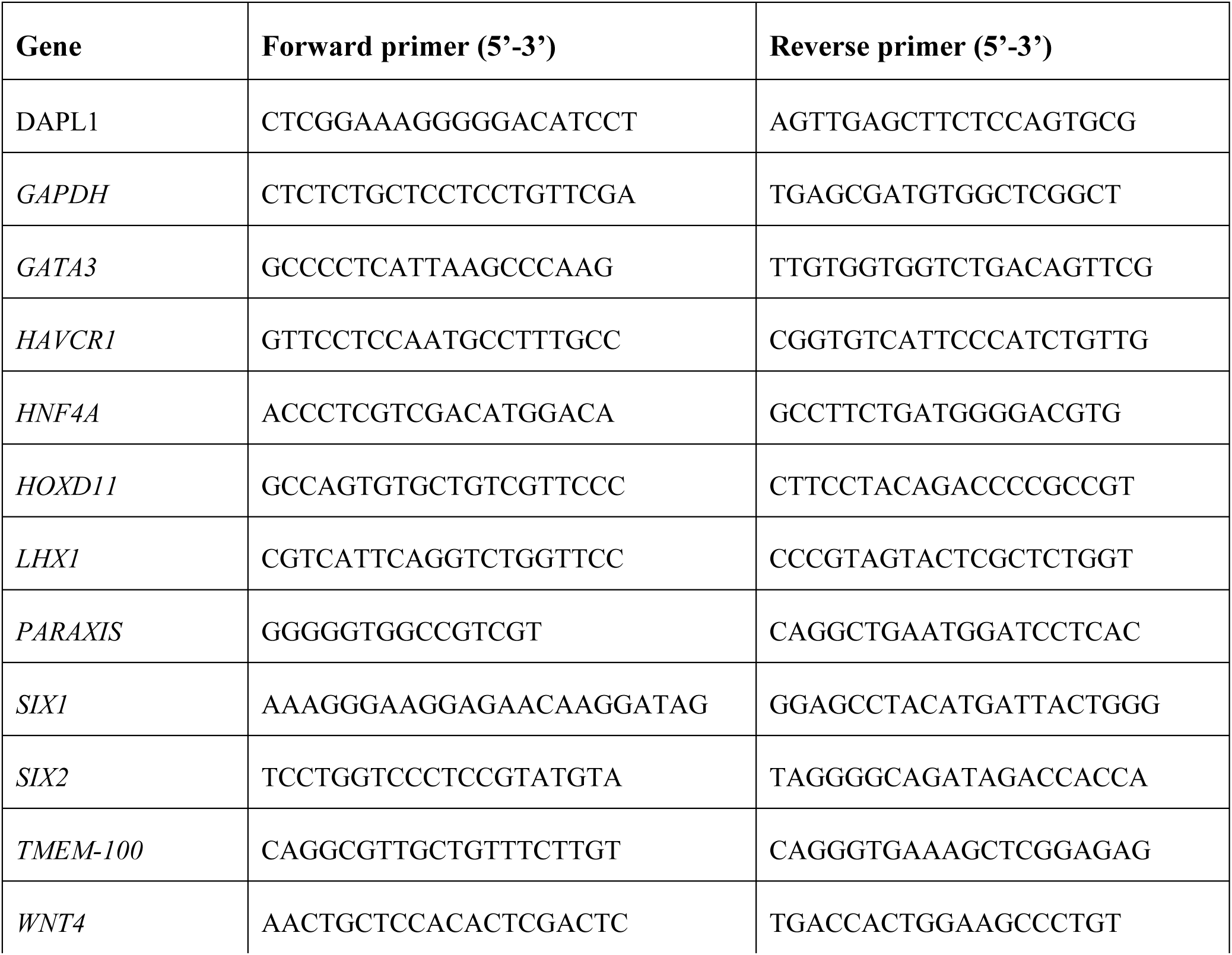
Forward and reverse primers used for qRT-PCR.

### Single cell RNA sequencing (scRNAseq) and dataset generation

The D13+12 dataset was generated using the CRL-2429/SIX2^Cre/Cre^:GAPDH^dual^ iPSC line. The D13 and D13+14 organoids were generated using the CRL1502.C32 with four replicates per time point, where each replicate was derived from an independent well. Cells were dissociated following previously published methods (Lawlor, *et al*., 2021). For the D13 and D13+14 samples, replicates were multiplexed following the method of Soeckius et al. (Stoeckius, *et al*., 2018). Cells were stained for 20 minutes on ice with 1µg of BioLegend TotalSeq-A anti-human hashtag oligo antibody (BioLegend TotalSeq-A0251 to A0258). Cells were washed 3 times then pooled at equal ratios for sequencing. A single library was generated for each suspension/condition, composed of equally sized pools of each replicate (Set 1 – 4). Libraries were generated following the standard 10x Chromium Next GEM Single Cell 3ʹ Reagent Kits v3.1 protocol except that ‘superloading’ of the 10x device was performed with ∼30k cells. Hash tag oligo (HTO) libraries were generated following the BioLegend manufacturer protocol. Sequencing was performed using an Illumina Novoseq.

10x mRNA libraries were demultiplexed using CellRanger (3.1.0) to generate matrices of UMI counts per cell. HTO libraries were demultiplexed using Cite-seq-count (1.4.3) to generate matrices of HTO counts per cell barcode. All data were loaded into Seurat (3.1.4) and HTO libraries were matched to mRNA libraries. Seurat was used to normalise HTO counts and determine cut-offs to assign HTO identity per cell using the *HTODemux* function with the positive.quantile parameter set at 0.99. HTO doublet and unassigned cells were removed, as were cells with mitochondrial content greater than 35% accounting for the increased metabolic activity of renal epithelium (Ransick, *et al*., 2019), number of genes per cell greater than 500 and the number of UMIs less than 100000, to obtain filtered datasets (D13 replicates: 3694 cells [A0251], 3545 cells [A0252], 3785 cells [A0253], 3641 cells [A0254]; D13+14 replicates: 3415 cells [A0255], 2350 cells [A0256], 2904 cells [A0257], 2578 cells [A0258]). The combined datasets contained a median of 3915 genes expressed per cell, with a median of 16352 UMI counts per cell.

### Analysis of scRNAseq datasets

Data was normalised using the SCTransform method (Hafemeister and Satija, 2019) including the regression of cell cycle scores. A 30 component Principal Component Analysis (PCA) was performed, followed by Uniform Manifold Approximation and Projection (UMAP) using these PCA components. Seurat’s graph-based clustering approach was used to identify, with resolutions of 0.7 (D13) and 0.5 (D13+14) chosen for downstream analysis. Marker analysis was performed using the Seurat *FindMarkers* function, using student’s t-test, limited to positive markers (i.e. increased expression within a cluster) above 0.25 log fold-change expressed in at least 10% of cells within a cluster. Marker lists were exported and cluster identities were determined by comparison with published human single cell data (Howden, *et al*., 2019) or Gene ontology analysis using ToppFun (https://toppgene.cchmc.org/enrichment.jsp). The PT cluster was isolated and reanalysed as above to further investigate any subpopulations.

The D13+12 dataset was integrated with an age- and line-matched published dataset (Howden, *et al*., 2019) using the anchor-based method within Seurat (Butler, *et al*., 2018; Stuart, *et al*., 2019). This integrated dataset was analysed as above, isolating the PT cluster and comparing gene expression of cells from both samples within this population.

For DevKidCC analyses, The D13 and D13p14 samples were analysed using DevKidCC (v0.0.3); a hierarchical set of machine-learning binary classifiers trained on a human fetal kidney reference dataset. The classified dataset was then compared to relevant existing single cell organoid datasets using the *DotPlotCompare* function.

For Azimuth analyses, cells were uploaded to the online Azimuth portal at https://app.azimuth.hubmapconsortium.org/app/human-fetus and instructions were followed as per the website for the analysis.

### Agarose bead-mediated morphogen signalling assay

Bioprinted patch organoids were generated and cultured as described previously prior to the addition of morphogen-soaked beads at D7+5 (Lawlor, *et al*., 2021). The day before bead addition, 100µL of Affi-Gel Blue Gel 100 – 200 mesh crosslinked agarose beads (Bio-Rad Laboratories, Hercules, CA), were washed 3 times in PBS via centrifugation. Washed beads were resuspended in 100µL of PBS (control) or 10µM IWR-1 (stock reconstituted according to manufacturer’s instructions; Sigma Aldrich) and incubated for 1 hour at room temperature prior to overnight storage at 4°C. On day 7+5, suspensions were agitated to resuspend beads and 0.3 µL was added to the centre of each patch organoid with the aid of a P2 pipette and dissecting microscope (Leica Microsystems, Wetzlar, Germany). Organoid media (TeSR-E6 [STEMCELL Technologies, Vancouver Canada]) was refreshed every second day prior to harvest at D7+9 for immunofluorescence.

### Quantification of tissue patterning changes in response to IWR soaked beads

Tissue patterning within the radius of beads was quantified using custom Python (3.10.2) scripts, with method as follows. Images (n = 3 per condition, IWR soaked and control) were loaded as Numpy (1.22.1) (Harris, *et al*., 2020) arrays using the Czifile library (2019.7.2) and masks of bead location were generated by manually segmenting each bead using the Napari (0.4.13) labels layer feature. Nephron segments were segmented by applying a gaussian filter to each channel (sigma of 5 pixels) followed by Otsu thresholding (for NPHS1 staining) or multi-otsu thresholding (for LTL, EPCAM) using the second threshold value. All processing was implemented using functions in scikit-image (0.19.2) (van der Walt, *et al*., 2014). The distance of each pixel in the image from the bead edge was calculated using the Euclidian distance transform in Scipy (1.7.3) (Virtanen, *et al*., 2020). These values were used to define the total region within 200 pixels of the bead surface, including the beads themselves. The percentage of pixels assigned to each nephron marker as a proportion of total nephron tissue (defined by the total pixels that were segmented as NPHS1 or EPCAM positive), within the 200 pixel region of each image was then calculated. Scipy was used to conduct t-tests, Matplotlib (3.5.1) was used to generate plots and Napari was used to generate composite images.

### Cisplatin toxicity assay

D13+14 PT-enhanced organoids were exposed through the basolateral compartment of the Transwell tissue culture plate (Corning Incorporated, Corning, NY) to 1mL per well of 20 µM Cisplatin (Accord Healthcare, Durham, NC), or an equivalent volume of PBS, in TeSR-E6 for 24 hours (37°C, 5% CO_2_ and 5% O_2_). Following incubation, organoids within Transwells were washed with PBS and harvested for flow cytometry as described above.

### Fluorescent substrate uptake assays

For albumin uptake assays, D13+14 PT-enhanced organoids (triplicate wells per condition) were incubated in TRITC albumin (1:1000, Sigma Aldrich) and anti-MEGALIN/LRP2 (1:500, pre-incubated with an alpaca Nano-secondary Alexa Fluor 647 secondary antibody diluted in TeSR-E6 culture media via the basolateral compartment of Transwell tissue culture plates and incubated overnight (37°C, 5% CO_2_ and 5% O_2_). Control organoids were incubated in secondary antibody alone. After incubation, plates containing organoids were washed in at least 3 changes of Hanks’ Balanced Salt Solution (HBSS; Thermo Fisher Scientific) for 30 minutes and live-imaged immediately using a ZEISS LSM 780 confocal microscope. For organic cation transport assays, D13+14 PT-enhanced organoids (triplicate wells per condition) were incubated in 4’,6-diamindino-2-phenylindole substrate (DAPI; 1:1000 [Thermo Fisher Scientific]) with 1:500 DRAQ7 dead cell label (Thermo Fisher Scientific]) diluted in TeSR-E6 for 1 hour (37°C, 5% CO_2_ and 5% O_2_). Control organoids were pre-incubated for 15 minutes in 100 µM Cimetidine inhibitor (Sigma Aldrich) prior to incubation for 1 hour in TeSR-E6 containing both inhibitor, substrate, and dead cell label (1:1000 DAPI, 1:500 DRAQ7, 100 µM Cimetidine). Following incubation, substrate and substrate + inhibitor solutions were replaced with HBSS and live-imaged immediately using a ZEISS LSM 780 confocal microscope.

### Viral infection assays

Standard and PT-enhanced organoids grown on Transwells were infected with 10^4^ tissue-culture infectious dose 50 (TCID_50_) of SARS-CoV-2 (Australia/VIC01/2020) in TeSR-E6 media added above the Transwell for 3 hours (virus titration experiments) or below the Transwell with a drop ontop of the organoid for 1 hour (virus localisation experiments). Following incubation (37°C and 5% CO_2_), the viral inoculum was removed and replaced with 1mL of plain TeSR-E6 medium beneath the Transwell as for typical organoid culture (Takasato, *et al*., 2016). Culture medium was collected on days 0, 2, 4, and 6 post-infection for viral titer quantification and replaced with fresh medium. Median TCID_50_ in supernatants were determined, as detailed below, by 10- fold serial dilution in Vero cells and calculated using the Reed and Muench method. Organoids were harvested at 6 days post-infection and fixed with 4% PFA fixation for immunofluorescence.

### Infectious virus titration (Median Tissue Culture Infectious Dose assay; TCID_50_)

Viral titrations were performed on confluent monolayers of Vero cells in 96-well plates. Wells were washed with plain minimum essential media (MEM) and replaced with 180µl of infection media (MEM, 50U/ml Penicillin, 50µg/ml Streptomycin, 2mM GlutaMax, 15mM HEPES and 1µg/ml TPCK-treated Trypsin). 20µl of the samples to be titred were added to four wells and 10-fold serial dilutions were made. Plates were incubated at 37°C and 5% CO_2_. Four days post-infection, SARS-CoV-2-induced cytopathic effect was assessed by microscopy.

### RT-qPCR for SARS-CoV-2 genome

RNA was extracted from supernatant culture media using the QIAamp 96 Virus QIAcube HT Kit (Qiagen). E-gene expression was determined using the SensiFAST Probe No-Rox One Step Kit (Bioline) and the following primers/probes: Fwd: 5’-ACAGGTACGTTAATAGTTAATAGCGT’-3, Rev: ATATTGCAGCAGTACGCACACA and Probe: FAM-ACACTAGCCATCCTTACTGCGCTTCG-BBQ. Viral genome copies were interpolated using a standard curve generated by using a plasmid vector containing the *E*-gene.

## Acknowledgements

This work was supported by the National Health and Medical Research Council (GNT1156440), the National Institutes of Health (UH3DK107344) and the Victorian State Government (DJPR/COVID-19). MHL is an NHMRC Senior Principal Research Fellow (GNT1136085). We acknowledge the Stafford Fox Medical Research Foundation MCRI genome editing facility for the generation of all pluripotent stem cell lines. We thank Maelle Le Moing and the Murdoch Children’s Research Institute Translational Genomics Unit for 10x single cell and hash-tag oligo library preparation and sequencing, and bulk-RNAseq sequencing; Matthew Burton and the Murdoch Children’s Research Institute Microscopy Core; Professor John Rasko and Dr Charles Bailey for providing the SLC6A19 antibody; Dr Dad Abu-Bonsrah for providing the dsRNA antibody; Professor Ralph Baric for providing the GFP-tagged SARS-CoV-2.

## Author Contributions

JMV, MHL, and KS contributed to experimental design and planning. JMV, KST, EG, RR, JN, SM, MS, and SEH performed experiments and developed reagents and methods. SBW and JMV performed bioinformatics analyses. KL performed image analyses. JMV, MHL, and SBW contributed to manuscript preparation. JMV and MHL wrote the manuscript.

## Data availability

All transcriptional profiling datasets have been submitted to GEO (GSE184928). These include scRNAseq from D13 monolayer differentiation, D13+14 PT-enhanced kidney organoids, and D13+12 PT-enhanced kidney organoid. Code and raw data for scRNAseq and image analyses are available through the Github repository (https://github.com/KidneyRegeneration/Vanslambrouck2022).

## Supplementary Figures and legends

**Supplementary Figure 1:**
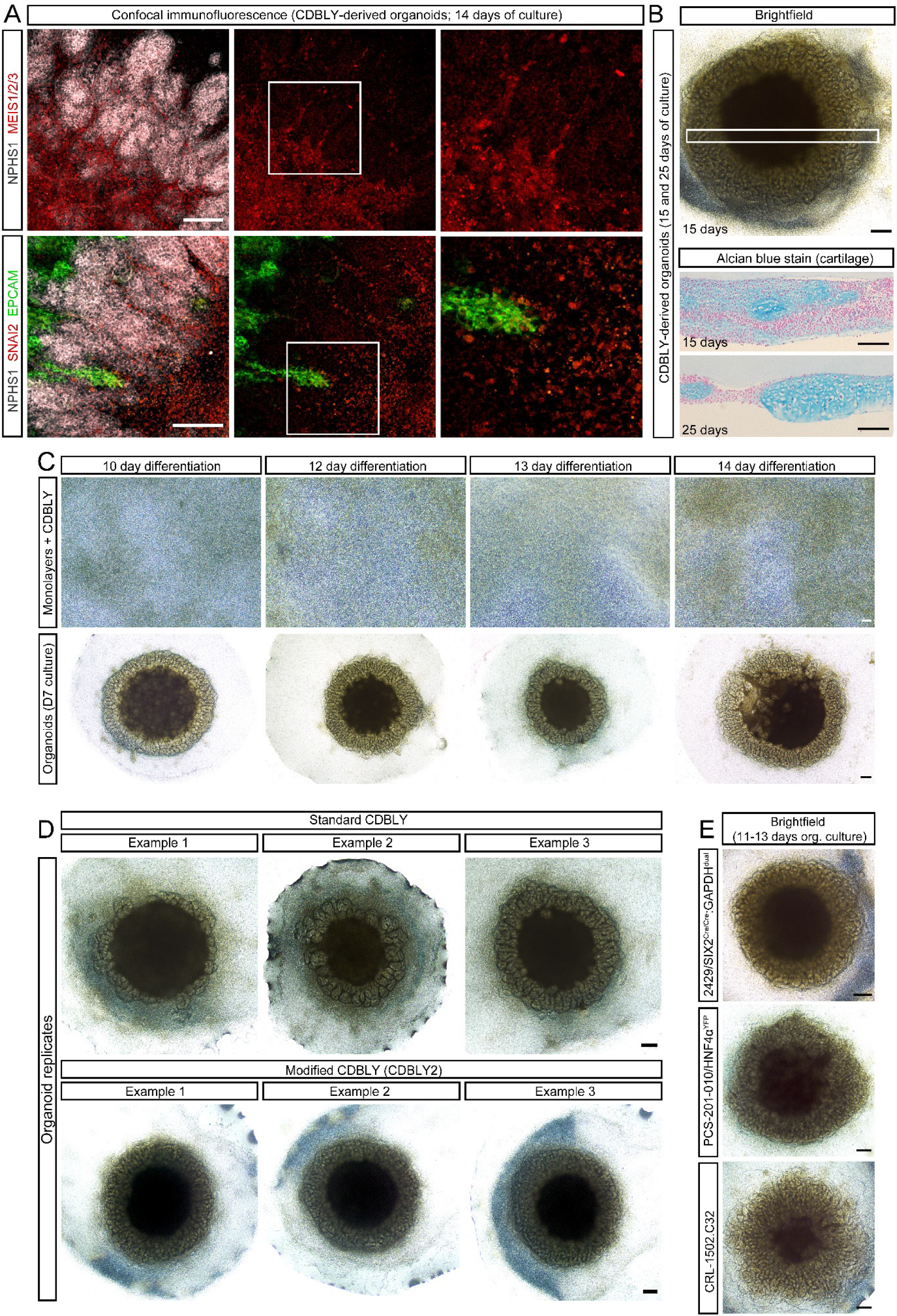
Analyses of central core region and morphology of organoids resulting from extended monolayer differentiation in multiple cell lines. **A.** Confocal immunofluorescence of stromal markers, MEIS1/2/3 (red; top panels) and SNAI2 (red; bottom panels) in central core region of organoids derived from CDBLY-exposed extended monolayer differentiations. Organoids are co-stained for podocytes (NPHS1; grey) and epithelium (EPCAM; green). Scale bars represent 100 µm. **B.** Representative brightfield image (top) of a day 13+15 organoid exposed to CDBLY at monolayer differentiation day 8. White box indicates approximate regions of cross sections shown in bottom panels stained with Alcian blue, indicating patchy cartilage formation in central core region (blue). Scale bars represent 200 µm. **C.** Brightfield images of CDBLY-exposed monolayer differentiations extended for 10, 12, 13, and 14 days and their resulting organoids. Scale bars represent 100 µm (monolayers) and 200 µm (organoids). **D.** Brightfield images showing 3 examples of representative organoid morphologies derived from CDBLY- (5 ng/mL BMP7) and CDBLY2-exposed (10 ng/mL BMP7) monolayer differentiations. Scale bars represent 200 µm. **E.** Brightfield images of D13+11 – D13+13 organoids generated from multiple iPSC lines using extended monolayer differentiation with 5 days x 6µM CHIR exposure (CRL1502.C32; parental iPSC line derived from fetal female skin fibroblasts (Briggs, *et al*., 2013), CRL-2429/SIX2^Cre/Cre^:GAPDH^dual^; lineage tracing reporter iPSC line originally derived from neonatal male foreskin fibroblasts (Howden, *et al*., 2019; Vanslambrouck, *et al*., 2019), and PCS-201-010/HNF4α^YFP^; PT-specific fluorescence reporter iPSC line originally derived from neonatal male skin fibroblasts (Vanslambrouck, *et al*., 2019). Scale bars represent 200 µm.

**Supplementary Figure 2:**
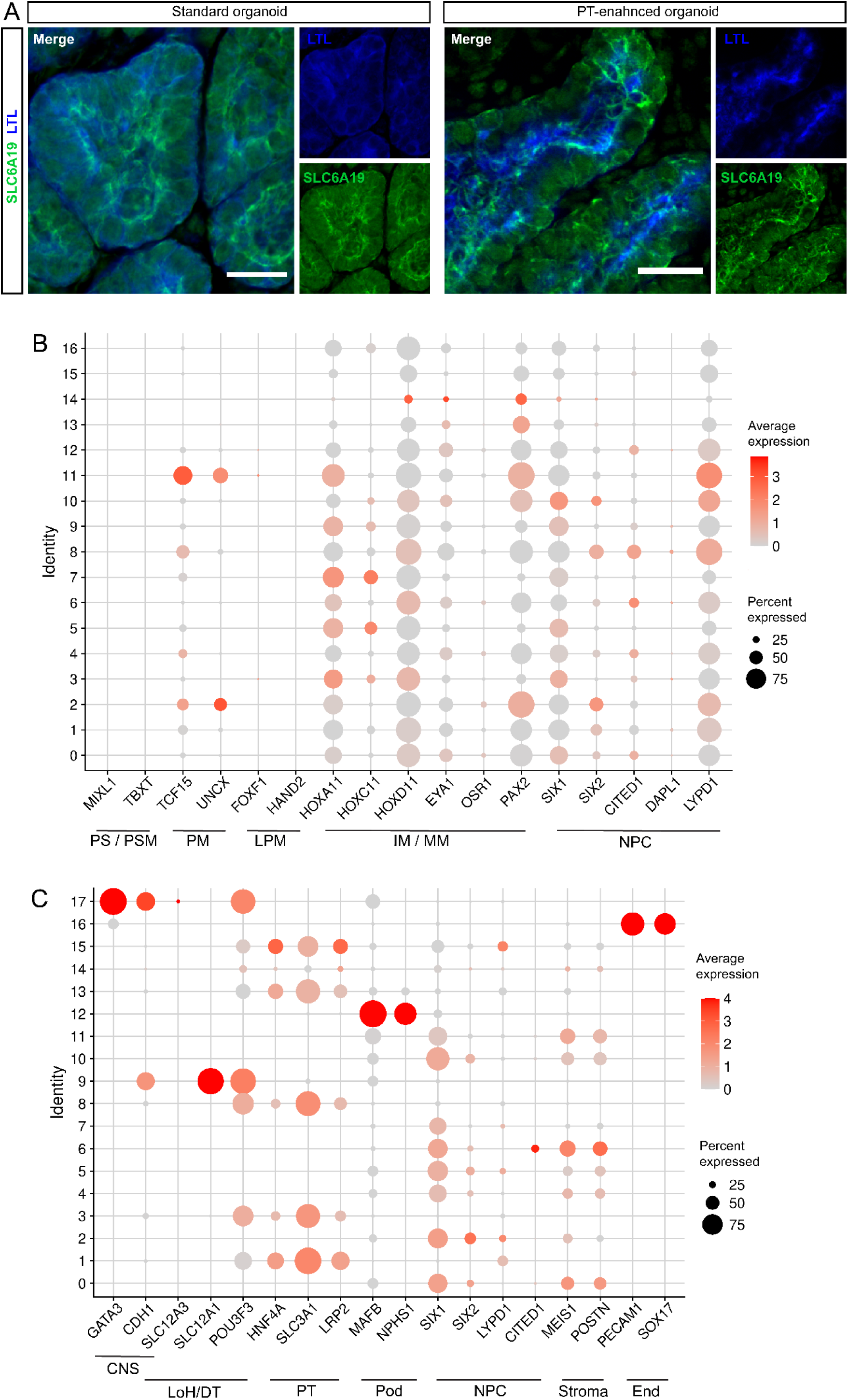
Brush border membrane marker visualisation and scRNAseq cluster marker analyses of D13 monolayers and D13+14 PT-enhanced organoids. **A.** High-resolution confocal microscopy depicting immunofluorescence for apical PT brush border membrane marker LTL (blue) and SLC6A19 (green) in D7+14 (standard) and D13+14 (PT-enhanced) organoids. Scale bars represent 20µm. **B.** Dot plot of D13 combined replicate samples showing expression of early mesenchymal markers preceding metanephric kidney formation across all resolved clusters. Abbreviations: primitive streak (PS), presomitic mesoderm (PSM), paraxial mesoderm (PM), lateral plate mesoderm (LPM), intermediate mesoderm (IM), metanephric mesenchyme (MM), nephron progenitor cells (NPC). **C.** Dotplot of D13+14 combined replicate samples showing expression of kidney-specific markers across all resolved clusters. Abbreviations: connecting segments (CNS), loop of Henle (LoH), distal tubule (DT), proximal tubule (PT), podocyte (Pod), nephron progenitor cell (NPC), endothelium (End).

**Supplementary Figure 3:**
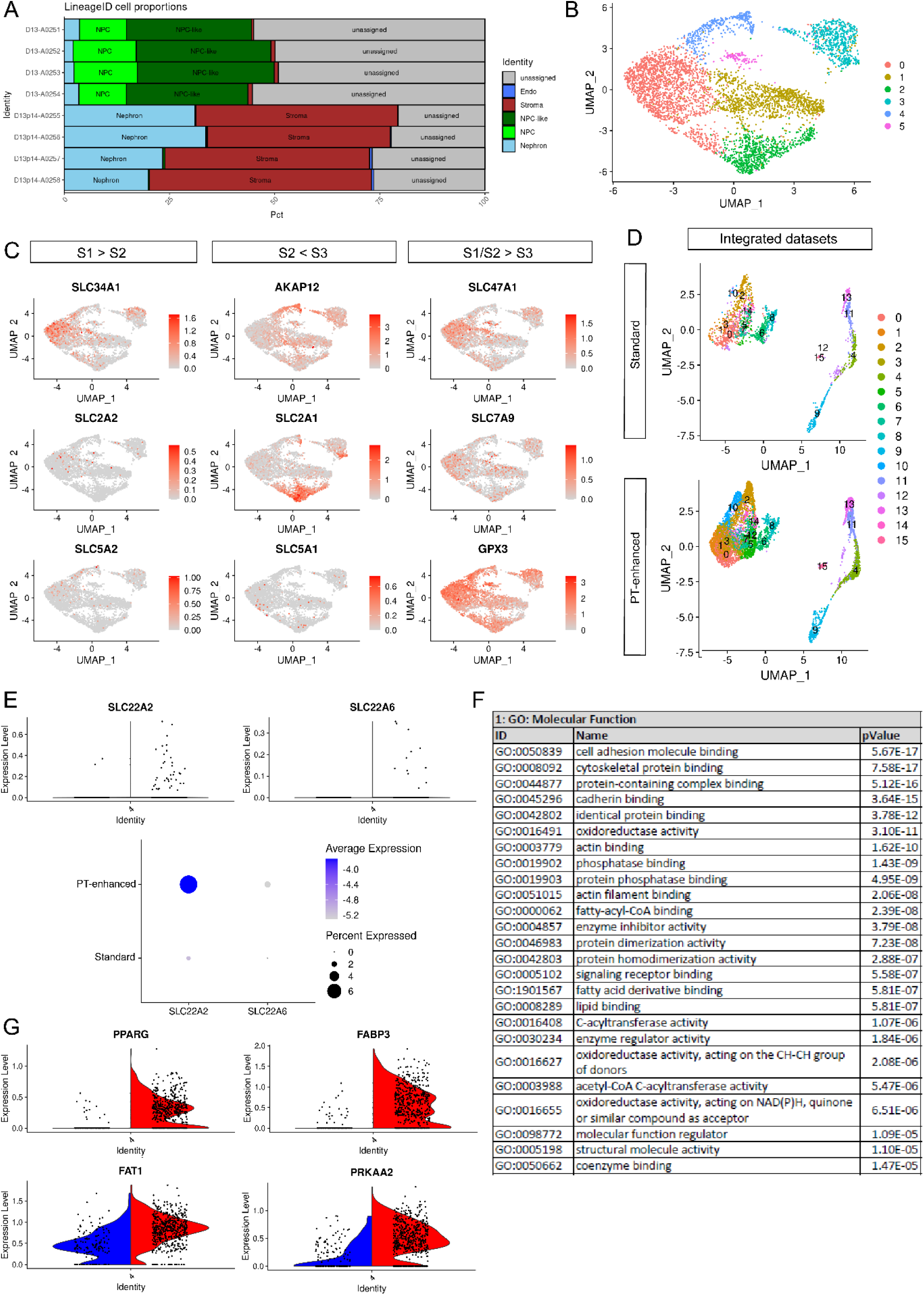
Comparison of D13 monolayer, PT-enhanced organoid, and published scRNAseq datasets. **A.** *ComparePlot* showing proportion of kidney cell types within each D13 and D13+14 replicate. **B.** UMAP plot of isolated PT clusters from D13+14 PT-enhanced organoids, re-clustered to resolve 6 distinct cell populations. **C.** UMAP plots showing the expression of S1, S2, and S3 segment markers within the isolated PT population of D13+14 organoids. **D.** UMAP plots from integrated analyses of PT-enhanced organoids and our existing standard organoid dataset (Howden, *et al*., 2019) (iPSC line- and age-matched). Clustering resolved 15 distinct cell populations for each sample in the integrated datasets. **E.** Violin (top panels) and dot plots (bottom panel) depicting *SLC22A2* and *SLC22A6* expression within the PT cluster of integrated PT-enhanced and standard organoid datasets from (**D**) (left and right on violin plots, respectively). **F.** Table depicting top 25 GO terms arising from unbiased ToppFun GO Molecular Function analyses of significantly differentially expressed genes between standard (blue, left) and PT-enhanced (red, right) organoid datasets from (**D**)**. G.** Violin plots comparing examples of genes involved in fatty acid metabolism in standard (blue, left) and PT-enhanced (red, right) organoid datasets from **(D**).

**Supplementary Figure 4:**
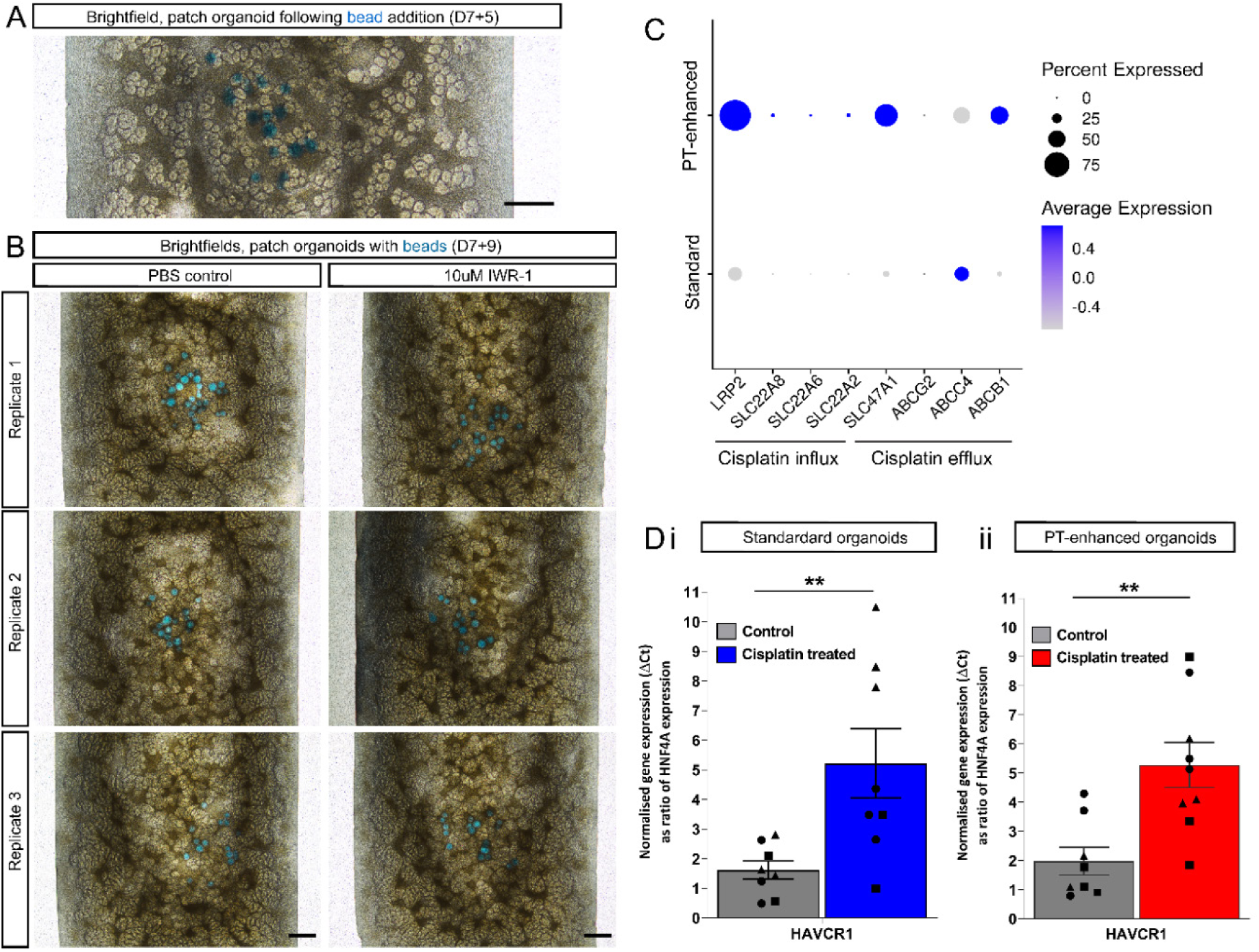
Defining the mechanism of nephron directionality using IWR-1-soaked agarose beads and exploring PT-enhanced organoid functionality through cisplatin response. **A.** Brightfield image of a D7+5 standard bioprinted patch organoid immediately after the addition of agarose beads (blue) depicting their contact with forming renal vesicle structures. Scale bar represents 200 µm. **C.** Brightfield images of D7+9 bioprinted patch organoids containing PSB-soaked or IWR1-soaked (left and right panels, respectively) beads (blue). Scale bars represent 200µm. **C.** scRNAseq dotplot comparing the expression of cisplatin influx and efflux transporters within the PT cluster of integrated PT-enhanced and existing standard organoid datasets (Howden, *et al*., 2019) (iPSC line-and age-matched). **D.** qRT-PCR analyses depicting KIM-1 gene (*HAVCR1*) expression in (**i**) standard (blue) and (**ii**) PT-enhanced (red) organoids treated with cisplatin, compared to their respective PBS-treated controls (grey). *HAVCR1* gene expression values are normalised to the housekeeping gene *GAPDH* (2^-ΔCt^) and expressed as a ratio of *HNF4A* to compensate for differences in proximal tubule proportion. Error bars represent SEM from n = 8 (control) and n = 9 (cisplatin-treated) biological replicates across 3 replicate experiments as indicated. Statistical significance was determined using an unpaired t test. Asterisks represent P values (**; P ≤ 0.01) adjusted for multiple comparisons using the Holm-Sidak method alpha = 0.05.

**Supplementary Figure 5:**
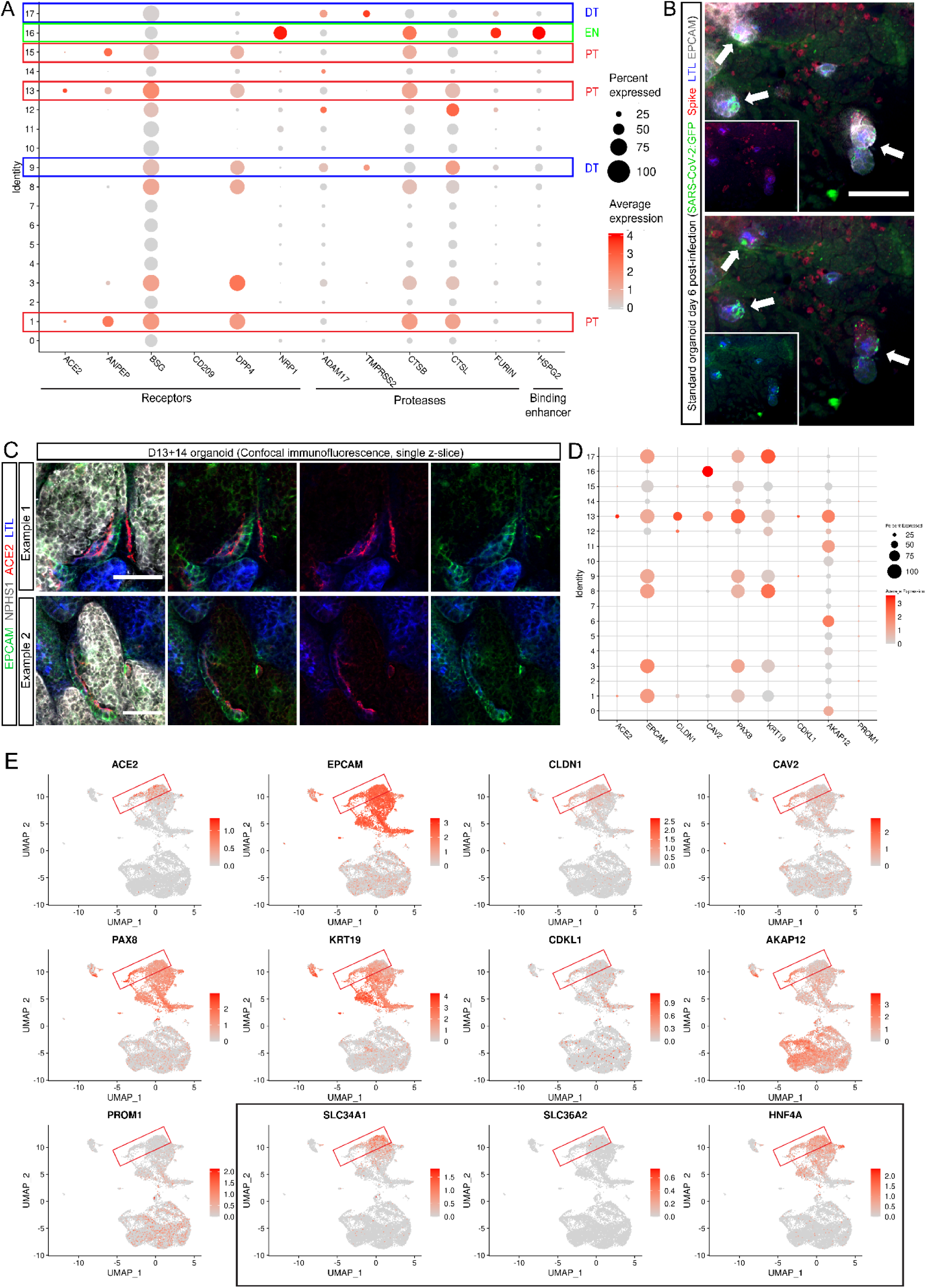
Expression of SARS-CoV-2 entry factors and infectious particles in kidney organoids. **A.** scRNAseq analysis of SARS-CoV-2 entry factor expression in D13+14 kidney organoids. Boxes outline proximal (red), distal (blue), and endothelial (green) clusters. **B.** Confocal immunofluorescence of a D13+20 organoid 6 days post-infection with GFP-tagged SARS-CoV-2 confirming the presence of GFP-positive (green) and spike protein-expressing (red) mature and replicating virus within EPCAM^+^/LTL positive (blue) PTs as well as the interstitium. Insets depict 2-channel overlays of larger merged images. Arrows indicate examples of viral GFP in LTL-positive tubules Scale bar represents 50µm. **C.** Confocal immunofluorescence of a D13+14 PT-enhanced organoid depicting apical ACE2 (red) expression on EPCAM-positive (green) cells entering the early portion of Bowman’s capsule surrounding NPHS1-positive (grey) podocytes of glomeruli. LTL (blue) marks PTs. Scale bars represent 50µm. **D-E.** Analyses of D13+14 scRNAseq dataset displayed as a dot plot (**D**) and feature plots (**E**), depicting expression of *ACE2* in clusters co-expressing markers of cuboidal and intermediate PECs, as well S1-specific markers of PT. Boxes in (**E**) highlight the key region of overlapping expression in the PT clusters.

**Supplementary Figure 6:**
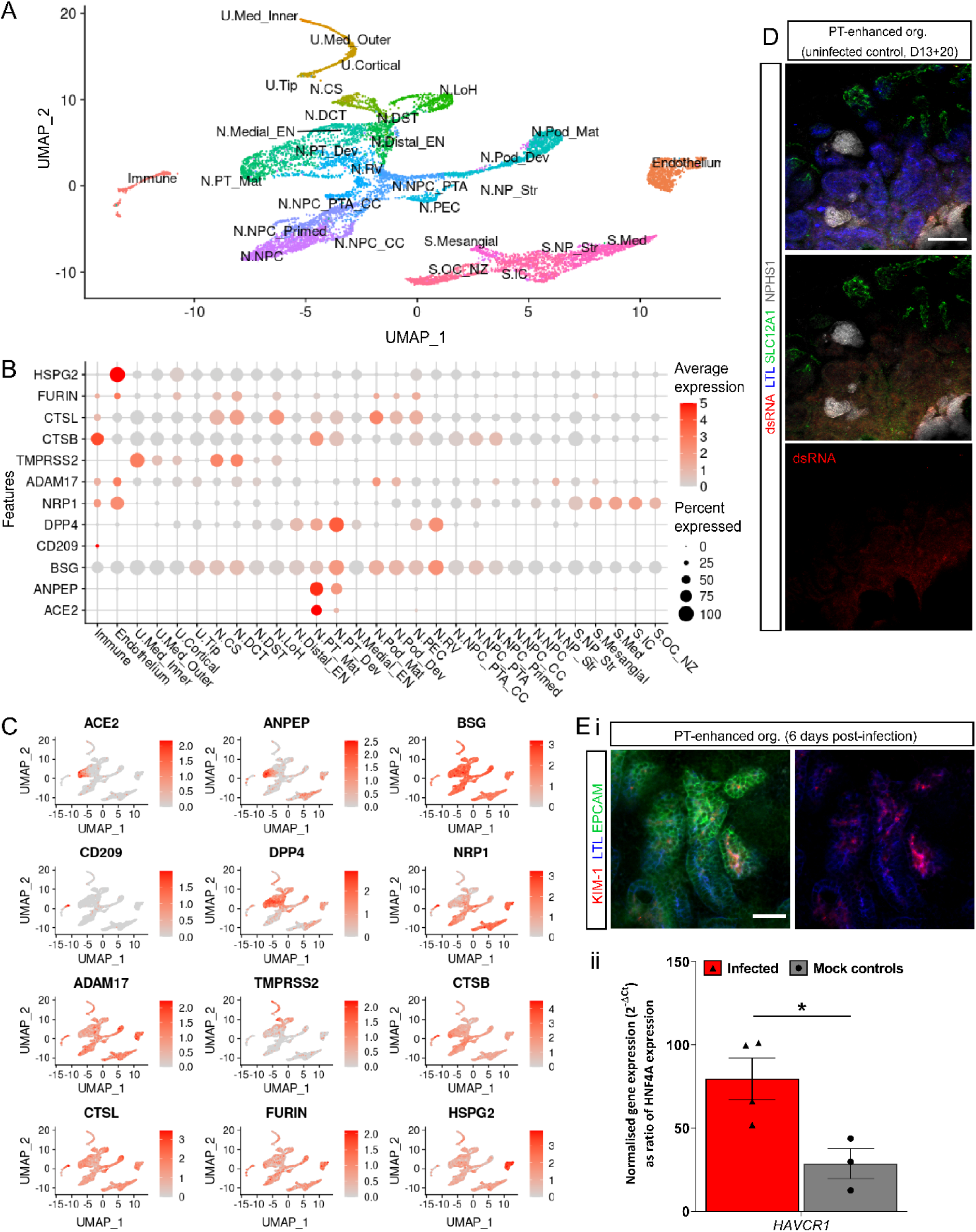
Distribution of SARS-CoV-2 entry factors in PT-enhanced kidney organoids. **A-C.** Single cell RNAseq analysis of existing week 11 – 18 mixed human fetal kidney reference datasets (Hochane, *et al*., 2019; Tran, *et al*., 2019; Holloway, *et al*., 2020) displayed in UMAP (**A** and **C**) and dot plot (**B**) formats confirming the resolution of distinct kidney cell clusters and the expression of SARS-CoV-2 entry factors within each cluster. Cluster abbreviations: ureteric (U), medullary (Med), nephron (N), connecting segment (CS), distal convoluted tubule (DCT), distal straight tubule (DST), loop of Henle (LoH), early nephron (EN), proximal tubule (PT), developing (Dev), maturing (Mat), podocyte (Pod), parietal epithelial cell (PEC), renal vesicle (RV), nephron progenitor cell (NPC), pre-tubular aggregate (PTA), cycling cells (CC), nephron progenitor (NP), stroma (S), stromal (Str), inner cortical (IC), outer cortical (OC), nephrogenic zone (NZ). **D.** Confocal immunofluorescence of a representative matched experimental control organoid for SARS-CoV-2 infection experiments (uninfected D13+20 organoid). Control organoids were stained for viral RNA (dsRNA; red), PT (LTL; blue), LoH (SLC12A1; green), and podocytes (NPHS1; grey). Scale bar represents 100µm. **Ei.** Confocal immunofluorescence of a PT-enhanced organoid 6 days post infection with SARS-CoV-2 depicting KIM-1 (red) expression within EPCAM-positive/LTL-positive (green/blue) PTs. Scale bar represents 50µm. **Eii.** qRT-PCR analysis depicting KIM-1 gene (*HAVCR1*) expression in infected (red) and mock control (grey; uninfected) PT-enhanced kidney organoids at 6 days post-infection. *HAVCR1* gene expression values are normalised to *GAPDH* housekeeping gene and expressed as a ratio of *HNF4A* expression to compensate for differenced in proximal tubule proportion. Error bars represent SEM from n = 4 (infected) and n = 3 (mock) biological replicate organoids. Statistical significance was determined using an unpaired t test. Asterisks represent P values (*; P ≤ 0.05) adjusted for multiple comparisons using the Holm-Sidak method alpha = 0.05.

## Supplementary Tables

**Supplementary Table 1:** Differentially expressed (DE) genes by cluster in day 13 (D13) monolayers derived from extended differentiation in CDBLY2 (see: Vanslambrouck JM et al_Supplementary Table 1).

**Supplementary Table 2:** Differentially expressed (DE) genes by cluster in D13+14 organoids derived from D13 monolayers (see: Vanslambrouck JM et al_Supplementary Table 2).

**Supplementary Table 3:** Quantification of nephron structures in organoids exposed to IWR1- soaked and PBS-soaked agarose beads (see: Vanslambrouck JM et al_Supplementary Table 3).

