## Supplementary Table 1 for "Enhanced metanephric specification to functional proximal tubule enables toxicity screening and infectious disease modelling in kidney organoids"

| **Cluster 0** | | |
|---|---|---|
| Gene | LogFC | pVal |
| PAX8 | 0.341722 | 0 |
| CDH6 | 0.326793 | 2.1E-252 |
| DDIT4 | 0.286203 | 7.3E-221 |

| **Cluster 1** | | |
| --- | --- | --- |
| Gene | LogFC | pVal |
| HIST1H4C | 1.471057 | 0 |
| TUBA1B | 0.742174 | 0 |
| PCLAF | 0.614055 | 0 |
| NUSAP1 | 0.607118 | 0 |
| HIST1H1D | 0.594679 | 0 |
| HMGB2 | 0.587079 | 0 |
| HIST1H1A | 0.583082 | 0 |
| TYMS | 0.559061 | 0 |
| CKS1B | 0.554846 | 0 |
| H2AFX | 0.553989 | 0 |
| TOP2A | 0.55328 | 0 |
| UBE2C | 0.533803 | 0 |
| RANBP1 | 0.529632 | 0 |
| RAD51AP1 | 0.487828 | 0 |
| UBE2T | 0.467349 | 0 |
| HIST1H1C | 0.461396 | 0 |
| TUBB | 0.461121 | 0 |
| BIRC5 | 0.458944 | 0 |
| AURKB | 0.44896 | 0 |
| TUBB4B | 0.441417 | 0 |
| CDK1 | 0.438404 | 0 |
| CENPF | 0.436111 | 0 |
| CKS2 | 0.426608 | 0 |
| H2AFZ | 0.425736 | 0 |
| HMGN2 | 0.425091 | 0 |
| SMC4 | 0.423804 | 0 |
| MAD2L1 | 0.413449 | 0 |
| HMGB1 | 0.413184 | 0 |
| STMN1 | 0.401246 | 0 |
| MKI67 | 0.395393 | 0 |
| KIFC1 | 0.390757 | 0 |
| DEK | 0.38078 | 0 |
| RRM2 | 0.377799 | 0 |
| SPC25 | 0.374375 | 0 |
| CENPW | 0.37315 | 0 |
| HSP90AA1 | 0.368847 | 0 |
| CDCA4 | 0.368551 | 0 |
| ITGB3BP | 0.366293 | 0 |
| PBK | 0.363226 | 0 |
| FABP5 | 0.360583 | 0 |
| NDC80 | 0.355685 | 0 |
| H1FX | 0.349957 | 0 |
| NASP | 0.348923 | 0 |
| MND1 | 0.347162 | 0 |
| IER2 | 0.342522 | 0 |
| PA2G4 | 0.342101 | 0 |
| ZWINT | 0.34148 | 0 |
| DLGAP5 | 0.338209 | 0 |
| TPX2 | 0.338004 | 0 |
| HIST1H1B | 0.336148 | 0 |
| UBE2S | 0.334718 | 0 |
| NUF2 | 0.334289 | 0 |
| SNRNP25 | 0.329749 | 0 |
| MCM7 | 0.327684 | 0 |

| | | |
|---|---|---|
| JUNB | 0.327422 | 9.6E-272 |
| PCNA | 0.326804 | 0 |
| CENPH | 0.322469 | 0 |
| TK1 | 0.319804 | 0 |
| SNRPB | 0.319627 | 0 |
| CENPK | 0.31782 | 0 |
| UBC | 0.317679 | 0 |
| MYBL2 | 0.314126 | 0 |
| CENPN | 0.312891 | 0 |
| TMSB15A | 0.312743 | 0 |
| NCAPG | 0.311669 | 0 |
| KIF22 | 0.309837 | 0 |
| DUT | 0.308756 | 1.7E-266 |
| GMNN | 0.308652 | 0 |
| H2AFY | 0.306226 | 0 |
| BUB3 | 0.305075 | 0 |
| FBXO5 | 0.300937 | 0 |
| RRM1 | 0.296969 | 0 |
| EGR1 | 0.296231 | 4.2E-242 |
| HNRNPAB | 0.295858 | 0 |
| SRRM1 | 0.295839 | 0 |
| PTTG1 | 0.293866 | 0 |
| CMC2 | 0.293709 | 0 |
| ORC6 | 0.292688 | 0 |
| HIST2H2A | 0.291379 | 0 |
| CDCA3 | 0.290243 | 1E-303 |
| GGH | 0.290142 | 0 |
| ANAPC11 | 0.288422 | 0 |
| KPNA2 | 0.287034 | 0 |
| SIVA1 | 0.284251 | 0 |
| KIF20B | 0.281166 | 0 |
| CENPM | 0.279778 | 0 |
| ALYREF | 0.27616 | 0 |
| DNMT1 | 0.275472 | 0 |
| SRSF7 | 0.273792 | 0 |
| FEN1 | 0.272153 | 1.4E-294 |
| DTYMK | 0.271123 | 0 |
| CLSPN | 0.269216 | 5.4E-290 |
| NUCKS1 | 0.268637 | 0 |
| H2AFV | 0.26854 | 0 |
| FOS | 0.267999 | 2E-202 |
| ERH | 0.267151 | 0 |
| CDT1 | 0.266726 | 8E-306 |
| HIST1H1E | 0.262637 | 0 |
| VRK1 | 0.25948 | 0 |
| SNRPG | 0.257336 | 0 |
| DUSP1 | 0.255089 | 1.4E-142 |
| CENPV | 0.25383 | 6.1E-275 |
| SRSF2 | 0.252871 | 0 |
| PSMC3 | 0.25257 | 0 |
| PSIP1 | 0.251935 | 0 |
| SUPT16H | 0.250391 | 7.4E-274 |

| **Cluster 2** | | |
|---|---|---|
| Gene | LogFC | pVal |
| PRPH | 1.070159 | 0 |
| AGT | 0.891092 | 0 |
| IGFBP5 | 0.782874 | 0 |
| CDKN1C | 0.745328 | 9.3E-225 |
| EMX2 | 0.646487 | 1.9E-255 |
| S100A1 | 0.643734 | 0 |
| TUBB2B | 0.637737 | 0 |
| GADD45B | 0.627962 | 0 |
| SPINT2 | 0.600106 | 0 |
| KRT18 | 0.598156 | 0 |
| BST2 | 0.598031 | 5E-296 |
| KHDRBS3 | 0.574302 | 3.9E-242 |
| CAST | 0.562439 | 5.8E-251 |
| PAX2 | 0.552636 | 0 |
| SCT | 0.552425 | 3E-161 |
| DBI | 0.516564 | 1E-278 |
| CST3 | 0.484193 | 0 |
| DUSP1 | 0.483802 | 6.1E-227 |
| APOE | 0.480318 | 1.9E-267 |
| KLK6 | 0.472431 | 0 |
| IGFBP7 | 0.469511 | 7.8E-163 |
| SAT1 | 0.449915 | 1.1E-226 |
| S100A4 | 0.445333 | 1.2E-119 |
| VAMP8 | 0.421174 | 0 |
| TM7SF2 | 0.408999 | 5.9E-291 |
| CADM1 | 0.407253 | 5E-291 |
| CD24 | 0.3966 | 2.7E-249 |
| CD9 | 0.383194 | 1E-215 |
| COTL1 | 0.378771 | 0 |
| S100A6 | 0.374378 | 7.2E-145 |
| LHX1 | 0.373697 | 1.2E-124 |
| HOXD1 | 0.372818 | 4.9E-207 |
| PDLIM1 | 0.36887 | 2.4E-239 |
| EGFL7 | 0.368458 | 3.2E-273 |
| JAG1 | 0.3668 | 1.4E-201 |
| UNCX | 0.365519 | 1E-126 |
| BCAM | 0.35883 | 2.6E-207 |
| UCHL1 | 0.349676 | 2.3E-154 |
| IFITM1 | 0.343097 | 1.7E-169 |
| PKDCC | 0.342682 | 3.7E-222 |
| UCP2 | 0.341127 | 1.5E-276 |
| JUNB | 0.340187 | 9.7E-144 |
| CLDN4 | 0.336916 | 6.3E-216 |
| CEBPD | 0.335492 | 3.3E-137 |
| EPHX1 | 0.33397 | 2.2E-228 |
| ITM2B | 0.331131 | 0 |
| ID4 | 0.32598 | 2.7E-267 |
| TSC22D3 | 0.325504 | 5.7E-176 |
| TFPI2 | 0.322197 | 1.45E-61 |
| HOXB7 | 0.319628 | 1.4E-175 |
| RGS2 | 0.314907 | 3.1E-157 |
| TSC22D1 | 0.313521 | 4.9E-118 |
| CALM1 | 0.311166 | 7.9E-176 |
| BTG2 | 0.306265 | 6.9E-165 |

| | | |
|---|---|---|
| ANXA2 | 0.305569 | 1.23E-80 |
| EMID1 | 0.298582 | 3E-200 |
| HES1 | 0.298485 | 2.58E-81 |
| KRT8 | 0.290056 | 3.7E-210 |
| CYBA | 0.288884 | 5.4E-188 |
| LYPD6B | 0.28685 | 7.4E-168 |
| IGFBP4 | 0.284828 | 6.16E-94 |
| TUBA1A | 0.283894 | 8.6E-121 |
| HLA-C | 0.28237 | 2.3E-197 |
| ADGRG2 | 0.282131 | 2.6E-162 |
| NKAIN4 | 0.280733 | 2.5E-149 |
| SLC32A1 | 0.28008 | 1.2E-206 |
| TRIM27 | 0.279044 | 3.9E-196 |
| SLC3A2 | 0.279025 | 1.7E-173 |
| IGFBP2 | 0.278194 | 4.1E-279 |
| KLF6 | 0.278145 | 2.4E-124 |
| CXXC5 | 0.275001 | 8.2E-250 |
| LINC02381 | 0.274416 | 2E-173 |
| CLIC1 | 0.273071 | 2.8E-210 |
| SOCS3 | 0.272286 | 5.4E-117 |
| NR2F6 | 0.271253 | 6.8E-293 |
| CITED2 | 0.267859 | 5.7E-144 |
| EPCAM | 0.26721 | 9.8E-117 |
| CHMP1B | 0.263976 | 3.7E-135 |
| RGS4 | 0.263934 | 3.1E-100 |
| SLC16A3 | 0.260711 | 2.3E-130 |
| SCUBE3 | 0.260111 | 3.3E-137 |
| CD63 | 0.258148 | 9.5E-273 |
| ID2 | 0.255147 | 1E-154 |
| HOXB-AS3 | 0.253591 | 8.21E-78 |
| NLRP1 | 0.250565 | 1.7E-124 |
| PRICKLE1 | 0.250373 | 3.8E-109 |

| **Cluster 3** | | |
|---|---|---|
| Gene | LogFC | pVal |
| VIM | 0.535081 | 0 |
| RGS5 | 0.46602 | 3.7E-119 |
| NPW | 0.426838 | 9.4E-180 |
| GPC3 | 0.379946 | 1.7E-131 |
| S100A10 | 0.350846 | 2.2E-114 |
| PCOLCE | 0.304173 | 2.5E-106 |
| AC004540. | 0.303402 | 7.05E-88 |
| SPATS2L | 0.2843 | 1.78E-93 |
| ROBO2 | 0.279279 | 5.9E-121 |
| TPM2 | 0.271774 | 4.9E-144 |
| UBE2E3 | 0.267724 | 4.6E-144 |
| CTGF | 0.261583 | 3.81E-28 |
| SSR3 | 0.257133 | 1.4E-123 |
| DES | 0.255938 | 7.33E-80 |
| COL1A1 | 0.253819 | 2.31E-88 |

| **Cluster 4** | | |
|---|---|---|
| Gene | LogFC | pVal |
| DUT | 0.603753 | 5.2E-251 |
| GINS2 | 0.561835 | 0 |
| MCM3 | 0.498824 | 2.2E-271 |
| PCNA | 0.478363 | 1.6E-289 |
| UNG | 0.418083 | 7.8E-211 |
| MCM4 | 0.407158 | 1.8E-211 |
| MCM7 | 0.390327 | 5E-230 |
| MCM6 | 0.387732 | 1.2E-202 |
| CDT1 | 0.386808 | 3.5E-196 |
| SLBP | 0.386671 | 2E-178 |
| MCM5 | 0.382956 | 2.4E-213 |
| PCLAF | 0.378793 | 8.3E-195 |
| CDC6 | 0.376249 | 2.5E-188 |
| FEN1 | 0.372938 | 4.4E-187 |
| CLSPN | 0.367888 | 1.1E-167 |
| NASP | 0.358488 | 2E-303 |
| HELLS | 0.357342 | 1.4E-180 |
| ORC6 | 0.350938 | 1.6E-207 |
| GMNN | 0.337277 | 1.2E-197 |
| MSH6 | 0.333903 | 1.4E-133 |
| DHFR | 0.328792 | 3.9E-137 |
| TK1 | 0.319929 | 1.1E-136 |
| CDCA7 | 0.317732 | 8E-162 |
| CHEK1 | 0.310843 | 7.3E-149 |
| RFC2 | 0.306963 | 1.7E-159 |
| DEK | 0.304549 | 1.6E-201 |
| E2F1 | 0.294905 | 3.5E-145 |
| TYMS | 0.293719 | 2.8E-138 |
| MCM10 | 0.27839 | 1.4E-126 |
| POLD3 | 0.27356 | 1E-122 |
| DTL | 0.27298 | 5E-137 |
| RFC4 | 0.270154 | 5.3E-123 |
| SNRNP25 | 0.270056 | 3.3E-115 |
| UHRF1 | 0.258579 | 1.6E-122 |

| **Cluster 5** | | |
|---|---|---|
| Gene | LogFC | pVal |
| TWIST1 | 1.103201 | 0 |
| PRRX1 | 0.871018 | 0 |
| VIM | 0.830146 | 0 |
| PCOLCE | 0.817552 | 0 |
| CDC42EP5 | 0.581643 | 0 |
| EMP3 | 0.575057 | 9E-303 |
| CALD1 | 0.554954 | 0 |
| A2M | 0.551243 | 1.4E-229 |
| GNG11 | 0.514068 | 4.5E-239 |
| BGN | 0.505751 | 7.8E-250 |
| COL1A2 | 0.477317 | 0 |
| LMO4 | 0.467604 | 1.2E-229 |
| COL3A1 | 0.4656 | 1.3E-204 |
| LIMCH1 | 0.457176 | 3.3E-221 |
| LGALS1 | 0.454195 | 2.9E-114 |
| PCDH9 | 0.453987 | 1.7E-193 |
| PDGFRA | 0.439963 | 5.5E-182 |
| MARCKS | 0.439479 | 0 |
| PTN | 0.432311 | 5.7E-100 |
| TPM2 | 0.431627 | 2E-262 |
| CLMP | 0.416105 | 2.4E-194 |
| IGDCC3 | 0.413201 | 6.9E-246 |
| PRRX2 | 0.409523 | 8.2E-151 |
| TAGLN | 0.406858 | 7.7E-106 |
| EVA1B | 0.405334 | 1.1E-216 |
| SERPINF1 | 0.40233 | 2.3E-183 |
| THSD7A | 0.391699 | 1.4E-173 |
| MFAP4 | 0.387725 | 1.2E-194 |
| CYR61 | 0.377108 | 2.34E-88 |
| AC103702. | 0.374206 | 4.8E-149 |
| POSTN | 0.370552 | 1.5E-147 |
| MYL9 | 0.366852 | 4.4E-129 |
| EMILIN1 | 0.366522 | 9.4E-190 |
| S100A11 | 0.364472 | 2.1E-282 |
| COL6A2 | 0.34741 | 3E-161 |
| TCF12 | 0.342789 | 1.5E-146 |
| FHL3 | 0.341633 | 1.4E-144 |
| SERPINH1 | 0.33992 | 9.3E-183 |
| RARRES2 | 0.337804 | 1.55E-92 |
| LIMA1 | 0.33721 | 1.2E-153 |
| MSX2 | 0.336857 | 5.2E-159 |
| TNC | 0.332323 | 2.1E-136 |
| TPD52L1 | 0.32839 | 9.64E-93 |
| MAGED2 | 0.327311 | 2.2E-196 |
| C12orf57 | 0.325818 | 9.5E-223 |
| ID1 | 0.324977 | 5.2E-167 |
| TPM4 | 0.31787 | 2.8E-144 |
| UBE2E3 | 0.314923 | 5.9E-175 |
| PTH1R | 0.312539 | 4E-144 |
| PCDH18 | 0.308682 | 3.3E-133 |
| MSX1 | 0.299065 | 8.65E-80 |
| HGF | 0.29768 | 1.66E-94 |
| OAF | 0.296293 | 6.4E-156 |
| CNN2 | 0.29604 | 4.2E-131 |

| | | |
|---|---|---|
| ACTB | 0.29037 | 1.3E-143 |
| HAPLN1 | 0.289558 | 7.3E-105 |
| GLT8D2 | 0.288097 | 2.8E-144 |
| FAM89A | 0.286087 | 2.5E-132 |
| LGALS3 | 0.285606 | 3.1E-112 |
| COL1A1 | 0.285417 | 6.5E-100 |
| AFF3 | 0.283146 | 2.2E-130 |
| RHOBTB3 | 0.28253 | 1.3E-115 |
| CTHRC1 | 0.281461 | 3.49E-57 |
| PIK3R1 | 0.276092 | 1.4E-108 |
| OLFML3 | 0.272738 | 1.4E-122 |
| STOM | 0.268992 | 7.6E-119 |
| IL11RA | 0.258784 | 4.9E-107 |
| HEY1 | 0.255941 | 7.72E-89 |
| LDB2 | 0.250755 | 2.8E-119 |

| Cluster 6 | | |
| --- | --- | --- |
| Gene | LogFC | pVal |
| PTTG1 | 0.870783 | 8E-180 |
| CDC20 | 0.773402 | 4.2E-138 |
| HMGB2 | 0.616724 | 6.15E-66 |
| CCNB2 | 0.60957 | 9.3E-128 |
| HSP90AA1 | 0.584261 | 0 |
| PTMS | 0.583786 | 0 |
| BIRC5 | 0.583079 | 8.7E-191 |
| UBE2S | 0.575758 | 1.73E-47 |
| CENPF | 0.571305 | 5.1E-127 |
| CCNB1 | 0.567303 | 1.69E-28 |
| HNRNPM | 0.50979 | 1.1E-251 |
| HMGN2 | 0.47231 | 2.2E-285 |
| HMGB3 | 0.47059 | 4.9E-133 |
| HOTAIRM1 | 0.467122 | 8.8E-135 |
| TUBA1C | 0.465062 | 1.9E-106 |
| HMGB1 | 0.449877 | 6.2E-294 |
| ODC1 | 0.440546 | 1.7E-125 |
| H2AFZ | 0.439758 | 1.6E-265 |
| NUCKS1 | 0.433433 | 7.2E-224 |
| HNRNPA2 | 0.426148 | 3.5E-276 |
| PPP1R14E | 0.408687 | 7.4E-208 |
| TROAP | 0.404713 | 3E-105 |
| HMMR | 0.399172 | 1.45E-46 |
| NDUFB4 | 0.398065 | 2.9E-233 |
| HSP90B1 | 0.396825 | 3.9E-213 |
| LDHA | 0.391601 | 3.8E-170 |
| STMN1 | 0.388838 | 1E-306 |
| PSMB6 | 0.384452 | 1.2E-207 |
| HMGN1 | 0.381511 | 1E-299 |
| OAZ1 | 0.380355 | 7.7E-199 |
| PSMA7 | 0.379054 | 1.6E-206 |
| CCT5 | 0.374009 | 6.2E-137 |
| UBB | 0.374009 | 4.6E-261 |
| CDKN3 | 0.370557 | 8.73E-47 |
| TMSB10 | 0.369309 | 0 |
| CALM3 | 0.36853 | 5.1E-174 |
| NDUFC1 | 0.366383 | 7.4E-157 |
| HDGF | 0.363688 | 2.5E-116 |
| DLGAP5 | 0.361694 | 1.93E-36 |
| GPX4 | 0.351092 | 2.8E-209 |
| RAN | 0.350667 | 2.6E-252 |
| HNRNPH3 | 0.339008 | 5E-122 |
| DYNLL1 | 0.335656 | 7.52E-65 |
| EIF5A | 0.334637 | 1.5E-147 |
| DCTN3 | 0.331033 | 6.2E-123 |
| LSM5 | 0.32787 | 2.39E-97 |
| PSMA4 | 0.32683 | 3.1E-137 |
| BANF1 | 0.325376 | 5E-190 |
| PRDX1 | 0.322636 | 3.1E-194 |
| TOP1 | 0.307681 | 3.18E-83 |
| TUBA1B | 0.304946 | 1.3E-198 |
| ZNF436-AS | 0.301363 | 2.01E-87 |
| MRFAP1 | 0.300451 | 8.9E-117 |
| SFPQ | 0.300318 | 1.76E-80 |

| | | |
|---|---|---|
| COX8A | 0.298675 | 7.5E-144 |
| MRPL51 | 0.294301 | 1.7E-124 |
| YWHAE | 0.293533 | 4.2E-203 |
| POMP | 0.293145 | 2.4E-129 |
| ILF2 | 0.292955 | 1E-111 |
| H2AFV | 0.29151 | 7E-125 |
| HNRNPK | 0.282376 | 1.7E-146 |
| ATP5MF | 0.281846 | 4.9E-167 |
| PSME2 | 0.281136 | 7.53E-90 |
| ANP32E | 0.280523 | 5.3E-58 |
| POLR2L | 0.278185 | 3.8E-114 |
| TPX2 | 0.273758 | 6.2E-17 |
| SLIRP | 0.271715 | 1.1E-106 |
| ATP5F1B | 0.271558 | 9E-127 |
| PSMB3 | 0.271064 | 2.4E-113 |
| LSM4 | 0.269957 | 4.2E-125 |
| NUF2 | 0.265638 | 6.27E-21 |
| SEM1 | 0.263245 | 5.9E-132 |
| BNIP3 | 0.262289 | 6.73E-49 |
| CSRP2 | 0.260897 | 4.1E-101 |
| ANAPC15 | 0.259943 | 4.48E-67 |
| TMSB4X | 0.259888 | 1.43E-85 |
| GNG5 | 0.258633 | 4.8E-152 |
| MORF4L2 | 0.258505 | 4.15E-50 |
| ILF3-DT | 0.258461 | 4.38E-59 |
| UBL5 | 0.257429 | 3.4E-157 |
| UQCR10 | 0.256868 | 2.6E-144 |
| DTYMK | 0.254165 | 8.25E-45 |
| AL118516. | 0.254013 | 4.09E-75 |
| FOXC2 | 0.252729 | 1.38E-39 |

| **Cluster 7** | | |
|---|---|---|
| Gene | LogFC | pVal |
| DKK1 | 2.106763 | 0 |
| TWIST1 | 1.536364 | 0 |
| NEFM | 1.483339 | 2.9E-230 |
| PRRX1 | 1.375432 | 0 |
| LGALS1 | 1.043799 | 1.3E-207 |
| VIM | 0.994352 | 0 |
| PRRX2 | 0.885141 | 0 |
| CALD1 | 0.878438 | 0 |
| CDC42EP5 | 0.852736 | 0 |
| THSD7A | 0.758156 | 9.9E-268 |
| EMP3 | 0.728631 | 0 |
| CLMP | 0.728317 | 4.4E-279 |
| COL3A1 | 0.69206 | 2.7E-248 |
| LIMCH1 | 0.685376 | 2.1E-271 |
| TAC1 | 0.67702 | 7.3E-104 |
| FRZB | 0.672259 | 2.1E-194 |
| MARCKS | 0.663053 | 0 |
| PCOLCE | 0.656755 | 0 |
| PDGFRA | 0.641996 | 4.6E-238 |
| BGN | 0.62559 | 8.2E-246 |
| TAGLN | 0.622221 | 6.49E-75 |
| VEGFD | 0.618828 | 8.1E-132 |
| MYL9 | 0.615729 | 3.4E-167 |
| FILIP1L | 0.612304 | 3.7E-135 |
| PCDH9 | 0.61197 | 3.6E-239 |
| TPM4 | 0.611108 | 3.3E-292 |
| MSX1 | 0.602925 | 8.9E-133 |
| COL1A2 | 0.602574 | 0 |
| A2M | 0.596003 | 2E-181 |
| GNG11 | 0.589801 | 5.8E-232 |
| CYR61 | 0.589121 | 2.2E-152 |
| COL6A2 | 0.585847 | 2.8E-255 |
| TCF12 | 0.584811 | 1.9E-239 |
| CLEC1A | 0.573592 | 2.3E-209 |
| LIMA1 | 0.56726 | 1.2E-223 |
| EVA1B | 0.560854 | 2.1E-273 |
| APCDD1 | 0.56055 | 1.9E-175 |
| RARRES2 | 0.552681 | 1.27E-77 |
| CTHRC1 | 0.542582 | 3.6E-117 |
| ACTB | 0.541941 | 0 |
| DNM3OS | 0.531335 | 5E-201 |
| PCDH18 | 0.524464 | 2.7E-192 |
| MEIS2 | 0.518489 | 4.3E-152 |
| ACAT2 | 0.516511 | 2.2E-172 |
| IL11RA | 0.505397 | 2.4E-199 |
| C12orf57 | 0.505392 | 1E-303 |
| FDPS | 0.497771 | 2E-216 |
| LIX1 | 0.496142 | 1.3E-121 |
| IGDCC3 | 0.488982 | 1.5E-230 |
| PTH1R | 0.482636 | 5.5E-205 |
| MFAP4 | 0.482611 | 5.6E-194 |
| CNN3 | 0.481458 | 6.2E-230 |
| MSMO1 | 0.479452 | 2.2E-152 |
| MAB21L2 | 0.473245 | 1.1E-131 |

| | | |
|---|---|---|
| TGFBI | 0.470139 | 1E-145 |
| NEFL | 0.464492 | 8.6E-102 |
| LGALS3 | 0.464419 | 3.1E-143 |
| IDI1 | 0.463451 | 7.3E-155 |
| CCND1 | 0.463288 | 1.7E-169 |
| FDFT1 | 0.46293 | 8.2E-215 |
| S100A11 | 0.458467 | 0 |
| BAALC | 0.449528 | 3.6E-151 |
| EMILIN1 | 0.446628 | 3.7E-189 |
| SERPINH1 | 0.44625 | 7.7E-198 |
| CNN2 | 0.444033 | 1.2E-195 |
| ID1 | 0.423182 | 3.7E-208 |
| PALLD | 0.420351 | 1.1E-152 |
| HMGCS1 | 0.418772 | 8.1E-138 |
| CTGF | 0.416967 | 1.15E-69 |
| OLFML3 | 0.410291 | 4E-158 |
| PAPSS1 | 0.400451 | 1.7E-140 |
| MSX2 | 0.398323 | 8.1E-119 |
| OAF | 0.39627 | 1.3E-171 |
| EDNRA | 0.395826 | 3.4E-139 |
| AES | 0.387462 | 2.2E-153 |
| TUBB6 | 0.387199 | 1.3E-119 |
| FTL | 0.379591 | 1.7E-104 |
| MAGED2 | 0.378304 | 1.6E-169 |
| JUND | 0.378229 | 6.7E-138 |
| VCL | 0.376895 | 6.8E-119 |
| SERPINF1 | 0.37561 | 3.4E-131 |
| SQLE | 0.373726 | 1.7E-138 |
| TPM2 | 0.372202 | 4.4E-164 |
| GLT8D2 | 0.366951 | 1.8E-152 |
| LDB2 | 0.366265 | 1.8E-147 |
| PMAIP1 | 0.358515 | 6.05E-95 |
| TNC | 0.347508 | 5.29E-94 |
| AFF3 | 0.345965 | 2.7E-142 |
| ITGA4 | 0.341505 | 3E-130 |
| DACT3 | 0.338968 | 1.1E-146 |
| HMGA2 | 0.332597 | 2.9E-145 |
| FHL3 | 0.332304 | 7.8E-126 |
| LMO4 | 0.331863 | 1.1E-103 |
| FBLN1 | 0.3313 | 1.6E-155 |
| COL5A2 | 0.329052 | 2E-122 |
| MDFI | 0.32354 | 1.7E-141 |
| COL1A1 | 0.322324 | 3.87E-84 |
| DDR2 | 0.321768 | 1.7E-137 |
| MAP1LC3A | 0.319271 | 2.4E-124 |
| MFAP2 | 0.315922 | 1.1E-132 |
| COL25A1 | 0.314612 | 6.4E-136 |
| EPB41L2 | 0.311347 | 1.1E-104 |
| HIST1H4C | 0.307356 | 2.54E-16 |
| NME4 | 0.305899 | 3.1E-159 |
| ZFHX4 | 0.304253 | 1.5E-120 |
| EMCN | 0.3042 | 2.08E-97 |
| HOXA10 | 0.303544 | 7.4E-102 |
| CPE | 0.299077 | 2.5E-96 |
| LRRC17 | 0.296759 | 3.1E-101 |
| THY1 | 0.295372 | 2.04E-91 |

| | | |
|---|---|---|
| ISG15 | 0.293581 | 5.7E-108 |
| CRTAP | 0.291359 | 9.1E-114 |
| CXCL14 | 0.290094 | 4.34E-35 |
| GNB4 | 0.289363 | 3.9E-108 |
| MECOM | 0.289115 | 4.4E-113 |
| MME | 0.286996 | 9.2E-102 |
| ANXA6 | 0.28667 | 1.5E-115 |
| POSTN | 0.281888 | 4.73E-84 |
| HNRNPH3 | 0.281596 | 4.9E-111 |
| FAM89A | 0.281464 | 9.2E-103 |
| B2M | 0.2805 | 3.7E-139 |
| MAP1B | 0.280084 | 5.48E-81 |
| VCAM1 | 0.278042 | 3.7E-101 |
| CPED1 | 0.277633 | 3.6E-114 |
| METTL9 | 0.274481 | 7.6E-130 |
| COL6A1 | 0.270398 | 1.09E-92 |
| PTN | 0.2703 | 6.6E-46 |
| NFIB | 0.269539 | 3.6E-119 |
| BASP1 | 0.268193 | 5.9E-94 |
| DDAH2 | 0.267082 | 2.9E-107 |
| NDUFA4 | 0.267027 | 1.5E-196 |
| SELENOM | 0.265577 | 2.7E-90 |
| RAP1B | 0.262861 | 1.77E-82 |
| SGK1 | 0.260245 | 3.42E-76 |
| EML4 | 0.260146 | 2.27E-88 |
| CYTL1 | 0.259985 | 6.83E-66 |
| ZEB1 | 0.259544 | 5.9E-108 |
| FLRT2 | 0.258834 | 8.6E-99 |
| FANCL | 0.256818 | 5.01E-91 |
| MYL12A | 0.256682 | 3.32E-95 |
| ZNF106 | 0.255428 | 4.84E-87 |
| GAS2 | 0.254981 | 1.77E-89 |
| MYL12B | 0.253458 | 1.75E-89 |
| JUN | 0.25138 | 8.34E-43 |
| SNAI2 | 0.251336 | 5.47E-54 |
| AC027031. | 0.250614 | 1.76E-79 |

| Cluster 8 | | |
|---|---|---|
| Gene | LogFC | pVal |
| UBE2C | 1.816757 | 0 |
| TUBB4B | 1.587338 | 0 |
| KPNA2 | 1.581853 | 0 |
| CENPF | 1.581666 | 0 |
| TUBA1C | 1.571858 | 0 |
| CCNB1 | 1.489109 | 0 |
| UBE2S | 1.465092 | 0 |
| TOP2A | 1.453513 | 0 |
| CDC20 | 1.350906 | 0 |
| PTTG1 | 1.32565 | 0 |
| CDKN3 | 1.258121 | 0 |
| CDK1 | 1.257747 | 0 |
| HMGB2 | 1.246891 | 0 |
| ARL6IP1 | 1.235075 | 0 |
| NUSAP1 | 1.199956 | 0 |
| ASPM | 1.1756 | 4.3E-285 |
| CCNB2 | 1.147924 | 0 |
| PLK1 | 1.125382 | 1E-304 |
| CENPA | 1.106829 | 0 |
| CKS2 | 1.101516 | 0 |
| BIRC5 | 1.065604 | 0 |
| DLGAP5 | 1.05673 | 0 |
| MKI67 | 1.041235 | 7.2E-260 |
| AURKA | 1.040293 | 1.4E-275 |
| TPX2 | 1.03596 | 2.2E-282 |
| CENPE | 1.033396 | 2.8E-263 |
| CKS1B | 1.01728 | 0 |
| SGO2 | 1.010206 | 5E-307 |
| CDCA3 | 1.007243 | 8.4E-271 |
| HMMR | 0.965771 | 1.1E-246 |
| NEK2 | 0.930168 | 2E-230 |
| AURKB | 0.927473 | 2E-296 |
| NUF2 | 0.925681 | 5.8E-277 |
| PIMREG | 0.903781 | 1.8E-259 |
| JPT1 | 0.891665 | 0 |
| GTSE1 | 0.876928 | 1.6E-268 |
| CALM2 | 0.862826 | 0 |
| CCNA2 | 0.833354 | 4E-233 |
| KIF20B | 0.82917 | 1.3E-242 |
| TUBA1B | 0.821851 | 0 |
| KNSTRN | 0.820663 | 2.1E-253 |
| H2AFX | 0.820302 | 1.2E-255 |
| PSRC1 | 0.812127 | 2.9E-248 |
| TROAP | 0.790452 | 9.5E-229 |
| CDCA8 | 0.788145 | 1.4E-218 |
| BUB3 | 0.756033 | 2.6E-273 |
| MAD2L1 | 0.732264 | 0 |
| TACC3 | 0.724522 | 1E-224 |
| NDC80 | 0.722694 | 5.8E-235 |
| KIFC1 | 0.721893 | 3.5E-252 |
| PBK | 0.703744 | 1.2E-217 |
| HMGB3 | 0.698142 | 3.5E-254 |
| SMC4 | 0.694349 | 5.6E-253 |
| CKAP2 | 0.690425 | 2.9E-189 |

| | | |
|---|---|---|
| DEPDC1 | 0.678591 | 5.1E-189 |
| KIF2C | 0.659272 | 4.4E-191 |
| TTK | 0.653976 | 7E-200 |
| KIF20A | 0.650448 | 1.6E-185 |
| BUB1 | 0.635186 | 1E-174 |
| PIF1 | 0.630853 | 1.2E-169 |
| SGO1 | 0.629629 | 2.5E-206 |
| KIF23 | 0.617034 | 4.3E-175 |
| DTYMK | 0.615211 | 3.6E-241 |
| KIF22 | 0.610762 | 7.1E-222 |
| MZT1 | 0.607223 | 4.8E-199 |
| MIS18BP1 | 0.590164 | 7.9E-183 |
| CENPW | 0.584906 | 1.9E-241 |
| CKAP2L | 0.579959 | 2E-176 |
| SKA2 | 0.56937 | 2.8E-232 |
| KIF4A | 0.562272 | 2.6E-160 |
| MXD3 | 0.562031 | 8.2E-146 |
| NUCKS1 | 0.557582 | 2.1E-240 |
| ECT2 | 0.556487 | 2.2E-157 |
| H1FX | 0.553612 | 9.9E-253 |
| H2AFZ | 0.544392 | 0 |
| DBF4 | 0.538435 | 4.9E-149 |
| UBE2T | 0.533459 | 1E-212 |
| NCAPG | 0.523504 | 1.6E-153 |
| HMGN2 | 0.51313 | 5.3E-274 |
| HJURP | 0.511665 | 1.4E-133 |
| G2E3 | 0.49797 | 4E-147 |
| DYNLL1 | 0.497621 | 1.1E-207 |
| CDCA2 | 0.487615 | 2.7E-131 |
| KNL1 | 0.48663 | 6.9E-130 |
| STMN1 | 0.485827 | 0 |
| CDC25C | 0.482306 | 2.2E-138 |
| HMGB1 | 0.481241 | 0 |
| KIF5B | 0.480755 | 1.1E-185 |
| KIF18A | 0.479441 | 2.1E-132 |
| SPC25 | 0.47873 | 9.3E-158 |
| FAM83D | 0.478081 | 7.8E-140 |
| CEP55 | 0.474915 | 9.9E-123 |
| UBALD2 | 0.473476 | 4.1E-124 |
| KIF14 | 0.46421 | 5.7E-122 |
| TUBB | 0.460073 | 0 |
| DEPDC1B | 0.458609 | 1.9E-135 |
| H2AFV | 0.453599 | 1.5E-211 |
| ARHGEF3[illegible] | 0.450447 | 2.9E-131 |
| DCTN3 | 0.446898 | 3.7E-166 |
| NCAPD2 | 0.437433 | 5E-140 |
| LSM5 | 0.436783 | 2.5E-175 |
| CKAP5 | 0.435611 | 4.5E-129 |
| PRC1 | 0.425765 | 1.4E-137 |
| RHEB | 0.420904 | 5.6E-156 |
| RAN | 0.418273 | 7.8E-281 |
| COX17 | 0.418056 | 8.9E-159 |
| RAD21 | 0.416459 | 3.3E-124 |
| RACGAP1 | 0.416174 | 1.3E-136 |
| NUDCD2 | 0.415568 | 8.8E-126 |
| SPDL1 | 0.411127 | 1.2E-117 |

| | | |
|---|---|---|
| CNIH4 | 0.408343 | 3.3E-138 |
| CENPN | 0.408278 | 1.1E-140 |
| HNRNPA2 | 0.40424 | 2.2E-186 |
| PLIN3 | 0.4023 | 2.5E-144 |
| GAS2L3 | 0.397106 | 8.7E-109 |
| REEP4 | 0.395275 | 8.4E-110 |
| LBR | 0.395062 | 3.4E-110 |
| RNF26 | 0.39168 | 6E-117 |
| OIP5 | 0.388481 | 2.2E-131 |
| HP1BP3 | 0.385645 | 4.2E-100 |
| TUBA1A | 0.378047 | 1.8E-118 |
| KIF11 | 0.376759 | 1.12E-92 |
| DDX39A | 0.372916 | 3.5E-110 |
| EMC9 | 0.371483 | 5.2E-112 |
| MORF4L2 | 0.369155 | 1.3E-141 |
| CEP70 | 0.36656 | 1.1E-107 |
| UBB | 0.363685 | 3.9E-190 |
| GPSM2 | 0.362775 | 2.47E-98 |
| VDAC3 | 0.35582 | 3.9E-127 |
| PARPBP | 0.354473 | 3.9E-109 |
| CCNF | 0.353421 | 2.4E-102 |
| CMC2 | 0.353216 | 1.4E-116 |
| BORA | 0.352451 | 1.1E-112 |
| LMNB1 | 0.348259 | 2.07E-88 |
| MRPL51 | 0.34582 | 1.1E-174 |
| DCAF7 | 0.342221 | 6.2E-122 |
| KMT5A | 0.340469 | 3.98E-84 |
| ILF2 | 0.337982 | 3.7E-156 |
| CALM3 | 0.337715 | 7E-130 |
| RNF5 | 0.336376 | 3.5E-109 |
| CD9 | 0.335077 | 4.16E-52 |
| BCL2L12 | 0.333556 | 1.1E-112 |
| CCDC18 | 0.332909 | 1.92E-95 |
| FBXO5 | 0.332762 | 3.22E-82 |
| HSP90AA1 | 0.33185 | 4.6E-126 |
| INCENP | 0.327573 | 7.57E-92 |
| ANP32E | 0.327132 | 6.82E-95 |
| HSP90B1 | 0.32672 | 3.1E-103 |
| RUVBL2 | 0.324593 | 1.1E-104 |
| HMG20B | 0.323781 | 1.03E-86 |
| BANF1 | 0.323543 | 4.9E-168 |
| CYCS | 0.323381 | 1E-104 |
| BUB1B | 0.321903 | 3.98E-85 |
| CCT5 | 0.320718 | 4.7E-119 |
| ARHGAP1 | 0.317609 | 1.36E-88 |
| MYEF2 | 0.316495 | 1.32E-80 |
| PHF19 | 0.31552 | 1.38E-85 |
| BRD8 | 0.313634 | 1.37E-77 |
| PSIP1 | 0.312329 | 1E-112 |
| GLRX5 | 0.311787 | 4.3E-119 |
| CDC27 | 0.309294 | 1.58E-85 |
| PPP1R35 | 0.308261 | 2.21E-84 |
| PTMS | 0.307899 | 3.1E-120 |
| MDH1 | 0.304377 | 4.2E-120 |
| ACTB | 0.300714 | 1.4E-159 |
| SPA17 | 0.299637 | 9.41E-83 |

| | | |
|---|---|---|
| SCLT1 | 0.294754 | 2.07E-74 |
| ASF1B | 0.294628 | 2.69E-86 |
| GOT1 | 0.29313 | 3.15E-84 |
| SAPCD2 | 0.290482 | 4.23E-84 |
| RTKN2 | 0.287627 | 4.59E-81 |
| SPAG5 | 0.284211 | 1.21E-78 |
| EEF1AKMT | 0.284062 | 1.02E-79 |
| PLGRKT | 0.278793 | 9.93E-83 |
| COX8A | 0.278104 | 4.9E-129 |
| NCAPH | 0.275841 | 1.36E-80 |
| NDE1 | 0.275056 | 1.77E-79 |
| ERH | 0.274498 | 3.3E-180 |
| GRK6 | 0.271901 | 1.98E-89 |
| ISCA2 | 0.271316 | 1.7E-103 |
| HSPA1A | 0.271031 | 1.28E-56 |
| FAM110A | 0.268758 | 8.8E-80 |
| CCDC34 | 0.267793 | 2.06E-82 |
| GTF2A2 | 0.26592 | 2.29E-98 |
| TNFAIP8L | 0.264883 | 8.8E-79 |
| RBM8A | 0.264552 | 1.3E-106 |
| LDHA | 0.264396 | 8.43E-70 |
| KIF15 | 0.260549 | 3.34E-71 |
| CHEK2 | 0.258774 | 1.81E-79 |
| IMMP1L | 0.256781 | 1.05E-83 |
| PRPSAP1 | 0.256196 | 2.56E-69 |
| TRIOBP | 0.255916 | 8.22E-73 |
| FOXM1 | 0.254855 | 3.78E-68 |
| GGH | 0.254669 | 2.25E-74 |
| ATP5F1B | 0.253457 | 3.2E-108 |
| SMC2 | 0.253251 | 2.26E-83 |
| UBC | 0.251779 | 1.13E-78 |

| **Cluster 9** | | |
|---|---|---|
| Gene | LogFC | pVal |
| MT1X | 2.416159 | 5.6E-215 |
| MT2A | 2.199162 | 4.2E-136 |
| MT1E | 1.718452 | 1.95E-42 |
| MT1G | 1.200161 | 1.76E-07 |
| VIM | 0.344877 | 9.34E-46 |
| PCOLCE | 0.334028 | 1.23E-31 |
| NPW | 0.282695 | 1.5E-19 |
| RGS5 | 0.271106 | 1.36E-15 |
| HAPLN1 | 0.264555 | 5.72E-14 |
| S100A10 | 0.258815 | 4.44E-16 |

| **Cluster 10** | | |
|---|---|---|
| Gene | LogFC | pVal |
| HIST1H4C | 0.762728 | 1.75E-60 |
| CENPF | 0.684526 | 3.72E-45 |
| MKI67 | 0.650163 | 1.72E-37 |
| CTNNB1 | 0.626874 | 3.92E-68 |
| TOP2A | 0.622797 | 2.15E-38 |
| LRRC75A | 0.609053 | 2.33E-62 |
| AC092069. | 0.578063 | 1.66E-55 |
| MT-ND4 | 0.555762 | 4.52E-54 |
| HNRNPH1 | 0.548354 | 2.27E-64 |
| SMC4 | 0.542207 | 5.87E-43 |
| PABPC1 | 0.505489 | 5.77E-61 |
| EGR1 | 0.488534 | 1.97E-25 |
| ZFP36L1 | 0.481064 | 1.82E-45 |
| TTC3 | 0.465063 | 7.53E-45 |
| CENPE | 0.4612 | 6.19E-22 |
| MTRNR2L | 0.460304 | 1.83E-26 |
| EIF2S3 | 0.456899 | 1.47E-39 |
| MT-ND2 | 0.455069 | 1.92E-43 |
| GOLGA4 | 0.449115 | 4.17E-39 |
| C1orf56 | 0.443134 | 7.59E-41 |
| SOX11 | 0.441962 | 9.63E-31 |
| KPNB1 | 0.426758 | 1.35E-38 |
| TPX2 | 0.417161 | 8.14E-21 |
| MT-ND1 | 0.413145 | 1.13E-38 |
| NUCKS1 | 0.403646 | 8.45E-43 |
| ASPM | 0.391104 | 3.44E-17 |
| BIRC5 | 0.38079 | 1.64E-21 |
| HMGA2 | 0.379229 | 2.35E-27 |
| TCAF1 | 0.375503 | 1.77E-27 |
| DLGAP5 | 0.375499 | 1.74E-20 |
| NIPBL | 0.374742 | 1.54E-24 |
| KTN1 | 0.365812 | 4.13E-30 |
| SETD5 | 0.364087 | 1.15E-28 |
| IRF2BP2 | 0.360607 | 3.34E-22 |
| GTF2I | 0.360147 | 6.22E-21 |
| SYNE2 | 0.350183 | 3.3E-16 |
| RAD21 | 0.35015 | 1.86E-19 |
| MAN1A2 | 0.349547 | 1.15E-26 |
| SET | 0.347019 | 4.99E-39 |
| UBE2C | 0.346359 | 1.63E-16 |
| SLIT2 | 0.343526 | 8.14E-21 |
| KIF5B | 0.340948 | 5.73E-22 |
| BPTF | 0.335823 | 7.56E-19 |
| NEAT1 | 0.333283 | 3.88E-17 |
| MT-ND5 | 0.331922 | 2.96E-31 |
| KIF20B | 0.328609 | 4.57E-17 |
| EIF4EBP2 | 0.326984 | 5.49E-17 |
| PAX2 | 0.32551 | 1.04E-15 |
| ZNF462 | 0.32526 | 4.23E-15 |
| ARID3A | 0.325014 | 8.63E-24 |
| RDH10 | 0.323413 | 4.54E-15 |
| OSBPL8 | 0.321791 | 4.87E-20 |
| TNRC6B | 0.320263 | 2.64E-17 |
| MEX3A | 0.320208 | 1.59E-18 |

| | | |
|---|---|---|
| ROCK2 | 0.320145 | 9.19E-17 |
| NR2F2 | 0.317036 | 2.62E-19 |
| GTSE1 | 0.316222 | 1.16E-15 |
| ROCK1 | 0.31496 | 2.53E-19 |
| CEP350 | 0.314766 | 5.09E-24 |
| MT-CO2 | 0.31252 | 6.19E-31 |
| ENAH | 0.308476 | 2.47E-16 |
| SGO2 | 0.308147 | 5.57E-12 |
| HIST1H1D | 0.30723 | 1.2E-10 |
| PPP1CB | 0.304076 | 3.55E-13 |
| CTBP1 | 0.304052 | 1.47E-20 |
| CCDC88A | 0.303876 | 1.39E-18 |
| ARID1A | 0.30192 | 6.15E-17 |
| B4GALT1 | 0.301113 | 1.17E-18 |
| TP53 | 0.300574 | 2.98E-21 |
| CKAP5 | 0.299714 | 3.27E-09 |
| ARL4C | 0.297498 | 1.46E-12 |
| IGDCC3 | 0.297214 | 1.26E-21 |
| MYBL2 | 0.295237 | 4.35E-15 |
| USP1 | 0.294244 | 6.48E-14 |
| MDM4 | 0.293403 | 4.77E-21 |
| BAZ1B | 0.293072 | 1.02E-13 |
| ANKRD11 | 0.290188 | 1.47E-14 |
| MIS18BP1 | 0.287472 | 1.2E-11 |
| UBE2S | 0.286819 | 1.07E-15 |
| ARL6IP1 | 0.286386 | 2.32E-11 |
| CELF1 | 0.286318 | 5.39E-16 |
| HNRNPUL | 0.28416 | 1.41E-13 |
| BRD4 | 0.283319 | 7.32E-17 |
| ADD3 | 0.283286 | 8.41E-17 |
| HIST1H1A | 0.283146 | 4.42E-10 |
| FANCD2 | 0.281708 | 1.7E-16 |
| EIF4G2 | 0.277658 | 4.72E-13 |
| NCAPD2 | 0.276935 | 6.65E-13 |
| CBX5 | 0.276202 | 2.69E-14 |
| FADS1 | 0.276079 | 3.01E-12 |
| FKBP9 | 0.275619 | 9.41E-18 |
| HOXC8 | 0.273601 | 3.83E-14 |
| SMC1A | 0.273567 | 7.39E-13 |
| PTP4A2 | 0.273196 | 6.83E-13 |
| CSDE1 | 0.271189 | 8.41E-11 |
| PLK1 | 0.269028 | 5.81E-08 |
| CAPZA1 | 0.268899 | 2.36E-13 |
| ATRX | 0.268655 | 6.44E-12 |
| BDP1 | 0.268398 | 3.64E-12 |
| ZNF292 | 0.268001 | 2.26E-13 |
| DACH1 | 0.266464 | 1.27E-11 |
| ZKSCAN1 | 0.266415 | 6.19E-14 |
| CDC42SE1 | 0.266375 | 4.7E-17 |
| TCF4 | 0.266354 | 1.56E-13 |
| PCM1 | 0.266119 | 1.42E-13 |
| SLC39A6 | 0.265484 | 7.9E-17 |
| SLK | 0.265289 | 2.02E-14 |
| ERGIC3 | 0.263749 | 1.56E-14 |
| CCNI | 0.263446 | 1.47E-15 |
| HDGFL3 | 0.263237 | 3.6E-13 |

| | | |
|---|---|---|
| MYH10 | 0.262886 | 0.000388 |
| NPEPPS | 0.261179 | 3.49E-08 |
| KIF23 | 0.26041 | 1.64E-10 |
| TRA2A | 0.259413 | 5.54E-13 |
| PXDN | 0.259367 | 6.25E-11 |
| DHFR | 0.259077 | 1.3E-10 |
| HOXC10 | 0.257857 | 7.56E-14 |
| HDAC2 | 0.257718 | 1.15E-10 |
| IGF2BP2 | 0.25768 | 7.51E-11 |
| ARHGAP5 | 0.256431 | 5.34E-11 |
| KIF14 | 0.25609 | 1.76E-10 |
| TOR1AIP2 | 0.255487 | 5.11E-15 |
| KHSRP | 0.255286 | 8.69E-10 |
| MYO10 | 0.255142 | 8.29E-10 |
| MT-CO3 | 0.254371 | 4.4E-22 |
| CEP78 | 0.253394 | 2.15E-12 |
| BRD3 | 0.253382 | 3.98E-13 |
| RB1CC1 | 0.252176 | 6.68E-13 |
| IPO5 | 0.251469 | 4.31E-07 |
| KMT2E | 0.251125 | 7.94E-10 |
| PCBP2 | 0.251066 | 5.27E-23 |
| HP1BP3 | 0.250445 | 4.87E-11 |
| PPP2R1A | 0.250157 | 6.39E-12 |

| **Cluster 11** | | |
|---|---|---|
| Gene | LogFC | pVal |
| EMX2 | 1.272424 | 4.1E-147 |
| SCT | 0.909476 | 5.17E-70 |
| BST2 | 0.887818 | 2.21E-87 |
| TSPAN1 | 0.848842 | 1.17E-90 |
| RAB11FIP1 | 0.843514 | 2.98E-87 |
| CLDN11 | 0.819234 | 6.86E-72 |
| TUBB2B | 0.770841 | 4.8E-97 |
| SPINT2 | 0.691339 | 4.9E-122 |
| SH3YL1 | 0.690863 | 5.79E-95 |
| PRICKLE1 | 0.671685 | 1.14E-79 |
| ADGRL4 | 0.665302 | 3.07E-66 |
| YBX3 | 0.644738 | 3.18E-85 |
| EMX2OS | 0.605115 | 2.56E-59 |
| LHX1 | 0.603335 | 1.59E-54 |
| SERPINE2 | 0.590396 | 8.06E-91 |
| IGFBP3 | 0.58705 | 3.94E-21 |
| TFPI2 | 0.558955 | 2.08E-25 |
| BASP1 | 0.548039 | 1.01E-74 |
| SEMA6A | 0.532258 | 2.55E-65 |
| SLC8A1 | 0.526535 | 8.83E-52 |
| DKK1 | 0.524921 | 3.61E-31 |
| ID4 | 0.500677 | 4.71E-73 |
| CAMK2N1 | 0.488875 | 3.36E-39 |
| FLI1 | 0.48643 | 3.34E-61 |
| SNHG8 | 0.483229 | 1.06E-70 |
| VAMP8 | 0.47825 | 2.33E-84 |
| CADM1 | 0.474276 | 1.45E-59 |
| IGFBP5 | 0.473133 | 8.58E-32 |
| PAX2 | 0.469795 | 6.97E-89 |
| LINC0230[illegible] | 0.462547 | 2.38E-22 |
| GFRA1 | 0.453848 | 9.06E-54 |
| KRT18 | 0.450584 | 9.86E-81 |
| LYPD6B | 0.443287 | 3.47E-62 |
| SCX | 0.441023 | 5.73E-44 |
| APOE | 0.440424 | 9.06E-66 |
| BCAM | 0.438318 | 3.47E-59 |
| IGFBP7 | 0.427238 | 8.35E-37 |
| TINAGL1 | 0.402624 | 1.08E-54 |
| ATP1B1 | 0.402495 | 2.4E-45 |
| APCDD1 | 0.401359 | 3.74E-36 |
| SUCO | 0.398469 | 1.74E-48 |
| CD47 | 0.392481 | 3.42E-38 |
| HOXD1 | 0.392135 | 2.72E-45 |
| S100A6 | 0.391664 | 4.34E-28 |
| KHDRBS3 | 0.385969 | 2.92E-34 |
| DMKN | 0.381129 | 2.15E-52 |
| SOD3 | 0.381118 | 2.43E-34 |
| LY6E | 0.380806 | 2.87E-57 |
| AIG1 | 0.37779 | 5.96E-51 |
| ALDH2 | 0.368441 | 5.67E-38 |
| TPD52L1 | 0.367994 | 3.24E-44 |
| S100A9 | 0.365549 | 1.64E-23 |
| BMP7 | 0.362134 | 1.46E-45 |
| ABCG2 | 0.361188 | 7.19E-26 |

| | | |
|---|---|---|
| PRCP | 0.360612 | 4.79E-30 |
| EGFL7 | 0.357641 | 2.79E-49 |
| SCUBE3 | 0.357406 | 1.78E-37 |
| PCDH17 | 0.356667 | 9.41E-33 |
| NFE2L3 | 0.355591 | 2.56E-39 |
| TM7SF2 | 0.35559 | 1.47E-45 |
| AP002884. | 0.354373 | 5.58E-44 |
| OLFM3 | 0.348594 | 3.09E-40 |
| CHCHD10 | 0.346636 | 2.21E-47 |
| CDKN1A | 0.339298 | 9.79E-40 |
| C2CD4B | 0.338383 | 1.44E-36 |
| EPCAM | 0.33403 | 1.38E-32 |
| GADD45B | 0.332674 | 1.76E-30 |
| SPINT1 | 0.330626 | 2.49E-49 |
| COL4A2 | 0.329489 | 1.38E-45 |
| SAT1 | 0.324531 | 1.23E-30 |
| LYPD1 | 0.324482 | 9.94E-29 |
| S100A13 | 0.322842 | 9.43E-52 |
| S100A4 | 0.32151 | 5.1E-15 |
| AVP | 0.320866 | 9.39E-36 |
| UCP2 | 0.319917 | 1.42E-48 |
| UCHL1 | 0.319759 | 1.03E-35 |
| DBI | 0.319358 | 1.41E-29 |
| IL4I1 | 0.319067 | 9.03E-51 |
| KTN1 | 0.31633 | 2.1E-53 |
| CCDC85B | 0.314583 | 2.35E-49 |
| CTHRC1 | 0.314495 | 4.02E-23 |
| PLS1 | 0.311165 | 3.1E-35 |
| SOX4 | 0.307567 | 7.93E-44 |
| NEBL | 0.306143 | 1.78E-40 |
| TFAP2A | 0.301319 | 1.81E-35 |
| NPDC1 | 0.30093 | 1.34E-43 |
| PDLIM1 | 0.300429 | 1.46E-25 |
| KCNE5 | 0.29894 | 2.19E-11 |
| CLDN7 | 0.298792 | 1.53E-43 |
| RBM38 | 0.297738 | 4.13E-26 |
| CD9 | 0.296247 | 4.94E-27 |
| CLINT1 | 0.291283 | 8.01E-35 |
| GYPC | 0.28554 | 6.46E-28 |
| CLDN23 | 0.284766 | 8.96E-30 |
| RARRES2 | 0.284251 | 0.000789 |
| NFKBIA | 0.282082 | 1.91E-32 |
| CSTB | 0.281786 | 1.78E-47 |
| ATP10D | 0.28098 | 1.58E-37 |
| COL4A1 | 0.280382 | 3.89E-36 |
| PURPL | 0.280304 | 1.67E-18 |
| ACSL4 | 0.279542 | 5.32E-30 |
| KIAA1217 | 0.277073 | 3.77E-34 |
| DUSP1 | 0.276117 | 1.59E-29 |
| FAM181B | 0.276072 | 7.87E-30 |
| SKIL | 0.276066 | 6.57E-31 |
| SLC25A5 | 0.276053 | 4.99E-57 |
| KLK6 | 0.275299 | 3.84E-27 |
| SPON1 | 0.273989 | 2.69E-31 |
| MMD | 0.272564 | 2.63E-31 |
| MAP1B | 0.270618 | 1.7E-14 |

| | | |
|---|---|---|
| LINC01116 | 0.270102 | 2.88E-32 |
| THBS1 | 0.269701 | 8.78E-24 |
| TPBG | 0.269452 | 5.64E-28 |
| LITAF | 0.269131 | 5.75E-43 |
| GNG11 | 0.266628 | 3.87E-19 |
| BIN1 | 0.266494 | 4.08E-39 |
| AP1M2 | 0.263823 | 3.75E-37 |
| MBNL2 | 0.262494 | 5.37E-26 |
| TMEM59L | 0.26046 | 3.11E-36 |
| PRDX6 | 0.25729 | 4.37E-33 |
| AP1S2 | 0.25542 | 1.94E-24 |
| RAB25 | 0.254856 | 9.45E-35 |
| TMC6 | 0.253594 | 7.51E-37 |
| EVA1B | 0.2527 | 3.25E-35 |
| PRPH | 0.252612 | 5E-16 |
| CYB5A | 0.250968 | 8.07E-37 |

| Cluster 12 | | |
|---|---|---|
| Gene | LogFC | pVal |
| DLK1 | 1.78169 | 7.9E-204 |
| CDH6 | 0.292898 | 5.53E-16 |
| PAX8 | 0.26219 | 5.31E-25 |

| **Cluster 13** | | |
|---|---|---|
| Gene | LogFC | pVal |
| XIST | 1.427671 | 1.15E-09 |
| MT-ND5 | 1.380745 | 6.2E-103 |
| MT-ND6 | 1.379577 | 5.59E-60 |
| MT-ATP6 | 1.375351 | 6.4E-115 |
| MT-ND1 | 1.318434 | 9.3E-100 |
| MT-CO3 | 1.314127 | 3.5E-117 |
| MT-CO1 | 1.311654 | 1E-115 |
| MT-CO2 | 1.260873 | 7.4E-116 |
| MT-CYB | 1.228742 | 2.6E-118 |
| MT-ND4 | 1.227463 | 3.4E-113 |
| MT-ND3 | 1.203451 | 1.2E-110 |
| MTRNR2L | 1.070695 | 1.99E-53 |
| MT-ND2 | 1.062846 | 8.71E-90 |
| NEAT1 | 0.971509 | 4.67E-28 |
| MT-ND4L | 0.943187 | 1.77E-47 |
| KCNQ1OT | 0.90727 | 5.86E-27 |
| WSB1 | 0.88314 | 2.91E-23 |
| MALAT1 | 0.807787 | 1.32E-39 |
| NKTR | 0.777621 | 2.27E-20 |
| DDX17 | 0.691286 | 5.72E-24 |
| GABPB1-A | 0.646313 | 2.54E-08 |
| MT-ATP8 | 0.632091 | 5.08E-15 |
| ARID1B | 0.603822 | 2.54E-13 |
| SFPQ | 0.600114 | 2.29E-08 |
| SLC38A2 | 0.582484 | 4.62E-12 |
| FTX | 0.579303 | 5.25E-11 |
| POLR2J3.1 | 0.573455 | 9.94E-14 |
| PLCG2 | 0.572161 | 0.002336 |
| VCAN | 0.570708 | 6.9E-08 |
| MTRNR2L8 | 0.569475 | 6.96E-09 |
| HMGA2 | 0.547539 | 1.32E-05 |
| LUC7L3 | 0.541712 | 2.03E-12 |
| RBM5 | 0.538599 | 2.93E-06 |
| OGA | 0.529305 | 2.12E-07 |
| PLXNB2 | 0.527806 | 7.48E-09 |
| ZMYM2 | 0.508651 | 1.85E-07 |
| HELLS | 0.501709 | 0.000112 |
| PRTG | 0.49581 | 9.23E-08 |
| FUS | 0.495763 | 5.67E-07 |
| CHD9 | 0.492224 | 0.000199 |
| HOXD9 | 0.491922 | 4.06E-06 |
| ADCY2 | 0.489705 | 4.13E-06 |
| PLOD1 | 0.489008 | 3.51E-05 |
| HNRNPU | 0.484328 | 2.42E-08 |
| ARGLU1 | 0.472876 | 8.69E-06 |
| MDM4 | 0.472561 | 2.02E-07 |
| PRRC2A | 0.46874 | 1.74E-07 |
| MSI2 | 0.465341 | 6.06E-07 |
| ZNF292 | 0.463879 | 7.41E-05 |
| ZNF117 | 0.462312 | 6.07E-08 |
| CCND2 | 0.461952 | 1.91E-05 |
| COL1A1 | 0.461248 | 0.005301 |
| CCNL2 | 0.457508 | 1.64E-06 |
| AGRN | 0.455328 | 1.25E-09 |

| | | |
|---|---|---|
| CCDC14 | 0.453921 | 3.06E-07 |
| SMG1 | 0.453626 | 3.73E-07 |
| GOLGA4 | 0.447409 | 0.000171 |
| HOXC8 | 0.447212 | 0.001781 |
| TSHZ2 | 0.443496 | 0.003111 |
| PCSK5 | 0.435856 | 0.0005 |
| KANSL1 | 0.434866 | 6.14E-06 |
| FBN2 | 0.429164 | 1.66E-05 |
| CEP78 | 0.425947 | 0.005029 |
| MRC2 | 0.425249 | 0.000188 |
| CLSTN1 | 0.422916 | 1.99E-05 |
| GOLGA8A | 0.41628 | 2.42E-06 |
| FRMD4A | 0.415205 | 0.021171 |
| IGF2BP1 | 0.408645 | 0.001515 |
| PCNX4 | 0.405615 | 0.000139 |
| N4BP2L2 | 0.405581 | 8.99E-05 |
| SYNE2 | 0.405195 | 0.000857 |
| PLPPR3 | 0.403519 | 0.013934 |
| DNM1 | 0.401331 | 0.000187 |
| RSRP1 | 0.394761 | 6.25E-05 |
| WDR6 | 0.394517 | 0.000542 |
| ATRX | 0.394498 | 0.005279 |
| DYNC1H1 | 0.394362 | 0.005119 |
| PRRC2B | 0.390697 | 7.2E-06 |
| INTS6 | 0.389695 | 0.003815 |
| TNRC6B | 0.388702 | 0.026268 |
| PNISR | 0.385929 | 2.53E-08 |
| PHIP | 0.384751 | 0.00298 |
| GTF2I | 0.384253 | 0.015857 |
| ZNF638 | 0.38385 | 0.024978 |
| SUGP2 | 0.383126 | 0.00394 |
| LENG8 | 0.377397 | 9.31E-06 |
| PXDN | 0.377185 | 0.000438 |
| AFF4 | 0.37524 | 0.03126 |
| LPP | 0.373739 | 0.001362 |
| ATM | 0.370343 | 0.001056 |
| ARFGAP1 | 0.36906 | 0.000253 |
| MKLN1 | 0.368094 | 0.000576 |
| BAZ2B | 0.368009 | 0.009874 |
| SACS | 0.361084 | 1.75E-05 |
| UBR5 | 0.360751 | 0.001519 |
| NFATC2IP | 0.358737 | 0.009527 |
| TMEM245 | 0.358386 | 8.3E-05 |
| TRIM44 | 0.357595 | 0.006654 |
| FLNA | 0.356644 | 0.014009 |
| HOXB3 | 0.354832 | 0.001675 |
| KIAA1109 | 0.351929 | 0.000426 |
| PCED1A | 0.351287 | 0.001238 |
| PWAR6 | 0.337712 | 0.002571 |
| SMO | 0.336368 | 0.000757 |
| SLIT3 | 0.33474 | 0.021872 |
| ZNF431 | 0.333976 | 0.008681 |
| GJC1 | 0.331304 | 0.002881 |
| ZNF445 | 0.328755 | 0.001782 |
| NISCH | 0.325983 | 0.002777 |
| TSC2 | 0.325289 | 0.001713 |

| | | |
|---|---|---|
| PLCG1 | 0.324734 | 0.000692 |
| ANKRD36( | 0.3235 | 0.000107 |
| EPHB3 | 0.322828 | 9.34E-05 |
| TANC1 | 0.322276 | 0.017565 |
| HSPG2 | 0.320826 | 0.015972 |
| TAOK1 | 0.320484 | 0.000505 |
| BRAF | 0.319093 | 0.024529 |
| OBSCN | 0.316071 | 9.14E-05 |
| MARCH6 | 0.315358 | 0.016061 |
| SEC61A1 | 0.313549 | 0.00736 |
| MFSD14C | 0.309267 | 0.004875 |
| KMT2C | 0.301278 | 0.018278 |
| SLIT1 | 0.297949 | 0.039924 |
| CACNB4 | 0.297638 | 0.015447 |
| FLNB | 0.296457 | 0.001896 |
| PUM2 | 0.294472 | 0.003114 |
| LPCAT1 | 0.286152 | 0.008797 |
| DMXL2 | 0.285645 | 0.006396 |
| WDR27 | 0.284883 | 0.007657 |
| CDK10 | 0.284523 | 0.010519 |
| INTS1 | 0.283053 | 0.001437 |
| SORL1 | 0.280035 | 0.010351 |
| PIGQ | 0.270883 | 0.03472 |
| SYMPK | 0.270092 | 0.01541 |
| PLXNA3 | 0.268362 | 0.011819 |
| FBXL19 | 0.263377 | 0.005491 |
| AL078639. | 0.26089 | 0.004346 |
| AC074117. | 0.252632 | 0.020288 |

| **Cluster 14** | | |
|---|---|---|
| Gene | LogFC | pVal |
| PLCG2 | 4.489875 | 2.07E-53 |
| TLE4 | 1.984942 | 3E-19 |
| NEAT1 | 1.871749 | 1.39E-17 |
| SOX4 | 1.810777 | 2.95E-19 |
| CCDC80 | 1.722924 | 0.000928 |
| MAFB | 1.708843 | 3.5E-06 |
| CSKMT | 1.687451 | 4.23E-11 |
| MALAT1 | 1.651339 | 1.88E-31 |
| AMD1 | 1.624192 | 2.69E-09 |
| HEXIM1 | 1.578122 | 3.46E-08 |
| HES1 | 1.575823 | 0.000186 |
| PPP1R10 | 1.562645 | 1.48E-09 |
| HIST1H2B | 1.550301 | 5.11E-07 |
| CHD9 | 1.43269 | 3.85E-06 |
| PLK2 | 1.432376 | 0.000104 |
| HIST1H2B | 1.416423 | 9.04E-05 |
| AC007952. | 1.40895 | 6.84E-06 |
| PIK3R3 | 1.347446 | 0.000158 |
| SAT1 | 1.321632 | 0.008168 |
| IER5L | 1.294069 | 0.003344 |
| Z93241.1 | 1.292127 | 1.36E-05 |
| JUN | 1.223547 | 0.027037 |
| AC110769. | 1.219883 | 0.000695 |
| AC110285. | 1.218471 | 0.002401 |
| CCNL2 | 1.209454 | 0.010036 |
| RND3 | 1.208983 | 0.001363 |
| LUC7L3 | 1.184026 | 8.56E-06 |
| IER5 | 1.070015 | 0.007554 |
| HIST1H2A | 1.058711 | 0.030226 |
| AL021453. | 1.05627 | 0.001012 |
| INTS6 | 1.046795 | 0.034606 |
| PCDH10 | 1.020896 | 0.017379 |
| ELF3 | 1.018491 | 0.032035 |
| AC025164. | 0.945331 | 0.009507 |

| Cluster 15 | | |
|---|---|---|
| Gene | LogFC | pVal |
| CCNB1 | 1.03967 | 5.51E-08 |
| UBE2C | 1.037628 | 4.53E-10 |
| TUBB4B | 1.014022 | 1.22E-10 |
| TUBA1C | 0.938241 | 7.97E-05 |
| KPNA2 | 0.936724 | 1.67E-05 |
| CDC20 | 0.921988 | 4.45E-05 |
| HMGB2 | 0.913893 | 4.18E-13 |
| NUSAP1 | 0.858284 | 1.17E-11 |
| UBE2S | 0.83116 | 9.87E-08 |
| DLGAP5 | 0.818921 | 2.25E-05 |
| BIRC5 | 0.803277 | 1.2E-10 |
| PTTG1 | 0.800901 | 5.52E-07 |
| CDKN3 | 0.800501 | 0.00857 |
| CDK1 | 0.784946 | 2.79E-10 |
| CCNB2 | 0.777061 | 1.55E-08 |
| CKS1B | 0.760829 | 3.9E-15 |
| AURKA | 0.72787 | 0.000148 |
| PLK1 | 0.71369 | 9.97E-05 |
| CKS2 | 0.707537 | 4.72E-07 |
| AURKB | 0.697509 | 5.13E-06 |
| H2AFX | 0.695265 | 1.16E-10 |
| CENPA | 0.641992 | 0.000173 |
| MAD2L1 | 0.603113 | 4.21E-10 |
| TUBA1B | 0.602611 | 4.7E-24 |
| CALM2 | 0.595548 | 0.005248 |
| TOP2A | 0.569651 | 0.001436 |
| PBK | 0.56571 | 0.000235 |
| STMN1 | 0.549469 | 1.68E-17 |
| PSRC1 | 0.544152 | 0.010764 |
| PIMREG | 0.53476 | 3.04E-05 |
| KIF2C | 0.524662 | 0.000138 |
| CDCA8 | 0.513004 | 0.028881 |
| CCNA2 | 0.509792 | 0.00355 |
| CDCA3 | 0.506662 | 0.00383 |
| BUB1 | 0.501294 | 0.031709 |
| BUB3 | 0.497541 | 0.000596 |
| JPT1 | 0.491802 | 0.003536 |
| DYNLL1 | 0.484789 | 5.36E-08 |
| UBE2T | 0.469409 | 3.4E-05 |
| HMGB3 | 0.468224 | 0.010971 |
| KIFC1 | 0.462237 | 0.000127 |
| HMGN2 | 0.452948 | 0.000366 |
| LDHA | 0.440798 | 7.38E-11 |
| H1FX | 0.43742 | 0.020402 |
| SKA2 | 0.432726 | 4.21E-06 |
| BNIP3 | 0.424541 | 0.01562 |
| PCLAF | 0.424138 | 8.8E-05 |
| HMGB1 | 0.420589 | 0.007215 |
| TUBB | 0.417732 | 6.28E-21 |
| H2AFZ | 0.415473 | 1.62E-10 |
| RAN | 0.402089 | 2.51E-10 |
| OIP5 | 0.400006 | 0.004096 |
| CENPW | 0.390976 | 0.019034 |
| RANBP1 | 0.380282 | 1.14E-06 |

| | | |
|---|---|---|
| CALM3 | 0.366747 | 0.027437 |
| ENO1 | 0.364787 | 2.63E-08 |
| SNRPB | 0.344696 | 0.002158 |
| RPL27A | 0.324215 | 7.74E-15 |
| TMSB15A | 0.321948 | 0.037002 |
| TPI1 | 0.3207 | 1.97E-07 |
| TYMS | 0.319934 | 0.028266 |
| PHPT1 | 0.313703 | 0.02983 |
| H2AFV | 0.30498 | 0.023584 |
| PKM | 0.288987 | 0.001706 |
| RPL31 | 0.283371 | 2.02E-07 |
| LSM3 | 0.272092 | 0.003264 |
| SRP9 | 0.269946 | 0.015173 |
| TMSB10 | 0.268959 | 2.94E-09 |
| MIF | 0.267666 | 0.001826 |
| YBX1 | 0.263346 | 0.000139 |
| RPL13A | 0.26121 | 1.78E-10 |
| SNRPD2 | 0.25793 | 0.018644 |
| SNRPE | 0.25237 | 0.048467 |
| RPL35 | 0.251215 | 7.67E-09 |
| HMGN1 | 0.250866 | 7.12E-07 |

| **Cluster 16** | | |
|---|---|---|
| Gene | LogFC | pVal |
| FABP5 | 1.791108 | 0.001016 |
| LIX1 | 1.524209 | 0.037029 |
| TGFBI | 1.334973 | 0.005672 |
| RARRES2 | 1.110524 | 0.030608 |
| SCRG1 | 1.085136 | 0.004889 |
| PMEL | 0.991106 | 0.040102 |
| PAX3 | 0.844757 | 2.17E-05 |
| SPARC | 0.818332 | 0.00257 |
| FOXD3-AS | 0.815492 | 0.000241 |
| TMSB4X | 0.781718 | 0.003015 |
| MEF2C | 0.775956 | 0.00283 |
| CCND1 | 0.759869 | 0.001239 |
| EMP3 | 0.631156 | 0.011734 |
| APOE | 0.609938 | 0.000276 |
| HEBP1 | 0.597791 | 6.38E-05 |
| COL9A3 | 0.565178 | 0.0152 |
| METRN | 0.523896 | 0.001568 |
| OAF | 0.522883 | 0.041036 |
| FBLN1 | 0.520682 | 0.000812 |
| HSP90AA1 | 0.487773 | 0.00098 |
| CCT5 | 0.432289 | 0.002799 |
| NME1 | 0.397116 | 0.041286 |
| CD63 | 0.384999 | 0.000649 |
