## Supplementary Table 2 for "Enhanced metanephric specification to functional proximal tubule enables toxicity screening and infectious disease modelling in kidney organoids"

| Cluster 0 | | |
|---|---|---|
| Gene | LogFC | pVal |
| OGN | 1.120957 | 0 |
| ASPN | 0.889002 | 2.4E-215 |
| CRABP1 | 0.859729 | 0 |
| CXCL12 | 0.833008 | 0 |
| LRRC17 | 0.756381 | 0 |
| PCP4 | 0.7375 | 5.1E-106 |
| MAB21L1 | 0.714256 | 0 |
| PRRX1 | 0.699879 | 0 |
| PDGFRA | 0.682258 | 0 |
| IFI44L | 0.64704 | 2.9E-215 |
| NNAT | 0.638728 | 0 |
| POSTN | 0.628416 | 1.5E-121 |
| KCNE4 | 0.62186 | 1.1E-239 |
| CTSC | 0.619523 | 2.21E-31 |
| CCDC80 | 0.616512 | 0 |
| OLFML3 | 0.603256 | 2.9E-294 |
| MFAP4 | 0.599227 | 0 |
| CYTL1 | 0.585367 | 4.8E-213 |
| VIM | 0.573567 | 0 |
| MAB21L2 | 0.566189 | 0 |
| MARCKS | 0.559709 | 0 |
| CDC42EP5 | 0.559592 | 0 |
| DCN | 0.554428 | 4.9E-101 |
| COL3A1 | 0.552015 | 0 |
| DKK1 | 0.538332 | 1.22E-68 |
| SCX | 0.533762 | 2E-214 |
| PCOLCE | 0.531982 | 0 |
| FN1 | 0.530657 | 4.4E-278 |
| IL11RA | 0.529981 | 0 |
| GRP | 0.516494 | 9E-108 |
| ETV1 | 0.5144 | 4.4E-209 |
| DDIT4 | 0.511478 | 4.3E-151 |
| CALD1 | 0.511121 | 0 |
| IGF1 | 0.511079 | 1.14E-92 |
| CNN3 | 0.510384 | 0 |
| MDFI | 0.508079 | 0 |
| NOVA1 | 0.505673 | 9.1E-132 |
| HOXD10 | 0.502979 | 2.8E-235 |
| COL1A2 | 0.502488 | 0 |
| CRABP2 | 0.501654 | 0 |
| SULT1E1 | 0.499432 | 1.62E-51 |
| NDUFA4L2 | 0.498565 | 9.25E-34 |
| COL1A1 | 0.495328 | 0 |
| LGALS1 | 0.486737 | 0 |
| CDH11 | 0.479279 | 1.4E-294 |
| MEG3 | 0.471302 | 1E-226 |
| CSRP2 | 0.470207 | 2.3E-130 |
| CHST2 | 0.464772 | 1.6E-149 |
| IFI44 | 0.462257 | 2.3E-206 |
| SOX11 | 0.455667 | 8.9E-152 |
| TWIST1 | 0.450323 | 1.2E-183 |
| IGFBP5 | 0.447456 | 1.8E-236 |
| SOSTDC1 | 0.445972 | 5.4E-106 |
| EFNA5 | 0.444304 | 2.2E-141 |

| | | |
|---|---|---|
| HGF | 0.444072 | 2.28E-81 |
| BMP5 | 0.435691 | 1.4E-153 |
| MFAP2 | 0.434823 | 2.4E-274 |
| MEST | 0.43389 | 0 |
| LRRC4C | 0.432941 | 2E-149 |
| CDO1 | 0.429788 | 2.3E-203 |
| EDNRA | 0.429041 | 2.7E-183 |
| PRRX2 | 0.427532 | 1.5E-243 |
| HOXD11 | 0.427418 | 5.8E-170 |
| RASL11B | 0.419024 | 6.1E-158 |
| TPBG | 0.41739 | 5.1E-133 |
| HNRNPA1 | 0.415937 | 0 |
| SIX1 | 0.413202 | 2.8E-189 |
| OAF | 0.411604 | 2.9E-199 |
| FIBIN | 0.408451 | 8.28E-97 |
| AKAP12 | 0.407417 | 2.7E-143 |
| CTSK | 0.402531 | 2.4E-192 |
| SELENOM | 0.399273 | 1.7E-201 |
| SEPT6 | 0.398892 | 2E-186 |
| FSTL1 | 0.398809 | 3.3E-202 |
| MEIS2 | 0.396333 | 1.7E-142 |
| ID3 | 0.395482 | 5.8E-123 |
| NFIA | 0.394163 | 3E-181 |
| ZFHX4 | 0.393079 | 2.1E-135 |
| C1orf54 | 0.392405 | 3.2E-224 |
| FZD2 | 0.392178 | 4.1E-243 |
| RBMS3 | 0.388432 | 3.1E-171 |
| NPM1 | 0.385137 | 0 |
| KCNJ2 | 0.385113 | 2.45E-71 |
| FAT4 | 0.379052 | 1.6E-143 |
| MIR99AHG | 0.378791 | 7.7E-154 |
| PDZRN4 | 0.377004 | 3.2E-178 |
| GAS1 | 0.374558 | 2E-185 |
| ITGA8 | 0.371978 | 7.2E-167 |
| HOXA10 | 0.371742 | 1.4E-109 |
| MMP2 | 0.3674 | 1.2E-179 |
| PLPP3 | 0.366034 | 1.5E-115 |
| GLT8D2 | 0.364695 | 5.1E-176 |
| IFITM3 | 0.36442 | 4.6E-234 |
| EBF3 | 0.362814 | 1E-137 |
| CLDN11 | 0.358967 | 4.51E-97 |
| RPS11 | 0.357118 | 0 |
| HMGA2 | 0.356407 | 1.8E-110 |
| CLMP | 0.35155 | 4.9E-157 |
| RFLNB | 0.351035 | 2.2E-139 |
| DNM3OS | 0.348986 | 9.3E-164 |
| PCDH7 | 0.348463 | 4.2E-106 |
| ASXL3 | 0.346161 | 3.1E-147 |
| ISLR | 0.342668 | 5.4E-160 |
| SELENOP | 0.341706 | 9.29E-80 |
| AP3S1 | 0.341379 | 3.3E-103 |
| MEOX1 | 0.341248 | 2.3E-153 |
| ZFP36L1 | 0.340924 | 4E-137 |
| MEIS1 | 0.340896 | 4.7E-144 |
| TPM2 | 0.340178 | 1.6E-257 |
| LRP1B | 0.337392 | 3E-145 |

| | | |
|---|---|---|
| RPL13A | 0.336939 | 0 |
| CPE | 0.335827 | 0 |
| KCTD12 | 0.335637 | 7.4E-104 |
| LIMCH1 | 0.333371 | 1.7E-115 |
| TNMD | 0.332998 | 8.38E-30 |
| EIF3F | 0.331376 | 2E-307 |
| PHACTR2 | 0.331249 | 1.74E-97 |
| PTGFR | 0.330092 | 1.3E-100 |
| MARCKSL1 | 0.329819 | 2E-304 |
| TTC3 | 0.328622 | 3E-147 |
| SLITRK6 | 0.324566 | 3E-90 |
| RPS12 | 0.324082 | 0 |
| LITAF | 0.32376 | 1E-122 |
| FAT3 | 0.323642 | 2.2E-128 |
| OGFRL1 | 0.322942 | 2.11E-76 |
| CCND1 | 0.32239 | 7.9E-124 |
| DACT3 | 0.322059 | 1.1E-149 |
| RPL21 | 0.319089 | 0 |
| RPL27A | 0.316917 | 0 |
| LAYN | 0.314679 | 8.7E-135 |
| RPL24 | 0.314099 | 0 |
| ID1 | 0.311512 | 3.43E-68 |
| RPS16 | 0.310493 | 0 |
| BOC | 0.309671 | 4.3E-131 |
| COL5A2 | 0.309258 | 1E-117 |
| RBFOX2 | 0.308636 | 2.8E-149 |
| DMD | 0.305477 | 1.14E-93 |
| PLS3 | 0.30543 | 1.3E-110 |
| C6orf48 | 0.3033 | 6.9E-157 |
| PLEKHO1 | 0.303273 | 5.9E-129 |
| CDH2 | 0.302922 | 1.48E-31 |
| COL6A2 | 0.299103 | 1.1E-139 |
| NDN | 0.29891 | 4.8E-125 |
| MARVELD1 | 0.298417 | 4.2E-125 |
| KCNQ1OT1 | 0.298232 | 5.82E-95 |
| C12orf57 | 0.297259 | 3.2E-163 |
| SESN3 | 0.296262 | 6.82E-94 |
| SPOCK3 | 0.292302 | 2.2E-69 |
| ADAMTS6 | 0.292265 | 2.89E-72 |
| CNRIP1 | 0.292218 | 4E-126 |
| RPL30 | 0.292166 | 0 |
| AFF3 | 0.291986 | 4.6E-116 |
| EMCN | 0.291942 | 6.27E-81 |
| CNTN3 | 0.291538 | 3.9E-108 |
| RPS20 | 0.291288 | 1E-300 |
| PRSS12 | 0.290693 | 5.52E-78 |
| RPS15A | 0.289139 | 0 |
| KCNE5 | 0.288954 | 2.4E-112 |
| RPL23A | 0.288915 | 0 |
| PCDH18 | 0.287908 | 1.59E-95 |
| PTX3 | 0.287657 | 2.4E-37 |
| SFRP2 | 0.287243 | 3.1E-253 |
| EIF3E | 0.28678 | 6.6E-189 |
| RPL22 | 0.286281 | 0 |
| RPL6 | 0.285946 | 0 |
| KAT6B | 0.285941 | 1.28E-87 |

| | | |
|---|---|---|
| H3F3B | 0.285925 | 6.2E-275 |
| GAS2 | 0.284743 | 1.4E-81 |
| MDK | 0.282642 | 2.3E-237 |
| PLPPR3 | 0.282442 | 5.3E-117 |
| SERPINH1 | 0.281878 | 3.38E-94 |
| TRIL | 0.280937 | 2E-107 |
| RPL3 | 0.280684 | 0 |
| PMEPA1 | 0.280326 | 2.3E-111 |
| WSB1 | 0.280176 | 1.99E-62 |
| FBN2 | 0.279631 | 7.22E-96 |
| EBPL | 0.278805 | 6.7E-113 |
| COL5A1 | 0.277911 | 2.2E-107 |
| ZNF703 | 0.276897 | 6.42E-94 |
| PHLDA1 | 0.27659 | 4.14E-76 |
| PCDH9 | 0.276576 | 8.44E-50 |
| FBLN1 | 0.276086 | 4.5E-136 |
| FLRT2 | 0.275779 | 2.5E-100 |
| PARM1 | 0.275059 | 1.16E-92 |
| GULP1 | 0.274953 | 7.36E-94 |
| NELL2 | 0.274389 | 9.97E-41 |
| AC027031.2 | 0.274373 | 1.62E-86 |
| NCAM1 | 0.273908 | 1.7E-103 |
| RPS3 | 0.273645 | 0 |
| RPS19 | 0.273634 | 0 |
| CIRBP | 0.273402 | 7.7E-192 |
| ANGPTL4 | 0.272065 | 8.93E-83 |
| SNAI2 | 0.271795 | 4.54E-72 |
| RPL23 | 0.269462 | 0 |
| AUTS2 | 0.269261 | 1.38E-95 |
| SOX6 | 0.268317 | 2.05E-84 |
| SSR2 | 0.268019 | 6.7E-167 |
| LDB2 | 0.266901 | 8.9E-107 |
| BEX3 | 0.26599 | 3.2E-150 |
| NACA | 0.264866 | 0 |
| TBX3 | 0.263813 | 4.94E-87 |
| ENC1 | 0.263173 | 9.05E-71 |
| CALCRL | 0.26317 | 2.73E-82 |
| NAV3 | 0.262297 | 3.25E-88 |
| MAP1B | 0.261782 | 5.97E-75 |
| SPARCL1 | 0.261449 | 7.87E-60 |
| OLFM2 | 0.261322 | 2.54E-77 |
| RPLP2 | 0.260591 | 0 |
| EEF1D | 0.258615 | 9.2E-201 |
| NAP1L1 | 0.258579 | 2.2E-160 |
| RPL10 | 0.257522 | 0 |
| TMEM98 | 0.257412 | 2.23E-76 |
| COL25A1 | 0.257228 | 1.78E-98 |
| RRBP1 | 0.256687 | 4.81E-81 |
| PLK2 | 0.255998 | 1.84E-69 |
| EMP3 | 0.25599 | 4.1E-141 |
| RPL35A | 0.255951 | 0 |
| RPL27 | 0.255913 | 0 |
| JAM2 | 0.255564 | 1.88E-84 |
| MEX3B | 0.255281 | 8.32E-87 |
| HOXA11 | 0.254299 | 2.56E-50 |
| EVL | 0.254279 | 3.4E-77 |

| | | |
|---|---|---|
| RPL15 | 0.253983 | 0 |
| INHBA | 0.253288 | 1.23E-68 |
| RPL7 | 0.251024 | 0 |

| **Cluster 1** | | |
| --- | --- | --- |
| Gene | LogFC | pVal |
| MT1H | 2.952466 | 0 |
| MT1G | 2.863548 | 0 |
| MT2A | 2.533624 | 0 |
| MT1E | 2.49489 | 0 |
| MT1X | 2.341066 | 0 |
| MT1F | 2.108348 | 0 |
| APOE | 1.771033 | 0 |
| S100A1 | 1.541138 | 0 |
| TSPAN1 | 1.417369 | 0 |
| BHMT | 1.346481 | 0 |
| TMEM176A | 1.32179 | 0 |
| ALDH1A1 | 1.307945 | 0 |
| FMO1 | 1.266311 | 0 |
| PCSK1N | 1.236654 | 0 |
| FTL | 1.21338 | 0 |
| TMEM176B | 1.188161 | 0 |
| SMIM24 | 1.182897 | 0 |
| SPP1 | 1.178926 | 0 |
| LINC01781 | 1.17382 | 0 |
| MPC2 | 1.169346 | 0 |
| AMN | 1.165283 | 0 |
| SLC39A4 | 1.134736 | 0 |
| EPCAM | 1.124298 | 0 |
| ANXA4 | 1.115282 | 0 |
| CLDN7 | 1.113229 | 0 |
| NIT2 | 1.089089 | 0 |
| CLU | 1.062081 | 0 |
| ATP1B1 | 1.043514 | 0 |
| HPN | 1.041529 | 0 |
| CLDN4 | 1.025508 | 0 |
| AGT | 1.01058 | 0 |
| PDZK1 | 1.006715 | 0 |
| RHOB | 0.990245 | 0 |
| CYB5A | 0.989207 | 0 |
| CLDN3 | 0.977949 | 0 |
| CYBA | 0.976494 | 0 |
| CRYL1 | 0.974507 | 0 |
| F10 | 0.972895 | 0 |
| CUBN | 0.971194 | 0 |
| RIDA | 0.965471 | 0 |
| AIG1 | 0.962968 | 0 |
| C1QTNF12 | 0.95609 | 0 |
| LGALS2 | 0.951102 | 0 |
| GPX3 | 0.944202 | 0 |
| CD9 | 0.942148 | 0 |
| BSG | 0.940663 | 0 |
| LAMTOR5 | 0.931403 | 0 |
| FLRT3 | 0.929701 | 0 |
| PTGR1 | 0.928771 | 0 |
| KRT18 | 0.92312 | 0 |
| ATP6V1F | 0.914166 | 0 |
| LGMN | 0.906395 | 0 |
| CAMK2N1 | 0.89245 | 0 |
| SLC16A4 | 0.88002 | 0 |

| | | |
|---|---|---|
| KRT8 | 0.879816 | 0 |
| TM7SF2 | 0.877801 | 0 |
| ITM2B | 0.869766 | 0 |
| DAB2 | 0.868677 | 0 |
| ETFB | 0.866831 | 0 |
| GLYATL1 | 0.865633 | 0 |
| CHCHD10 | 0.859806 | 0 |
| TINAG | 0.858448 | 0 |
| C19orf33 | 0.850462 | 0 |
| PPP1R1A | 0.847788 | 0 |
| SERPINA1 | 0.845293 | 0 |
| SLC3A1 | 0.829539 | 0 |
| NPC2 | 0.829078 | 0 |
| TUBB4B | 0.825906 | 0 |
| CITED2 | 0.821488 | 0 |
| PBLD | 0.820628 | 0 |
| SLC9A3R1 | 0.809576 | 0 |
| CST3 | 0.809262 | 0 |
| KIF12 | 0.807402 | 0 |
| MT-ND3 | 0.807168 | 0 |
| EZR | 0.805826 | 0 |
| KLF6 | 0.804865 | 0 |
| HLA-A | 0.804513 | 0 |
| KCNJ15 | 0.803799 | 0 |
| TMEM256 | 0.803765 | 0 |
| DBI | 0.801578 | 0 |
| EPS8L2 | 0.78448 | 0 |
| CD151 | 0.781773 | 0 |
| CYS1 | 0.780988 | 0 |
| GSTP1 | 0.778288 | 0 |
| ATP5IF1 | 0.777751 | 0 |
| TMBIM6 | 0.772281 | 0 |
| SLC44A4 | 0.772031 | 0 |
| SMIM1 | 0.771564 | 0 |
| VAMP8 | 0.771493 | 0 |
| CLEC18A | 0.768737 | 0 |
| ACAA2 | 0.765832 | 0 |
| IL32 | 0.763832 | 0 |
| IMPA2 | 0.760196 | 0 |
| SLC39A5 | 0.75792 | 0 |
| CD68 | 0.751525 | 0 |
| MSRB1 | 0.745302 | 0 |
| CLIC1 | 0.739809 | 0 |
| PNP | 0.738892 | 0 |
| PPP1R16A | 0.736601 | 0 |
| AFP | 0.729619 | 0 |
| UGT2B7 | 0.727443 | 0 |
| SLC37A4 | 0.727367 | 0 |
| SPINT1 | 0.72697 | 0 |
| ASS1 | 0.716747 | 0 |
| MT-ND4 | 0.713575 | 0 |
| HLA-C | 0.712224 | 0 |
| PCBD1 | 0.70984 | 0 |
| RBPMS | 0.706434 | 0 |
| DPP4 | 0.704618 | 0 |
| PDGFA | 0.703194 | 0 |

| | | |
|---|---|---|
| GATM | 0.696905 | 0 |
| MGST3 | 0.696118 | 0 |
| GPX4 | 0.694325 | 0 |
| C11orf54 | 0.694155 | 0 |
| CKB | 0.692969 | 0 |
| CIDEB | 0.692747 | 0 |
| SPINT2 | 0.691034 | 0 |
| MDH1 | 0.690898 | 0 |
| BIN1 | 0.688562 | 0 |
| MT-ND2 | 0.688527 | 0 |
| RARRES2 | 0.686029 | 5E-301 |
| ASAH1 | 0.677373 | 0 |
| HDHD3 | 0.675668 | 0 |
| PGAM2 | 0.674639 | 1.6E-271 |
| MST1 | 0.673651 | 0 |
| CDKN1C | 0.673138 | 0 |
| FTH1 | 0.672648 | 0 |
| AKR1C3 | 0.672224 | 0 |
| CYSTM1 | 0.670172 | 0 |
| ACAT1 | 0.669548 | 0 |
| SMS | 0.669415 | 0 |
| FXYD2 | 0.667613 | 0 |
| CD63 | 0.666552 | 0 |
| GPD1 | 0.664202 | 0 |
| TMEM150/ | 0.664165 | 0 |
| CEBPD | 0.661654 | 0 |
| S100A14 | 0.657079 | 0 |
| MT-CYB | 0.656591 | 0 |
| AGPAT2 | 0.656391 | 0 |
| CDHR5 | 0.656004 | 0 |
| UGCG | 0.655353 | 0 |
| SLC51B | 0.654264 | 0 |
| APOA1 | 0.651894 | 1.4E-270 |
| MRPS36 | 0.65049 | 0 |
| DPYS | 0.648904 | 0 |
| CLEC18B | 0.645503 | 0 |
| LHX1 | 0.643591 | 0 |
| LDHB | 0.640731 | 0 |
| MT-CO2 | 0.640347 | 0 |
| GAL3ST1 | 0.639344 | 0 |
| RNF181 | 0.639129 | 0 |
| SERPINE2 | 0.636652 | 0 |
| FABP3 | 0.635058 | 0 |
| S100A10 | 0.630402 | 0 |
| SLC25A5 | 0.626639 | 0 |
| TSTD1 | 0.623456 | 0 |
| PPP1R14C | 0.621059 | 0 |
| APOM | 0.619277 | 0 |
| EMID1 | 0.617536 | 0 |
| DCDC2 | 0.61571 | 0 |
| VTN | 0.613478 | 0 |
| MT-CO3 | 0.612741 | 0 |
| CHPT1 | 0.610376 | 0 |
| ACSM2A | 0.610279 | 0 |
| MT-ATP6 | 0.607305 | 0 |
| SLC27A2 | 0.602872 | 0 |

| | | |
|---|---|---|
| DPEP1 | 0.599696 | 0 |
| ODC1 | 0.598492 | 0 |
| CYP27A1 | 0.596955 | 0 |
| PRR13 | 0.596007 | 0 |
| IGFBP7 | 0.590555 | 0 |
| GSTK1 | 0.589038 | 0 |
| SDC4 | 0.588291 | 0 |
| CTSB | 0.586941 | 0 |
| LRPAP1 | 0.586668 | 0 |
| NEU1 | 0.583712 | 0 |
| GCHFR | 0.582293 | 0 |
| CA2 | 0.57625 | 0 |
| STRADB | 0.574122 | 0 |
| EBP | 0.573073 | 0 |
| EMX2 | 0.572773 | 0 |
| CDK2AP2 | 0.57077 | 0 |
| NR2F6 | 0.570391 | 0 |
| MYO6 | 0.568062 | 0 |
| S100A11 | 0.567504 | 0 |
| RGS14 | 0.566797 | 0 |
| GIPC2 | 0.563091 | 0 |
| ABHD11 | 0.559474 | 0 |
| RAB25 | 0.556513 | 0 |
| PDLIM1 | 0.554146 | 0 |
| CDH6 | 0.551182 | 0 |
| JUNB | 0.548596 | 0 |
| DNPH1 | 0.548027 | 0 |
| QPRT | 0.545853 | 0 |
| GGH | 0.545576 | 0 |
| FOLR1 | 0.544345 | 0 |
| TXNDC17 | 0.543371 | 0 |
| ECH1 | 0.543236 | 0 |
| MT-ND1 | 0.542137 | 0 |
| TNFRSF12 | 0.540281 | 1.1E-164 |
| CALM3 | 0.539106 | 0 |
| HNF1B | 0.537728 | 0 |
| FTCD | 0.536753 | 0 |
| PRELID1 | 0.536362 | 0 |
| SCRN2 | 0.53598 | 0 |
| HEXB | 0.534769 | 0 |
| ACO2 | 0.533216 | 0 |
| TMEM141 | 0.53263 | 0 |
| ERICH5 | 0.531458 | 0 |
| CCDC198 | 0.530463 | 0 |
| PRDX1 | 0.528205 | 0 |
| PSAT1 | 0.51809 | 7E-298 |
| PLIN2 | 0.516215 | 0 |
| IER2 | 0.514842 | 0 |
| MT-ND4L | 0.512913 | 0 |
| SERPINF1 | 0.512125 | 8.2E-230 |
| CHST13 | 0.51156 | 0 |
| ERBB3 | 0.509645 | 0 |
| ACAA1 | 0.509233 | 0 |
| MSRB2 | 0.50726 | 0 |
| PEPD | 0.507096 | 0 |
| TST | 0.494922 | 0 |

| | | |
|---|---|---|
| RHOC | 0.492885 | 0 |
| OCIAD2 | 0.492818 | 0 |
| HGD | 0.492322 | 0 |
| S100A16 | 0.492034 | 0 |
| CXXC5 | 0.491618 | 0 |
| LPCAT3 | 0.49161 | 0 |
| TSPAN12 | 0.491285 | 0 |
| ELF3 | 0.491127 | 0 |
| APLP2 | 0.489683 | 0 |
| EPHX2 | 0.489591 | 0 |
| PLEKHA1 | 0.485193 | 0 |
| SLC6A13 | 0.48424 | 0 |
| GLRX | 0.482287 | 0 |
| PTGDS | 0.482179 | 3.55E-31 |
| ECI1 | 0.481872 | 0 |
| MARVELD | 0.480664 | 0 |
| LACTB2 | 0.478949 | 0 |
| AGTRAP | 0.477288 | 0 |
| GALM | 0.47715 | 0 |
| ADH5 | 0.476661 | 0 |
| HES4 | 0.47641 | 0 |
| FGFR4 | 0.475455 | 0 |
| COX6A1 | 0.473466 | 0 |
| ATP5MC3 | 0.473385 | 0 |
| PRSS8 | 0.471949 | 0 |
| PCK1 | 0.47069 | 0 |
| PON2 | 0.469881 | 0 |
| COX5B | 0.468612 | 0 |
| ENPP3 | 0.468488 | 0 |
| TMEM37 | 0.467374 | 0 |
| BRI3 | 0.466834 | 0 |
| HMGCL | 0.466628 | 0 |
| CISD1 | 0.465487 | 0 |
| RTN4 | 0.463165 | 0 |
| MGAT4B | 0.461687 | 0 |
| SSTR2 | 0.461526 | 0 |
| ADH6 | 0.460593 | 0 |
| ALDH6A1 | 0.459514 | 0 |
| GSTO1 | 0.458966 | 0 |
| GLTPD2 | 0.458699 | 0 |
| JUN | 0.457819 | 0 |
| TECR | 0.453985 | 0 |
| FJX1 | 0.453335 | 4.5E-292 |
| ZFP36 | 0.451361 | 5.8E-290 |
| A4GALT | 0.444008 | 0 |
| ACADVL | 0.443379 | 0 |
| ABRACL | 0.443014 | 0 |
| TXN | 0.442894 | 0 |
| ERI3 | 0.442043 | 0 |
| AKR7A2 | 0.440801 | 0 |
| KLHDC8B | 0.438223 | 0 |
| ATP5F1B | 0.437469 | 0 |
| VIL1 | 0.436208 | 0 |
| ATP5ME | 0.432351 | 0 |
| BTG2 | 0.429935 | 1.3E-293 |
| SERPINI1 | 0.428052 | 3.5E-273 |

| | | |
|---|---|---|
| TRAP1 | 0.426559 | 0 |
| ARHGAP1 | 0.42609 | 0 |
| RNF5 | 0.425087 | 0 |
| TUBG1 | 0.423211 | 0 |
| NDUFA5 | 0.423055 | 0 |
| ANK3 | 0.42195 | 0 |
| FAHD1 | 0.419156 | 0 |
| KIF21A | 0.417506 | 0 |
| CXADR | 0.417025 | 0 |
| PLPP1 | 0.416624 | 0 |
| ATOX1 | 0.415187 | 0 |
| HIBADH | 0.413386 | 0 |
| PMM1 | 0.413139 | 0 |
| SLC7A7 | 0.412407 | 1.1E-274 |
| PPFIBP1 | 0.411709 | 9.7E-265 |
| KLK1 | 0.410685 | 8.1E-224 |
| GLS | 0.40943 | 0 |
| SAT1 | 0.408625 | 0 |
| NUCB1 | 0.407226 | 0 |
| MT-CO1 | 0.406081 | 0 |
| SIGIRR | 0.405599 | 0 |
| ANPEP | 0.403257 | 0 |
| ECHS1 | 0.403129 | 0 |
| TNFSF12 | 0.401917 | 0 |
| HES1 | 0.400566 | 7.9E-134 |
| SYAP1 | 0.400449 | 0 |
| STX3 | 0.398866 | 0 |
| BLVRA | 0.398861 | 0 |
| COA3 | 0.398539 | 0 |
| CD24 | 0.397668 | 0 |
| NDUFB10 | 0.397319 | 0 |
| SETD3 | 0.396659 | 0 |
| DMTN | 0.396365 | 0 |
| ACADS | 0.394929 | 0 |
| TCEA3 | 0.394398 | 0 |
| AK4 | 0.394119 | 0 |
| LMAN2 | 0.39378 | 0 |
| SEPHS2 | 0.392847 | 0 |
| NINJ1 | 0.392709 | 0 |
| SUCLG1 | 0.391343 | 0 |
| SLC35D2 | 0.391119 | 0 |
| PAX8 | 0.390586 | 0 |
| UQCR10 | 0.390584 | 0 |
| ABCC6 | 0.39047 | 0 |
| ASRGL1 | 0.390381 | 0 |
| CERS2 | 0.389373 | 0 |
| CLPTM1L | 0.389202 | 0 |
| CLRN3 | 0.3891 | 0 |
| CALM1 | 0.389078 | 0 |
| AIFM1 | 0.387471 | 0 |
| GRN | 0.38724 | 0 |
| CAPG | 0.386919 | 0 |
| HSPE1 | 0.386747 | 0 |
| SLC25A4 | 0.386672 | 0 |
| NIPSNAP1 | 0.386562 | 0 |
| MYL12B | 0.386024 | 0 |

| | | |
|---|---|---|
| RAB29 | 0.385498 | 0 |
| TGIF1 | 0.385105 | 5.1E-291 |
| NDUFB9 | 0.385017 | 0 |
| SLC44A3 | 0.38391 | 0 |
| BMP4 | 0.383301 | 1E-303 |
| LINC00671 | 0.382772 | 5.2E-272 |
| ACOT13 | 0.382697 | 0 |
| TSPAN33 | 0.381114 | 0 |
| HSD17B14 | 0.380738 | 0 |
| ENTPD2 | 0.380417 | 0 |
| COL18A1 | 0.380058 | 0 |
| CD46 | 0.379752 | 0 |
| TNFSF10 | 0.379091 | 1.8E-145 |
| GPR160 | 0.378593 | 0 |
| SLC2A4RG | 0.376995 | 0 |
| TMSB4X | 0.376787 | 0 |
| CIB1 | 0.375998 | 0 |
| PLLP | 0.375998 | 0 |
| HSPB1 | 0.375605 | 4.8E-243 |
| RAB3IP | 0.375399 | 0 |
| C1orf210 | 0.374624 | 0 |
| CTSL | 0.374523 | 0 |
| GK | 0.374401 | 0 |
| CYB5R3 | 0.374206 | 0 |
| TUBA1C | 0.373147 | 0 |
| CSTB | 0.372862 | 0 |
| HOOK1 | 0.371965 | 0 |
| AQP6 | 0.371799 | 6.2E-293 |
| TMEM205 | 0.371667 | 0 |
| CXCL14 | 0.369382 | 3.2E-135 |
| MSMO1 | 0.366716 | 7.3E-207 |
| MITF | 0.366597 | 0 |
| ARMT1 | 0.366541 | 0 |
| CRYZ | 0.365553 | 0 |
| ECHDC2 | 0.365153 | 0 |
| CLDN2 | 0.365121 | 2.6E-177 |
| CRYAB | 0.364059 | 1.3E-241 |
| BCAP31 | 0.36162 | 0 |
| ATP6AP1 | 0.358821 | 0 |
| OCEL1 | 0.357473 | 0 |
| PPA1 | 0.357408 | 0 |
| TAGLN2 | 0.356922 | 9.9E-200 |
| VPS28 | 0.356422 | 0 |
| COX5A | 0.353794 | 0 |
| SH3YL1 | 0.35247 | 0 |
| METRNL | 0.351583 | 0 |
| SNX7 | 0.351537 | 0 |
| ATP6V1G1 | 0.34965 | 0 |
| CLDN6 | 0.348966 | 7E-161 |
| P2RX4 | 0.348357 | 0 |
| GGT1 | 0.347386 | 0 |
| GSTM4 | 0.347028 | 0 |
| GLYAT | 0.34661 | 2.6E-232 |
| HAGH | 0.346281 | 0 |
| GALE | 0.345971 | 0 |
| PCBP1 | 0.344925 | 0 |

| | | |
|---|---|---|
| MGST2 | 0.342829 | 0 |
| YBX3 | 0.34269 | 0 |
| ETHE1 | 0.342369 | 6E-307 |
| PARD6B | 0.342153 | 0 |
| UGT8 | 0.342119 | 0 |
| PRKAB1 | 0.342087 | 8.5E-253 |
| MYL6 | 0.341872 | 0 |
| STXBP2 | 0.341739 | 0 |
| DPY30 | 0.340846 | 0 |
| LSR | 0.340514 | 0 |
| F12 | 0.340513 | 1.2E-283 |
| FAM20C | 0.340429 | 0 |
| BCAM | 0.340237 | 0 |
| SERPINF2 | 0.339033 | 5E-303 |
| ATP5MC1 | 0.338077 | 0 |
| ARL6IP5 | 0.337217 | 0 |
| SORBS2 | 0.336886 | 0 |
| HOXB7 | 0.336573 | 0 |
| HHLA2 | 0.336201 | 0 |
| PLEKHJ1 | 0.334736 | 0 |
| ACSM3 | 0.334528 | 0 |
| CD164 | 0.33301 | 0 |
| FUOM | 0.332579 | 2.4E-285 |
| CCNG1 | 0.331971 | 0 |
| ATRAID | 0.33188 | 0 |
| DYNLT1 | 0.331809 | 0 |
| EIF1 | 0.331799 | 0 |
| SYPL1 | 0.330696 | 0 |
| C12orf75 | 0.330474 | 0 |
| ATP5PF | 0.330237 | 0 |
| YWHAH | 0.329951 | 2.7E-200 |
| FGFR3 | 0.329061 | 0 |
| NDUFC1 | 0.327766 | 0 |
| ATP5F1D | 0.327595 | 0 |
| PPDPF | 0.326761 | 0 |
| CLDN10 | 0.325892 | 1E-298 |
| ZDHHC12 | 0.325508 | 0 |
| KRT19 | 0.324694 | 3.6E-177 |
| METTL9 | 0.324551 | 0 |
| MSLN | 0.323634 | 1E-268 |
| APRT | 0.323589 | 0 |
| PHB | 0.323295 | 0 |
| PLOD2 | 0.32286 | 1.7E-260 |
| UGT3A1 | 0.322544 | 1.1E-209 |
| CNDP2 | 0.322446 | 0 |
| COA1 | 0.322026 | 0 |
| CARHSP1 | 0.321988 | 0 |
| LRP2 | 0.321954 | 2.1E-280 |
| BCAT2 | 0.321781 | 4.4E-280 |
| LEFTY1 | 0.321746 | 2E-120 |
| BLVRB | 0.319623 | 0 |
| RGS9 | 0.319295 | 0 |
| GOT1 | 0.318626 | 0 |
| ALDH2 | 0.318496 | 1E-297 |
| NDFIP2 | 0.317697 | 0 |
| RAB5C | 0.317601 | 0 |

| | | |
|---|---|---|
| MPST | 0.317437 | 0 |
| VMP1 | 0.316878 | 1.8E-216 |
| RBM47 | 0.316848 | 0 |
| UBL5 | 0.316725 | 0 |
| ACSM2B | 0.316207 | 2.5E-263 |
| ADI1 | 0.315452 | 2.7E-270 |
| SNX4 | 0.315434 | 0 |
| GDF15 | 0.315409 | 1.8E-168 |
| HSDL2 | 0.315258 | 0 |
| TM7SF3 | 0.315187 | 5E-305 |
| PANCR | 0.314457 | 7.2E-292 |
| MFSD10 | 0.314197 | 0 |
| PHYHD1 | 0.314147 | 0 |
| TMED4 | 0.313715 | 0 |
| SLC25A1 | 0.313277 | 0 |
| RHOA | 0.313191 | 0 |
| PRDX6 | 0.312856 | 0 |
| SCP2 | 0.312823 | 0 |
| FKBP2 | 0.311635 | 0 |
| AEBP1 | 0.310934 | 4.8E-251 |
| SLC5A8 | 0.310887 | 5.5E-225 |
| PGRMC1 | 0.309646 | 0 |
| PTTG1IP | 0.309128 | 0 |
| CDH16 | 0.308435 | 2E-305 |
| CTSA | 0.307771 | 1.3E-276 |
| DBNDD1 | 0.307568 | 4.4E-264 |
| GPR137B | 0.306198 | 0 |
| MRPL32 | 0.305959 | 1E-296 |
| GJB1 | 0.305223 | 1E-303 |
| CTSZ | 0.305203 | 4.3E-242 |
| PROS1 | 0.305001 | 8.2E-289 |
| RDH10 | 0.304312 | 1.2E-270 |
| ATP1B3 | 0.304012 | 4E-235 |
| ATP5MD | 0.303446 | 0 |
| GUSB | 0.302516 | 0 |
| LRRC19 | 0.301787 | 3.7E-257 |
| BNIP3 | 0.301161 | 6.2E-228 |
| SDHD | 0.300697 | 0 |
| STARD10 | 0.300208 | 0 |
| FAM131C | 0.300075 | 3.8E-233 |
| MAP7 | 0.299787 | 1E-303 |
| ARID3A | 0.299671 | 1.5E-289 |
| BPHL | 0.299498 | 1E-295 |
| FBP1 | 0.29903 | 2.9E-238 |
| ORAI3 | 0.298963 | 1E-296 |
| NOP10 | 0.298751 | 0 |
| ARPC3 | 0.298455 | 0 |
| SDHA | 0.298212 | 0 |
| PLBD1 | 0.297585 | 1.3E-279 |
| MID1IP1 | 0.297569 | 1.8E-212 |
| HNF4A | 0.297195 | 5.6E-243 |
| SLC3A2 | 0.297071 | 0 |
| VAT1 | 0.296802 | 5E-298 |
| LAMTOR2 | 0.295903 | 0 |
| ENTPD5 | 0.295807 | 1E-303 |
| MISP3 | 0.294774 | 2.8E-293 |

| | | |
|---|---|---|
| SKAP2 | 0.294709 | 5E-301 |
| TOLLIP | 0.294552 | 0 |
| METTL27 | 0.294351 | 4E-267 |
| TRAPPC6/ | 0.294169 | 0 |
| PAIP2B | 0.294137 | 4.2E-261 |
| TXNIP | 0.294113 | 4.6E-226 |
| PALM | 0.293717 | 7.9E-277 |
| BHLHE40 | 0.293004 | 2.8E-235 |
| EIF6 | 0.292983 | 0 |
| MT-ND5 | 0.291097 | 1.9E-265 |
| MAP1LC3E | 0.290907 | 2.2E-278 |
| ATP5F1C | 0.290307 | 0 |
| AZGP1 | 0.290283 | 7.8E-174 |
| RDH5 | 0.290169 | 6.5E-266 |
| NEAT1 | 0.287845 | 1.4E-233 |
| PRR15L | 0.287306 | 1.3E-251 |
| ESPN | 0.286958 | 5E-230 |
| NDUFA1 | 0.286487 | 0 |
| GRTP1 | 0.286435 | 4E-296 |
| CTSV | 0.286071 | 5.9E-166 |
| MNS1 | 0.285992 | 2.6E-185 |
| SPINT1-AS | 0.285526 | 4.1E-281 |
| RENBP | 0.285475 | 2.4E-276 |
| HOOK2 | 0.284842 | 2.1E-281 |
| ATP5PD | 0.28474 | 0 |
| RPS27L | 0.284502 | 0 |
| MIR4458H | 0.284031 | 7.4E-253 |
| TFEC | 0.283565 | 2.3E-249 |
| TTC38 | 0.282276 | 6E-278 |
| REEP6 | 0.281824 | 5.1E-218 |
| RHOBTB1 | 0.281209 | 6.1E-239 |
| STEAP1 | 0.28079 | 1.8E-279 |
| PLEKHB2 | 0.280597 | 5E-303 |
| CTSD | 0.280224 | 1E-296 |
| CLDN19 | 0.28001 | 2.5E-253 |
| CAPZA2 | 0.278935 | 0 |
| GNG11 | 0.278596 | 2.2E-240 |
| F2RL1 | 0.278227 | 8.6E-262 |
| GHITM | 0.27557 | 0 |
| MGMT | 0.275556 | 3.9E-264 |
| MARCH9 | 0.275538 | 7.1E-274 |
| TMEM219 | 0.275482 | 0 |
| IER3IP1 | 0.27507 | 0 |
| GFER | 0.274894 | 3E-214 |
| TPT1 | 0.274654 | 0 |
| CYC1 | 0.274638 | 4.1E-280 |
| DYNC2LI1 | 0.273476 | 2.4E-271 |
| NDUFA2 | 0.273145 | 0 |
| METTL7B | 0.272413 | 3.2E-221 |
| PRKAR1B | 0.272403 | 4.9E-201 |
| MRPL57 | 0.272259 | 0 |
| FCRLB | 0.271593 | 1.4E-252 |
| MCCD1 | 0.271401 | 1.9E-169 |
| SURF1 | 0.270984 | 1.2E-274 |
| LAMP2 | 0.270861 | 1E-274 |
| MCUR1 | 0.270608 | 1.4E-284 |

| | | |
|---|---|---|
| MAPK3 | 0.270584 | 9.9E-253 |
| BDH2 | 0.270574 | 1E-279 |
| AC004540. | 0.270475 | 5.2E-235 |
| SLC30A1 | 0.27025 | 2.7E-263 |
| MT1M | 0.269142 | 1.5E-77 |
| COX17 | 0.269069 | 7.2E-278 |
| ETFA | 0.268312 | 3.3E-279 |
| DSEL | 0.267989 | 2.1E-151 |
| YIF1A | 0.267449 | 7.9E-279 |
| PITX2 | 0.267209 | 3E-223 |
| DDT | 0.266766 | 4.6E-291 |
| HEXA | 0.26625 | 5.7E-276 |
| TRAPPC4 | 0.26614 | 1E-299 |
| FUCA2 | 0.263757 | 4.7E-263 |
| ACMSD | 0.26364 | 7.9E-217 |
| ACY3 | 0.262892 | 3.8E-178 |
| PXMP2 | 0.262853 | 1.1E-226 |
| AQP3 | 0.262752 | 2E-203 |
| FAM107B | 0.262597 | 5.8E-238 |
| BEX2 | 0.26225 | 3.1E-228 |
| AGPAT3 | 0.262129 | 2.2E-229 |
| RTKN | 0.261809 | 1E-262 |
| PSME1 | 0.261768 | 0 |
| COMT | 0.261274 | 2.6E-293 |
| RIT1 | 0.261055 | 5.3E-239 |
| UCP2 | 0.260849 | 2.3E-198 |
| STARD3NI | 0.260723 | 2.6E-281 |
| MTCH2 | 0.26039 | 9.4E-266 |
| GPR89B | 0.25999 | 5.4E-250 |
| RGL3 | 0.259709 | 7.5E-252 |
| COX7B | 0.259077 | 1E-267 |
| USH1C | 0.258812 | 0 |
| C4orf3 | 0.25847 | 4.6E-213 |
| HINT1 | 0.258224 | 0 |
| SMPDL3A | 0.258106 | 5.2E-203 |
| OGG1 | 0.257827 | 2.3E-244 |
| SHC2 | 0.257188 | 4.5E-237 |
| MRPL37 | 0.256799 | 1.2E-259 |
| SDHC | 0.256713 | 0 |
| IRX3 | 0.256306 | 5E-110 |
| DHRS4L2 | 0.254722 | 5.7E-218 |
| VPS35 | 0.254254 | 4.4E-278 |
| PLA2G12B | 0.254188 | 2.6E-227 |
| SLC22A5 | 0.254091 | 8.4E-241 |
| LINC02381 | 0.253865 | 4.3E-188 |
| DHRS1 | 0.253712 | 3E-250 |
| ACP5 | 0.253361 | 9.1E-177 |
| STK17A | 0.253118 | 1.8E-210 |
| SEMA5A | 0.253073 | 2.2E-144 |
| UGT2A3 | 0.253071 | 2E-193 |
| ARL1 | 0.252922 | 3.8E-293 |
| ZDHHC4 | 0.252683 | 2.8E-269 |
| COBLL1 | 0.251923 | 2.6E-186 |
| PCK2 | 0.251856 | 9.4E-241 |
| DUSP9 | 0.25147 | 6.7E-173 |
| NDUFAF3 | 0.25145 | 1.9E-254 |

| | | |
|---|---|---|
| PSMA2 | 0.251243 | 5.4E-167 |
| PEBP1 | 0.251234 | 2.8E-280 |
| NDUFS7 | 0.251161 | 2.1E-282 |
| CLEC18C | 0.251042 | 1.6E-230 |
| ECI2 | 0.250797 | 3.2E-270 |
| MMP24OS | 0.250511 | 2.7E-239 |
| ABHD14B | 0.25017 | 5.6E-242 |
| UQCRC1 | 0.250088 | 9.9E-250 |
| LTC4S | 0.25003 | 3.6E-218 |

| **Cluster 2** | | |
|---|---|---|
| Gene | LogFC | pVal |
| COL2A1 | 2.044258 | 0 |
| CNMD | 1.752884 | 0 |
| COL9A1 | 1.486764 | 4E-296 |
| MATN4 | 1.431244 | 1.5E-168 |
| C2orf40 | 1.348341 | 0 |
| COL9A2 | 1.334779 | 0 |
| EPYC | 1.32573 | 2.15E-67 |
| COL9A3 | 1.311346 | 0 |
| SFRP2 | 1.27332 | 0 |
| MIA | 1.267857 | 0 |
| DLK1 | 1.223813 | 1.7E-213 |
| MGP | 1.220136 | 0 |
| COL11A1 | 0.997224 | 1.3E-281 |
| IGFBP5 | 0.94689 | 3.4E-192 |
| PMP22 | 0.882652 | 0 |
| FRZB | 0.874031 | 7.2E-213 |
| PTN | 0.85791 | 0 |
| ITM2A | 0.832525 | 2.5E-111 |
| CPE | 0.823591 | 0 |
| HAPLN1 | 0.810947 | 2.1E-140 |
| SOX9 | 0.806524 | 1.1E-274 |
| RFLNA | 0.748417 | 6.4E-288 |
| COL11A2 | 0.706957 | 6.9E-162 |
| SOX5 | 0.695264 | 4.1E-213 |
| SCRG1 | 0.693449 | 2.63E-82 |
| FIBIN | 0.688815 | 1.2E-123 |
| MEST | 0.655894 | 0 |
| WWP2 | 0.647523 | 1.3E-178 |
| PEG10 | 0.646898 | 4.7E-103 |
| COL1A2 | 0.631037 | 0 |
| S100B | 0.627492 | 8.6E-106 |
| POMC | 0.620059 | 1.67E-95 |
| CTHRC1 | 0.60767 | 2.3E-157 |
| GDF10 | 0.602649 | 1.5E-141 |
| MEG3 | 0.598109 | 1.2E-179 |
| CAPN6 | 0.595389 | 1.6E-184 |
| TSC22D1 | 0.583822 | 3.8E-252 |
| SOX6 | 0.575951 | 1.2E-209 |
| PRELP | 0.567966 | 2.4E-146 |
| C1QTNF3 | 0.544926 | 2.09E-71 |
| KLF2 | 0.544642 | 4E-115 |
| JDP2 | 0.542905 | 3.5E-175 |
| DHRS3 | 0.515398 | 4.6E-165 |
| COL8A1 | 0.514621 | 2.06E-82 |
| TSPO | 0.513854 | 6.6E-233 |
| SOD3 | 0.505765 | 4.2E-119 |
| DNM3OS | 0.503042 | 3.9E-152 |
| GRP | 0.502367 | 3.99E-60 |
| C9orf3 | 0.500731 | 1.2E-146 |
| CDC42EP3 | 0.493312 | 3.3E-99 |
| CCDC34 | 0.485478 | 7.71E-92 |
| RHOBTB3 | 0.485353 | 8E-154 |
| FZD2 | 0.484791 | 4E-198 |
| PPIC | 0.482914 | 2.5E-123 |

| | | |
|---|---|---|
| FGFR1 | 0.476516 | 1.6E-169 |
| NDRG2 | 0.471513 | 3.2E-170 |
| COL12A1 | 0.466564 | 1.05E-81 |
| RGCC | 0.461079 | 1.97E-96 |
| RGS2 | 0.449894 | 1.13E-85 |
| BOC | 0.440795 | 1.1E-159 |
| GCNT1 | 0.439793 | 1.7E-125 |
| NFIB | 0.43378 | 1.6E-177 |
| PRICKLE1 | 0.424208 | 9E-122 |
| MMP2 | 0.423874 | 2.5E-123 |
| GPC3 | 0.422331 | 7.4E-109 |
| SNAI2 | 0.419724 | 2.01E-85 |
| RARG | 0.414453 | 1.1E-147 |
| C1QL1 | 0.414082 | 1.22E-74 |
| COL8A2 | 0.411516 | 4.7E-135 |
| FABP7 | 0.404448 | 9.1E-66 |
| COL1A1 | 0.404227 | 1.7E-236 |
| FGFRL1 | 0.402572 | 2.7E-107 |
| COL27A1 | 0.401443 | 3.1E-102 |
| SORBS2 | 0.400664 | 3.91E-32 |
| IL11RA | 0.400104 | 2.6E-189 |
| SERTAD4 | 0.399824 | 1E-114 |
| FBXO2 | 0.396326 | 6.98E-85 |
| TRPS1 | 0.395743 | 6.14E-93 |
| OAF | 0.3934 | 9.1E-116 |
| GAS1 | 0.392888 | 2.3E-118 |
| GAS2 | 0.390965 | 6.3E-107 |
| RPS12 | 0.389396 | 0 |
| CYR61 | 0.389323 | 5.22E-69 |
| LIMCH1 | 0.389196 | 2.05E-97 |
| SIX1 | 0.385543 | 5.19E-92 |
| DACT1 | 0.384855 | 4.3E-114 |
| ENPP2 | 0.384082 | 1.84E-74 |
| GPM6B | 0.383915 | 3.3E-106 |
| LMCD1 | 0.382763 | 3.7E-108 |
| RNF175 | 0.380613 | 6.7E-107 |
| KCNQ1OT | 0.37783 | 8.9E-119 |
| SNHG8 | 0.377453 | 1.1E-115 |
| PPP1R1B | 0.368726 | 4.08E-78 |
| MECOM | 0.367212 | 3E-111 |
| CRTAP | 0.365339 | 3.4E-130 |
| SERTAD4- | 0.364378 | 5.35E-94 |
| PLAC9 | 0.361972 | 3.51E-64 |
| TPR | 0.360263 | 4.79E-31 |
| FOXC1 | 0.353141 | 1.01E-69 |
| PCOLCE2 | 0.347659 | 4.12E-72 |
| COL6A2 | 0.345902 | 1.4E-154 |
| THSD4 | 0.345436 | 1.72E-88 |
| CDH11 | 0.34405 | 1.1E-127 |
| CKAP4 | 0.344016 | 2.97E-98 |
| CALU | 0.340701 | 1.04E-97 |
| EMILIN1 | 0.340093 | 3.7E-125 |
| RPS11 | 0.338773 | 0 |
| MDFI | 0.337924 | 1.8E-174 |
| SMOC2 | 0.337762 | 4.97E-65 |
| MEOX2 | 0.335158 | 2.81E-77 |

| | | |
|---|---|---|
| ELN | 0.333693 | 8.39E-81 |
| HMCN1 | 0.333513 | 4.98E-70 |
| PIEZO2 | 0.332476 | 5.07E-78 |
| RPL15 | 0.331617 | 0 |
| SELENOM | 0.33133 | 5.2E-110 |
| HNRNPA1 | 0.328867 | 0 |
| SIX2 | 0.328613 | 2.91E-73 |
| RPL13A | 0.327776 | 0 |
| ZFHX4 | 0.326681 | 7.75E-74 |
| RPS13 | 0.326455 | 0 |
| FOS | 0.326078 | 4.81E-18 |
| RCN2 | 0.326057 | 2.3E-118 |
| RRBP1 | 0.325728 | 4.3E-102 |
| VCAN | 0.325586 | 6.47E-84 |
| APP | 0.322218 | 1.12E-78 |
| SPARC | 0.318074 | 1.1E-122 |
| UCMA | 0.315936 | 1.3E-29 |
| IGF2.1 | 0.315345 | 3.66E-71 |
| PCDH18 | 0.315327 | 1.53E-64 |
| RPL27A | 0.310672 | 0 |
| BCAT1 | 0.31029 | 9.56E-74 |
| RPS19 | 0.30955 | 0 |
| SRM | 0.308702 | 8.4E-71 |
| COL14A1 | 0.308305 | 6.74E-35 |
| RPL21 | 0.307624 | 0 |
| CCND2 | 0.304522 | 2.98E-86 |
| MAB21L2 | 0.304439 | 9.7E-123 |
| RPS27A | 0.302468 | 0 |
| FBLN1 | 0.302403 | 3.44E-77 |
| C1orf21 | 0.302001 | 1.7E-76 |
| IGFBP2 | 0.301166 | 1.7E-136 |
| RPL30 | 0.299998 | 0 |
| GSN | 0.298294 | 1.04E-72 |
| NPM1 | 0.298099 | 0 |
| C6orf48 | 0.297986 | 1.3E-126 |
| EIF3H | 0.294576 | 8.7E-148 |
| RPL18A | 0.292916 | 0 |
| LUM | 0.291342 | 6.16E-20 |
| RPL34 | 0.289931 | 0 |
| ID3 | 0.287309 | 1.28E-64 |
| CNN3 | 0.285092 | 1.8E-135 |
| C4orf48 | 0.28446 | 3.72E-70 |
| ITGA10 | 0.284151 | 1.94E-68 |
| RPS16 | 0.282779 | 1E-306 |
| LSAMP | 0.282008 | 4.73E-71 |
| RPL6 | 0.281085 | 0 |
| RPS24 | 0.280101 | 0 |
| TNS2 | 0.280087 | 1.25E-66 |
| CADM1 | 0.279935 | 8.75E-69 |
| RPL32 | 0.279067 | 0 |
| SERPINH1 | 0.277847 | 1.8E-69 |
| EPB41L2 | 0.277661 | 8.4E-71 |
| MFAP4 | 0.277542 | 4.9E-112 |
| SCG5 | 0.276321 | 2.59E-43 |
| PRDM6 | 0.275114 | 1.09E-60 |
| CLDN11 | 0.275089 | 6.24E-28 |

| | | |
|---|---|---|
| C1GALT1 | 0.274737 | 1.32E-43 |
| RPS4X | 0.273715 | 9.3E-178 |
| CD44 | 0.273646 | 8.31E-61 |
| CAVIN1 | 0.273193 | 4.73E-73 |
| RPS3A | 0.272385 | 0 |
| CALD1 | 0.27226 | 3.72E-87 |
| FGFR2 | 0.271589 | 4.22E-60 |
| EPB41L4A | 0.270897 | 1.54E-73 |
| RPS23 | 0.270724 | 0 |
| ACAN | 0.269769 | 6.7E-58 |
| ERG | 0.268673 | 9.52E-73 |
| MEX3B | 0.267039 | 1.86E-48 |
| PPP3CA | 0.267017 | 9.67E-43 |
| PHACTR2 | 0.266435 | 1.1E-48 |
| CDC42EP5 | 0.266199 | 9.9E-121 |
| RPLP0 | 0.265551 | 1.5E-245 |
| STK26 | 0.265304 | 9.27E-58 |
| ZFAS1 | 0.265067 | 1.3E-119 |
| C12orf57 | 0.264373 | 4.93E-82 |
| GLI3 | 0.264063 | 6.31E-60 |
| SRGAP1 | 0.264001 | 1.68E-57 |
| RPLP1 | 0.263173 | 0 |
| RPL22 | 0.262278 | 1E-290 |
| RPL10 | 0.262203 | 0 |
| FBLN2 | 0.262127 | 2.9E-61 |
| SETBP1 | 0.262124 | 5.15E-56 |
| RPSA | 0.261702 | 6.4E-284 |
| ISM1 | 0.26071 | 6.37E-52 |
| EEF1A1 | 0.259198 | 0 |
| THBS1 | 0.258172 | 2.04E-54 |
| ID1 | 0.257667 | 9.3E-41 |
| RPS25 | 0.257227 | 2.3E-240 |
| PAPSS2 | 0.256703 | 1.53E-50 |
| SSR2 | 0.255818 | 2.6E-128 |
| MPPED2 | 0.254551 | 3.5E-59 |
| MIR99AHG | 0.254159 | 3.03E-39 |
| P4HA1 | 0.253519 | 1.41E-35 |
| PDGFRA | 0.252594 | 2.18E-78 |
| DAAM1 | 0.252328 | 1.88E-36 |
| EEF1D | 0.25064 | 6.8E-156 |

| Cluster 3 | | |
|---|---|---|
| Gene | LogFC | pVal |
| CLDN4 | 1.420317 | 0 |
| SPP1 | 1.141706 | 1.5E-240 |
| EPCAM | 1.121877 | 0 |
| S100A14 | 1.070768 | 0 |
| CLDN3 | 1.067829 | 0 |
| RHOB | 1.062102 | 0 |
| CLDN7 | 1.036889 | 0 |
| ATP1B1 | 1.034226 | 0 |
| KLF6 | 1.015481 | 0 |
| ANXA4 | 1.011461 | 0 |
| CKB | 0.966025 | 0 |
| FLRT3 | 0.949427 | 0 |
| TMEM176/ | 0.944043 | 0 |
| EZR | 0.935523 | 0 |
| CLU | 0.932357 | 0 |
| TMEM176I | 0.930224 | 0 |
| MPC2 | 0.920837 | 0 |
| ODC1 | 0.919717 | 1.4E-294 |
| TSPAN1 | 0.905436 | 1.3E-119 |
| UGT2B7 | 0.903208 | 0 |
| CD24 | 0.891561 | 0 |
| CITED2 | 0.890484 | 0 |
| DPP4 | 0.875006 | 0 |
| CYBA | 0.866605 | 0 |
| CLDN6 | 0.860548 | 4.4E-225 |
| SMIM24 | 0.857564 | 0 |
| CAMK2N1 | 0.855488 | 0 |
| SMS | 0.8358 | 0 |
| KRT8 | 0.832555 | 0 |
| KRT18 | 0.821766 | 0 |
| AGPAT2 | 0.812838 | 0 |
| CYS1 | 0.80229 | 0 |
| PDZK1 | 0.800219 | 0 |
| LINC02381 | 0.798493 | 6.5E-218 |
| PPP1R1A | 0.792723 | 0 |
| HPN | 0.787345 | 0 |
| PCSK1N | 0.783043 | 2.8E-220 |
| AMN | 0.780984 | 0 |
| IGFBP7 | 0.778486 | 0 |
| FXYD2 | 0.777217 | 4.1E-292 |
| KCNJ15 | 0.773441 | 0 |
| ITM2B | 0.763305 | 0 |
| VAMP8 | 0.759019 | 0 |
| PLEKHA1 | 0.753608 | 0 |
| LINC01781 | 0.751801 | 1.3E-284 |
| SLC44A4 | 0.75165 | 0 |
| RBPMS | 0.750897 | 0 |
| CD151 | 0.738185 | 0 |
| CLIC1 | 0.73718 | 0 |
| APLP2 | 0.734884 | 0 |
| DCDC2 | 0.72983 | 1.4E-270 |
| PTGR1 | 0.727768 | 9.2E-285 |
| HOTAIRM1 | 0.723812 | 7.9E-185 |
| BSG | 0.718884 | 0 |

| | | |
|---|---|---|
| PDGFA | 0.714465 | 2.4E-269 |
| IL32 | 0.710458 | 1.68E-80 |
| ELF3 | 0.696468 | 3E-183 |
| TXNIP | 0.6962 | 3.1E-277 |
| DAB2 | 0.691765 | 1E-298 |
| AIG1 | 0.688685 | 0 |
| JUN | 0.684838 | 0 |
| SPINT2 | 0.680724 | 0 |
| CCDC198 | 0.678139 | 4.8E-234 |
| GSTP1 | 0.677398 | 0 |
| KIF12 | 0.676156 | 0 |
| C19orf33 | 0.674612 | 1.7E-140 |
| PCBD1 | 0.662597 | 0 |
| ACAA2 | 0.661232 | 0 |
| IMPA2 | 0.660081 | 0 |
| SPINT1 | 0.655262 | 7E-304 |
| UGCG | 0.653792 | 2E-222 |
| LGALS2 | 0.652756 | 4.3E-249 |
| RAB25 | 0.650261 | 1.8E-276 |
| TMBIM6 | 0.650181 | 0 |
| LHX1 | 0.649849 | 9.8E-192 |
| PAX8 | 0.647734 | 6E-296 |
| CST3 | 0.641224 | 0 |
| KRT19 | 0.640826 | 4.1E-201 |
| MYO6 | 0.6402 | 1.9E-259 |
| APOE | 0.640107 | 0 |
| CEBPD | 0.63878 | 3.3E-237 |
| S100A11 | 0.638633 | 0 |
| RTN4 | 0.634125 | 0 |
| CYSTM1 | 0.630883 | 0 |
| TNFRSF12 | 0.630014 | 2.07E-88 |
| IRX1 | 0.624019 | 1.57E-79 |
| RIDA | 0.623661 | 0 |
| CD9 | 0.621679 | 4.5E-259 |
| EMID1 | 0.619978 | 1.1E-254 |
| CRYL1 | 0.616207 | 0 |
| SERPINE2 | 0.611958 | 2.3E-232 |
| HNF1B | 0.611694 | 1.1E-237 |
| F10 | 0.609461 | 9.8E-166 |
| SLC3A1 | 0.609322 | 1.3E-196 |
| SOX4 | 0.60475 | 4E-175 |
| CDKN1C | 0.59985 | 1.8E-160 |
| GAL3ST1 | 0.598026 | 8.6E-258 |
| FMO1 | 0.597717 | 2.1E-132 |
| ERBB4 | 0.597307 | 1.6E-152 |
| SDC4 | 0.589185 | 9.5E-226 |
| EMX2 | 0.588376 | 5.4E-235 |
| MGST3 | 0.588193 | 0 |
| CDK2AP2 | 0.578864 | 1.5E-181 |
| IRX3 | 0.576369 | 4.3E-132 |
| CYB5A | 0.574048 | 0 |
| HOOK1 | 0.57208 | 3E-200 |
| NR2F6 | 0.568593 | 0 |
| ADAMTS1 | 0.567645 | 5.3E-144 |
| TINAG | 0.567144 | 7.7E-154 |
| PRKAB1 | 0.566418 | 8.5E-191 |

| | | |
|---|---|---|
| MRPS36 | 0.563491 | 0 |
| TSTD1 | 0.562031 | 8.5E-141 |
| PLOD2 | 0.552954 | 1.5E-202 |
| RDH10 | 0.55291 | 1.3E-168 |
| IER2 | 0.548349 | 1.3E-226 |
| COL18A1 | 0.543208 | 3.7E-231 |
| ERBB3 | 0.542599 | 3.6E-192 |
| CLEC18A | 0.540579 | 3.7E-209 |
| SLC9A3R1 | 0.539765 | 7.2E-248 |
| C1QTNF12 | 0.538031 | 1.3E-151 |
| ABHD11 | 0.535632 | 4.4E-190 |
| TGIF1 | 0.53494 | 8.8E-167 |
| MITF | 0.533781 | 4.4E-189 |
| TUBB2B | 0.531706 | 6.9E-182 |
| PPP1R14C | 0.531264 | 9.6E-193 |
| CXADR | 0.530534 | 5.3E-214 |
| CD63 | 0.528932 | 0 |
| HES1 | 0.527818 | 5.18E-84 |
| TM7SF2 | 0.527385 | 4.8E-228 |
| EPS8L2 | 0.526329 | 8E-207 |
| PNP | 0.525964 | 2.7E-198 |
| NPC2 | 0.524137 | 0 |
| CLEC18B | 0.522974 | 5.7E-218 |
| MT-ND3 | 0.522857 | 0 |
| JUNB | 0.522146 | 1E-150 |
| DBI | 0.521702 | 0 |
| CRYAB | 0.520384 | 1.3E-112 |
| C12orf75 | 0.520127 | 4.9E-224 |
| SLC27A2 | 0.519496 | 1.1E-144 |
| ALDH1A1 | 0.519144 | 1.5E-126 |
| FTL | 0.514757 | 0 |
| LGMN | 0.512584 | 2.6E-165 |
| CXXC5 | 0.512057 | 0 |
| HLA-A | 0.511491 | 6.1E-129 |
| ATP5IF1 | 0.510493 | 0 |
| S100A16 | 0.510004 | 2E-220 |
| NIT2 | 0.509629 | 3.3E-168 |
| LRPAP1 | 0.508862 | 4.6E-292 |
| S100A10 | 0.508851 | 6.4E-285 |
| MGLL | 0.506215 | 2.9E-136 |
| MGAT4B | 0.50574 | 1.1E-230 |
| ERICH5 | 0.505703 | 3.6E-181 |
| LAMTOR5 | 0.501424 | 0 |
| CUBN | 0.496743 | 2.2E-111 |
| YWHAH | 0.496029 | 6.3E-120 |
| C11orf54 | 0.494897 | 2.5E-186 |
| COBLL1 | 0.493328 | 1.7E-152 |
| PON2 | 0.491932 | 1.3E-203 |
| KIF21A | 0.491357 | 7.8E-152 |
| MT-ND4 | 0.489276 | 0 |
| BIN1 | 0.484011 | 5.1E-194 |
| RHOA | 0.482338 | 2.2E-254 |
| LAPTM4B | 0.482161 | 1.7E-266 |
| TMSB4X | 0.481233 | 2.4E-165 |
| CALM3 | 0.480907 | 8.5E-249 |
| SLC16A4 | 0.477843 | 4.7E-140 |

| | | |
|---|---|---|
| ABRACL | 0.474918 | 6.9E-202 |
| UGT8 | 0.474 | 1.6E-158 |
| AP1M2 | 0.473864 | 2.3E-137 |
| FJX1 | 0.47365 | 1.76E-88 |
| NEU1 | 0.473592 | 2.3E-179 |
| DYNC2LI1 | 0.472388 | 1E-177 |
| ZBTB20 | 0.472175 | 3.4E-133 |
| CAPG | 0.469586 | 1.4E-191 |
| HGD | 0.469325 | 2.1E-161 |
| PRR13 | 0.469118 | 6.3E-209 |
| C1orf210 | 0.468911 | 1.1E-179 |
| RAB3IP | 0.467371 | 3.3E-165 |
| ATP6V1F | 0.467368 | 1.1E-270 |
| COMT | 0.467033 | 4.8E-180 |
| POU3F3 | 0.465453 | 9.5E-139 |
| SYAP1 | 0.460751 | 1.1E-170 |
| LMO4 | 0.460247 | 1.2E-106 |
| PERP | 0.460223 | 3.3E-138 |
| CD46 | 0.458867 | 1.7E-216 |
| MDH1 | 0.458641 | 6.5E-283 |
| MSRB1 | 0.458409 | 8.7E-167 |
| TAGLN2 | 0.457646 | 6.48E-86 |
| HLA-C | 0.457353 | 1.3E-158 |
| CHPT1 | 0.455993 | 3.1E-181 |
| CIDEB | 0.454838 | 1.4E-133 |
| MYL12B | 0.454794 | 0 |
| ANK3 | 0.454706 | 4.9E-140 |
| SKIL | 0.452849 | 2.1E-126 |
| SLC44A3 | 0.451933 | 3E-156 |
| DYNLT1 | 0.449667 | 7.6E-249 |
| LSR | 0.448414 | 5.8E-172 |
| NDUFA5 | 0.447564 | 1.2E-278 |
| FAM131C | 0.44654 | 7.8E-141 |
| SLC6A13 | 0.446088 | 2.6E-150 |
| HOXD8 | 0.446047 | 2E-109 |
| SNX7 | 0.444523 | 1.3E-179 |
| ID4 | 0.444222 | 6.8E-103 |
| PGRMC1 | 0.443032 | 1E-303 |
| STX3 | 0.439037 | 2.1E-167 |
| SLC37A4 | 0.43884 | 2.7E-173 |
| TMEM150A | 0.438426 | 2.1E-151 |
| TSPAN12 | 0.435207 | 2.7E-162 |
| LDHB | 0.435133 | 3E-274 |
| MRPL32 | 0.434575 | 2.5E-139 |
| DST | 0.434494 | 5.3E-105 |
| OCIAD2 | 0.434282 | 3.7E-150 |
| GNG11 | 0.43057 | 5.2E-146 |
| MAP1LC3B | 0.430379 | 4.1E-166 |
| KLF5 | 0.428337 | 6.1E-120 |
| DMTN | 0.42831 | 5.4E-138 |
| DPEP1 | 0.427625 | 1.7E-128 |
| GLYATL1 | 0.427388 | 9.7E-102 |
| TSPAN33 | 0.426577 | 7.1E-162 |
| HDHD3 | 0.426541 | 4.7E-148 |
| FTH1 | 0.426476 | 0 |
| MYL12A | 0.426147 | 4.6E-208 |

| | | |
|---|---|---|
| PBLD | 0.425591 | 1.1E-138 |
| RHOBTB1 | 0.425569 | 1E-141 |
| BMP4 | 0.423514 | 4E-120 |
| ELSPBP1 | 0.423103 | 7.84E-89 |
| PRSS8 | 0.422568 | 6.3E-149 |
| PSMA2 | 0.421587 | 4.5E-109 |
| AKIRIN1 | 0.420807 | 1.7E-149 |
| ARID3A | 0.420557 | 1.9E-157 |
| MYL6 | 0.419824 | 0 |
| NEAT1 | 0.418734 | 3.4E-108 |
| DSP | 0.415172 | 1.1E-129 |
| SEMA5A | 0.415131 | 8.49E-90 |
| RGS14 | 0.415075 | 4.4E-123 |
| CXCL14 | 0.412635 | 6.33E-55 |
| SEPHS2 | 0.412427 | 9.1E-157 |
| DBNDD1 | 0.411901 | 2.6E-131 |
| SH3YL1 | 0.41127 | 8.3E-143 |
| MARVELD | 0.411086 | 9.6E-119 |
| MT-CO3 | 0.411059 | 0 |
| HOXB7 | 0.410555 | 2.6E-131 |
| ECH1 | 0.410043 | 9E-180 |
| RDH11 | 0.408395 | 4.2E-124 |
| LPCAT3 | 0.407925 | 8.1E-147 |
| ATP6V1G1 | 0.407892 | 3E-297 |
| MT-CO2 | 0.407029 | 0 |
| EGFL7 | 0.405635 | 1E-120 |
| CDHR5 | 0.405624 | 4.1E-120 |
| SYPL1 | 0.405398 | 2.4E-193 |
| HSPB1 | 0.403682 | 1.07E-83 |
| LMAN2 | 0.40286 | 4.4E-205 |
| SQSTM1 | 0.402293 | 2.9E-120 |
| SKAP2 | 0.401719 | 3.4E-129 |
| PCBP1 | 0.401686 | 3.3E-209 |
| ARPC3 | 0.400777 | 2.4E-265 |
| UNCX | 0.400535 | 1.2E-120 |
| CTXN1 | 0.399095 | 6.6E-143 |
| DSC2 | 0.398582 | 1.6E-136 |
| GPX4 | 0.397339 | 1.6E-180 |
| CISD1 | 0.395953 | 3.1E-166 |
| GADD45A | 0.39557 | 2.1E-202 |
| MT-ND4L | 0.394459 | 1.3E-198 |
| COA3 | 0.393315 | 8.6E-189 |
| METTL9 | 0.393132 | 1.7E-207 |
| RBM47 | 0.392676 | 2E-130 |
| ETFB | 0.391986 | 1.3E-178 |
| METRNL | 0.389364 | 1.6E-135 |
| PTTG1IP | 0.388884 | 1.7E-185 |
| SLC39A5 | 0.387963 | 3.3E-138 |
| QPRT | 0.386904 | 1.8E-135 |
| CAPN1 | 0.386162 | 2.5E-150 |
| FAM107B | 0.384958 | 3.4E-123 |
| GRN | 0.383983 | 2.4E-130 |
| RGL3 | 0.383823 | 3E-128 |
| CERS2 | 0.383422 | 1.1E-153 |
| SAT1 | 0.383266 | 1.94E-80 |
| CNDP2 | 0.382407 | 2.3E-147 |

| | | |
|---|---|---|
| PRR15L | 0.382393 | 1.3E-122 |
| PARD6B | 0.381968 | 7.6E-127 |
| MSRB2 | 0.381454 | 1.6E-143 |
| DPY30 | 0.380771 | 2.2E-184 |
| PDLIM1 | 0.380393 | 8.54E-80 |
| MIA2 | 0.380294 | 5.1E-131 |
| VMP1 | 0.37842 | 5.6E-111 |
| YPEL2 | 0.378212 | 7.3E-113 |
| PLPP1 | 0.377952 | 1.9E-131 |
| AGT | 0.376556 | 2.59E-55 |
| CDH6 | 0.375408 | 1.38E-98 |
| GIPC2 | 0.37425 | 4.2E-108 |
| BCAP31 | 0.372562 | 3E-181 |
| PMM1 | 0.372555 | 2.2E-130 |
| SLC39A4 | 0.371633 | 3.44E-58 |
| MXRA8 | 0.371381 | 1.2E-172 |
| A4GALT | 0.371351 | 2.9E-117 |
| SERPINA1 | 0.369724 | 1.03E-48 |
| CRYZ | 0.369446 | 7.4E-119 |
| TMEM50A | 0.367126 | 1.9E-177 |
| CD164 | 0.366281 | 1.1E-181 |
| FGFR4 | 0.365482 | 4.2E-126 |
| STARD10 | 0.365011 | 6.2E-137 |
| IER3IP1 | 0.364741 | 1.5E-166 |
| ASPH | 0.363579 | 2.2E-125 |
| HES4 | 0.36209 | 1.9E-118 |
| LAMA1 | 0.361715 | 1.4E-117 |
| GALM | 0.360474 | 2.7E-114 |
| KCNJ16 | 0.359824 | 5.7E-117 |
| PURPL | 0.359271 | 2.04E-90 |
| RNF5 | 0.358979 | 3.6E-171 |
| AKR1C3 | 0.357499 | 2.78E-81 |
| TNFSF10 | 0.356725 | 8.52E-26 |
| OTUD1 | 0.356508 | 4.74E-91 |
| MAP7 | 0.355143 | 1E-116 |
| UQCR10 | 0.355009 | 7.2E-243 |
| GIPC1 | 0.354426 | 2.2E-124 |
| LRRK2 | 0.354409 | 1.2E-111 |
| NIPSNAP1 | 0.353667 | 5.8E-159 |
| UBL5 | 0.353141 | 8.4E-259 |
| FOLR1 | 0.351974 | 4.58E-95 |
| IRX5 | 0.351866 | 1.52E-64 |
| ASAH1 | 0.351501 | 3.1E-133 |
| DNM1 | 0.351339 | 3.6E-118 |
| MT-ATP6 | 0.349264 | 1.3E-263 |
| STARD3NI | 0.349052 | 2.9E-129 |
| MKNK2 | 0.34862 | 2.5E-124 |
| CCNG1 | 0.34859 | 2.9E-125 |
| ECI1 | 0.347806 | 3.5E-158 |
| NOP10 | 0.347532 | 5.1E-188 |
| ANXA3 | 0.347104 | 2.36E-65 |
| CYP27A1 | 0.346814 | 1.13E-98 |
| YBX3 | 0.344641 | 1.6E-135 |
| ACAT1 | 0.344342 | 6.1E-123 |
| MT-ND2 | 0.343256 | 2.7E-164 |
| RNF181 | 0.343023 | 1.6E-152 |

| | | |
|---|---|---|
| ARID5B | 0.34292 | 2.5E-108 |
| SLC35D2 | 0.342726 | 1.5E-109 |
| CAPZA2 | 0.342709 | 7.7E-152 |
| RHOC | 0.342326 | 8.4E-118 |
| EIF6 | 0.341555 | 1E-166 |
| ARHGAP1 | 0.341461 | 7.1E-118 |
| MT-CYB | 0.341406 | 2.3E-208 |
| TAX1BP1 | 0.341225 | 2.9E-127 |
| PPFIBP1 | 0.340967 | 4.12E-76 |
| GSTK1 | 0.340494 | 3.8E-141 |
| ARHGAP2 | 0.337602 | 1.77E-94 |
| VIL1 | 0.337312 | 2.8E-105 |
| FHOD3 | 0.337157 | 2.7E-100 |
| SNCA | 0.336252 | 6.9E-105 |
| STXBP2 | 0.336163 | 2.3E-126 |
| COX6A1 | 0.33546 | 3.7E-243 |
| SINHCAF | 0.335415 | 5.7E-100 |
| BTG2 | 0.335357 | 2.36E-74 |
| COL4A2 | 0.334616 | 3.4E-146 |
| TPM1 | 0.333825 | 1.8E-106 |
| FAHD1 | 0.333577 | 4.7E-114 |
| LLGL2 | 0.333468 | 6E-118 |
| GLRX | 0.333452 | 1.64E-98 |
| MID1IP1 | 0.331785 | 1.58E-88 |
| TUBA1C | 0.331616 | 2.3E-158 |
| CHST13 | 0.330886 | 1.25E-79 |
| CIB1 | 0.330785 | 1.3E-136 |
| LRRC19 | 0.330634 | 4.2E-103 |
| AC004540. | 0.330615 | 1.2E-99 |
| BTG1 | 0.330168 | 2.46E-71 |
| TMEM59 | 0.329276 | 3.3E-181 |
| SLC5A8 | 0.327715 | 6.69E-74 |
| FGFR3 | 0.327216 | 2.4E-101 |
| JUP | 0.327184 | 8E-116 |
| ARMT1 | 0.326655 | 2.1E-113 |
| ACADS | 0.326567 | 8.7E-112 |
| ZDHHC12 | 0.326421 | 6.8E-121 |
| SPATS2L | 0.325968 | 5.1E-117 |
| BHLHE40 | 0.325818 | 3.19E-83 |
| SIGIRR | 0.325459 | 1.5E-108 |
| EFNA1 | 0.324635 | 2.61E-91 |
| PTPN13 | 0.324559 | 2.04E-86 |
| MFSD10 | 0.324279 | 2.1E-119 |
| ZFP36 | 0.324123 | 2.68E-70 |
| RTN3 | 0.323433 | 1.2E-135 |
| VPS28 | 0.323337 | 9.3E-161 |
| TMEM205 | 0.323306 | 2.1E-122 |
| SLC25A4 | 0.32221 | 5.2E-120 |
| CYB5R3 | 0.321782 | 3.3E-126 |
| PODXL2 | 0.321141 | 1.06E-88 |
| EIF1 | 0.320729 | 3.6E-287 |
| ECHDC2 | 0.3206 | 1.3E-113 |
| PSME1 | 0.320576 | 3E-141 |
| NDUFB9 | 0.320324 | 2.5E-111 |
| PALM | 0.320111 | 1.6E-101 |
| HMGCL | 0.318957 | 7.4E-125 |

| | | |
|---|---|---|
| SYNE2 | 0.31809 | 5.85E-89 |
| SHC2 | 0.316169 | 4.39E-92 |
| SORBS2 | 0.316012 | 2E-105 |
| ARL1 | 0.315253 | 5.6E-136 |
| HOOK2 | 0.314902 | 1.2E-113 |
| GPRC5C | 0.314562 | 8.76E-91 |
| ADH6 | 0.313948 | 3.59E-83 |
| FKBP2 | 0.313073 | 6.2E-127 |
| ATP6V0B | 0.312638 | 1.5E-145 |
| SERPINF2 | 0.311372 | 2.43E-96 |
| BHMT | 0.311221 | 4.65E-65 |
| ACAA1 | 0.3103 | 5.4E-99 |
| TXN | 0.310242 | 1.1E-172 |
| FNBP1L | 0.310045 | 3.5E-99 |
| PAWR | 0.309825 | 6.7E-103 |
| DDAH2 | 0.309655 | 8E-143 |
| SPINT1-AS | 0.309475 | 7.1E-100 |
| CDC42 | 0.309437 | 2E-139 |
| TECR | 0.309277 | 5E-142 |
| AIFM1 | 0.309108 | 1.05E-98 |
| NDUFC1 | 0.307731 | 1.9E-162 |
| TMEM125 | 0.307558 | 1.27E-95 |
| TST | 0.306566 | 4.1E-101 |
| MT-ND5 | 0.306104 | 1.1E-169 |
| TP53I13 | 0.306067 | 1.6E-103 |
| CTDSPL | 0.30509 | 3.8E-99 |
| ACYP1 | 0.304722 | 1.34E-87 |
| RASSF7 | 0.30457 | 1.29E-94 |
| ARL6IP5 | 0.304402 | 2E-132 |
| PLBD1 | 0.304039 | 7.87E-94 |
| ATP1B3 | 0.303574 | 1.52E-82 |
| STRADB | 0.303554 | 7.6E-99 |
| JAG1 | 0.303042 | 1.26E-82 |
| NFKBIA | 0.302407 | 2.18E-39 |
| CRELD1 | 0.302159 | 5.1E-107 |
| MST1 | 0.302135 | 2.3E-81 |
| DDRGK1 | 0.302003 | 3.3E-107 |
| GSTO1 | 0.301962 | 8E-102 |
| HIBADH | 0.301386 | 1.7E-103 |
| TXNDC12 | 0.301311 | 2.4E-115 |
| MET | 0.300942 | 1.28E-93 |
| EIF5 | 0.300752 | 8.2E-129 |
| VAV3 | 0.300674 | 3.4E-85 |
| MYEF2 | 0.299887 | 2.18E-94 |
| DUSP9 | 0.299839 | 3.71E-65 |
| NUCB1 | 0.299295 | 6.7E-112 |
| BEX2 | 0.298695 | 1.28E-92 |
| TRAPPC4 | 0.298188 | 3.4E-103 |
| PLEKHJ1 | 0.297147 | 2.8E-116 |
| PPDPF | 0.29672 | 1.2E-130 |
| AC011043. | 0.296193 | 4.64E-96 |
| PRDX1 | 0.296004 | 8.3E-146 |
| SLC25A5 | 0.295506 | 7.6E-125 |
| ECHS1 | 0.295448 | 7.5E-137 |
| KIF9 | 0.295091 | 2.3E-102 |
| P4HTM | 0.294871 | 2.7E-102 |

| | | |
|---|---|---|
| REEP5 | 0.294788 | 5.1E-117 |
| HOXA9 | 0.294355 | 1.14E-71 |
| CNIH4 | 0.293931 | 3.9E-105 |
| IER5L | 0.293722 | 1.69E-59 |
| VPS35 | 0.293132 | 1.6E-104 |
| MARCH9 | 0.29289 | 2.88E-91 |
| GSPT1 | 0.292521 | 1.05E-96 |
| MALAT1 | 0.292379 | 2.9E-162 |
| ATRAID | 0.292081 | 5.9E-151 |
| BLVRA | 0.292057 | 3.22E-89 |
| NECTIN2 | 0.2918 | 8.2E-105 |
| ARF1 | 0.29153 | 3.5E-187 |
| CAPNS1 | 0.291061 | 1.9E-129 |
| VAT1 | 0.290989 | 4.1E-97 |
| CIAPIN1 | 0.290837 | 1.5E-102 |
| SRSF5 | 0.290259 | 1.3E-136 |
| ENPP3 | 0.289955 | 5.5E-73 |
| SESTD1 | 0.289565 | 1.74E-73 |
| SMTNL2 | 0.289488 | 5.2E-100 |
| TPM3 | 0.289324 | 5.2E-111 |
| PITX2 | 0.289122 | 4.42E-84 |
| LINC0095[illegible] | 0.288582 | 4.07E-89 |
| COX5B | 0.288044 | 1.9E-191 |
| TMEM37 | 0.28768 | 1.92E-75 |
| ENTPD5 | 0.287334 | 1.54E-82 |
| ACSM3 | 0.28726 | 5.58E-72 |
| ZDHHC4 | 0.287043 | 3.2E-105 |
| HEXB | 0.286725 | 1.97E-77 |
| LHX1-DT | 0.286702 | 2.91E-86 |
| HAGH | 0.286447 | 3.4E-104 |
| SSFA2 | 0.286417 | 5.07E-76 |
| MTSS1 | 0.286395 | 2.4E-86 |
| ACO2 | 0.285703 | 1.4E-99 |
| MLEC | 0.285382 | 2.1E-107 |
| DAZAP2 | 0.285363 | 3.3E-99 |
| MINDY2 | 0.284836 | 5.49E-82 |
| PAX2 | 0.284464 | 1.51E-84 |
| PLP2 | 0.284363 | 2.49E-60 |
| TMED4 | 0.28422 | 6.9E-104 |
| NAT14 | 0.284196 | 1.23E-94 |
| ZFP36L2 | 0.284069 | 5.65E-90 |
| STAP2 | 0.284053 | 2.2E-91 |
| COL4A1 | 0.283692 | 1.6E-118 |
| SETD3 | 0.283405 | 6.76E-92 |
| DOK6 | 0.282874 | 6.15E-87 |
| BMP3 | 0.282729 | 4.5E-44 |
| ATOX1 | 0.282542 | 8.1E-127 |
| SUPT4H1 | 0.282532 | 1.7E-112 |
| DNPH1 | 0.282425 | 2.3E-105 |
| ERP29 | 0.282248 | 1.1E-127 |
| MCUR1 | 0.281714 | 2.3E-103 |
| ARF6 | 0.281426 | 3.05E-91 |
| RTKN | 0.281291 | 9.38E-86 |
| RGS9 | 0.280558 | 9.48E-89 |
| COX17 | 0.280398 | 8.36E-93 |
| LACTB2 | 0.279742 | 2.43E-64 |

| | | |
|---|---|---|
| LINC01320 | 0.278274 | 8.04E-72 |
| PLLP | 0.278117 | 7.71E-76 |
| ABCC6 | 0.277813 | 3.53E-84 |
| DDR1 | 0.277745 | 2.2E-101 |
| PPP1R15A | 0.277545 | 3.13E-76 |
| FBRSL1 | 0.277531 | 6.82E-80 |
| ATP6V1D | 0.277453 | 1.9E-103 |
| LAMTOR2 | 0.277012 | 2E-112 |
| ERI3 | 0.276842 | 1.1E-101 |
| CDC34 | 0.276769 | 2.64E-90 |
| TMEM219 | 0.276744 | 3.4E-108 |
| PATJ | 0.276735 | 4.53E-81 |
| SUMF2 | 0.276641 | 1.69E-83 |
| FMC1 | 0.276282 | 6.63E-84 |
| CLPTM1L | 0.275708 | 5.7E-100 |
| CSTB | 0.275619 | 1.8E-107 |
| ARHGEF28 | 0.275544 | 3E-86 |
| GTF2H5 | 0.275382 | 2E-88 |
| RASSF6 | 0.274618 | 9.64E-80 |
| SNX29 | 0.27421 | 9.81E-81 |
| LAMB1 | 0.274043 | 1.08E-91 |
| PLEKHB2 | 0.273951 | 3.48E-92 |
| TMEM159 | 0.2739 | 5.7E-83 |
| NDFIP2 | 0.273819 | 6.39E-90 |
| DSTN | 0.273703 | 1.4E-125 |
| SELENOK | 0.273569 | 4.1E-115 |
| CLRN3 | 0.272973 | 8.48E-80 |
| TRAPPC6A | 0.272712 | 5E-91 |
| NDUFB10 | 0.272627 | 1.2E-148 |
| GCA | 0.272384 | 4.38E-85 |
| RPAIN | 0.271861 | 6.6E-106 |
| BPHL | 0.271859 | 2.14E-85 |
| SHISA5 | 0.271537 | 1.69E-84 |
| TMSB10 | 0.271381 | 5.3E-165 |
| PGF | 0.271254 | 3.69E-74 |
| DHRS7 | 0.270958 | 3.92E-83 |
| NUMB | 0.27091 | 1.03E-88 |
| DYNLRB1 | 0.270076 | 1.2E-157 |
| MBD2 | 0.270052 | 2.52E-85 |
| BRI3 | 0.269158 | 2.6E-138 |
| CAP1 | 0.269106 | 3.7E-102 |
| JAK1 | 0.268639 | 6.2E-69 |
| CRACR2B | 0.268542 | 5.6E-70 |
| GLOD4 | 0.268383 | 2.41E-87 |
| TXNDC17 | 0.267967 | 5.8E-119 |
| VAMP3 | 0.26725 | 3.09E-93 |
| ARL4A | 0.266299 | 1.88E-54 |
| HOXB8 | 0.266267 | 1.35E-76 |
| SEC23B | 0.266025 | 5.88E-59 |
| MVP | 0.265934 | 2.52E-82 |
| NDUFB4 | 0.265904 | 1E-131 |
| EPHX2 | 0.265203 | 1.08E-85 |
| PPP1R9A | 0.265145 | 6.95E-75 |
| HIP1R | 0.264997 | 1.15E-78 |
| YWHAZ | 0.264431 | 2.48E-94 |
| CD68 | 0.263878 | 1.38E-65 |

| | | |
|---|---|---|
| PPP1R16A | 0.263878 | 1.81E-75 |
| GPD1 | 0.263855 | 8.56E-68 |
| HOXD1 | 0.26369 | 1.84E-60 |
| REXO2 | 0.263584 | 1.8E-85 |
| SCP2 | 0.263296 | 2E-106 |
| SPSB3 | 0.263093 | 1.76E-79 |
| GRB7 | 0.26253 | 5.16E-80 |
| RIT1 | 0.262472 | 8.02E-72 |
| GOT1 | 0.261881 | 3.26E-76 |
| TUBB4B | 0.261571 | 2.2E-167 |
| LMBRD1 | 0.261555 | 1.21E-80 |
| DDX5 | 0.261503 | 3.2E-90 |
| TMEM123 | 0.261056 | 6.62E-79 |
| CCNDBP1 | 0.260935 | 4.33E-90 |
| CD2AP | 0.260449 | 7.31E-84 |
| TSG101 | 0.260331 | 8.08E-95 |
| CARHSP1 | 0.260181 | 1.37E-82 |
| BMP2 | 0.260173 | 1.43E-56 |
| CDC42SE2 | 0.259994 | 2.47E-74 |
| APOPT1 | 0.259946 | 7.2E-103 |
| TMED5 | 0.25977 | 3.67E-85 |
| ZFYVE21 | 0.259728 | 1.28E-97 |
| UBAC2 | 0.259431 | 6.14E-86 |
| ELOVL1 | 0.259118 | 1.03E-73 |
| COX20 | 0.259096 | 6.92E-78 |
| TRIM24 | 0.257651 | 1.15E-64 |
| ADI1 | 0.257414 | 2.37E-69 |
| CDH16 | 0.257159 | 4.15E-78 |
| TMEM179B | 0.256987 | 4.12E-86 |
| KRTCAP2 | 0.256385 | 2E-100 |
| SNX4 | 0.256219 | 3.25E-79 |
| SIM1 | 0.256179 | 9.24E-75 |
| FBXL5 | 0.256166 | 6.51E-78 |
| MT-ND1 | 0.25614 | 1.3E-152 |
| TRAM1 | 0.255544 | 6.93E-86 |
| PFN1 | 0.255371 | 1.4E-146 |
| ZNF83 | 0.255331 | 3.06E-80 |
| MIR4458H | 0.254904 | 9.06E-69 |
| B4GALT1 | 0.254881 | 1.08E-47 |
| ORMDL2 | 0.25479 | 5.21E-81 |
| COL4A3BP | 0.254501 | 3.36E-71 |
| RTL8A | 0.254443 | 2.65E-90 |
| CRYM | 0.253807 | 1.08E-71 |
| ZNF704 | 0.253274 | 2.65E-69 |
| BCL7A | 0.253142 | 2.51E-74 |
| ERICH1 | 0.252816 | 5.74E-69 |
| AFP | 0.252602 | 1.48E-43 |
| COX5A | 0.252539 | 6.1E-111 |
| SDCBP | 0.252042 | 1.03E-83 |
| ACADVL | 0.251919 | 8.19E-86 |
| NDFIP1 | 0.251864 | 2.3E-109 |
| MOB3B | 0.251718 | 1.22E-72 |
| CHCHD10 | 0.251674 | 5.57E-96 |
| MAF1 | 0.251199 | 5.28E-84 |
| P2RX4 | 0.251193 | 2.49E-66 |
| ATP6V0E1 | 0.250816 | 6.9E-118 |

| | | |
|---|---|---|
| TMEM38B | 0.250453 | 4.94E-54 |
| SLC39A1 | 0.250412 | 3.84E-79 |
| PLIN2 | 0.250101 | 1.88E-47 |

| Cluster 4 | | |
|---|---|---|
| Gene | LogFC | pVal |
| CENPF | 1.631444 | 4.91E-94 |
| HMGB2 | 1.593274 | 1.2E-177 |
| UBE2C | 1.536531 | 5.83E-75 |
| PTTG1 | 1.354618 | 4.04E-63 |
| TOP2A | 1.290174 | 6.6E-84 |
| KPNA2 | 1.271392 | 3.22E-61 |
| PCLAF | 1.228754 | 3.1E-242 |
| TUBA1B | 1.162153 | 0 |
| CCNB1 | 1.145625 | 5.31E-51 |
| UBE2S | 1.144448 | 3.83E-68 |
| NUSAP1 | 1.110646 | 1.1E-89 |
| CKS2 | 1.085816 | 1.02E-68 |
| TYMS | 1.056343 | 9.4E-214 |
| CDK1 | 1.028435 | 3.84E-91 |
| CCNB2 | 1.019209 | 2.27E-56 |
| TPX2 | 1.002556 | 4.33E-75 |
| H2AFZ | 0.993345 | 0 |
| MKI67 | 0.985544 | 1.58E-61 |
| CDC20 | 0.952493 | 1.57E-53 |
| HMGN2 | 0.951308 | 0 |
| BIRC5 | 0.933331 | 4.26E-74 |
| TMSB15A | 0.928172 | 6.7E-260 |
| H2AFX | 0.924247 | 1.6E-109 |
| ASPM | 0.916265 | 7.75E-55 |
| GINS2 | 0.912125 | 1.4E-161 |
| DLGAP5 | 0.896869 | 5.11E-55 |
| CKAP2 | 0.867805 | 8.76E-69 |
| CENPE | 0.864194 | 8.19E-52 |
| CKS1B | 0.855071 | 2.1E-134 |
| MAD2L1 | 0.837256 | 2.1E-128 |
| NASP | 0.83068 | 2.3E-217 |
| TUBA1C | 0.794798 | 1.05E-11 |
| DUT | 0.789735 | 4.64E-96 |
| CDKN3 | 0.786551 | 9.24E-61 |
| STMN1 | 0.779824 | 0 |
| CENPW | 0.754133 | 1.8E-134 |
| HMGB1 | 0.75183 | 5E-262 |
| DEK | 0.748653 | 3.4E-224 |
| ZWINT | 0.742304 | 5.5E-158 |
| SMC4 | 0.720683 | 1.1E-106 |
| PCNA | 0.710211 | 6.9E-118 |
| HMGB3 | 0.70574 | 3.22E-79 |
| UBE2T | 0.702086 | 2.1E-125 |
| MIS18BP1 | 0.701879 | 1.38E-94 |
| LMNB1 | 0.701552 | 4.4E-130 |
| CENPA | 0.69339 | 3.75E-40 |
| ARL6IP1 | 0.691644 | 8.51E-13 |
| TUBB | 0.690663 | 0 |
| NUCKS1 | 0.677781 | 9.3E-120 |
| LGALS1 | 0.675048 | 2.8E-162 |
| KIF2C | 0.672389 | 1.62E-58 |
| SGO2 | 0.669963 | 1.14E-60 |
| KIF20B | 0.658813 | 3.74E-73 |
| PSIP1 | 0.653212 | 8E-165 |

| | | |
|---|---|---|
| DTYMK | 0.649251 | 3.9E-109 |
| MZT1 | 0.648188 | 1.96E-64 |
| MCM7 | 0.644135 | 1E-118 |
| CENPK | 0.641522 | 5.8E-139 |
| ORC6 | 0.641023 | 1.3E-132 |
| AURKB | 0.638809 | 2.3E-46 |
| PIMREG | 0.620348 | 1.08E-55 |
| HELLS | 0.619036 | 7.7E-110 |
| SKA2 | 0.60942 | 3.11E-78 |
| H2AFV | 0.607641 | 2.4E-114 |
| JPT1 | 0.602556 | 1.69E-33 |
| CCNA2 | 0.594413 | 5.94E-49 |
| TK1 | 0.592497 | 2.4E-100 |
| AURKA | 0.591224 | 3.59E-35 |
| HMMR | 0.591061 | 4.58E-42 |
| NDC80 | 0.59099 | 6.33E-59 |
| MXD3 | 0.588538 | 1.78E-50 |
| RANBP1 | 0.585128 | 8.1E-133 |
| NUDT1 | 0.582495 | 1E-128 |
| CDT1 | 0.579921 | 1.9E-125 |
| PLK1 | 0.578282 | 1.3E-35 |
| MCM3 | 0.577828 | 1.18E-84 |
| SMC2 | 0.574872 | 6.8E-108 |
| RRM1 | 0.571179 | 5.9E-122 |
| CLSPN | 0.570277 | 7E-102 |
| CALM2 | 0.565033 | 7.73E-53 |
| DNMT1 | 0.560331 | 6.14E-97 |
| TMPO | 0.556908 | 4.4E-102 |
| TUBB4B | 0.555396 | 1.23E-11 |
| CENPU | 0.555329 | 2.1E-103 |
| TUBB6 | 0.553418 | 6.49E-55 |
| TACC3 | 0.551951 | 1.16E-54 |
| RAN | 0.551422 | 2.1E-172 |
| GMNN | 0.549884 | 1.3E-108 |
| SNRNP25 | 0.549065 | 6.9E-118 |
| CDCA8 | 0.543451 | 8.81E-47 |
| NUF2 | 0.543386 | 5.3E-47 |
| ANP32E | 0.53805 | 6.87E-68 |
| KIF23 | 0.530268 | 6.15E-43 |
| CDCA3 | 0.528081 | 7.54E-34 |
| KIFC1 | 0.526938 | 9.13E-55 |
| CENPH | 0.525633 | 2.07E-98 |
| BUB3 | 0.521645 | 2.3E-50 |
| HNRNPA2 | 0.519207 | 2.4E-101 |
| MND1 | 0.516899 | 2.31E-90 |
| TROAP | 0.511057 | 4.88E-47 |
| PTN | 0.508012 | 3E-139 |
| MCM5 | 0.505748 | 3.94E-89 |
| GTSE1 | 0.501785 | 2.67E-47 |
| KIF20A | 0.501648 | 3.43E-42 |
| NNAT | 0.498786 | 3E-156 |
| NEK2 | 0.497701 | 3.92E-40 |
| MCM4 | 0.497134 | 3.61E-89 |
| KIF11 | 0.49259 | 9.89E-52 |
| CKAP2L | 0.489821 | 6.14E-57 |
| CRABP1 | 0.486928 | 1.7E-184 |

| | | |
|---|---|---|
| FABP5 | 0.485868 | 1.82E-76 |
| EZH2 | 0.480432 | 1.28E-86 |
| RFC4 | 0.479853 | 8.33E-91 |
| SGO1 | 0.477131 | 2.85E-56 |
| RAD21 | 0.474036 | 5.64E-49 |
| ATAD2 | 0.471381 | 2.75E-83 |
| KNSTRN | 0.471064 | 4.65E-40 |
| SNRPD1 | 0.470829 | 3E-118 |
| DEPDC1 | 0.467717 | 1.29E-46 |
| LSM5 | 0.465251 | 3.1E-81 |
| CENPM | 0.463341 | 4.81E-85 |
| HIST1H4C | 0.463247 | 3.76E-29 |
| ITGB3BP | 0.462388 | 2.8E-87 |
| KIF22 | 0.462002 | 2.27E-53 |
| CENPN | 0.461412 | 2.53E-68 |
| RAD51AP1 | 0.459225 | 3.5E-87 |
| ECT2 | 0.458687 | 1.96E-44 |
| RMI2 | 0.455209 | 2.91E-80 |
| PSRC1 | 0.452904 | 2.65E-34 |
| CBX5 | 0.451823 | 7.06E-97 |
| ANP32B | 0.447599 | 4.7E-125 |
| RRM2 | 0.446178 | 1.02E-47 |
| CBX1 | 0.443937 | 1.98E-77 |
| MARCKS | 0.441389 | 4.3E-132 |
| DNAJC9 | 0.432804 | 1.39E-75 |
| FBLN1 | 0.426015 | 1.8E-102 |
| PRC1 | 0.424366 | 3.98E-49 |
| NT5DC2 | 0.423982 | 6.96E-85 |
| ZNF738 | 0.41916 | 2.86E-70 |
| TPM2 | 0.418198 | 4.8E-125 |
| TTK | 0.417652 | 1.5E-42 |
| C19orf48 | 0.41524 | 5.94E-74 |
| SRSF2 | 0.415109 | 1E-73 |
| DBF4 | 0.414086 | 2.33E-44 |
| KNL1 | 0.413997 | 6.64E-41 |
| FEN1 | 0.412988 | 4.02E-73 |
| UHRF1 | 0.411289 | 7.87E-80 |
| EMP2 | 0.409998 | 2.73E-79 |
| SRSF7 | 0.408197 | 6.49E-83 |
| CSRP2 | 0.406851 | 1.72E-50 |
| HNRNPD | 0.402642 | 1.88E-84 |
| USP1 | 0.401784 | 1.8E-72 |
| TMEM97 | 0.400268 | 2.41E-77 |
| MIS18A | 0.397314 | 1.28E-84 |
| SLBP | 0.395546 | 1.03E-50 |
| MCM6 | 0.395165 | 2.8E-70 |
| E2F1 | 0.394946 | 3.44E-73 |
| LSM4 | 0.394552 | 8.72E-83 |
| PTMA | 0.393167 | 0 |
| PTX3 | 0.392935 | 2.53E-34 |
| SPC25 | 0.391541 | 6.62E-56 |
| ALYREF | 0.391157 | 5.75E-74 |
| CENPX | 0.389287 | 7.07E-56 |
| LSM3 | 0.389055 | 6.92E-72 |
| KIF4A | 0.385394 | 7.45E-42 |
| DHFR | 0.385066 | 9.82E-49 |

| | | |
|---|---|---|
| SAE1 | 0.384231 | 1.29E-76 |
| SNRPB | 0.384169 | 1.97E-79 |
| ILF2 | 0.383857 | 1.02E-76 |
| RFC2 | 0.379827 | 6.36E-57 |
| PRKDC | 0.379733 | 8.88E-52 |
| DDX39A | 0.378353 | 2.34E-48 |
| CENPV | 0.378177 | 1.03E-47 |
| FBXO5 | 0.377647 | 5.89E-43 |
| NPW | 0.375115 | 2.1E-51 |
| HNRNPR | 0.374636 | 8.22E-93 |
| SRSF3 | 0.37243 | 7.7E-107 |
| SUPT16H | 0.3686 | 7.69E-60 |
| SNRPE | 0.365908 | 4.65E-75 |
| HJURP | 0.365417 | 8.71E-42 |
| KIF15 | 0.362172 | 7.06E-50 |
| BUB1 | 0.361148 | 9.12E-36 |
| DKK1 | 0.360537 | 1.39E-11 |
| NCAPG | 0.360395 | 1.66E-62 |
| PRRX1 | 0.358662 | 2.74E-97 |
| CDK4 | 0.357115 | 3.29E-44 |
| ASF1B | 0.356865 | 1.34E-56 |
| RPL39L | 0.356655 | 1.21E-66 |
| CHEK1 | 0.356117 | 2.74E-62 |
| RPA2 | 0.35497 | 1.91E-55 |
| SFPQ | 0.352928 | 2.9E-36 |
| GLRX5 | 0.352886 | 6.43E-36 |
| NMU | 0.350349 | 6.19E-22 |
| CDCA2 | 0.348406 | 8.24E-35 |
| SMC3 | 0.348325 | 3.7E-56 |
| CDC6 | 0.347858 | 1.2E-63 |
| SIVA1 | 0.347406 | 1.34E-57 |
| ERH | 0.346709 | 4.16E-76 |
| CNTLN | 0.345479 | 5.51E-58 |
| PSMC3 | 0.345456 | 3.61E-63 |
| KIF18A | 0.344942 | 3.86E-35 |
| LIG1 | 0.343319 | 1.26E-47 |
| RACGAP1 | 0.341261 | 4.38E-36 |
| CKAP5 | 0.340365 | 9.77E-30 |
| CDCA4 | 0.340221 | 1.95E-53 |
| GPSM2 | 0.339 | 7.21E-35 |
| SPC24 | 0.337079 | 5.89E-45 |
| DSN1 | 0.336943 | 6.01E-62 |
| EIF4EBP1 | 0.335256 | 1.43E-39 |
| HSPB11 | 0.334 | 2.17E-46 |
| PRRX2 | 0.333906 | 1.85E-62 |
| BANF1 | 0.331982 | 1.19E-64 |
| HP1BP3 | 0.331933 | 4.31E-24 |
| PHF19 | 0.329441 | 7.94E-50 |
| NME1 | 0.329077 | 1.54E-35 |
| LDHA | 0.328362 | 3.14E-61 |
| MCM2 | 0.32824 | 1.34E-55 |
| COMMD4 | 0.324947 | 8.73E-50 |
| HNRNPM | 0.324688 | 5.37E-54 |
| PCP4 | 0.323303 | 1.03E-11 |
| PSME2 | 0.322751 | 4.12E-37 |
| CHAF1A | 0.32259 | 4.42E-57 |

| | | |
|---|---|---|
| SNRPG | 0.322333 | 1.04E-47 |
| TMEM160 | 0.321974 | 1.86E-53 |
| BUB1B | 0.321653 | 1.76E-38 |
| CCDC34 | 0.321477 | 3.25E-47 |
| C1QL1 | 0.320008 | 7.69E-36 |
| NDE1 | 0.319457 | 1.19E-30 |
| NCL | 0.316356 | 3.7E-58 |
| H1FX | 0.316259 | 2.11E-41 |
| TUBA1A | 0.316057 | 6.67E-65 |
| RAC3 | 0.31592 | 4.86E-48 |
| TRIM59 | 0.313131 | 2.47E-29 |
| LMNB2 | 0.312407 | 1.25E-48 |
| HPF1 | 0.311642 | 1.97E-54 |
| PHGDH | 0.309174 | 2.6E-51 |
| PIF1 | 0.308297 | 1.25E-24 |
| POLD3 | 0.307689 | 3.24E-44 |
| TWIST1 | 0.307601 | 8.62E-40 |
| MSH6 | 0.306561 | 2.73E-39 |
| CBX3 | 0.30647 | 2.74E-44 |
| HSPA8 | 0.306165 | 7.31E-49 |
| CRABP2 | 0.304759 | 4.95E-52 |
| MAB21L2 | 0.304509 | 4.32E-62 |
| PCOLCE | 0.304355 | 1.27E-59 |
| NAP1L1 | 0.301817 | 7.65E-73 |
| HIST1H1A | 0.300684 | 9.47E-18 |
| IFI44L | 0.300237 | 8.11E-25 |
| RFC3 | 0.299461 | 2.27E-52 |
| DIAPH3 | 0.29829 | 2.82E-45 |
| PPIH | 0.298216 | 2.29E-48 |
| PTGES3 | 0.297612 | 1.59E-52 |
| SEPT7 | 0.297611 | 2.66E-52 |
| MRPL51 | 0.297034 | 4.28E-32 |
| B2M | 0.296901 | 1.16E-22 |
| HIST1H1D | 0.295309 | 4.58E-22 |
| COL1A1 | 0.295227 | 7.6E-104 |
| KIF14 | 0.293871 | 1.58E-29 |
| HMGN1 | 0.292608 | 3.82E-75 |
| CENPJ | 0.292477 | 7.87E-51 |
| WDR76 | 0.291526 | 2.09E-50 |
| DTL | 0.291313 | 3.57E-54 |
| ATAD5 | 0.289572 | 1.02E-56 |
| SAP30 | 0.289396 | 4.11E-31 |
| VRK1 | 0.288684 | 3.92E-49 |
| PSMC3IP | 0.28823 | 1.89E-50 |
| CNTRL | 0.287662 | 2.43E-32 |
| NME4 | 0.286939 | 3.79E-41 |
| HDGF | 0.286773 | 1.92E-17 |
| HNRNPAB | 0.286331 | 1.47E-31 |
| SRSF10 | 0.286201 | 1.89E-48 |
| CASP8AP2 | 0.285745 | 9.98E-43 |
| LBR | 0.285548 | 7.23E-22 |
| C12orf57 | 0.285278 | 4.95E-60 |
| HNRNPA1 | 0.28505 | 1.4E-191 |
| BRCA1 | 0.28505 | 6.61E-53 |
| CMC2 | 0.285006 | 2.05E-30 |
| MAD2L2 | 0.28486 | 1.88E-44 |

| | | |
|---|---|---|
| RHEB | 0.284553 | 1.32E-30 |
| MAGOHB | 0.283431 | 2.42E-39 |
| DNAJA1 | 0.282594 | 3.06E-39 |
| DCN | 0.282297 | 3.73E-08 |
| CTHRC1 | 0.281488 | 1.61E-28 |
| CDCA7 | 0.280545 | 1.66E-55 |
| ESCO2 | 0.27941 | 3.81E-40 |
| HMGXB4 | 0.279154 | 3.93E-42 |
| TPM4 | 0.279066 | 6.16E-37 |
| MCM10 | 0.278717 | 1.31E-48 |
| OIP5 | 0.278677 | 5.22E-43 |
| RBMX | 0.278353 | 5.66E-27 |
| TMEM106( | 0.278312 | 4.9E-30 |
| SNRNP40 | 0.277795 | 7.69E-39 |
| RPA3 | 0.277542 | 3.58E-32 |
| CACYBP | 0.277311 | 2.38E-32 |
| SET | 0.276982 | 3.2E-45 |
| ZNF714 | 0.276262 | 8.63E-43 |
| NCAPD2 | 0.275663 | 1.19E-27 |
| CEP55 | 0.273696 | 6.16E-32 |
| HNRNPA3 | 0.273257 | 1.51E-46 |
| SPDL1 | 0.273027 | 3.14E-32 |
| FAM83D | 0.272951 | 2.45E-26 |
| EMP3 | 0.272815 | 1.54E-49 |
| HACD3 | 0.27227 | 1.04E-32 |
| CCDC18 | 0.271984 | 8.97E-36 |
| TCEAL7 | 0.270303 | 3.26E-33 |
| ACAT2 | 0.269235 | 2.79E-25 |
| MAB21L1 | 0.26847 | 5.57E-34 |
| PLEKHO1 | 0.268242 | 1.03E-44 |
| CDK5RAP: | 0.267812 | 4.13E-33 |
| RBBP7 | 0.267737 | 8.37E-25 |
| GLO1 | 0.26669 | 8.95E-29 |
| H3F3A | 0.265904 | 5.7E-166 |
| PTMS | 0.265751 | 1.4E-11 |
| ITGAE | 0.264013 | 4.72E-27 |
| TRA2B | 0.263882 | 2.42E-36 |
| G2E3 | 0.263126 | 5.62E-22 |
| HNRNPH3 | 0.262812 | 5.14E-36 |
| COL1A2 | 0.262806 | 1.7E-81 |
| CNN3 | 0.262481 | 1.96E-53 |
| MZT2B | 0.261681 | 5.22E-44 |
| GPX8 | 0.260933 | 2.93E-36 |
| MGME1 | 0.260895 | 8.92E-36 |
| UQCC2 | 0.260393 | 7.26E-28 |
| IGF1 | 0.259687 | 6.17E-05 |
| MYBL2 | 0.259577 | 2.32E-32 |
| RPS20 | 0.259518 | 1.86E-84 |
| HMGA1 | 0.259297 | 3.44E-24 |
| RPL21 | 0.257522 | 2.6E-162 |
| FUS | 0.256548 | 2.13E-45 |
| ARHGAP1 | 0.256234 | 1.11E-31 |
| FIGNL1 | 0.255323 | 8.54E-41 |
| METRN | 0.254277 | 1.05E-21 |
| FANCL | 0.25426 | 1.66E-32 |
| XRCC2 | 0.254044 | 3.11E-43 |

| | | |
|---|---|---|
| RPLP0 | 0.254015 | 3.6E-104 |
| POC1A | 0.253778 | 2.64E-44 |
| HSP90AA1 | 0.25087 | 1.8E-30 |

| **Cluster 5** | | |
|---|---|---|
| Gene | LogFC | pVal |
| PRG4 | 2.737562 | 0 |
| TNMD | 0.955932 | 4.9E-100 |
| ITM2A | 0.905894 | 1.3E-131 |
| GAS2 | 0.878122 | 1.6E-236 |
| SFRP2 | 0.849199 | 8.5E-277 |
| MGP | 0.812647 | 2.8E-202 |
| CNMD | 0.70529 | 2.7E-172 |
| PTN | 0.677333 | 0 |
| C2orf40 | 0.661756 | 1.2E-165 |
| COL2A1 | 0.64337 | 5E-165 |
| PEG10 | 0.601759 | 6.7E-117 |
| TRPS1 | 0.587234 | 9.5E-149 |
| COL9A3 | 0.575323 | 4.7E-232 |
| OGN | 0.566324 | 6.5E-103 |
| POMC | 0.565634 | 6.85E-68 |
| COL1A1 | 0.558093 | 0 |
| GAS1 | 0.552749 | 2.2E-172 |
| NFIB | 0.522558 | 2.1E-193 |
| COL1A2 | 0.521193 | 0 |
| ZFHX4 | 0.506524 | 2E-113 |
| DLK1 | 0.501457 | 1.45E-59 |
| COL9A2 | 0.493537 | 3.2E-179 |
| SCX | 0.489468 | 2.74E-94 |
| MAB21L1 | 0.473989 | 2.9E-105 |
| COL11A1 | 0.456798 | 9.3E-103 |
| EBF3 | 0.439014 | 4.8E-92 |
| CDH11 | 0.435755 | 2.8E-132 |
| RGCC | 0.433513 | 2.12E-51 |
| SCG5 | 0.428249 | 4.33E-70 |
| PMP22 | 0.428053 | 5.8E-109 |
| NEFL | 0.427268 | 2.76E-29 |
| FN1 | 0.426184 | 4.5E-61 |
| COL9A1 | 0.422095 | 1.06E-83 |
| PRR16 | 0.419547 | 9.31E-72 |
| MEG3 | 0.417168 | 6.09E-90 |
| SRGAP1 | 0.413391 | 2.86E-76 |
| BOC | 0.412881 | 6.9E-108 |
| MIA | 0.411314 | 6.78E-69 |
| PDGFRA | 0.410225 | 4.9E-119 |
| RPS12 | 0.409313 | 0 |
| CAPN6 | 0.408089 | 1.66E-75 |
| MAB21L2 | 0.40274 | 8.5E-128 |
| RND3 | 0.395856 | 4.17E-64 |
| NPY | 0.387082 | 2.51E-22 |
| CPE | 0.386447 | 6.7E-209 |
| HMCN1 | 0.385351 | 2.08E-80 |
| ADAMTS6 | 0.383283 | 6.76E-57 |
| ELN | 0.38134 | 1.8E-90 |
| RFLNA | 0.38104 | 6.54E-83 |
| EIF3H | 0.377305 | 6.8E-155 |
| FGFR1 | 0.372763 | 2.75E-73 |
| PTX3 | 0.369344 | 1.25E-37 |
| PPIC | 0.367608 | 1.59E-74 |
| TSPO | 0.355331 | 5.2E-106 |

| | | |
|---|---|---|
| GRID2 | 0.353212 | 3.46E-40 |
| RPS11 | 0.349863 | 2.5E-281 |
| NDRG2 | 0.346002 | 4.57E-72 |
| CTHRC1 | 0.344028 | 1.98E-60 |
| FZD2 | 0.343619 | 2.39E-91 |
| FRZB | 0.342867 | 8.84E-61 |
| SCRG1 | 0.340955 | 1.78E-22 |
| CADM1 | 0.338981 | 2.94E-82 |
| RUNX1T1 | 0.338584 | 1.78E-70 |
| RNF175 | 0.338434 | 2.36E-73 |
| RPS19 | 0.338251 | 0 |
| SERTAD4- | 0.336154 | 3.2E-55 |
| FOXC1 | 0.335397 | 4.04E-55 |
| PCDH7 | 0.332669 | 7.03E-43 |
| EEF1D | 0.329577 | 1.9E-171 |
| HNRNPA1 | 0.328374 | 4.1E-267 |
| TPR | 0.325201 | 2.63E-34 |
| MEST | 0.321564 | 1.2E-143 |
| SOX5 | 0.320771 | 1.12E-47 |
| RPL18A | 0.320504 | 2E-305 |
| LHFPL6 | 0.316939 | 4.91E-53 |
| SNHG8 | 0.31674 | 8.69E-72 |
| GLI3 | 0.315838 | 9.27E-65 |
| NEFM | 0.313917 | 1.71E-15 |
| MEX3B | 0.313373 | 6.2E-58 |
| EMILIN1 | 0.312524 | 6.15E-91 |
| RPL15 | 0.312119 | 0 |
| IFI44L | 0.312033 | 4.1E-53 |
| RPSA | 0.311267 | 1.8E-267 |
| MDFI | 0.311038 | 9.8E-120 |
| NNAT | 0.310593 | 1.11E-92 |
| COL12A1 | 0.310245 | 3.09E-37 |
| IL11RA | 0.309497 | 1.2E-103 |
| FOXP1 | 0.309487 | 2.21E-29 |
| PDZRN4 | 0.308909 | 3.69E-65 |
| IGFBP5 | 0.308034 | 1.07E-44 |
| CNN3 | 0.307232 | 4.5E-108 |
| RPS27A | 0.304376 | 0 |
| CCND2 | 0.304245 | 2.74E-66 |
| PLAC9 | 0.303416 | 2.19E-42 |
| RPL21 | 0.303118 | 0 |
| DNM3OS | 0.300717 | 2.91E-60 |
| RPL22 | 0.300077 | 4.4E-280 |
| RPL13A | 0.299526 | 8.8E-281 |
| TPBG | 0.298233 | 2.81E-38 |
| PRELP | 0.297461 | 2.47E-46 |
| SERTAD4 | 0.295978 | 2.02E-52 |
| RPS13 | 0.295913 | 0 |
| COL8A2 | 0.295541 | 2E-63 |
| RPL19 | 0.294238 | 1.6E-293 |
| NPM1 | 0.293582 | 9.9E-267 |
| RPS3A | 0.292468 | 0 |
| RPL34 | 0.292234 | 5E-296 |
| CIRBP | 0.292133 | 6.6E-122 |
| RPL30 | 0.291719 | 0 |
| MEOX2 | 0.291659 | 1.31E-45 |

| | | |
|---|---|---|
| MARCKS | 0.29 | 1.2E-106 |
| C9orf3 | 0.289793 | 1.34E-45 |
| SOX9 | 0.289461 | 5.57E-46 |
| CKAP4 | 0.288949 | 7.74E-54 |
| RPL32 | 0.286389 | 0 |
| RPL6 | 0.286264 | 0 |
| RPLP1 | 0.285035 | 1E-302 |
| SLC8A1 | 0.283354 | 7E-48 |
| RPS16 | 0.282356 | 2.2E-272 |
| PCOLCE | 0.2822 | 2.05E-70 |
| NETO2 | 0.279982 | 8.47E-20 |
| NPW | 0.278357 | 4.77E-65 |
| KCNMA1 | 0.278143 | 4.8E-46 |
| COL8A1 | 0.276809 | 4.88E-22 |
| RPL10 | 0.276519 | 3.1E-275 |
| MFAP4 | 0.273609 | 3.77E-98 |
| EBPL | 0.272154 | 7.9E-66 |
| IGF2.1 | 0.271943 | 4.35E-44 |
| CD83 | 0.271747 | 1.9E-40 |
| CA10 | 0.271255 | 2.67E-44 |
| RPS24 | 0.270837 | 0 |
| RPL10A | 0.270086 | 5.8E-243 |
| RPL37 | 0.269758 | 8E-265 |
| RPL27A | 0.269538 | 4.1E-219 |
| NACA | 0.269328 | 3.3E-234 |
| RPS4X | 0.26764 | 3.9E-142 |
| TNFAIP8 | 0.26655 | 1.88E-40 |
| RPS23 | 0.26622 | 3.6E-286 |
| RPL11 | 0.266001 | 2.7E-253 |
| RGS2 | 0.265602 | 2.18E-31 |
| SELENOM | 0.265488 | 5.31E-56 |
| NOP53 | 0.264515 | 8.51E-65 |
| CRTAP | 0.263565 | 4.27E-61 |
| SETBP1 | 0.262401 | 2.66E-41 |
| KAT6B | 0.259828 | 4.75E-35 |
| RPL13 | 0.258002 | 1.1E-287 |
| EEF1A1 | 0.257861 | 1E-305 |
| NRXN1 | 0.257734 | 1.35E-23 |
| LGALS1 | 0.257491 | 1.2E-165 |
| GPM6B | 0.256755 | 4.6E-39 |
| CDC42EP5 | 0.255546 | 2.95E-91 |
| EEF1B2 | 0.254883 | 4.31E-84 |
| SOX6 | 0.25316 | 2.26E-39 |
| RPS8 | 0.252718 | 3.1E-284 |
| C1QL1 | 0.251946 | 1.03E-36 |
| RPLP0 | 0.251661 | 5.7E-183 |
| ISLR | 0.251269 | 2.35E-52 |

| **Cluster 6** | | |
|---|---|---|
| Gene | LogFC | pVal |
| COL21A1 | 1.43836 | 4.5E-267 |
| KRT17 | 1.366761 | 8.62E-75 |
| SLC26A7 | 1.193772 | 7.7E-157 |
| SPARCL1 | 1.184145 | 1.4E-149 |
| ISL1 | 1.181172 | 3.9E-205 |
| COL3A1 | 1.146317 | 0 |
| CRABP1 | 1.026399 | 0 |
| PRSS12 | 0.843786 | 4.6E-111 |
| NEFM | 0.831248 | 1.98E-60 |
| SELENOP | 0.830553 | 5.4E-172 |
| VIM | 0.796654 | 2.4E-260 |
| POSTN | 0.796211 | 1.44E-48 |
| CPE | 0.794298 | 0 |
| ZNF385D | 0.787914 | 6.4E-149 |
| IGF1 | 0.785842 | 4.9E-40 |
| OSR2 | 0.758493 | 7.9E-113 |
| LMO4 | 0.73612 | 2.1E-133 |
| UBE2E3 | 0.732971 | 1.2E-176 |
| SIM2 | 0.721441 | 4E-124 |
| NRP2 | 0.703043 | 1.71E-92 |
| SOX11 | 0.697251 | 4E-130 |
| TSHZ2 | 0.696883 | 8.6E-127 |
| LGALS1 | 0.693063 | 0 |
| TSC22D1 | 0.670024 | 2.58E-84 |
| SULF1 | 0.66951 | 2.7E-85 |
| NNAT | 0.663234 | 4.8E-222 |
| LUM | 0.658413 | 3.23E-33 |
| NTRK2 | 0.647378 | 4.6E-107 |
| COL5A2 | 0.642768 | 4.4E-129 |
| PRRX1 | 0.637416 | 4.5E-267 |
| TGFB2 | 0.63422 | 2.2E-114 |
| TBX2 | 0.631412 | 1.7E-120 |
| LRRC17 | 0.626151 | 1.1E-113 |
| SHOX2 | 0.624435 | 6.89E-85 |
| AKAP12 | 0.623215 | 2.02E-94 |
| AMD1 | 0.620787 | 1.5E-120 |
| TUBA1A | 0.610767 | 2.7E-148 |
| TSHZ1 | 0.604602 | 4.3E-102 |
| NELL2 | 0.600982 | 5.82E-79 |
| HAND2 | 0.597211 | 5.4E-104 |
| ID3 | 0.594788 | 2.2E-131 |
| IGFBP3 | 0.589066 | 1.37E-43 |
| TUBB2B | 0.584321 | 1.9E-108 |
| BASP1 | 0.541897 | 1.3E-172 |
| KCTD12 | 0.540164 | 7.9E-80 |
| MLLT11 | 0.537257 | 1.5E-103 |
| ID2 | 0.536379 | 1.58E-45 |
| MFAP4 | 0.534582 | 1.5E-148 |
| PTX3 | 0.523269 | 2.75E-38 |
| DKK2 | 0.504196 | 1.58E-64 |
| SFRP1 | 0.503815 | 8.3E-54 |
| CAPN6 | 0.500518 | 1.85E-57 |
| COX6C | 0.50005 | 7.7E-180 |
| RBP1 | 0.49545 | 2.07E-70 |

| | | |
|---|---|---|
| DCX | 0.495121 | 1.56E-66 |
| TCF4 | 0.491491 | 6.31E-95 |
| NFIA | 0.490972 | 1.1E-105 |
| IQGAP2 | 0.49071 | 5.57E-71 |
| MRPS6 | 0.484949 | 2.9E-119 |
| MIR99AHG | 0.482585 | 2.1E-104 |
| HOXA11 | 0.480866 | 4.12E-65 |
| NKAIN4 | 0.479655 | 1.54E-78 |
| ZNF503 | 0.478548 | 1.15E-77 |
| EBF1 | 0.475409 | 2.52E-90 |
| MGP | 0.475323 | 8.8E-109 |
| DOK6 | 0.474388 | 1.51E-66 |
| ANXA2 | 0.474276 | 5E-157 |
| NOVA1 | 0.472532 | 6.54E-75 |
| FIBIN | 0.463014 | 2.2E-44 |
| ZADH2 | 0.461378 | 1.82E-79 |
| CXCL14 | 0.455535 | 2.8E-14 |
| COL1A2 | 0.453883 | 0 |
| BMP5 | 0.44813 | 5.42E-67 |
| NFIB | 0.445436 | 3.9E-113 |
| IGFBP4 | 0.443763 | 9.38E-89 |
| CEP126 | 0.442835 | 6.19E-57 |
| COL1A1 | 0.440844 | 1.7E-274 |
| GLT8D2 | 0.4403 | 2.4E-99 |
| CTSC | 0.436885 | 1.95E-27 |
| PFN2 | 0.432646 | 2.1E-67 |
| IL11RA | 0.430059 | 7E-139 |
| FSTL1 | 0.429825 | 5.6E-82 |
| OLFM2 | 0.428411 | 4.91E-88 |
| PDE5A | 0.418073 | 4.67E-63 |
| C1orf54 | 0.413738 | 1.37E-96 |
| PDZRN4 | 0.412133 | 1.17E-69 |
| CDC42EP5 | 0.407939 | 3.4E-118 |
| MMP2 | 0.407415 | 2.35E-80 |
| CRABP2 | 0.401499 | 4.87E-77 |
| THBS4 | 0.401435 | 4.58E-35 |
| SH3BGRL3 | 0.401161 | 5.6E-77 |
| OPRK1 | 0.401046 | 2.7E-47 |
| OLFML3 | 0.39867 | 4.24E-86 |
| RNF175 | 0.398122 | 6.24E-75 |
| VCAN | 0.396264 | 2.53E-48 |
| NREP | 0.393196 | 7.22E-79 |
| C11orf96 | 0.385599 | 6.01E-46 |
| TMEM100 | 0.382493 | 7.32E-35 |
| HMGA2 | 0.378521 | 4.02E-65 |
| MEIS3 | 0.377666 | 1.46E-60 |
| MDK | 0.37534 | 6.5E-214 |
| MEST | 0.373522 | 7.24E-49 |
| MFAP2 | 0.365919 | 2.54E-89 |
| PBX3 | 0.365715 | 7.26E-37 |
| IFI27L2 | 0.361809 | 7.14E-49 |
| CCDC144N | 0.361008 | 6.16E-42 |
| ID1 | 0.359416 | 3.01E-53 |
| PRRX2 | 0.35941 | 1.09E-65 |
| CITED1 | 0.358185 | 1.54E-35 |
| SLC5A3 | 0.357402 | 6.3E-51 |

| | | |
|---|---|---|
| COL6A3 | 0.354123 | 9.79E-52 |
| CXCL12 | 0.352628 | 4.36E-50 |
| C12orf57 | 0.351946 | 3.1E-104 |
| MEIS1 | 0.350858 | 2.11E-55 |
| GPC3 | 0.350082 | 1.71E-41 |
| SPRY1 | 0.348595 | 3.34E-21 |
| MAB21L2 | 0.348373 | 1.54E-45 |
| PPP3CA | 0.344029 | 5.82E-37 |
| EMILIN1 | 0.343762 | 5.38E-79 |
| ZIC2 | 0.340225 | 6.64E-28 |
| EPB41L2 | 0.339546 | 3.27E-49 |
| TBX3 | 0.336736 | 5.77E-45 |
| MAP2 | 0.335132 | 2.01E-51 |
| SESTD1 | 0.334073 | 1.45E-38 |
| CALD1 | 0.332349 | 2.9E-107 |
| TPM4 | 0.330015 | 6.29E-53 |
| IFITM3 | 0.328294 | 5.51E-74 |
| FBLN1 | 0.327646 | 6.93E-68 |
| PRICKLE1 | 0.327497 | 9.19E-51 |
| SYNPO2 | 0.326867 | 1.81E-50 |
| PLK2 | 0.325502 | 3.19E-37 |
| WLS | 0.32068 | 3.81E-48 |
| CD81 | 0.320051 | 6.9E-149 |
| LITAF | 0.319664 | 2.35E-56 |
| HNRNPA1 | 0.319529 | 7.2E-230 |
| TWIST2 | 0.314701 | 1.76E-45 |
| PGM5-AS1 | 0.312454 | 5.92E-16 |
| ETV1 | 0.312114 | 4.91E-35 |
| SLIT3 | 0.310443 | 1.35E-48 |
| PDGFRA | 0.309991 | 1.19E-60 |
| CREB5 | 0.309654 | 1.72E-36 |
| KCNQ1OT | 0.307957 | 4.72E-64 |
| RHOU | 0.307231 | 1.11E-35 |
| EDNRA | 0.305681 | 3.61E-35 |
| MARCKSL | 0.305109 | 3.5E-148 |
| PKIG | 0.301808 | 9.73E-35 |
| GSN | 0.301344 | 3.02E-48 |
| ZEB2 | 0.301022 | 8.57E-28 |
| HOXA10 | 0.300088 | 2.94E-36 |
| RGS10 | 0.299729 | 1.67E-28 |
| PLPPR3 | 0.299312 | 6.03E-48 |
| CTHRC1 | 0.299123 | 5.46E-29 |
| SESN3 | 0.298829 | 3.59E-41 |
| LIMS1 | 0.29514 | 2.36E-35 |
| SIX1 | 0.294339 | 2.13E-36 |
| NPM1 | 0.293902 | 5.4E-197 |
| GABRA2 | 0.293841 | 9.08E-37 |
| LDB2 | 0.292618 | 1.11E-39 |
| DUSP4 | 0.291922 | 3.96E-43 |
| TTN | 0.291365 | 6.01E-37 |
| PCDH18 | 0.291214 | 5.01E-43 |
| CDK6 | 0.290496 | 2E-35 |
| TSPO | 0.289927 | 1.75E-51 |
| ACTG1 | 0.289346 | 5E-111 |
| RPS20 | 0.28879 | 3.5E-151 |
| B2M | 0.287471 | 1.42E-41 |

| | | |
|---|---|---|
| CTBP2 | 0.287051 | 7.55E-35 |
| PLAC9 | 0.287032 | 1.79E-25 |
| RBFOX2 | 0.285773 | 5.43E-51 |
| FAM213A | 0.284598 | 2.19E-31 |
| CCND1 | 0.284416 | 6.2E-44 |
| MAP1B | 0.283676 | 1.65E-38 |
| SPTBN1 | 0.281469 | 2.17E-35 |
| TOX3 | 0.28117 | 5.97E-21 |
| SEMA3A | 0.280891 | 8.69E-27 |
| IFITM2 | 0.280783 | 2.54E-36 |
| TTC3 | 0.277747 | 2.87E-61 |
| MARCKS | 0.276438 | 1.2E-143 |
| MDFI | 0.273333 | 1.44E-85 |
| RND3 | 0.272849 | 8.91E-23 |
| CALM2 | 0.272512 | 4.2E-47 |
| TMTC1 | 0.272364 | 8.68E-44 |
| FAM110B | 0.271667 | 1.84E-29 |
| PCP4 | 0.271555 | 0.015249 |
| GJA1 | 0.270765 | 1.31E-30 |
| EFEMP1 | 0.26947 | 2.9E-24 |
| CTSK | 0.268273 | 8.67E-46 |
| PLEKHO1 | 0.267626 | 2.82E-47 |
| ANGPT1 | 0.26698 | 2.89E-15 |
| CCDC141 | 0.266765 | 4.75E-41 |
| PCDH10 | 0.266709 | 7.93E-28 |
| PDE1A | 0.266437 | 2.95E-27 |
| ITGA4 | 0.265585 | 6.17E-33 |
| NPY1R | 0.2655 | 4.96E-18 |
| FUT8 | 0.263269 | 2.3E-28 |
| MAGED2 | 0.26222 | 1.13E-71 |
| PTN | 0.26167 | 1.23E-90 |
| FXYD6 | 0.260604 | 3.23E-33 |
| SOCS1 | 0.257293 | 1.51E-29 |
| NUPR1 | 0.257165 | 2.38E-27 |
| CCL2 | 0.25612 | 9.01E-12 |
| HGF | 0.255547 | 3.52E-30 |
| STMN1 | 0.254674 | 1.8E-102 |
| CIRBP | 0.254602 | 9.47E-97 |
| COL6A2 | 0.25354 | 2.79E-59 |
| PDZRN3 | 0.253306 | 1.45E-38 |
| PRKG1 | 0.252388 | 7.36E-32 |
| ZNF428 | 0.250627 | 5.86E-54 |

| Cluster 7 | | |
|---|---|---|
| Gene | LogFC | pVal |
| NEFL | 1.164986 | 8.13E-96 |
| CRABP1 | 0.944004 | 0 |
| NFIB | 0.861157 | 9.3E-281 |
| PTN | 0.856483 | 1.6E-280 |
| GAS2 | 0.839707 | 3E-131 |
| NEFM | 0.802196 | 1.77E-46 |
| FOXP1 | 0.800552 | 2.2E-123 |
| LHX9 | 0.740281 | 6.8E-116 |
| NPY | 0.722159 | 3.46E-39 |
| MGP | 0.699213 | 1.4E-121 |
| GAS1 | 0.680756 | 6.3E-158 |
| PRRX1 | 0.674381 | 4.1E-170 |
| CD83 | 0.651469 | 9.3E-99 |
| MARCKS | 0.624556 | 7.2E-198 |
| GRID2 | 0.623568 | 1.61E-74 |
| TRPS1 | 0.617066 | 6E-112 |
| COL1A1 | 0.602312 | 1E-298 |
| MAB21L2 | 0.594469 | 1.2E-168 |
| LHFPL6 | 0.592169 | 1.5E-102 |
| RND3 | 0.576579 | 8.65E-69 |
| LGALS1 | 0.575578 | 0 |
| RGCC | 0.56941 | 2.13E-70 |
| PRR16 | 0.532929 | 2.18E-83 |
| COL14A1 | 0.529965 | 6.31E-55 |
| RAP2B | 0.502695 | 3.28E-61 |
| HNRNPA1 | 0.488307 | 0 |
| RPS12 | 0.469262 | 0 |
| EBF3 | 0.456675 | 7.9E-80 |
| PTX3 | 0.451828 | 6.3E-46 |
| COL1A2 | 0.437417 | 4E-240 |
| ELN | 0.429774 | 2.16E-67 |
| FAM181B | 0.428276 | 6.83E-69 |
| IGFBP2 | 0.417724 | 2.2E-100 |
| EIF3H | 0.410741 | 7E-137 |
| FLRT2 | 0.406143 | 2.28E-46 |
| COL3A1 | 0.40537 | 4.54E-64 |
| GLI3 | 0.401295 | 1.08E-61 |
| PCDH8 | 0.39238 | 3.92E-48 |
| EEF1D | 0.391524 | 9.1E-194 |
| RPSA | 0.382747 | 1.8E-261 |
| BOC | 0.379967 | 1.76E-67 |
| RPS19 | 0.379746 | 0 |
| RPL18A | 0.378284 | 6.1E-289 |
| TSPO | 0.37632 | 2.02E-92 |
| FOXP2 | 0.37482 | 2.88E-49 |
| PBX3 | 0.371888 | 3.28E-55 |
| RPL13A | 0.367252 | 0 |
| NBL1 | 0.365022 | 1.56E-42 |
| MDFIC | 0.360717 | 2.22E-46 |
| NPM1 | 0.3536 | 3E-269 |
| KCTD12 | 0.352181 | 4.91E-50 |
| RPS11 | 0.347983 | 1.8E-243 |
| IFI44L | 0.347348 | 2.72E-61 |
| TNFAIP8 | 0.344547 | 3.02E-48 |

| | | |
|---|---|---|
| COL11A1 | 0.344056 | 4.49E-53 |
| FABP5 | 0.34171 | 2.45E-68 |
| RERG | 0.338884 | 1.12E-44 |
| RPL37 | 0.338473 | 8.1E-275 |
| RPL19 | 0.338295 | 2.5E-294 |
| RPL21 | 0.33435 | 8.6E-287 |
| RPS13 | 0.331744 | 4E-297 |
| MEX3B | 0.33135 | 7.08E-44 |
| COL5A2 | 0.3286 | 5.3E-45 |
| RPLP0 | 0.326308 | 5.1E-216 |
| FGFR1 | 0.324223 | 1.53E-57 |
| GDF10 | 0.323534 | 6.62E-36 |
| CDH11 | 0.323417 | 1.88E-72 |
| RPL15 | 0.323269 | 4E-305 |
| CCND2 | 0.321767 | 2.65E-49 |
| SCX | 0.31695 | 1.22E-39 |
| RPLP1 | 0.316064 | 1.6E-291 |
| EIF3F | 0.315042 | 5.1E-110 |
| RPL6 | 0.314716 | 1.4E-272 |
| ALCAM | 0.314085 | 1.15E-21 |
| SMOC2 | 0.313848 | 1.81E-32 |
| RPL27A | 0.312699 | 4.8E-211 |
| TWIST1 | 0.31237 | 9.6E-32 |
| RPL22 | 0.311019 | 1.2E-190 |
| NEXN | 0.310088 | 2.28E-37 |
| SULF2 | 0.307285 | 3.88E-29 |
| FBLN1 | 0.305556 | 2.09E-56 |
| EEF1B2 | 0.305311 | 5.47E-95 |
| RNF175 | 0.305159 | 2.7E-54 |
| RPS3A | 0.305048 | 2.4E-254 |
| RPL32 | 0.303556 | 1.1E-253 |
| EIF3E | 0.303502 | 1.4E-99 |
| TOX3 | 0.303087 | 1.57E-30 |
| RPL23 | 0.301684 | 3.4E-171 |
| GALNTL6 | 0.301519 | 3.21E-39 |
| MAB21L1 | 0.301056 | 8.19E-31 |
| EMILIN1 | 0.300199 | 8.33E-59 |
| NACA | 0.299825 | 2.4E-202 |
| CRABP2 | 0.296346 | 8.93E-60 |
| RPL34 | 0.295903 | 6.6E-228 |
| GLT8D2 | 0.295386 | 3.78E-43 |
| RPS16 | 0.295014 | 5.3E-193 |
| CIRBP | 0.294842 | 2.9E-106 |
| RPS8 | 0.294741 | 2.3E-240 |
| IGFBP4 | 0.294324 | 7.5E-59 |
| RPS25 | 0.293277 | 2.1E-211 |
| C3orf58 | 0.293056 | 9.14E-30 |
| RPL13 | 0.291885 | 7.8E-213 |
| RPL7 | 0.290319 | 2.1E-179 |
| IL11RA | 0.289839 | 2.93E-68 |
| RPS27A | 0.289637 | 2.1E-258 |
| RUNX1T1 | 0.287033 | 4.73E-38 |
| BCL11A | 0.286781 | 2.02E-26 |
| TPM2 | 0.286442 | 9.1E-114 |
| NRXN1 | 0.28596 | 5.93E-23 |
| AP1S2 | 0.285238 | 6.43E-22 |

| | | |
|---|---|---|
| NOP53 | 0.284703 | 7.63E-53 |
| RPL4 | 0.284392 | 5.4E-158 |
| RPL24 | 0.282637 | 3.9E-185 |
| BASP1 | 0.28184 | 2.23E-62 |
| SCG5 | 0.281838 | 2.04E-24 |
| EBF1 | 0.281071 | 3.64E-24 |
| RPL10A | 0.280768 | 2.7E-160 |
| SNHG8 | 0.279459 | 2.91E-45 |
| EPHA4 | 0.27718 | 2.14E-25 |
| ZIC2 | 0.276712 | 3.38E-26 |
| RPL11 | 0.275417 | 3.6E-210 |
| NFIX | 0.274842 | 9.69E-39 |
| RPL14 | 0.272943 | 4.2E-181 |
| MGST1 | 0.272677 | 6.85E-26 |
| FBN2 | 0.27255 | 2.36E-37 |
| RPS27 | 0.271461 | 1.8E-217 |
| RPS20 | 0.270332 | 1.7E-124 |
| EPB41L2 | 0.269739 | 1.04E-26 |
| NPW | 0.26905 | 8.26E-48 |
| RPL30 | 0.268251 | 1.1E-228 |
| FZD2 | 0.267995 | 1.27E-48 |
| NNAT | 0.266886 | 3.35E-58 |
| RPS15 | 0.265432 | 2.6E-185 |
| RSL1D1 | 0.263895 | 2.8E-39 |
| ZFP64 | 0.262626 | 6.01E-32 |
| LTBP1 | 0.261147 | 2.04E-22 |
| MFAP2 | 0.259264 | 1.23E-46 |
| SCRG1 | 0.259208 | 2.91E-11 |
| AUTS2 | 0.258764 | 7E-32 |
| KAT6B | 0.257695 | 2.52E-27 |
| PRRX2 | 0.25754 | 4.33E-34 |
| MLLT3 | 0.256852 | 1.48E-26 |
| RPL39 | 0.256458 | 2.7E-176 |
| RPS24 | 0.255445 | 5.4E-196 |
| RPL23A | 0.253407 | 6.2E-167 |
| EBPL | 0.252562 | 9.22E-40 |
| MPDZ | 0.25221 | 6.54E-26 |
| PRSS23 | 0.250746 | 3.55E-21 |
| RPS9 | 0.25056 | 7.1E-177 |
| ITM2A | 0.250395 | 1.17E-11 |
| SERTAD4 | 0.25003 | 3.26E-27 |

| Cluster 8 | | |
|---|---|---|
| Gene | LogFC | pVal |
| KRT19 | 1.814878 | 0 |
| CKB | 1.78116 | 0 |
| BNIP3 | 1.75586 | 0 |
| PGK1 | 1.405863 | 0 |
| ENO1 | 1.337875 | 0 |
| NDUFA4L2 | 1.133272 | 5.2E-103 |
| PKM | 1.087311 | 0 |
| LDHA | 1.074249 | 0 |
| TPI1 | 1.067454 | 0 |
| ANXA4 | 1.057146 | 0 |
| C4orf3 | 1.005532 | 0 |
| PTGDS | 1.002644 | 4.22E-39 |
| SLC2A1 | 0.994874 | 4E-178 |
| ODC1 | 0.985744 | 1.1E-188 |
| ATP1B1 | 0.920045 | 0 |
| GAPDH | 0.900445 | 0 |
| TMEM176A | 0.898796 | 1.1E-265 |
| EPCAM | 0.89847 | 0 |
| CAMK2N1 | 0.894849 | 0 |
| SLC16A3 | 0.887598 | 2.6E-227 |
| AGPAT2 | 0.887053 | 0 |
| PLOD2 | 0.872116 | 1.4E-261 |
| FAM162A | 0.867529 | 2.1E-290 |
| S100A14 | 0.85591 | 2E-137 |
| CITED2 | 0.843522 | 2.6E-255 |
| IGFBP2 | 0.843254 | 1.4E-168 |
| IRX3 | 0.835331 | 5.8E-144 |
| CLEC18A | 0.833457 | 7.1E-212 |
| MIF | 0.822151 | 0 |
| HPN | 0.817588 | 0 |
| P4HA1 | 0.808749 | 3.9E-246 |
| RHOB | 0.800416 | 1.5E-195 |
| CDKN1C | 0.796725 | 5.7E-134 |
| CLEC18B | 0.792199 | 4.4E-215 |
| SERPINE2 | 0.789636 | 1.1E-202 |
| SMS | 0.753868 | 2.3E-239 |
| TMEM176B | 0.75293 | 1.1E-249 |
| SPP1 | 0.741147 | 7E-138 |
| APOE | 0.7396 | 1.6E-251 |
| MPC2 | 0.734219 | 0 |
| C1QTNF12 | 0.729594 | 8.6E-162 |
| LGALS2 | 0.727981 | 1.3E-206 |
| GPI | 0.723025 | 3.6E-190 |
| EMID1 | 0.722059 | 3.7E-221 |
| CD24 | 0.712331 | 1E-307 |
| FTL | 0.711022 | 0 |
| RBPMS | 0.706206 | 8.9E-284 |
| JUNB | 0.690998 | 2.3E-137 |
| ENO2 | 0.687142 | 1.2E-158 |
| EZR | 0.685908 | 1.3E-224 |
| KCNJ15 | 0.68175 | 1E-191 |
| NPC2 | 0.681231 | 0 |
| PPP1R16A | 0.680803 | 1.7E-152 |
| SMIM24 | 0.678061 | 3E-245 |

| | | |
|---|---|---|
| CLDN7 | 0.6707 | 2.3E-234 |
| CYBA | 0.666719 | 7E-295 |
| SLC3A1 | 0.653497 | 9.2E-162 |
| NR2F6 | 0.653432 | 9E-295 |
| VDAC2 | 0.652497 | 4.4E-268 |
| PDGFA | 0.648008 | 4.4E-156 |
| CLDN3 | 0.64472 | 5.5E-278 |
| PDZK1 | 0.643882 | 5.4E-228 |
| AK4 | 0.642672 | 5E-185 |
| BSG | 0.638547 | 0 |
| EMX2 | 0.63489 | 4.7E-175 |
| ALDOA | 0.632171 | 2.9E-186 |
| NDRG1 | 0.630874 | 1.2E-113 |
| PPP1R1A | 0.625982 | 1.6E-212 |
| BCAM | 0.624344 | 3E-297 |
| SPINT2 | 0.623652 | 0 |
| SLC44A4 | 0.610081 | 1.2E-202 |
| PCBD1 | 0.608954 | 0 |
| SDC4 | 0.608187 | 2.9E-158 |
| FBXO17 | 0.604898 | 2.9E-205 |
| TMEM256 | 0.60219 | 1.8E-169 |
| SLC27A2 | 0.600884 | 4.3E-127 |
| RAB25 | 0.600327 | 2.2E-143 |
| CLDN4 | 0.600304 | 1.5E-144 |
| FXYD2 | 0.597058 | 6.8E-160 |
| PFKP | 0.596157 | 3.2E-140 |
| FMO1 | 0.590789 | 4.1E-117 |
| FTH1 | 0.585045 | 0 |
| S100A1 | 0.585036 | 2.32E-77 |
| HES4 | 0.584086 | 6.6E-132 |
| CCDC198 | 0.582888 | 5.8E-163 |
| CEBPD | 0.581534 | 5.2E-168 |
| IGFBP7 | 0.581102 | 6.6E-277 |
| CSTB | 0.580516 | 2E-172 |
| PPP1R14C | 0.577787 | 2.5E-141 |
| PLEKHA1 | 0.574621 | 8.7E-188 |
| CLIC1 | 0.574064 | 0 |
| HOOK1 | 0.574007 | 3.6E-166 |
| SNHG7 | 0.571026 | 4.82E-98 |
| ID4 | 0.570569 | 2.35E-83 |
| ADAMTS1 | 0.570209 | 8.79E-92 |
| NEU1 | 0.558785 | 2.3E-144 |
| PGAM1 | 0.557919 | 1E-241 |
| SMIM1 | 0.557642 | 2.1E-143 |
| CD151 | 0.555518 | 1.3E-203 |
| UGT2B7 | 0.555437 | 5.7E-157 |
| EPS8L2 | 0.552446 | 2.2E-175 |
| EIF1 | 0.550902 | 2.5E-261 |
| TPD52L1 | 0.548282 | 2.68E-59 |
| ARID3A | 0.547221 | 1.1E-162 |
| ANKRD37 | 0.545746 | 1.8E-125 |
| APLP2 | 0.545017 | 2.3E-189 |
| LINC01781 | 0.54189 | 6.2E-144 |
| ITM2B | 0.535429 | 0 |
| IER2 | 0.534727 | 2.6E-156 |
| TMEM92 | 0.534569 | 4.05E-82 |

| | | |
|---|---|---|
| CHCHD10 | 0.533089 | 1.1E-148 |
| HINT1 | 0.530486 | 0 |
| ACAA2 | 0.526482 | 8.6E-225 |
| SLC39A4 | 0.524281 | 2.2E-80 |
| CYB5A | 0.52397 | 1.7E-233 |
| GPR146 | 0.520505 | 6.68E-56 |
| MSRB1 | 0.519205 | 2E-145 |
| CYSTM1 | 0.518198 | 1.2E-187 |
| ACOT13 | 0.515972 | 3.8E-133 |
| CD9 | 0.511435 | 3.2E-155 |
| TSTD1 | 0.510701 | 1.55E-93 |
| TMCC1 | 0.510021 | 3.6E-101 |
| TMEM141 | 0.50889 | 9.8E-133 |
| PAX8 | 0.507759 | 1.7E-165 |
| UNCX | 0.506053 | 3.79E-69 |
| C1orf43 | 0.504118 | 5.6E-171 |
| RIDA | 0.503577 | 1.3E-179 |
| MARVELD | 0.501137 | 2.4E-121 |
| BRI3 | 0.498929 | 4.7E-190 |
| CCNG1 | 0.49662 | 9.7E-150 |
| TMBIM6 | 0.495443 | 0 |
| RIT1 | 0.495235 | 3.2E-108 |
| MYO6 | 0.491795 | 1.5E-132 |
| DUSP9 | 0.491705 | 1.15E-82 |
| HNF1B | 0.490775 | 4.2E-131 |
| CCNB1IP1 | 0.490045 | 1.8E-140 |
| KRT18 | 0.487992 | 1.8E-177 |
| ZFAS1 | 0.484502 | 2.8E-128 |
| DARS | 0.483374 | 5.6E-140 |
| F10 | 0.480726 | 1.4E-105 |
| CXXC5 | 0.480699 | 1.7E-198 |
| MFSD10 | 0.478927 | 8E-151 |
| FGFR3 | 0.477005 | 9.2E-118 |
| AIG1 | 0.473833 | 5.6E-195 |
| CD46 | 0.47183 | 3.4E-170 |
| ATP6V1F | 0.471657 | 1.2E-186 |
| MIR210HG | 0.467176 | 1.2E-128 |
| SYAP1 | 0.466553 | 3.5E-122 |
| VEGFB | 0.463478 | 3.7E-153 |
| POU3F3 | 0.461359 | 2.3E-111 |
| CLDN6 | 0.459041 | 1.12E-64 |
| EGFL7 | 0.458879 | 2.08E-96 |
| LHX1 | 0.457757 | 1.3E-101 |
| KRT8 | 0.454733 | 3E-181 |
| ENPP3 | 0.454683 | 1.7E-105 |
| GPX4 | 0.454358 | 4.2E-189 |
| CYS1 | 0.453608 | 1.3E-152 |
| VEGFA | 0.452944 | 3.1E-122 |
| PLIN2 | 0.452404 | 4.26E-74 |
| CALM3 | 0.451499 | 1.6E-165 |
| KLF6 | 0.451166 | 1.05E-74 |
| CLU | 0.448123 | 2.5E-179 |
| DPP4 | 0.443386 | 2.7E-117 |
| CLEC18C | 0.442899 | 4.69E-91 |
| QPRT | 0.43945 | 1.9E-148 |
| PRSS8 | 0.437621 | 3.7E-118 |

| | | |
|---|---|---|
| JUN | 0.437289 | 7.51E-81 |
| FAM120AC | 0.436864 | 1.02E-66 |
| TXNIP | 0.431333 | 2.7E-95 |
| FLRT3 | 0.430725 | 8E-106 |
| KDELR1 | 0.427716 | 3.9E-191 |
| IMPA2 | 0.426937 | 1.3E-146 |
| GLYATL1 | 0.425489 | 5.08E-69 |
| LGMN | 0.423086 | 5.6E-105 |
| BIN1 | 0.421438 | 4.5E-117 |
| RTN4 | 0.420106 | 9.9E-177 |
| ASAH1 | 0.418677 | 8.8E-112 |
| PNRC1 | 0.4186 | 7.1E-157 |
| CARHSP1 | 0.415347 | 2.2E-125 |
| RNF181 | 0.412928 | 1.4E-148 |
| TOMM20 | 0.412814 | 1.4E-149 |
| RNASET2 | 0.412074 | 1.95E-83 |
| MXRA7 | 0.410784 | 2.61E-97 |
| DDIT4 | 0.410424 | 1.69E-65 |
| NEAT1 | 0.4095 | 5.57E-91 |
| VKORC1 | 0.408816 | 2.2E-138 |
| XBP1 | 0.407296 | 8.5E-102 |
| CRACR2B | 0.407147 | 7.64E-79 |
| MGST3 | 0.405241 | 2.1E-200 |
| SPINT1 | 0.404354 | 1.5E-137 |
| SLC9A3R1 | 0.403845 | 2.8E-150 |
| ATP5IF1 | 0.403729 | 1.5E-254 |
| SINHCAF | 0.402442 | 3.7E-103 |
| SLC6A8 | 0.402255 | 6.01E-36 |
| STX3 | 0.400818 | 2.2E-118 |
| LAMTOR5 | 0.395906 | 3.1E-177 |
| LAPTM4B | 0.394832 | 1.1E-139 |
| CRYAB | 0.394593 | 5.75E-37 |
| DYNC2LI1 | 0.393933 | 3.2E-111 |
| TXN | 0.393373 | 1.6E-151 |
| LSR | 0.392343 | 4.5E-113 |
| S100A10 | 0.392112 | 2.4E-155 |
| PCBP1 | 0.391743 | 5.4E-165 |
| LINC01320 | 0.391177 | 3.54E-84 |
| VIL1 | 0.391099 | 5.8E-102 |
| TSPAN4 | 0.391096 | 9.71E-98 |
| DPCD | 0.390809 | 8.1E-107 |
| DMTN | 0.389036 | 1.26E-94 |
| COL18A1 | 0.388972 | 4.3E-123 |
| MT-ND6 | 0.388858 | 4.75E-66 |
| PURPL | 0.386705 | 4.29E-79 |
| ALKBH7 | 0.385588 | 3.2E-118 |
| HOTAIRM1 | 0.384537 | 2.85E-54 |
| JAG1 | 0.382132 | 2.46E-91 |
| INSIG2 | 0.381493 | 7.43E-84 |
| ERBB3 | 0.381207 | 2.9E-102 |
| MRPS36 | 0.379406 | 3.8E-176 |
| PERP | 0.377952 | 6.55E-78 |
| SNCA | 0.377628 | 8.62E-71 |
| SKAP2 | 0.37718 | 9.59E-94 |
| DAB2 | 0.376065 | 8.1E-107 |
| SLC37A4 | 0.371973 | 5.5E-103 |

| | | |
|---|---|---|
| HMGCL | 0.370868 | 2.2E-110 |
| C19orf33 | 0.370172 | 3.1E-55 |
| FJX1 | 0.369657 | 1.35E-53 |
| VAT1 | 0.369467 | 9.8E-105 |
| ERBB4 | 0.369316 | 8.7E-71 |
| CIDEB | 0.368813 | 8.59E-85 |
| TINAG | 0.368363 | 7.07E-78 |
| AMN | 0.368262 | 8.3E-109 |
| C11orf54 | 0.368197 | 1.4E-114 |
| DDT | 0.367686 | 6.1E-112 |
| DPY30 | 0.367543 | 1.4E-138 |
| BNIP3L | 0.367448 | 5.3E-119 |
| WDR54 | 0.366722 | 1.33E-89 |
| CYP27A1 | 0.366503 | 6.55E-73 |
| CUBN | 0.360849 | 2.26E-75 |
| DBI | 0.360818 | 7.3E-151 |
| CD68 | 0.360739 | 5.02E-72 |
| CXADR | 0.360733 | 4.8E-106 |
| LGI4 | 0.359885 | 4.15E-74 |
| ENTPD2 | 0.35837 | 7.41E-82 |
| HOOK2 | 0.358125 | 1.91E-91 |
| GCHFR | 0.357335 | 1.04E-71 |
| TUBB2B | 0.353195 | 2.05E-65 |
| ZFP36 | 0.352534 | 3.68E-61 |
| PKN1 | 0.352363 | 4.5E-100 |
| COBLL1 | 0.351525 | 1.23E-68 |
| DSP | 0.35134 | 2.08E-75 |
| LRRC19 | 0.350498 | 4.4E-83 |
| LINC02381 | 0.350471 | 2.65E-78 |
| RPL41 | 0.350346 | 2E-194 |
| POLR1D | 0.350223 | 9.9E-104 |
| PDLIM1 | 0.349883 | 6.82E-67 |
| EIF4A2 | 0.348889 | 2.3E-125 |
| DCDC2 | 0.347057 | 2.12E-90 |
| FBRSL1 | 0.347019 | 1.46E-91 |
| RNF5 | 0.346497 | 2.1E-120 |
| HES1 | 0.346455 | 2.84E-32 |
| PPFIBP1 | 0.346237 | 2.41E-49 |
| TMEM134 | 0.345527 | 2.9E-101 |
| CD63 | 0.345452 | 3.2E-225 |
| PMM1 | 0.344997 | 2.9E-79 |
| PRR13 | 0.344984 | 1.1E-115 |
| ELSPBP1 | 0.34402 | 3.62E-52 |
| TUBB4B | 0.340884 | 1.8E-151 |
| TSPAN33 | 0.340796 | 3.12E-88 |
| SLC6A13 | 0.340703 | 2.88E-65 |
| GAL3ST1 | 0.339493 | 2.3E-102 |
| IRX5 | 0.339225 | 2.53E-47 |
| MSMO1 | 0.338986 | 2.94E-41 |
| NOP10 | 0.338049 | 4.3E-134 |
| SLC39A1 | 0.337116 | 7.16E-82 |
| ARMT1 | 0.334792 | 9.13E-89 |
| MDH1 | 0.333259 | 1.5E-141 |
| RASSF7 | 0.332487 | 5.79E-83 |
| CHPT1 | 0.332388 | 1.55E-95 |
| NDUFA5 | 0.331451 | 2.5E-148 |

| | | |
|---|---|---|
| TMEM150/ | 0.331345 | 3.1E-85 |
| HOXB7 | 0.330484 | 8.43E-85 |
| GNG11 | 0.330374 | 5.43E-76 |
| TKT | 0.329728 | 1.36E-87 |
| BHLHE40 | 0.32967 | 7.06E-72 |
| DSC2 | 0.329403 | 1.9E-84 |
| CCDC151 | 0.326409 | 3.79E-67 |
| ERO1A | 0.326361 | 7.19E-75 |
| JUND | 0.325782 | 3.1E-106 |
| SH3YL1 | 0.325703 | 2.65E-80 |
| GSTP1 | 0.325256 | 6.7E-137 |
| GALNT14 | 0.32308 | 1.92E-78 |
| LMO4 | 0.323042 | 3.4E-44 |
| PRKAB1 | 0.323025 | 2.69E-74 |
| PLLP | 0.322639 | 5.45E-66 |
| RTN3 | 0.32009 | 5.08E-96 |
| TXNDC17 | 0.319506 | 1.46E-89 |
| HSBP1L1 | 0.319129 | 2.61E-67 |
| LDHB | 0.318864 | 2.8E-186 |
| TCIM | 0.31805 | 8.05E-43 |
| RNF7 | 0.318 | 1.1E-132 |
| FOS | 0.317627 | 5.3E-46 |
| RPL36 | 0.317537 | 3.5E-160 |
| UQCR10 | 0.316363 | 1.8E-167 |
| TCEA3 | 0.315334 | 3.33E-58 |
| CD164 | 0.313614 | 2.4E-100 |
| VAMP8 | 0.312259 | 1.2E-126 |
| SAT1 | 0.312235 | 1.36E-67 |
| LRPAP1 | 0.31145 | 7.5E-117 |
| ARF1 | 0.31137 | 7.9E-144 |
| AP1M2 | 0.310599 | 1.08E-68 |
| COX7C | 0.310545 | 2.7E-178 |
| PNCK | 0.309432 | 6.92E-27 |
| ARL1 | 0.308761 | 4.6E-99 |
| EGR1 | 0.308663 | 5.53E-49 |
| COX7A2L | 0.307625 | 1E-129 |
| FXYD1 | 0.307047 | 1.98E-35 |
| PNP | 0.306721 | 1.21E-73 |
| RHOA | 0.30584 | 4.64E-79 |
| WBP2 | 0.30549 | 8.89E-77 |
| PTTG1IP | 0.304747 | 7.91E-90 |
| ORAI3 | 0.304638 | 1.11E-69 |
| ELF3 | 0.304188 | 3.06E-56 |
| YWHAH | 0.30207 | 2.71E-38 |
| TMEM258 | 0.301483 | 4E-123 |
| C12orf75 | 0.300261 | 9.76E-80 |
| C1orf210 | 0.300113 | 3.74E-78 |
| OCEL1 | 0.299803 | 4.94E-77 |
| ASPH | 0.299051 | 3.82E-70 |
| NDUFC1 | 0.299001 | 4.3E-133 |
| PHPT1 | 0.298681 | 9.77E-98 |
| HOMER3 | 0.298222 | 1.97E-73 |
| RETREG2 | 0.298219 | 4.1E-74 |
| MXI1 | 0.297733 | 7.63E-71 |
| CRELD1 | 0.297276 | 1.67E-77 |
| CTDSPL | 0.296224 | 1.41E-77 |

| | | |
|---|---|---|
| FGFR4 | 0.295525 | 6.65E-75 |
| PRDX4 | 0.295437 | 2.46E-80 |
| MGLL | 0.29517 | 6.32E-58 |
| FNBP1L | 0.295128 | 5.78E-70 |
| ARHGAP2 | 0.29492 | 3.92E-60 |
| ERP29 | 0.294423 | 2.7E-111 |
| EFNA1 | 0.293394 | 5.69E-62 |
| STARD10 | 0.292736 | 4.83E-81 |
| AC087482. | 0.292489 | 3.21E-44 |
| CFAP36 | 0.292366 | 1.78E-74 |
| BTG2 | 0.291553 | 1.81E-36 |
| C12orf76 | 0.291458 | 1.23E-70 |
| GIPC2 | 0.290936 | 6.37E-55 |
| GCA | 0.289039 | 1.32E-67 |
| NDUFA1 | 0.288969 | 1.7E-119 |
| FAM131C | 0.28799 | 3.22E-62 |
| ECI1 | 0.28727 | 5.08E-92 |
| ECHS1 | 0.286999 | 4.2E-107 |
| IER5L | 0.286598 | 1.28E-41 |
| BMP2 | 0.285634 | 8.15E-48 |
| JUP | 0.285352 | 3.47E-74 |
| MALAT1 | 0.285173 | 8.6E-78 |
| PFKL | 0.284601 | 6.49E-72 |
| HGD | 0.284051 | 9.82E-70 |
| CREG1 | 0.283762 | 1.19E-66 |
| SYPL1 | 0.283397 | 3.58E-83 |
| RPS7 | 0.283349 | 1.2E-187 |
| IER3IP1 | 0.283093 | 5.39E-88 |
| ARHGDIG | 0.282267 | 5.74E-43 |
| PALM | 0.282259 | 4.4E-74 |
| UQCRH | 0.282206 | 8.19E-95 |
| TRAPPC6 | 0.282012 | 1.35E-75 |
| PLP2 | 0.281976 | 3.71E-37 |
| COMT | 0.281964 | 8.67E-74 |
| GPRC5C | 0.281735 | 2.72E-67 |
| BMP4 | 0.280459 | 5.5E-54 |
| DPPA4 | 0.280411 | 9.55E-49 |
| PPDPF | 0.279047 | 4.49E-92 |
| S100A16 | 0.276966 | 7.48E-72 |
| COX5B | 0.276415 | 1E-148 |
| RPAIN | 0.276334 | 1.99E-78 |
| YPEL2 | 0.276203 | 1.38E-58 |
| CRYL1 | 0.275837 | 3.76E-93 |
| SNX7 | 0.275734 | 8.46E-77 |
| ATP5MC2 | 0.275643 | 7.49E-97 |
| AC090204. | 0.275611 | 1.42E-52 |
| MIR4458H | 0.275308 | 1.78E-56 |
| ARHGAP1 | 0.275166 | 6.2E-60 |
| LINC0095 | 0.27467 | 2.48E-64 |
| MNS1 | 0.274207 | 8.91E-33 |
| VDAC1 | 0.274034 | 3.2E-119 |
| COX20 | 0.273993 | 1.27E-57 |
| LAMB1 | 0.273687 | 4.36E-68 |
| EBP | 0.27366 | 2.46E-64 |
| UGT8 | 0.273205 | 4.84E-63 |
| TNFSF10 | 0.273054 | 7.54E-18 |

| | | |
|---|---|---|
| STXBP2 | 0.272764 | 8.41E-74 |
| COX17 | 0.271508 | 2.04E-70 |
| HOXD1 | 0.271458 | 8E-50 |
| HSPD1 | 0.271197 | 1.01E-93 |
| FUT11 | 0.270479 | 3.41E-65 |
| NDUFB5 | 0.269842 | 2.82E-88 |
| SETD3 | 0.269371 | 2.01E-57 |
| LAMA1 | 0.269156 | 3.18E-58 |
| RDH10 | 0.269073 | 2.47E-53 |
| CDHR5 | 0.268613 | 8.49E-55 |
| MAP7 | 0.268301 | 1.45E-62 |
| NXPH4 | 0.267808 | 1.98E-45 |
| MINDY2 | 0.267777 | 4.73E-65 |
| OTUD1 | 0.26737 | 3.4E-46 |
| ARPC3 | 0.265838 | 2.2E-127 |
| UGCG | 0.265791 | 1.33E-75 |
| PARD6B | 0.265271 | 4.45E-64 |
| MGAT4B | 0.265121 | 6.4E-75 |
| HIGD2A | 0.265049 | 3.53E-75 |
| RBM47 | 0.264757 | 3.59E-64 |
| PHYHIPL | 0.264207 | 4.78E-56 |
| PAWR | 0.264159 | 3.1E-68 |
| TSPAN12 | 0.263959 | 7.85E-61 |
| HILPDA | 0.263698 | 3.91E-36 |
| PTGR1 | 0.263174 | 2.92E-73 |
| VAMP3 | 0.261604 | 3.11E-67 |
| ATRAID | 0.261483 | 3.4E-99 |
| MRPL17 | 0.261457 | 5.1E-65 |
| MET | 0.261347 | 2.08E-65 |
| CISD1 | 0.261339 | 1.02E-72 |
| MMP24OS | 0.260177 | 1.41E-69 |
| TBCA | 0.259953 | 1E-119 |
| GALM | 0.259901 | 4.87E-53 |
| PODXL2 | 0.259726 | 4.24E-55 |
| GRB7 | 0.25971 | 4.43E-62 |
| ZDHHC4 | 0.259247 | 8.08E-70 |
| YBX3 | 0.259017 | 5.81E-78 |
| UBL5 | 0.257924 | 1.8E-139 |
| LETMD1 | 0.257528 | 2.87E-59 |
| MITF | 0.257439 | 4.39E-57 |
| FAM69B | 0.257075 | 7.77E-58 |
| SF3B6 | 0.257064 | 5.43E-94 |
| TECR | 0.257007 | 4.95E-73 |
| JAGN1 | 0.25682 | 1.17E-69 |
| CTXN1 | 0.256803 | 6.14E-61 |
| LLGL2 | 0.254757 | 1.78E-60 |
| OCIAD2 | 0.254347 | 5.72E-59 |
| EPB41L4A | 0.25431 | 1.69E-34 |
| METTL9 | 0.253824 | 6.87E-96 |
| METTL26 | 0.253653 | 2.24E-61 |
| HIP1R | 0.253355 | 7.74E-59 |
| DSG2 | 0.253306 | 6.06E-58 |
| SESTD1 | 0.2531 | 5.85E-41 |
| ZNF770 | 0.25269 | 1.39E-49 |
| MRPL32 | 0.2524 | 1.72E-57 |
| ERICH1 | 0.252346 | 3.92E-54 |

| | | |
|---|---|---|
| ANAPC16 | 0.251886 | 1.23E-92 |
| M6PR | 0.251383 | 1.8E-64 |
| CYB5R3 | 0.251378 | 9.14E-66 |
| PFKFB4 | 0.250982 | 3.85E-59 |
| TSC22D3 | 0.250979 | 2.25E-16 |
| AKR1C3 | 0.250784 | 7.15E-34 |
| EIF1B | 0.250608 | 2.9E-76 |
| KIF12 | 0.250244 | 1.88E-90 |

| **Cluster 9** | | |
|---|---|---|
| Gene | LogFC | pVal |
| MAL | 2.20683 | 0 |
| SLC12A1 | 2.02602 | 1E-297 |
| DEFB1 | 1.977465 | 7.88E-78 |
| IRX1 | 1.58237 | 0 |
| WFDC2 | 1.50691 | 2.4E-143 |
| LINC02381 | 1.403269 | 5E-308 |
| IGFBP7 | 1.387547 | 0 |
| TACSTD2 | 1.340223 | 7.6E-181 |
| S100A14 | 1.314657 | 1.6E-261 |
| FXYD2 | 1.228295 | 1.1E-192 |
| HOXD8 | 1.223111 | 0 |
| TUBB2B | 1.223023 | 0 |
| CLDN10 | 1.19092 | 6.7E-122 |
| ERBB4 | 1.140732 | 4.4E-272 |
| UCHL1 | 1.122379 | 8.8E-276 |
| DUSP9 | 1.061077 | 3.9E-157 |
| VAV3 | 1.045494 | 1.9E-252 |
| CKB | 1.029845 | 0 |
| CD24 | 1.024404 | 0 |
| CLDN6 | 1.017603 | 1.8E-276 |
| NR0B1 | 1.007941 | 1E-150 |
| IRX2 | 0.976158 | 3.3E-166 |
| MUC1 | 0.928526 | 1.8E-177 |
| CLDN4 | 0.917975 | 0 |
| ATP1B1 | 0.910942 | 0 |
| TMPRSS4 | 0.909427 | 2.3E-186 |
| HOTAIRM1 | 0.909271 | 1.5E-208 |
| CYS1 | 0.888848 | 4.6E-287 |
| CEBPD | 0.869861 | 2.7E-240 |
| KIF21A | 0.86903 | 8.6E-126 |
| ATP6V0B | 0.865213 | 4.7E-213 |
| SAT1 | 0.857972 | 7.4E-156 |
| CLDN3 | 0.856805 | 1E-298 |
| TFAP2B | 0.844867 | 3.7E-191 |
| KCNJ1 | 0.810924 | 4.95E-74 |
| CITED2 | 0.803095 | 2.3E-270 |
| DNER | 0.799477 | 3.8E-171 |
| MECOM | 0.796827 | 8.9E-122 |
| MGLL | 0.782742 | 7.6E-200 |
| AP1M2 | 0.77919 | 2.8E-225 |
| EPCAM | 0.770114 | 0 |
| POU3F3 | 0.764636 | 6.9E-203 |
| PERP | 0.756822 | 6.9E-212 |
| COMT | 0.739325 | 1.3E-221 |
| LINC01116 | 0.739116 | 1.1E-171 |
| HOXA9 | 0.729257 | 5.9E-184 |
| KLHL13 | 0.712139 | 6.4E-154 |
| COL18A1 | 0.712087 | 2.5E-224 |
| APLP2 | 0.706879 | 2.4E-246 |
| GPRC5C | 0.703456 | 2.2E-171 |
| CLDN7 | 0.702573 | 3.1E-246 |
| CPM | 0.698645 | 7.9E-123 |
| MTURN | 0.69839 | 9.5E-177 |
| PPDPF | 0.695603 | 7.2E-275 |

| | | |
|---|---|---|
| MGST3 | 0.693745 | 1.4E-253 |
| CYSTM1 | 0.690981 | 9.7E-250 |
| GAL3ST1 | 0.687731 | 3.5E-193 |
| RAB25 | 0.684667 | 1.1E-232 |
| ATP1A1 | 0.684152 | 1.2E-76 |
| UNCX | 0.677841 | 4E-175 |
| CLCN5 | 0.672145 | 3E-147 |
| ANXA11 | 0.67187 | 9.7E-173 |
| RDH10 | 0.66756 | 5.8E-133 |
| CAPG | 0.666757 | 7.9E-193 |
| AKIRIN1 | 0.666084 | 2.1E-206 |
| CAMK2N1 | 0.665155 | 7E-271 |
| PLCB1 | 0.647029 | 1.9E-131 |
| CA12 | 0.640385 | 5.31E-83 |
| IDH2 | 0.636714 | 3.29E-92 |
| PRR15L | 0.632227 | 1.1E-162 |
| HES1 | 0.628582 | 2E-121 |
| S100A16 | 0.6266 | 1.4E-179 |
| HOXB-AS3 | 0.625473 | 1.9E-119 |
| PTGR1 | 0.62394 | 6.9E-138 |
| NDUFA5 | 0.621294 | 1.6E-235 |
| KRT8 | 0.620232 | 1.1E-205 |
| CRACR2B | 0.616264 | 2.5E-131 |
| NUDT4 | 0.61292 | 7.82E-97 |
| COA3 | 0.612919 | 1.7E-188 |
| ARHGAP2 | 0.611095 | 4.6E-148 |
| NEAT1 | 0.610115 | 6E-101 |
| SH3YL1 | 0.607226 | 1.7E-186 |
| STARD10 | 0.607157 | 2.8E-166 |
| KRT19 | 0.60176 | 5.8E-125 |
| KCNJ16 | 0.592468 | 1.2E-132 |
| ACPP | 0.589576 | 1.92E-86 |
| GSTM3 | 0.58863 | 1.6E-154 |
| SLC25A4 | 0.587846 | 7.7E-145 |
| RTN4 | 0.58689 | 8.6E-261 |
| S100A13 | 0.58385 | 6.5E-160 |
| GNG11 | 0.5838 | 8E-166 |
| MT-CO1 | 0.581075 | 1.3E-249 |
| HOXB2 | 0.580406 | 2E-139 |
| SPINT1 | 0.580015 | 1.8E-192 |
| SCIN | 0.57988 | 4E-134 |
| UGT8 | 0.57811 | 1.5E-139 |
| RDH11 | 0.574236 | 1.8E-177 |
| PCBD1 | 0.570637 | 2.7E-254 |
| HOXB6 | 0.570388 | 2.4E-131 |
| CLDN16 | 0.567234 | 3.16E-60 |
| METTL9 | 0.558962 | 3.4E-202 |
| AC004540. | 0.558529 | 8.1E-120 |
| ELF3 | 0.552897 | 7.1E-113 |
| ARHGAP2 | 0.552654 | 1.2E-135 |
| PRSS8 | 0.551428 | 1.9E-142 |
| PAX8 | 0.550755 | 1.7E-153 |
| LLGL2 | 0.550383 | 3.4E-142 |
| AGPAT2 | 0.549238 | 2.7E-196 |
| C4orf48 | 0.548643 | 1.3E-149 |
| CLIC1 | 0.547665 | 3.8E-269 |

| | | |
|---|---|---|
| ITM2C | 0.544275 | 1.1E-209 |
| FKBP4 | 0.544139 | 1.4E-129 |
| CD9 | 0.54099 | 3.2E-165 |
| ATP6V0E1 | 0.537792 | 9.9E-173 |
| MT-CO2 | 0.536182 | 1.6E-281 |
| BMP3 | 0.535044 | 2.49E-88 |
| PTH1R | 0.533173 | 2.3E-72 |
| CPS1 | 0.529986 | 8.72E-56 |
| GIPC1 | 0.522883 | 1.3E-150 |
| SNCG | 0.521868 | 1.81E-49 |
| GLRX | 0.521163 | 7.3E-119 |
| PLEKHA5 | 0.521103 | 7.8E-124 |
| EMX2 | 0.519778 | 2.7E-158 |
| UQCR10 | 0.518912 | 6E-213 |
| TSPAN33 | 0.518653 | 3.2E-128 |
| SPINT2 | 0.517558 | 0 |
| RGL3 | 0.512227 | 6.1E-134 |
| AMIGO2 | 0.51222 | 2.03E-92 |
| HOOK1 | 0.510346 | 3.1E-143 |
| ADGRG1 | 0.508126 | 5.9E-119 |
| COBLL1 | 0.505115 | 2E-123 |
| VAMP8 | 0.504332 | 2.2E-225 |
| DSTN | 0.499272 | 4.2E-198 |
| RBM47 | 0.498023 | 3.1E-116 |
| TSTD1 | 0.493776 | 3.84E-84 |
| CTXN1 | 0.490335 | 5.6E-123 |
| PAX2 | 0.485493 | 1.3E-114 |
| ESRRG | 0.48421 | 1.7E-104 |
| KRT18 | 0.481465 | 1.1E-204 |
| SEPHS2 | 0.481453 | 1.5E-156 |
| DCDC2 | 0.480465 | 4.1E-122 |
| CRYL1 | 0.480421 | 1.1E-133 |
| HOXA-AS2 | 0.479399 | 1.7E-105 |
| PTPN13 | 0.476498 | 3.8E-106 |
| BLNK | 0.476181 | 9.3E-122 |
| S100A11 | 0.473227 | 4.9E-137 |
| ATP6V0A4 | 0.471998 | 2.89E-86 |
| MINDY2 | 0.471307 | 2.4E-123 |
| S100A6 | 0.470478 | 1.08E-33 |
| CYCS | 0.46401 | 2.9E-111 |
| DSP | 0.463164 | 6.94E-96 |
| C5orf38 | 0.462606 | 9.19E-94 |
| RABAC1 | 0.461492 | 2.5E-131 |
| MPC1 | 0.461339 | 2.3E-104 |
| RHOB | 0.460665 | 6.8E-157 |
| MITF | 0.460414 | 4.3E-128 |
| TMEM37 | 0.45975 | 8.09E-80 |
| DMKN | 0.459472 | 6.6E-115 |
| MYEF2 | 0.458684 | 1.5E-112 |
| NAT14 | 0.458449 | 1.4E-138 |
| TFCP2L1 | 0.458179 | 1.86E-96 |
| NDUFB5 | 0.457805 | 2.4E-147 |
| MIR503HG | 0.456646 | 9.6E-111 |
| DDR1 | 0.456578 | 2.9E-124 |
| HNF1B | 0.456228 | 2E-127 |
| ADI1 | 0.455668 | 2.8E-100 |

| | | |
|---|---|---|
| ABRACL | 0.455617 | 1.6E-129 |
| REEP5 | 0.455549 | 2.2E-140 |
| TMEM38B | 0.454535 | 6E-118 |
| COL6A1 | 0.454182 | 7.51E-48 |
| NDUFB9 | 0.453016 | 2.2E-113 |
| UBAC2 | 0.449608 | 3.7E-110 |
| CST3 | 0.448779 | 8.4E-155 |
| TMEM213 | 0.447777 | 5.03E-73 |
| SLC4A4 | 0.447291 | 6.29E-95 |
| MT-ND4 | 0.446213 | 3E-218 |
| MT-CO3 | 0.445003 | 6E-213 |
| LAPTM4B | 0.444778 | 1.7E-147 |
| CYB5A | 0.444311 | 4.1E-189 |
| ARPC3 | 0.441723 | 8.4E-182 |
| ACADVL | 0.440322 | 2.2E-100 |
| TXNIP | 0.439568 | 1.1E-109 |
| CD151 | 0.437715 | 1.4E-192 |
| COX5A | 0.437395 | 1.3E-110 |
| CA4 | 0.436748 | 4.96E-76 |
| DYNC2LI1 | 0.43667 | 9.5E-113 |
| APMAP | 0.435432 | 3.8E-106 |
| NDFIP1 | 0.433857 | 1.4E-129 |
| NCOA7 | 0.432822 | 3.98E-77 |
| MT-CYB | 0.429433 | 6E-154 |
| CXXC5 | 0.428311 | 4.9E-184 |
| TM7SF2 | 0.428266 | 1.4E-129 |
| SNCA | 0.428069 | 9.77E-92 |
| SDC4 | 0.42795 | 1.1E-114 |
| CLDN19 | 0.426631 | 3.1E-102 |
| PPP1R1A | 0.426088 | 4.3E-129 |
| CTDSPL | 0.425954 | 1.7E-111 |
| LDHB | 0.423366 | 1.2E-275 |
| GOLM1 | 0.422485 | 1.4E-103 |
| SUCLG1 | 0.422118 | 1.4E-99 |
| LMBRD1 | 0.421417 | 3.24E-97 |
| CEBPA | 0.420988 | 1.29E-92 |
| RASSF7 | 0.420807 | 6.3E-113 |
| SNRPN | 0.420161 | 9.9E-125 |
| TRAPPC6/ | 0.416197 | 3.2E-115 |
| IER5L | 0.415418 | 1.66E-69 |
| JUN | 0.415374 | 4.5E-120 |
| CPNE8 | 0.414717 | 1.37E-61 |
| EZR | 0.413402 | 4.2E-139 |
| MAGEF1 | 0.40952 | 5E-107 |
| TRAF4 | 0.409191 | 8.8E-112 |
| HLA-DMB | 0.409137 | 5.04E-87 |
| ITM2B | 0.408577 | 1.8E-255 |
| MRPS36 | 0.408141 | 1.7E-136 |
| COX20 | 0.40678 | 7.7E-105 |
| CDH1 | 0.403379 | 4.23E-97 |
| ZNF44 | 0.40192 | 2.44E-55 |
| TMBIM6 | 0.401168 | 1.3E-243 |
| CNDP2 | 0.400457 | 5.6E-99 |
| PON2 | 0.400236 | 8.6E-125 |
| PNP | 0.398624 | 1.3E-84 |
| IER3 | 0.398204 | 8.16E-54 |

| | | |
|---|---|---|
| ATP6AP2 | 0.397549 | 7.8E-113 |
| TUSC3 | 0.397309 | 2.1E-115 |
| NR2F6 | 0.396635 | 4.2E-168 |
| HSP90AA1 | 0.396575 | 3.8E-135 |
| PBDC1 | 0.39586 | 1.61E-90 |
| OCLN | 0.39537 | 9.66E-86 |
| PLP2 | 0.393481 | 4.54E-80 |
| PRSS35 | 0.391894 | 1.09E-89 |
| EPN1 | 0.391673 | 1.1E-105 |
| CAPZA2 | 0.391487 | 6.3E-111 |
| MLF1 | 0.3901 | 9.74E-89 |
| ATP5F1B | 0.389762 | 1.3E-107 |
| EGFL7 | 0.388793 | 1.63E-70 |
| GCA | 0.388462 | 5.41E-93 |
| HOXB7 | 0.388215 | 7.11E-98 |
| PNKD | 0.388062 | 4.9E-111 |
| ARID3A | 0.38683 | 6.19E-98 |
| S100A4 | 0.386386 | 8.3E-39 |
| MOCS2 | 0.384691 | 3.73E-91 |
| MRPL33 | 0.384108 | 2.2E-125 |
| HIPK2 | 0.383548 | 5.46E-88 |
| BSG | 0.383329 | 2.8E-202 |
| UPP1 | 0.382746 | 1.14E-86 |
| ABHD17C | 0.38268 | 4.1E-82 |
| PHGDH | 0.381489 | 2.2E-110 |
| KCTD1 | 0.381349 | 1.62E-85 |
| TCIM | 0.378571 | 1.8E-62 |
| PARD6B | 0.377365 | 9.63E-85 |
| ERICH1 | 0.377105 | 2.35E-72 |
| CAPS | 0.376938 | 4.24E-81 |
| NPC2 | 0.375037 | 4.4E-161 |
| MYL12B | 0.374959 | 4E-137 |
| PHACTR1 | 0.374474 | 8.57E-71 |
| HOXD1 | 0.374338 | 2E-83 |
| SINHCAF | 0.37429 | 1.5E-84 |
| RAB3IP | 0.373672 | 2.05E-89 |
| METRNL | 0.371817 | 3.7E-96 |
| CXADR | 0.371208 | 3.43E-90 |
| IMPA2 | 0.370891 | 6.1E-103 |
| PRDX3 | 0.369353 | 7E-124 |
| DBI | 0.368908 | 1.8E-122 |
| CYBA | 0.368535 | 4.2E-128 |
| GDF11 | 0.366409 | 3.89E-75 |
| HIGD1A | 0.365648 | 6.69E-89 |
| MIEN1 | 0.365595 | 1.1E-100 |
| ALKAL2 | 0.364352 | 9.98E-64 |
| PIM3 | 0.364321 | 3.41E-76 |
| AP002884. | 0.364227 | 5.24E-70 |
| CCDC160 | 0.362637 | 8.87E-87 |
| HOXD9 | 0.36209 | 2.23E-76 |
| DDAH2 | 0.36087 | 3.2E-125 |
| ABHD11 | 0.360774 | 6.16E-87 |
| DPP4 | 0.360547 | 2.89E-88 |
| LINC0095{ | 0.360023 | 1.54E-82 |
| ZNF467 | 0.359985 | 6.28E-94 |
| NFKBIA | 0.359601 | 2.98E-63 |

| | | |
|---|---|---|
| TSC22D2 | 0.358258 | 3.83E-60 |
| MGAT4B | 0.358052 | 2.3E-101 |
| FXYD6 | 0.35682 | 2E-86 |
| ATP1B3 | 0.356299 | 6.06E-74 |
| IRF2BPL | 0.35623 | 6.08E-74 |
| GOT1 | 0.355662 | 1.23E-68 |
| SH3GLB2 | 0.355564 | 1.36E-86 |
| PAWR | 0.355239 | 8.74E-94 |
| IER2 | 0.354349 | 7.8E-101 |
| BCAM | 0.354072 | 3.1E-149 |
| ID4 | 0.353823 | 7.61E-56 |
| HOXA7 | 0.353393 | 5.25E-79 |
| PCSK1N | 0.35288 | 2.43E-52 |
| EEF1E1 | 0.352616 | 2.43E-95 |
| YWHAZ | 0.35227 | 8.64E-95 |
| CTSL | 0.352079 | 1.37E-40 |
| TMEM123 | 0.350864 | 1.6E-88 |
| IVNS1ABP | 0.347495 | 4.92E-72 |
| ATP6V1G1 | 0.346994 | 3.7E-144 |
| FXYD3 | 0.346739 | 6.63E-52 |
| NDUFB2 | 0.345931 | 5.1E-119 |
| MARVELD | 0.345776 | 1.07E-91 |
| SKAP2 | 0.344612 | 1.47E-72 |
| KRAS | 0.344158 | 9.71E-88 |
| CAPN1 | 0.344075 | 9.25E-89 |
| HOXC4 | 0.344071 | 3.98E-77 |
| CD47 | 0.343827 | 1.33E-62 |
| SPINT1-AS | 0.343821 | 2.44E-78 |
| MARCKSL | 0.343732 | 2.5E-198 |
| VPS25 | 0.34297 | 3.15E-95 |
| MT-ND5 | 0.342808 | 3.4E-112 |
| MT-ND4L | 0.342566 | 4.5E-99 |
| ATP5MC1 | 0.342444 | 3.25E-71 |
| NDUFA3 | 0.342074 | 2.2E-102 |
| RPAIN | 0.341799 | 1.7E-112 |
| COX5B | 0.341609 | 7.3E-106 |
| ANK3 | 0.341266 | 7.05E-81 |
| RTN3 | 0.340476 | 1.96E-89 |
| LSR | 0.340383 | 1.63E-86 |
| NDUFA8 | 0.339961 | 4.15E-82 |
| SLC29A1 | 0.339335 | 1.84E-82 |
| GRB14 | 0.338796 | 9.75E-74 |
| PTS | 0.33809 | 5.46E-86 |
| ZMAT3 | 0.33789 | 5.22E-59 |
| EPB41L5 | 0.337466 | 3.82E-86 |
| PRKAB1 | 0.337446 | 1.08E-81 |
| ATP6V1A | 0.336631 | 2.32E-78 |
| SMTNL2 | 0.336623 | 4.43E-82 |
| SLC3A2 | 0.336548 | 9.1E-66 |
| AIF1L | 0.336133 | 7.25E-80 |
| MT-ATP6 | 0.336039 | 7.2E-146 |
| SSFA2 | 0.335602 | 5.05E-71 |
| LRPAP1 | 0.335148 | 2.1E-115 |
| PLD3 | 0.334999 | 7.5E-99 |
| DAZAP2 | 0.334473 | 7.12E-90 |
| CD46 | 0.334466 | 5.16E-92 |

| | | |
|---|---|---|
| TMEM167/ | 0.333839 | 2.11E-88 |
| TOM1L1 | 0.333747 | 1.17E-73 |
| MYL6 | 0.333333 | 1.5E-140 |
| SCNN1A | 0.333065 | 9.94E-80 |
| ATP6V0E2 | 0.332322 | 1.32E-78 |
| ADGRF1 | 0.332264 | 1.11E-69 |
| MTX2 | 0.331596 | 9E-104 |
| TMEM125 | 0.330971 | 2.42E-78 |
| YWHAB | 0.330541 | 5.7E-106 |
| COX7B | 0.330146 | 4.36E-87 |
| KIF12 | 0.330019 | 6.1E-100 |
| NAPA | 0.329976 | 2.16E-78 |
| MAP9 | 0.329723 | 2.48E-79 |
| JUP | 0.329511 | 1.11E-86 |
| ARL1 | 0.328968 | 9.8E-104 |
| BCL7A | 0.328744 | 5.11E-79 |
| TMEM159 | 0.328647 | 2.46E-67 |
| SCP2 | 0.328265 | 7.55E-84 |
| PODXL2 | 0.328184 | 8.89E-70 |
| CDH3 | 0.326547 | 1.3E-85 |
| NDUFS6 | 0.32592 | 7.99E-96 |
| HOXB9 | 0.325625 | 2.59E-65 |
| SEC23B | 0.325358 | 1.84E-71 |
| ENSA | 0.325057 | 4.12E-90 |
| TPST1 | 0.323782 | 1.21E-79 |
| SEC11C | 0.323129 | 3.38E-77 |
| MTSS1 | 0.321749 | 2.37E-75 |
| STMP1 | 0.321738 | 3.1E-113 |
| TMEM50A | 0.321474 | 3.78E-87 |
| SLC25A39 | 0.320695 | 9.54E-77 |
| TEX264 | 0.319932 | 5.19E-75 |
| MRPS6 | 0.319276 | 3.37E-93 |
| UBL5 | 0.318833 | 8.6E-127 |
| RBBP8 | 0.318211 | 4.25E-60 |
| ARHGEF2 | 0.318096 | 7.5E-80 |
| LMAN2 | 0.317989 | 2.2E-103 |
| CMTM6 | 0.317914 | 4.66E-82 |
| HEBP2 | 0.317675 | 7.29E-82 |
| LRRK2 | 0.317483 | 2.7E-57 |
| NDUFB3 | 0.317207 | 1.25E-81 |
| LYPLA1 | 0.316676 | 1.98E-73 |
| FMC1 | 0.316033 | 4.72E-66 |
| MPC2 | 0.315926 | 4.7E-142 |
| BICC1 | 0.314164 | 2.82E-50 |
| LAMA1 | 0.314111 | 1.07E-68 |
| CACNB4 | 0.313372 | 3.24E-64 |
| DEPTOR | 0.312649 | 6.05E-67 |
| MMP24OS | 0.312628 | 1.59E-72 |
| FKBP2 | 0.312372 | 1.19E-91 |
| MCRIP2 | 0.312278 | 4.3E-63 |
| COX6A1 | 0.312019 | 9E-134 |
| TMEM205 | 0.311644 | 4.15E-73 |
| ITGB6 | 0.311586 | 1.43E-64 |
| HOXD11 | 0.311421 | 3.05E-65 |
| HINT2 | 0.311344 | 3.1E-75 |
| PATZ1 | 0.310981 | 1.81E-72 |

| | | |
|---|---|---|
| TLCD1 | 0.310764 | 2.75E-75 |
| HOXB8 | 0.310542 | 2.12E-70 |
| COX6B1 | 0.30952 | 8.5E-99 |
| SRI | 0.309021 | 1.86E-80 |
| ORAI2 | 0.308425 | 2.9E-68 |
| GBE1 | 0.307959 | 1.04E-66 |
| PLEKHB2 | 0.307333 | 9.35E-67 |
| P4HTM | 0.307043 | 6.01E-82 |
| SCTR | 0.305345 | 3.2E-62 |
| DPY30 | 0.304937 | 1.24E-96 |
| FREM2 | 0.30453 | 1.14E-55 |
| MST1 | 0.303909 | 5.23E-63 |
| HPRT1 | 0.303564 | 4.65E-72 |
| COQ9 | 0.303532 | 2.23E-80 |
| LNX1 | 0.301229 | 1.5E-66 |
| LAP3 | 0.30114 | 9.38E-62 |
| MAD2L2 | 0.300491 | 6.27E-67 |
| ATP5F1C | 0.300176 | 1.85E-83 |
| MBD2 | 0.300057 | 3.01E-69 |
| CDK2AP2 | 0.29972 | 7.58E-76 |
| POLR2L | 0.299666 | 3.05E-89 |
| CISD1 | 0.299665 | 1.23E-63 |
| SYAP1 | 0.299545 | 3.56E-64 |
| TAX1BP1 | 0.29914 | 6.94E-81 |
| CASR | 0.29885 | 1.46E-63 |
| COX17 | 0.298667 | 3.14E-75 |
| TNFRSF12 | 0.298459 | 8.47E-37 |
| PPP1CA | 0.298454 | 9.28E-76 |
| HDAC1 | 0.298412 | 1.89E-70 |
| UGT2B7 | 0.298196 | 5.92E-54 |
| MAL2 | 0.297637 | 7.33E-79 |
| SDHD | 0.29756 | 1.31E-77 |
| TMEM59 | 0.297441 | 3.1E-80 |
| MYO6 | 0.29736 | 1.12E-84 |
| MRPL34 | 0.297143 | 1.57E-69 |
| RRAGD | 0.296997 | 2.19E-59 |
| EVA1B | 0.295904 | 3.98E-78 |
| AK3 | 0.294747 | 9.38E-64 |
| C19orf70 | 0.294513 | 6.86E-73 |
| MYL12A | 0.294488 | 1.23E-93 |
| ATP6AP1 | 0.294395 | 1.48E-79 |
| IQGAP1 | 0.294019 | 1.99E-68 |
| NDUFA4 | 0.29372 | 3.4E-95 |
| KIAA1217 | 0.293694 | 8.54E-58 |
| TRAPPC1 | 0.292758 | 1.31E-87 |
| TPM1 | 0.292218 | 2.9E-102 |
| EMC10 | 0.292136 | 3.81E-48 |
| SPSB3 | 0.291911 | 1.75E-72 |
| CCNDBP1 | 0.291727 | 1.29E-73 |
| NBDY | 0.290973 | 1.56E-85 |
| HOXB3 | 0.290948 | 1.21E-67 |
| PAPSS2 | 0.290016 | 5.11E-58 |
| TMEM179B | 0.288813 | 8.14E-70 |
| HOXA3 | 0.28877 | 1.81E-53 |
| NDUFV2 | 0.28832 | 5.74E-78 |
| MAPK10 | 0.287968 | 2.37E-68 |

| | | |
|---|---|---|
| EMC6 | 0.287674 | 1.79E-73 |
| CD2AP | 0.287632 | 1.25E-71 |
| MT-ND3 | 0.287192 | 8.2E-185 |
| ARSD | 0.287113 | 2.21E-57 |
| PRCP | 0.28687 | 6.18E-47 |
| MINOS1 | 0.286863 | 3.29E-64 |
| SYPL1 | 0.286284 | 6.07E-70 |
| NECTIN2 | 0.286018 | 9.89E-68 |
| GSKIP | 0.285692 | 2.9E-70 |
| UQCRFS1 | 0.285257 | 4.83E-82 |
| UQCR11 | 0.285071 | 1.72E-80 |
| CHCHD5 | 0.284659 | 4.09E-65 |
| PDHB | 0.284494 | 6.23E-73 |
| MFHAS1 | 0.284225 | 4.98E-63 |
| TBCB | 0.284225 | 2.9E-80 |
| DNAJA2 | 0.283825 | 6.3E-66 |
| TADA3 | 0.283376 | 9.43E-70 |
| GRN | 0.283285 | 6.31E-74 |
| PRRG2 | 0.283078 | 3.8E-62 |
| TPD52 | 0.282938 | 2.26E-56 |
| TUBA4A | 0.282629 | 9.79E-57 |
| HIBADH | 0.281334 | 3.36E-57 |
| PKP4 | 0.281004 | 8.09E-54 |
| VWA1 | 0.280827 | 2.14E-63 |
| ACSM3 | 0.280618 | 2.89E-64 |
| JTB | 0.280485 | 1.74E-80 |
| AHCYL1 | 0.280408 | 2.69E-64 |
| PPCS | 0.280066 | 3.46E-69 |
| CIAPIN1 | 0.279667 | 3.23E-68 |
| ARRDC3 | 0.279639 | 4.41E-61 |
| REXO2 | 0.279459 | 1.16E-51 |
| ATRAID | 0.279416 | 2.62E-81 |
| TSG101 | 0.278999 | 1.18E-72 |
| ZDHHC12 | 0.278929 | 4.22E-67 |
| AKAP9 | 0.278669 | 2.04E-55 |
| CDH16 | 0.278439 | 5.31E-60 |
| ACSL4 | 0.278436 | 3.81E-42 |
| GABARAP | 0.278403 | 3.53E-97 |
| AFDN | 0.278173 | 6.44E-62 |
| GHITM | 0.278009 | 2.52E-64 |
| SYNGR2 | 0.277702 | 1.86E-67 |
| PLAGL1 | 0.277353 | 7.22E-51 |
| ABHD12 | 0.276775 | 1.25E-62 |
| PTPRF | 0.276622 | 2.39E-56 |
| GNAI1 | 0.276583 | 1.9E-42 |
| PIGH | 0.276496 | 9.98E-58 |
| NOP10 | 0.275995 | 4.59E-85 |
| MRPL36 | 0.275742 | 1.41E-59 |
| B4GALT1 | 0.275529 | 3.01E-36 |
| TMEM72 | 0.27552 | 2.27E-55 |
| CIB1 | 0.275391 | 9.11E-70 |
| DYNLRB1 | 0.275361 | 1.51E-97 |
| NARS | 0.275325 | 5.52E-61 |
| TRIM8 | 0.275232 | 2.63E-71 |
| FAM96B | 0.275196 | 3.6E-87 |
| IFT57 | 0.274686 | 6.23E-77 |

| | | |
|---|---|---|
| AC079630. | 0.27444 | 3.03E-40 |
| STAP2 | 0.274246 | 5.68E-55 |
| ZFYVE21 | 0.274141 | 1.73E-69 |
| KIAA1522 | 0.273853 | 4.1E-57 |
| PSAP | 0.273236 | 1.42E-72 |
| IRX3 | 0.272194 | 3.8E-32 |
| MTRNR2L | 0.272046 | 6.7E-46 |
| TXNDC12 | 0.271861 | 1.01E-73 |
| CDC34 | 0.271846 | 9.82E-67 |
| ZDHHC14 | 0.271056 | 9.04E-60 |
| RNF13 | 0.270335 | 6.44E-62 |
| MAPKAPK | 0.270189 | 4.45E-63 |
| SFT2D1 | 0.269843 | 2.33E-72 |
| ACYP1 | 0.268536 | 4.23E-58 |
| CDC42SE1 | 0.268166 | 4.16E-48 |
| PPP1CB | 0.267749 | 1.76E-51 |
| SMDT1 | 0.267324 | 2.62E-72 |
| SLC35F2 | 0.267158 | 2.38E-60 |
| HPN | 0.267036 | 2.6E-136 |
| TCTA | 0.266925 | 5.79E-55 |
| RHOA | 0.26692 | 1.88E-84 |
| TMBIM4 | 0.266496 | 6.79E-61 |
| TIMM17B | 0.266466 | 1.51E-80 |
| FIS1 | 0.265377 | 1.22E-72 |
| DARS | 0.265245 | 1.38E-62 |
| NIPSNAP2 | 0.264741 | 1.58E-62 |
| CLDN23 | 0.264649 | 3.83E-48 |
| SLC44A3 | 0.263897 | 4.25E-55 |
| SDHAF3 | 0.263716 | 2.39E-59 |
| B3GNT2 | 0.263364 | 2.63E-54 |
| COA5 | 0.263302 | 2.72E-68 |
| SLC44A2 | 0.26306 | 3.64E-57 |
| FAM84A | 0.262551 | 1.34E-54 |
| NDUFA2 | 0.262382 | 1.02E-78 |
| PTTG1IP | 0.262279 | 1.89E-69 |
| SELENOT | 0.262261 | 9.22E-68 |
| KLF5 | 0.261822 | 8.77E-49 |
| EIF5 | 0.261615 | 2.28E-74 |
| TSPAN12 | 0.261403 | 2.22E-75 |
| TUFM | 0.261308 | 1.16E-69 |
| ROGDI | 0.261261 | 8.2E-53 |
| UQCRC1 | 0.260939 | 1.99E-58 |
| ANP32E | 0.26083 | 1.25E-73 |
| ARPC5 | 0.260774 | 4.27E-77 |
| RHBDD2 | 0.260381 | 4.59E-54 |
| FGF9 | 0.260312 | 1.42E-56 |
| ACTR3 | 0.259951 | 2.62E-57 |
| GLB1 | 0.259598 | 2.41E-61 |
| DPM3 | 0.259527 | 5.6E-60 |
| CDC42 | 0.258816 | 2.65E-75 |
| IER3IP1 | 0.258468 | 3.5E-70 |
| GCC2 | 0.258372 | 2.15E-53 |
| UNC50 | 0.258196 | 5.9E-66 |
| TUBA1C | 0.258126 | 1.45E-72 |
| RPN2 | 0.257819 | 1.11E-59 |
| CGN | 0.257489 | 4.57E-60 |

| | | |
|---|---|---|
| SLC38A1 | 0.257362 | 9.94E-57 |
| TMEM9B | 0.256654 | 6.64E-59 |
| GTF2H5 | 0.256642 | 5.64E-67 |
| CLCNKB | 0.256582 | 1.89E-40 |
| VDAC1 | 0.256417 | 5.47E-63 |
| CCDC151 | 0.256403 | 1.38E-52 |
| HADHB | 0.256079 | 1.61E-56 |
| PNPLA2 | 0.255912 | 1.95E-54 |
| TMEM8A | 0.255659 | 2.8E-53 |
| SERTAD3 | 0.255659 | 4.31E-51 |
| DSC2 | 0.255649 | 2.09E-52 |
| ERP29 | 0.25541 | 3.32E-69 |
| SBDS | 0.255396 | 2.78E-71 |
| COX11 | 0.255382 | 5.25E-53 |
| ECHDC2 | 0.255133 | 3.38E-62 |
| C1orf43 | 0.254664 | 1.77E-83 |
| ITGA2 | 0.254516 | 2.04E-65 |
| ELF1 | 0.254143 | 3.34E-55 |
| NDUFS7 | 0.254138 | 2.33E-60 |
| TSPAN15 | 0.253909 | 4.18E-56 |
| NDUFC2 | 0.253852 | 7.96E-62 |
| GCGR | 0.253841 | 2.91E-57 |
| HOXD3 | 0.253484 | 4.03E-61 |
| NDUFA1 | 0.253451 | 2.09E-75 |
| LYPLAL1 | 0.253188 | 1.9E-51 |
| LGR4 | 0.252883 | 8.09E-55 |
| WNT7B | 0.252572 | 1.88E-46 |
| EPS8L1 | 0.25229 | 1.28E-56 |
| ATP6V0D1 | 0.252151 | 2.56E-59 |
| DAPK1 | 0.251785 | 2.04E-50 |
| ECI1 | 0.251759 | 3.76E-58 |
| NDUFB4 | 0.251697 | 2.72E-78 |
| ATP5MD | 0.251056 | 1.4E-71 |
| SMIM14 | 0.250206 | 1.2E-57 |
| VAMP2 | 0.250107 | 1.56E-62 |
| SLC6A13 | 0.250005 | 2.23E-60 |

| **Cluster 10** | | |
|---|---|---|
| Gene | LogFC | pVal |
| HIST1H4C | 2.816922 | 0 |
| HIST1H1D | 1.960687 | 0 |
| HIST1H1A | 1.934971 | 0 |
| HIST1H1B | 1.820312 | 0 |
| HMGB2 | 1.629472 | 0 |
| PCLAF | 1.619736 | 0 |
| TOP2A | 1.580108 | 0 |
| TUBA1B | 1.529124 | 0 |
| UBE2C | 1.466712 | 0 |
| CENPF | 1.463478 | 1E-300 |
| H2AFX | 1.461447 | 0 |
| NUSAP1 | 1.44605 | 2.1E-292 |
| MKI67 | 1.443801 | 8.3E-278 |
| CDK1 | 1.435278 | 0 |
| TYMS | 1.431279 | 0 |
| H2AFZ | 1.303014 | 0 |
| HIST1H1C | 1.268023 | 5.6E-233 |
| HMGN2 | 1.232288 | 0 |
| BIRC5 | 1.145819 | 5.3E-270 |
| SMC4 | 1.127544 | 1.6E-232 |
| RRM2 | 1.123774 | 1.7E-229 |
| TPX2 | 1.078586 | 1.4E-220 |
| TMSB15A | 1.075963 | 3.5E-285 |
| HIST2H2A | 1.070939 | 8E-213 |
| ZWINT | 1.042038 | 6.4E-252 |
| HIST1H1E | 1.024272 | 6.4E-221 |
| MAD2L1 | 1.004487 | 1.3E-233 |
| MXD3 | 0.987987 | 8.4E-184 |
| UBE2T | 0.985233 | 9.4E-248 |
| TUBB | 0.977692 | 0 |
| STMN1 | 0.970675 | 0 |
| HMGB1 | 0.967534 | 0 |
| CKS2 | 0.957537 | 1.6E-169 |
| PTTG1 | 0.944217 | 5.7E-173 |
| CKS1B | 0.940543 | 2E-233 |
| AURKB | 0.927093 | 2E-202 |
| DEK | 0.909648 | 0 |
| CENPW | 0.905965 | 3.1E-213 |
| ATAD2 | 0.892737 | 2.3E-183 |
| NDC80 | 0.882486 | 3.5E-179 |
| UBE2S | 0.875789 | 2.3E-178 |
| SPC25 | 0.865209 | 4E-190 |
| ASPM | 0.860675 | 2.3E-134 |
| DLGAP5 | 0.835882 | 9.6E-147 |
| H2AFV | 0.831539 | 1.9E-275 |
| CENPK | 0.825063 | 8.6E-197 |
| RAD51AP1 | 0.823627 | 1.4E-184 |
| KIFC1 | 0.81445 | 9.1E-180 |
| CDT1 | 0.811597 | 3.7E-171 |
| ESCO2 | 0.807561 | 2E-164 |
| CDCA4 | 0.804812 | 6.6E-204 |
| FBXO5 | 0.801562 | 2.6E-182 |
| DUT | 0.78898 | 2E-146 |
| NASP | 0.787828 | 9.5E-249 |

| | | |
|---|---|---|
| KPNA2 | 0.784977 | 5.1E-144 |
| CKAP2 | 0.784662 | 1.9E-164 |
| SMC2 | 0.774676 | 6.7E-192 |
| MIS18BP1 | 0.773378 | 2.3E-160 |
| TK1 | 0.763897 | 3.9E-173 |
| CENPM | 0.753765 | 1.4E-165 |
| KIF11 | 0.746958 | 5.7E-145 |
| CCNB2 | 0.746835 | 4.4E-136 |
| MND1 | 0.739931 | 4E-160 |
| KIF20B | 0.736289 | 1.9E-140 |
| CENPE | 0.732319 | 1E-116 |
| PCNA | 0.730314 | 9.6E-149 |
| CDCA3 | 0.724432 | 3.3E-126 |
| RRM1 | 0.722782 | 1.9E-181 |
| CDKN3 | 0.717646 | 3E-134 |
| PTN | 0.713025 | 0 |
| ITGB3BP | 0.712979 | 4.6E-180 |
| DTYMK | 0.691484 | 3.1E-196 |
| CLSPN | 0.689636 | 8.8E-128 |
| EZH2 | 0.685899 | 2.5E-149 |
| KIF2C | 0.682969 | 7.6E-140 |
| RANBP1 | 0.680985 | 2.4E-215 |
| CENPH | 0.678215 | 8.1E-143 |
| ORC6 | 0.676389 | 1.3E-164 |
| TMPO | 0.675401 | 3.4E-154 |
| PSIP1 | 0.674669 | 1.7E-198 |
| ANP32E | 0.665296 | 5.1E-175 |
| CKAP2L | 0.657996 | 1.2E-136 |
| GTSE1 | 0.654244 | 9.7E-136 |
| DNMT1 | 0.654041 | 5E-155 |
| SPC24 | 0.653752 | 6.5E-140 |
| NUDT1 | 0.652281 | 3E-163 |
| CENPU | 0.641184 | 9.3E-137 |
| LMNB1 | 0.638096 | 6.4E-144 |
| KIF23 | 0.63703 | 2.1E-123 |
| CCNB1 | 0.636882 | 6.94E-91 |
| USP1 | 0.635753 | 6.7E-154 |
| HELLS | 0.634518 | 2.1E-143 |
| RAD21 | 0.633259 | 3.3E-143 |
| TACC3 | 0.633064 | 4.2E-128 |
| NCAPG | 0.630185 | 1.1E-123 |
| CENPN | 0.626058 | 3.4E-138 |
| GINS2 | 0.621833 | 2.3E-128 |
| CENPA | 0.619127 | 6.15E-91 |
| SGO2 | 0.6175 | 4.8E-100 |
| NUCKS1 | 0.615954 | 3.6E-200 |
| DNAJC9 | 0.615465 | 4.9E-146 |
| KIF15 | 0.614725 | 7.8E-116 |
| SNRNP25 | 0.612565 | 8.2E-156 |
| SGO1 | 0.612526 | 1.1E-126 |
| GMNN | 0.612375 | 7.6E-153 |
| KIF22 | 0.60856 | 8.7E-133 |
| NUF2 | 0.599224 | 2.1E-120 |
| KNL1 | 0.597271 | 3.4E-107 |
| HIST1H3B | 0.597 | 1.9E-121 |
| DIAPH3 | 0.585067 | 5.4E-121 |

| | | |
|---|---|---|
| HMGB3 | 0.584562 | 1.6E-127 |
| TMEM106( | 0.58381 | 1.1E-136 |
| PIMREG | 0.579418 | 2.2E-111 |
| CBX5 | 0.574463 | 2.6E-178 |
| CCDC34 | 0.573264 | 2.3E-133 |
| HIST1H2A | 0.564758 | 3.61E-96 |
| HNRNPA2 | 0.562211 | 2.5E-214 |
| FBLN1 | 0.558026 | 7.8E-180 |
| SAE1 | 0.55425 | 3.7E-127 |
| CCNA2 | 0.553511 | 5.6E-112 |
| MYBL2 | 0.55312 | 2.4E-111 |
| NNAT | 0.551323 | 1.5E-191 |
| ALYREF | 0.540147 | 1.6E-131 |
| ASF1B | 0.538558 | 2.1E-115 |
| EMP2 | 0.538291 | 6.3E-112 |
| SKA2 | 0.5371 | 2.5E-117 |
| SMC3 | 0.53446 | 2.7E-141 |
| ANP32B | 0.533711 | 8.5E-192 |
| CMC2 | 0.532755 | 1.4E-134 |
| BUB1B | 0.530768 | 2.1E-102 |
| LSM4 | 0.528909 | 5.5E-142 |
| TUBB6 | 0.528248 | 1.34E-93 |
| CDCA5 | 0.527717 | 6.9E-117 |
| MCM7 | 0.522864 | 2.4E-108 |
| H1FX | 0.522474 | 7E-138 |
| ANLN | 0.508241 | 6.4E-102 |
| HNRNPR | 0.503481 | 3.5E-190 |
| CDCA8 | 0.501122 | 2.97E-86 |
| NT5DC2 | 0.497469 | 7.7E-140 |
| CALM2 | 0.496793 | 1.6E-176 |
| TUBB4B | 0.4951 | 1.8E-127 |
| CDC20 | 0.495012 | 1.44E-77 |
| RFC4 | 0.491464 | 5E-113 |
| FOXM1 | 0.488134 | 2.4E-105 |
| RHEB | 0.483254 | 6.2E-130 |
| HMMR | 0.479446 | 1.2E-80 |
| FAM111A | 0.479382 | 9.54E-84 |
| BUB3 | 0.478819 | 4.42E-91 |
| DHFR | 0.469849 | 9.04E-86 |
| MGME1 | 0.464719 | 9.22E-97 |
| MIS18A | 0.462084 | 4.5E-102 |
| DBF4 | 0.457651 | 1.41E-77 |
| COMMD4 | 0.456086 | 6.3E-106 |
| FEN1 | 0.455897 | 6.75E-98 |
| PTMA | 0.4545 | 6E-302 |
| CDK4 | 0.451908 | 3.9E-126 |
| NCAPD2 | 0.450183 | 4.35E-93 |
| TTK | 0.449635 | 5.2E-87 |
| PRKDC | 0.448391 | 4.5E-100 |
| CDK5RAP! | 0.447597 | 7.22E-82 |
| SIVA1 | 0.447518 | 3.9E-130 |
| PSMC3 | 0.447017 | 1.2E-116 |
| SUPT16H | 0.444171 | 1.7E-116 |
| LMNB2 | 0.439625 | 2.5E-94 |
| DEPDC1 | 0.439577 | 1.94E-81 |
| CHEK1 | 0.436931 | 3.37E-85 |

| | | |
|---|---|---|
| HNRNPD | 0.435631 | 4.9E-123 |
| ECT2 | 0.435383 | 8.26E-75 |
| KIF20A | 0.432917 | 1.55E-72 |
| ATAD5 | 0.431859 | 1.22E-91 |
| RACGAP1 | 0.431201 | 6.43E-89 |
| CDCA2 | 0.430108 | 6.39E-77 |
| PRR11 | 0.429967 | 8.18E-72 |
| POLD3 | 0.429919 | 1.58E-83 |
| PKMYT1 | 0.426929 | 6.97E-92 |
| JPT1 | 0.426198 | 3.85E-72 |
| LIG1 | 0.425378 | 1.05E-87 |
| MELK | 0.424351 | 1.24E-81 |
| SAC3D1 | 0.422768 | 9.9E-97 |
| SRSF3 | 0.419272 | 1.3E-175 |
| RFC2 | 0.417479 | 5.08E-83 |
| HMGXB4 | 0.41721 | 1.47E-97 |
| CENPX | 0.41653 | 3.29E-86 |
| KIF4A | 0.414284 | 7.97E-80 |
| AURKA | 0.411242 | 4.24E-58 |
| SRSF2 | 0.410907 | 2.7E-105 |
| SMC1A | 0.410903 | 2.84E-71 |
| BARD1 | 0.410686 | 2.37E-74 |
| TMEM97 | 0.409769 | 1.64E-84 |
| SKA3 | 0.407856 | 5.13E-87 |
| PRC1 | 0.40657 | 4.54E-83 |
| CHAF1A | 0.403851 | 1.32E-83 |
| DDX39A | 0.400859 | 5.81E-74 |
| DSN1 | 0.40079 | 5.69E-85 |
| HIST1H2A | 0.399914 | 1.53E-82 |
| BRCA1 | 0.399557 | 6.19E-76 |
| CKAP5 | 0.396115 | 8.16E-80 |
| VRK1 | 0.395027 | 1.76E-82 |
| PHGDH | 0.392144 | 1.9E-114 |
| NSD2 | 0.391397 | 3.72E-74 |
| PSRC1 | 0.390864 | 4.29E-54 |
| BANF1 | 0.389184 | 6.9E-141 |
| CRABP1 | 0.387917 | 2.4E-123 |
| PTX3 | 0.38771 | 4.58E-45 |
| HJURP | 0.387154 | 2.06E-72 |
| E2F8 | 0.383862 | 2.97E-75 |
| RMI2 | 0.383154 | 2.49E-81 |
| SRSF10 | 0.38077 | 1.17E-91 |
| CNTLN | 0.380639 | 2.04E-82 |
| RFC3 | 0.379825 | 5.8E-74 |
| CDKN2D | 0.377815 | 4.47E-58 |
| TROAP | 0.376259 | 4.47E-56 |
| LGALS1 | 0.376096 | 9.6E-133 |
| CBX1 | 0.376062 | 1.2E-109 |
| FABP5 | 0.37492 | 3.1E-86 |
| MCM10 | 0.374696 | 1.65E-76 |
| SYNE2 | 0.369784 | 3.68E-65 |
| B2M | 0.367437 | 2.02E-74 |
| H3F3A | 0.366963 | 3.9E-264 |
| SRSF7 | 0.366923 | 7.3E-109 |
| C19orf48 | 0.365607 | 9.87E-80 |
| HIST1H4E | 0.365321 | 4.37E-71 |

| | | |
|---|---|---|
| MCM3 | 0.364611 | 4.27E-62 |
| SNRPB | 0.364276 | 1.1E-100 |
| CBX3 | 0.363769 | 2.8E-108 |
| SKA1 | 0.363504 | 1.2E-61 |
| EMILIN1 | 0.362892 | 1.9E-103 |
| LRR1 | 0.362089 | 6.53E-71 |
| MZT1 | 0.361146 | 5.96E-67 |
| ZNF738 | 0.359006 | 7.12E-73 |
| NCAPH | 0.356794 | 1.96E-77 |
| POC1A | 0.35661 | 4.5E-86 |
| NRM | 0.356408 | 1.39E-65 |
| HMGN1 | 0.356339 | 4.3E-151 |
| SNRPG | 0.355822 | 7.72E-95 |
| CEP78 | 0.35511 | 3.1E-66 |
| CDC25C | 0.354195 | 4.26E-78 |
| COL1A1 | 0.352963 | 4.7E-178 |
| MCM4 | 0.352065 | 3.56E-58 |
| HNRNPA3 | 0.350845 | 2.8E-111 |
| HP1BP3 | 0.349747 | 6.03E-72 |
| LSM3 | 0.349625 | 3.02E-88 |
| TPM2 | 0.349602 | 2.8E-135 |
| BLM | 0.349411 | 1.45E-69 |
| SNRPD1 | 0.349266 | 1.3E-100 |
| RAN | 0.348993 | 5.4E-116 |
| C21orf58 | 0.347352 | 1.25E-73 |
| CENPL | 0.346419 | 5.76E-60 |
| CENPJ | 0.345363 | 2.89E-68 |
| RPL39L | 0.344668 | 5.13E-64 |
| HSPB11 | 0.342807 | 2.83E-68 |
| PBK | 0.340783 | 3.72E-47 |
| NAP1L1 | 0.339851 | 7.1E-127 |
| CENPV | 0.338687 | 2.9E-53 |
| PLK1 | 0.338528 | 3.21E-48 |
| GGH | 0.337122 | 7.1E-70 |
| XRCC2 | 0.336062 | 5.05E-64 |
| CSRP2 | 0.335868 | 4.18E-55 |
| SPAG5 | 0.334596 | 6.78E-65 |
| CEP152 | 0.334504 | 7.49E-67 |
| EMP3 | 0.333616 | 3.5E-97 |
| PCOLCE | 0.332755 | 3.08E-78 |
| NFIC | 0.332695 | 1.9E-75 |
| SAP30 | 0.330586 | 1.85E-60 |
| SEPT10 | 0.327338 | 4.22E-57 |
| PRRX1 | 0.32731 | 3.32E-90 |
| CTHRC1 | 0.327119 | 5.27E-50 |
| CCDC14 | 0.325678 | 1.37E-70 |
| ERH | 0.325653 | 9.9E-123 |
| KIF18A | 0.324404 | 6.35E-54 |
| RHNO1 | 0.323448 | 3.77E-61 |
| RBBP7 | 0.32341 | 6.98E-61 |
| MAD2L2 | 0.323227 | 1.36E-69 |
| SPDL1 | 0.32183 | 1.16E-58 |
| PHF19 | 0.321136 | 2.13E-61 |
| H2AFY | 0.319595 | 8.31E-73 |
| TPGS2 | 0.319161 | 3.29E-65 |
| E2F7 | 0.318102 | 4.54E-64 |

| | | |
|---|---|---|
| COL9A3 | 0.317813 | 1.65E-70 |
| KNSTRN | 0.316094 | 4.61E-48 |
| BAZ1B | 0.315563 | 5.02E-64 |
| TWIST1 | 0.315419 | 1.48E-48 |
| HNRNPAB | 0.314385 | 9.15E-58 |
| XRCC5 | 0.312693 | 5.51E-92 |
| HADH | 0.311666 | 1.46E-61 |
| NCAPH2 | 0.310394 | 3.35E-62 |
| PTGES3 | 0.310047 | 3.13E-96 |
| NCAPG2 | 0.309921 | 6.61E-59 |
| MMS22L | 0.309691 | 7.32E-53 |
| FANCD2 | 0.309674 | 4.6E-61 |
| PRTFDC1 | 0.309504 | 3.69E-54 |
| CCDC18 | 0.308599 | 4.49E-52 |
| ANAPC11 | 0.308444 | 2.23E-58 |
| HIRIP3 | 0.308242 | 4.89E-54 |
| FANCI | 0.306904 | 2.2E-62 |
| PLEKHO1 | 0.306653 | 3.21E-76 |
| DKK1 | 0.306503 | 2.02E-06 |
| ZNF93 | 0.306278 | 1.08E-50 |
| NSMCE4A | 0.305141 | 8.99E-63 |
| RPA2 | 0.304092 | 4.27E-52 |
| EXOSC8 | 0.303097 | 5.27E-57 |
| TRIP13 | 0.302583 | 9.9E-63 |
| CDC45 | 0.302009 | 4.97E-59 |
| UQCC2 | 0.300766 | 1.25E-62 |
| SEPT7 | 0.300594 | 6.3E-102 |
| CSE1L | 0.300512 | 4.22E-61 |
| PMP22 | 0.299996 | 2.52E-58 |
| MAB21L2 | 0.299426 | 1.8E-75 |
| RPA3 | 0.298525 | 1.27E-62 |
| MZT2B | 0.298342 | 1.2E-104 |
| PRIM1 | 0.298154 | 2.79E-55 |
| FUS | 0.297633 | 1.85E-81 |
| NFIB | 0.297267 | 1.65E-70 |
| HIST1H2B | 0.296832 | 1.14E-55 |
| MASTL | 0.296435 | 4.1E-59 |
| MAB21L1 | 0.295955 | 6.39E-48 |
| WDR34 | 0.295457 | 1.04E-56 |
| HIST1H3G | 0.295142 | 2.74E-57 |
| TMEM237 | 0.294218 | 3.26E-55 |
| TCF19 | 0.292889 | 7.74E-61 |
| OIP5 | 0.292632 | 9.96E-61 |
| EEF1B2 | 0.292135 | 8.6E-110 |
| BRIP1 | 0.291967 | 1.01E-57 |
| NUP62 | 0.289595 | 2.26E-55 |
| C1QL1 | 0.289335 | 2.02E-33 |
| RTKN2 | 0.289257 | 1.13E-51 |
| UHRF1 | 0.288846 | 7.46E-54 |
| ARHGAP1 | 0.286478 | 2.16E-52 |
| RFC1 | 0.285959 | 5.7E-53 |
| SMCHD1 | 0.285847 | 5.9E-55 |
| KDELC2 | 0.285816 | 3.82E-48 |
| MDFI | 0.285725 | 2.28E-92 |
| SNRNP40 | 0.28565 | 3.01E-54 |
| GAS2 | 0.285605 | 8.35E-35 |

| | | |
|---|---|---|
| RALY | 0.285135 | 5.02E-66 |
| BUB1 | 0.285107 | 1.03E-46 |
| FANCL | 0.284598 | 1.56E-52 |
| PSMC3IP | 0.283483 | 2.65E-55 |
| ARL6IP1 | 0.283469 | 2.54E-24 |
| TCEAL7 | 0.282524 | 2.12E-44 |
| WDR76 | 0.281755 | 1.98E-49 |
| COL1A2 | 0.281355 | 3.9E-158 |
| HSP90AA1 | 0.281174 | 6.35E-68 |
| HAUS8 | 0.280565 | 2.39E-52 |
| FANCG | 0.279982 | 1.96E-54 |
| TPR | 0.279849 | 2.03E-57 |
| PRRX2 | 0.279189 | 2.93E-49 |
| CDC6 | 0.279095 | 1.75E-53 |
| NSD3 | 0.2789 | 1.74E-64 |
| WDHD1 | 0.278764 | 1.65E-49 |
| HNRNPH3 | 0.278486 | 1.5E-73 |
| XPO1 | 0.277584 | 2.45E-50 |
| RAC3 | 0.276422 | 1.23E-48 |
| ELN | 0.276382 | 1.41E-54 |
| SCARA3 | 0.276029 | 7.43E-51 |
| GPSM2 | 0.275257 | 7.53E-46 |
| YBX1 | 0.274797 | 2.5E-126 |
| PARP2 | 0.274551 | 6.88E-47 |
| TPM4 | 0.273562 | 3.56E-53 |
| MYH10 | 0.273553 | 5.7E-54 |
| SLBP | 0.273517 | 1.6E-33 |
| DTL | 0.27342 | 3.35E-52 |
| EVL | 0.271559 | 2.2E-65 |
| MAGOHB | 0.270787 | 1.64E-52 |
| CEP135 | 0.270503 | 1.19E-50 |
| RNASEH2( | 0.269505 | 4.29E-44 |
| HPF1 | 0.268969 | 1.35E-54 |
| SHCBP1 | 0.26848 | 1.82E-48 |
| RBBP4 | 0.268342 | 1.53E-57 |
| HDGFL3 | 0.26771 | 4.09E-49 |
| ILF2 | 0.266481 | 4.37E-85 |
| SIX1 | 0.266421 | 2.23E-47 |
| FIGNL1 | 0.266332 | 2.26E-50 |
| CASP8AP2 | 0.265283 | 6.26E-43 |
| E2F1 | 0.263436 | 4.1E-43 |
| BCL2L12 | 0.263299 | 3.94E-48 |
| HNRNPA0 | 0.263211 | 5.21E-63 |
| SNRPE | 0.262746 | 5.78E-74 |
| RPLP1 | 0.262578 | 2.6E-190 |
| MRPL51 | 0.262278 | 2.36E-73 |
| ACAT2 | 0.261775 | 4.99E-34 |
| SNRPD3 | 0.26037 | 1.18E-73 |
| INCENP | 0.260182 | 9.39E-49 |
| SSRP1 | 0.260058 | 1.15E-45 |
| BRCA2 | 0.260001 | 1.61E-48 |
| NCAM1 | 0.259862 | 5.69E-42 |
| CEP57L1 | 0.25768 | 2.01E-43 |
| EXOSC9 | 0.257401 | 1.67E-45 |
| TIMELESS | 0.257347 | 7.56E-45 |
| IDH2 | 0.256452 | 1.06E-47 |

| | | |
|---|---|---|
| PAXX | 0.256259 | 6.4E-50 |
| CENPQ | 0.255781 | 2.11E-49 |
| TRIM59 | 0.255403 | 5.93E-42 |
| PCP4 | 0.254923 | 2.77E-05 |
| PXMP2 | 0.254913 | 5.73E-40 |
| HIST1H2B | 0.254767 | 1.06E-45 |
| CEP295 | 0.254066 | 4.36E-40 |
| LSM5 | 0.250736 | 2.56E-54 |
| C1QTNF2 | 0.250677 | 1.91E-42 |
| TRA2B | 0.250465 | 9.45E-52 |
| PPIH | 0.250188 | 1.89E-49 |
| CTCF | 0.250177 | 4.14E-39 |
| FXYD6 | 0.250076 | 5.13E-41 |

| **Cluster 11** | | |
|---|---|---|
| Gene | LogFC | pVal |
| TAGLN | 2.367005 | 4.3E-176 |
| ACTA2 | 2.085162 | 7.04E-77 |
| THY1 | 1.543925 | 2.1E-167 |
| COL6A3 | 1.530743 | 1.6E-211 |
| MMP9 | 1.410596 | 9.36E-39 |
| LGALS3 | 1.400125 | 1.1E-130 |
| TPM2 | 1.265594 | 2.8E-220 |
| BGN | 1.192649 | 2.3E-148 |
| TIMP1 | 1.171592 | 1.5E-140 |
| TPM4 | 1.090253 | 3E-199 |
| STMN2 | 0.954643 | 1.1E-27 |
| C11orf96 | 0.947892 | 6.82E-65 |
| CALD1 | 0.873836 | 8.5E-192 |
| PDGFRB | 0.872583 | 7.28E-98 |
| AC083967. | 0.868885 | 1.09E-19 |
| NPW | 0.867549 | 1.53E-91 |
| RGS5 | 0.849974 | 5.61E-18 |
| LGALS1 | 0.832938 | 1.3E-191 |
| EDNRA | 0.829848 | 1.8E-123 |
| SULF1 | 0.824952 | 7.55E-84 |
| SEPT11 | 0.821407 | 2.1E-155 |
| TSHZ2 | 0.789312 | 2.2E-128 |
| PLAC9 | 0.762317 | 6.17E-53 |
| COL4A1 | 0.75943 | 1.95E-61 |
| CSRP1 | 0.758411 | 8.5E-87 |
| TMSB4X | 0.758053 | 1.3E-132 |
| HES4 | 0.754571 | 3.86E-97 |
| SPARCL1 | 0.751997 | 7.94E-64 |
| ACTB | 0.738651 | 4E-173 |
| MYL9 | 0.720923 | 6.29E-91 |
| NR2F2 | 0.711482 | 7.69E-97 |
| LMNA | 0.699274 | 1.91E-92 |
| HEY1 | 0.669637 | 3.35E-59 |
| CHST2 | 0.666084 | 1.13E-80 |
| ARPC5 | 0.665515 | 3.93E-80 |
| FILIP1L | 0.656607 | 8.06E-38 |
| ZEB2 | 0.655179 | 1.71E-81 |
| ANXA1 | 0.653158 | 3.06E-41 |
| MAP1B | 0.650944 | 7.65E-75 |
| IFITM3 | 0.647982 | 3E-124 |
| COL6A2 | 0.635883 | 1.4E-123 |
| MGP | 0.633067 | 4.39E-69 |
| VIM | 0.622289 | 1.7E-120 |
| CALM1 | 0.621334 | 1.36E-78 |
| TPM1 | 0.618122 | 5.47E-44 |
| SLC26A7 | 0.617971 | 1.4E-30 |
| KCNE4 | 0.604612 | 4.86E-56 |
| ARPC2 | 0.60431 | 1.4E-111 |
| TGFB1I1 | 0.595735 | 3.51E-92 |
| MIR4435-2 | 0.579652 | 7.37E-44 |
| MCAM | 0.57695 | 2.83E-47 |
| NGFR | 0.563337 | 2.32E-67 |
| COL3A1 | 0.55276 | 1.2E-112 |
| LAMA4 | 0.544337 | 1.35E-73 |

| | | |
|---|---|---|
| CYTOR | 0.536817 | 2.79E-56 |
| PGM5-AS1 | 0.531747 | 3.42E-13 |
| CAVIN3 | 0.530508 | 1.05E-55 |
| RAP1B | 0.530144 | 1.82E-90 |
| ACTN1 | 0.528941 | 1.42E-76 |
| PRKG1 | 0.522514 | 9.14E-72 |
| TUBA1A | 0.518825 | 8.57E-46 |
| FHL3 | 0.515673 | 7.63E-75 |
| ZYX | 0.51457 | 3.19E-75 |
| KHDRBS3 | 0.51186 | 1.8E-29 |
| CNN3 | 0.50746 | 4E-112 |
| CDC42EP5 | 0.4965 | 1.7E-123 |
| KRT17 | 0.493601 | 2.38E-15 |
| EBF1 | 0.491497 | 2.19E-63 |
| CRABP2 | 0.486692 | 1.63E-46 |
| PFN1 | 0.482176 | 3.12E-91 |
| SERPINH1 | 0.480971 | 9.23E-97 |
| MIR181A1I | 0.475926 | 3.33E-58 |
| SEMA5A | 0.475058 | 8.47E-40 |
| ALDH1A2 | 0.474592 | 1.32E-23 |
| SEPT7 | 0.471297 | 7.4E-112 |
| PFN2 | 0.469382 | 2.04E-51 |
| SELENOW | 0.466881 | 9.94E-84 |
| CD248 | 0.466234 | 2.01E-65 |
| TNFRSF1A | 0.465752 | 6.73E-61 |
| B2M | 0.463923 | 1.19E-55 |
| COL1A1 | 0.46343 | 3.3E-201 |
| FKBP1A | 0.462994 | 1.87E-77 |
| VCL | 0.461858 | 1.15E-62 |
| SH3BGRL3 | 0.45949 | 2.22E-75 |
| MYLK | 0.457236 | 7.93E-61 |
| FABP5 | 0.456112 | 2.87E-28 |
| EIF4EBP1 | 0.453281 | 2.56E-63 |
| CCL2 | 0.448137 | 5.03E-07 |
| IGF2.1 | 0.447343 | 1.04E-53 |
| OLFML3 | 0.447169 | 6.62E-60 |
| IFITM2 | 0.446923 | 8.82E-53 |
| MEIS3 | 0.446699 | 1.17E-55 |
| PRSS35 | 0.442884 | 3.76E-15 |
| PCOLCE | 0.442546 | 6.4E-62 |
| TUBB6 | 0.44054 | 3.35E-53 |
| S100A11 | 0.433592 | 1.97E-82 |
| TUBA1B | 0.433065 | 4.63E-43 |
| C9orf16 | 0.432774 | 1.45E-74 |
| NRGN | 0.425901 | 4.65E-49 |
| FLNA | 0.422965 | 3.96E-58 |
| LOXL2 | 0.419417 | 1.6E-54 |
| TTYH1 | 0.418656 | 6.41E-30 |
| IFI16 | 0.418577 | 1.49E-55 |
| MAP4K4 | 0.417819 | 1.44E-54 |
| CPE | 0.416817 | 6.69E-95 |
| CCDC102B | 0.415638 | 2.23E-24 |
| ACTG1 | 0.414663 | 1.2E-103 |
| MAGED2 | 0.41281 | 1.35E-83 |
| AKAP12 | 0.412682 | 9.07E-57 |
| PCDH18 | 0.410819 | 4.06E-51 |

| | | |
|---|---|---|
| COTL1 | 0.410617 | 9.94E-45 |
| TNC | 0.410201 | 6.6E-19 |
| RASL12 | 0.409214 | 4.51E-49 |
| TIMP3 | 0.408618 | 3.04E-18 |
| LRRC4C | 0.40737 | 7.9E-21 |
| FERMT2 | 0.406848 | 8.25E-51 |
| GLIPR2 | 0.405738 | 1.89E-50 |
| PLVAP | 0.405239 | 4.97E-25 |
| CTHRC1 | 0.404261 | 6.9E-21 |
| GSN | 0.403727 | 1.15E-64 |
| SRM | 0.398926 | 7.76E-61 |
| GBP1 | 0.397855 | 3.84E-39 |
| PDLIM1 | 0.396778 | 1.8E-17 |
| CFL1 | 0.394861 | 2E-110 |
| BZW1 | 0.393529 | 9.67E-68 |
| CTSC | 0.390614 | 3.84E-15 |
| PHLDA1 | 0.387139 | 1.99E-34 |
| INAFM1 | 0.386643 | 8.35E-43 |
| EMP2 | 0.38551 | 2.59E-46 |
| TBX2 | 0.383827 | 1.97E-28 |
| MYL12A | 0.383654 | 9.61E-52 |
| ANXA2 | 0.382543 | 1.86E-46 |
| NOTCH3 | 0.378686 | 1.25E-53 |
| TSPAN5 | 0.376479 | 5.49E-39 |
| SDC2 | 0.374827 | 8.14E-43 |
| PRSS23 | 0.37406 | 1.51E-27 |
| IER3 | 0.373184 | 3.12E-07 |
| HIC1 | 0.372852 | 1.04E-34 |
| EIF5A | 0.372666 | 1.11E-49 |
| COL4A2 | 0.37214 | 3.34E-23 |
| MTCH1 | 0.3719 | 1.03E-64 |
| GUCY1B1 | 0.371072 | 3.62E-40 |
| IGFBP3 | 0.369861 | 1.82E-05 |
| KCNJ8 | 0.369135 | 8.09E-21 |
| RBM3 | 0.364949 | 4.26E-65 |
| FOXS1 | 0.364923 | 9.91E-39 |
| PLCL1 | 0.363248 | 3.1E-37 |
| ETS1 | 0.362802 | 9.94E-47 |
| CD63 | 0.361199 | 8.33E-75 |
| SEMA3A | 0.357746 | 3.91E-44 |
| C1orf54 | 0.35671 | 1.27E-59 |
| LIMA1 | 0.354141 | 5.19E-37 |
| ENG | 0.353647 | 1.71E-51 |
| NHP2 | 0.349116 | 8.54E-47 |
| LY6E | 0.346834 | 2.16E-37 |
| PTGIR | 0.346476 | 3.19E-40 |
| LHFPL6 | 0.343971 | 4.04E-37 |
| TFPI | 0.343229 | 2.12E-31 |
| MEST | 0.341696 | 3.28E-76 |
| A2M | 0.341035 | 4.96E-36 |
| CNN2 | 0.339479 | 3.56E-43 |
| ACTG2 | 0.338875 | 2.84E-10 |
| GPM6B | 0.336087 | 2.18E-34 |
| VCAN | 0.333918 | 5.42E-24 |
| ERRFI1 | 0.333342 | 3.8E-24 |
| EHD2 | 0.333308 | 7.01E-49 |

| | | |
|---|---|---|
| MLLT11 | 0.331043 | 1.46E-37 |
| CLEC11A | 0.330208 | 3.83E-14 |
| COL6A1 | 0.328587 | 8.71E-38 |
| MAFB | 0.32824 | 1.61E-23 |
| SEPT9 | 0.326883 | 2.33E-39 |
| CTGF | 0.326396 | 6.63E-10 |
| AHNAK | 0.324405 | 2.53E-45 |
| ACAT2 | 0.323047 | 1.66E-13 |
| CRMP1 | 0.322944 | 1.25E-39 |
| VCAM1 | 0.32024 | 5.01E-19 |
| MAP1LC3A | 0.318468 | 8.79E-40 |
| EMCN | 0.315431 | 3.76E-30 |
| SPARC | 0.312881 | 2.03E-48 |
| RSU1 | 0.311068 | 1.24E-34 |
| NUPR1 | 0.310884 | 3.08E-23 |
| NME1 | 0.309996 | 3.51E-26 |
| MARCKS | 0.309274 | 6.61E-91 |
| PHLDA2 | 0.309189 | 2.34E-26 |
| DKK1 | 0.304812 | 2.06E-06 |
| APOC1 | 0.304789 | 6.71E-27 |
| YBX1 | 0.303373 | 1.47E-70 |
| TSC22D4 | 0.302457 | 3.83E-34 |
| IFI27L2 | 0.301806 | 1.08E-29 |
| SSBP4 | 0.300598 | 1.1E-38 |
| SH2D3C | 0.299368 | 1.31E-44 |
| EMP3 | 0.297269 | 9.27E-49 |
| RPL22L1 | 0.296787 | 1.25E-27 |
| SHMT2 | 0.296104 | 2.91E-44 |
| RHOC | 0.295937 | 6.2E-54 |
| ACTR3 | 0.29593 | 1.87E-32 |
| PSMA7 | 0.293544 | 2.69E-53 |
| PDE1A | 0.293294 | 8.88E-20 |
| TLN1 | 0.292968 | 6.63E-33 |
| RNF24 | 0.292742 | 8.69E-37 |
| CDC42EP1 | 0.291104 | 7.29E-37 |
| FJX1 | 0.290411 | 2.12E-16 |
| MYADM | 0.28933 | 4.13E-21 |
| SPON2 | 0.289314 | 4.04E-14 |
| RAMP2 | 0.286752 | 2.28E-37 |
| VASN | 0.285467 | 8.45E-20 |
| SMARCA5 | 0.285434 | 2.12E-24 |
| LARP6 | 0.284273 | 1.36E-29 |
| TCEAL7 | 0.284225 | 4.72E-30 |
| RPLP1 | 0.282242 | 4.9E-115 |
| SNAI2 | 0.282213 | 7.16E-26 |
| EGFR | 0.281476 | 1.31E-30 |
| BASP1 | 0.281116 | 4.91E-35 |
| LIMD2 | 0.281108 | 6.3E-32 |
| ISG15 | 0.278833 | 8.86E-33 |
| CD81 | 0.278684 | 1.31E-52 |
| NDUFA4L2 | 0.277913 | 3.94E-15 |
| C12orf57 | 0.276978 | 1.74E-58 |
| CYGB | 0.276902 | 2.44E-21 |
| EMILIN1 | 0.275776 | 1.21E-42 |
| L1TD1 | 0.275461 | 2.95E-26 |
| NRP2 | 0.275081 | 4.3E-13 |

| | | |
|---|---|---|
| EIF4A1 | 0.274079 | 1.22E-35 |
| GNAI2 | 0.273679 | 1.35E-41 |
| KCNQ2 | 0.273389 | 2.36E-29 |
| EPHX1 | 0.273373 | 2.88E-33 |
| HNRNPAB | 0.273191 | 5.34E-23 |
| DRAP1 | 0.27254 | 9.98E-37 |
| PMEPA1 | 0.272119 | 1.54E-27 |
| ITGA4 | 0.271836 | 8.42E-33 |
| MYC | 0.270945 | 3.9E-21 |
| SLIT3 | 0.268596 | 4.22E-30 |
| NDUFS5 | 0.267645 | 2.99E-57 |
| CARMN | 0.267342 | 7.59E-35 |
| TGFB1 | 0.266744 | 1.93E-28 |
| ATP2B1 | 0.266224 | 5.63E-19 |
| LITAF | 0.266164 | 2.16E-35 |
| ROCK2 | 0.265488 | 9.63E-26 |
| ANXA6 | 0.265337 | 9.44E-38 |
| YWHAB | 0.26335 | 6.01E-45 |
| PPIB | 0.262583 | 1.98E-32 |
| NTAN1 | 0.26166 | 5.24E-36 |
| TPM3 | 0.261098 | 1.85E-27 |
| RPS18 | 0.260361 | 1.29E-69 |
| POMP | 0.259796 | 3.78E-31 |
| SHC1 | 0.259251 | 9.52E-34 |
| NREP | 0.257984 | 1.32E-17 |
| TAGLN2 | 0.257667 | 8.69E-09 |
| PDS5B | 0.257265 | 1.43E-30 |
| PHLDA3 | 0.257089 | 3.62E-32 |
| DCLK1 | 0.256698 | 4.53E-29 |
| RPL12 | 0.256064 | 4.4E-52 |
| ARPC4 | 0.254733 | 2.42E-31 |
| SLC25A6 | 0.254563 | 1.7E-42 |
| TLE1 | 0.254464 | 2.76E-24 |
| INKA1 | 0.253549 | 1.33E-31 |
| RPL27 | 0.25352 | 2.32E-92 |
| DES | 0.252463 | 7E-13 |
| VASP | 0.252249 | 6.69E-21 |
| ITGA5 | 0.251921 | 2.87E-28 |
| STOM | 0.251351 | 1.39E-20 |
| BCL7C | 0.250777 | 1.67E-28 |
| AP2S1 | 0.250145 | 7.1E-26 |

| Cluster 12 | | |
|---|---|---|
| Gene | LogFC | pVal |
| NPHS2 | 3.420277 | 6.8E-244 |
| TCF21 | 2.981554 | 1.8E-155 |
| MAFB | 2.725076 | 3E-293 |
| ANXA1 | 2.615424 | 1.7E-164 |
| CPXM1 | 2.552334 | 9E-208 |
| SOST | 2.550826 | 5.2E-169 |
| BST2 | 2.527497 | 1.5E-173 |
| GADD45A | 2.470151 | 7.6E-161 |
| PODXL | 2.343459 | 6.7E-166 |
| PTPRO | 2.241664 | 2.8E-167 |
| CTGF | 2.189224 | 3.05E-45 |
| PLTP | 2.157053 | 8.3E-159 |
| AIF1 | 2.110502 | 1E-146 |
| BCAM | 2.092466 | 8E-202 |
| CLIC5 | 2.05414 | 1.2E-171 |
| WT1 | 1.949174 | 7.5E-183 |
| DDN | 1.90585 | 3.7E-99 |
| SBSPON | 1.893734 | 1.4E-142 |
| TJP1 | 1.789538 | 8.1E-173 |
| MYL9 | 1.737985 | 6.9E-154 |
| FOXC2 | 1.679741 | 1.3E-151 |
| MXRA8 | 1.660304 | 2.9E-159 |
| ITM2B | 1.638011 | 6.8E-146 |
| TNNI1 | 1.631773 | 1.34E-97 |
| CLDN5 | 1.628381 | 2.1E-132 |
| SPOCK2 | 1.621108 | 1.8E-124 |
| HTRA1 | 1.615404 | 1E-129 |
| MPP5 | 1.548671 | 1.4E-106 |
| TGFBR3 | 1.525531 | 8.7E-115 |
| SLC9A3R2 | 1.51958 | 9.2E-145 |
| ST3GAL6 | 1.496063 | 7.6E-115 |
| PTH1R | 1.478964 | 4.1E-122 |
| ST6GALNA | 1.476619 | 2.4E-106 |
| SPINT2 | 1.473674 | 7.7E-157 |
| VEGFA | 1.466294 | 9.2E-110 |
| TPPP3 | 1.447171 | 7.41E-97 |
| AQP3 | 1.437687 | 3.47E-95 |
| TARID | 1.406499 | 5.9E-103 |
| MIR503HG | 1.399527 | 1.26E-87 |
| FOXD1 | 1.377615 | 1.64E-78 |
| NES | 1.362524 | 1.34E-96 |
| PDLIM2 | 1.340267 | 6.4E-116 |
| EIF3M | 1.337966 | 5.1E-141 |
| MAGI2 | 1.313203 | 3.6E-94 |
| NPHS1 | 1.30385 | 8E-110 |
| EPHX1 | 1.285016 | 8.4E-108 |
| TNNT2 | 1.275022 | 1.01E-73 |
| TYRO3 | 1.250391 | 2E-101 |
| DUSP23 | 1.24083 | 4.91E-82 |
| SEPT11 | 1.234092 | 2.7E-120 |
| TSPAN2 | 1.214252 | 8.57E-67 |
| MME | 1.206933 | 3.26E-47 |
| ENPEP | 1.180365 | 6.87E-77 |
| ARHGEF3 | 1.177253 | 3.24E-88 |

| | | |
|---|---|---|
| IGFBP7 | 1.17041 | 2.74E-72 |
| SAMD11 | 1.160541 | 2.42E-53 |
| TSC22D3 | 1.156032 | 6.76E-72 |
| CAST | 1.147908 | 3.78E-95 |
| TUBB2A | 1.145675 | 1.63E-95 |
| ITIH5 | 1.144381 | 5.21E-75 |
| PDE4DIP | 1.125784 | 1.04E-88 |
| STON2 | 1.125544 | 3.25E-87 |
| S100A6 | 1.124258 | 3.6E-68 |
| KCNH2 | 1.11189 | 2.22E-98 |
| VAMP5 | 1.111286 | 4.08E-87 |
| LEPROT | 1.096015 | 8.43E-95 |
| MRGPRF | 1.095811 | 1.61E-79 |
| VAMP8 | 1.083827 | 7.3E-103 |
| NECTIN2 | 1.075402 | 1.26E-92 |
| SELENOP | 1.07302 | 1.5E-104 |
| SERINC5 | 1.069036 | 2.86E-78 |
| ARMH4 | 1.054796 | 2.01E-57 |
| PLA2R1 | 1.048003 | 6.06E-65 |
| PLD3 | 1.028013 | 1.72E-97 |
| SLC48A1 | 1.027763 | 8.88E-78 |
| SEMA3B | 1.023094 | 1.78E-79 |
| GAS6 | 1.021838 | 4.34E-93 |
| TMEM178/ | 1.019926 | 1.12E-84 |
| PON2 | 1.005131 | 3.18E-79 |
| ANXA2 | 0.982865 | 4.86E-78 |
| CTNNAL1 | 0.977085 | 6.49E-80 |
| SDC2 | 0.958154 | 1.96E-76 |
| P3H2 | 0.947818 | 1.48E-81 |
| PRDX6 | 0.938344 | 1.74E-88 |
| ARHGAP2 | 0.926581 | 5.88E-59 |
| THSD7A | 0.926225 | 2.14E-50 |
| PLAC1 | 0.925314 | 3.03E-66 |
| RBMS1 | 0.923427 | 9.07E-76 |
| EFNB1 | 0.922586 | 2.44E-74 |
| ITGA3 | 0.920858 | 1.32E-64 |
| IQGAP2 | 0.91959 | 6.44E-57 |
| FRMD1 | 0.912325 | 1.37E-66 |
| DACH1 | 0.910781 | 1.98E-72 |
| ZFP36L2 | 0.903303 | 3.54E-60 |
| DDR1 | 0.901245 | 1.92E-75 |
| PABPN1 | 0.896241 | 3.19E-72 |
| COMT | 0.891485 | 2.48E-77 |
| TSPAN3 | 0.891024 | 1.39E-71 |
| GSN | 0.890567 | 6.09E-84 |
| ARHGAP2 | 0.889763 | 4.99E-70 |
| VAMP2 | 0.884847 | 2.28E-68 |
| RETREG1 | 0.870642 | 1.31E-62 |
| TRIM8 | 0.86677 | 7.44E-72 |
| ROBO2 | 0.865214 | 6.76E-61 |
| NPW | 0.86251 | 5.8E-19 |
| TMEM266 | 0.847232 | 1.98E-63 |
| SPARC | 0.843369 | 2.72E-72 |
| RRAS | 0.836834 | 3.29E-71 |
| HIPK2 | 0.819111 | 2.5E-48 |
| KIRREL1 | 0.818191 | 2.74E-64 |

| | | |
|---|---|---|
| CTTNBP2 | 0.81512 | 1.13E-63 |
| NTNG1 | 0.812326 | 3.52E-40 |
| LEFTY1 | 0.811051 | 2.91E-30 |
| F2R | 0.810276 | 4.73E-53 |
| CD59 | 0.808889 | 1.47E-64 |
| TCEAL3 | 0.803812 | 3.44E-54 |
| RASSF8-A | 0.802378 | 1.48E-57 |
| SRGAP1 | 0.801857 | 3.38E-61 |
| VASN | 0.800793 | 8.99E-57 |
| TCEAL2 | 0.796764 | 5.49E-38 |
| RASSF8 | 0.791736 | 3.8E-57 |
| HIST3H2A | 0.788854 | 3.77E-51 |
| FYN | 0.786246 | 8.58E-57 |
| CD2AP | 0.782594 | 2.41E-48 |
| S100A4 | 0.780648 | 2.95E-30 |
| KLK6 | 0.779058 | 3.28E-50 |
| GPC3 | 0.778525 | 8.31E-42 |
| F3 | 0.775368 | 3.23E-46 |
| RIPOR1 | 0.774058 | 2.15E-62 |
| UNC119 | 0.767302 | 6.07E-62 |
| TM4SF1 | 0.761357 | 3.15E-24 |
| AIF1L | 0.760176 | 1.18E-68 |
| REEP5 | 0.759817 | 6.31E-63 |
| KRT19 | 0.755148 | 3.47E-25 |
| ACTN4 | 0.745835 | 2.16E-42 |
| FOXD2 | 0.744962 | 8.33E-52 |
| C6orf203 | 0.744549 | 4.26E-56 |
| G0S2 | 0.742328 | 2.52E-13 |
| NPNT | 0.741489 | 2.53E-37 |
| PAPPA | 0.737395 | 2.04E-52 |
| CDC14A | 0.733192 | 2.43E-55 |
| CGNL1 | 0.732001 | 1.87E-59 |
| CRB2 | 0.731357 | 1.4E-54 |
| PUF60 | 0.730777 | 8.17E-59 |
| SLC9A3R1 | 0.730048 | 1.45E-67 |
| CD151 | 0.728272 | 1.6E-80 |
| CALM2 | 0.721592 | 7.03E-63 |
| TESC | 0.719868 | 3.63E-30 |
| TRAC | 0.716862 | 2.45E-47 |
| PLCE1 | 0.714454 | 2.23E-49 |
| STT3B | 0.714093 | 4.12E-43 |
| EMP3 | 0.712457 | 2.82E-51 |
| HMGN3 | 0.710743 | 5.33E-60 |
| FOXC1 | 0.70937 | 6.87E-59 |
| HOXD1 | 0.708443 | 2.81E-37 |
| NRIP2 | 0.7053 | 3.41E-45 |
| AC092813. | 0.702306 | 7.23E-46 |
| MCUB | 0.700521 | 6.6E-51 |
| SNCA | 0.685343 | 3.02E-55 |
| LOXL1 | 0.685286 | 5.84E-40 |
| B4GAT1 | 0.684726 | 1.45E-47 |
| DSC2 | 0.684429 | 2.13E-48 |
| SQOR | 0.674243 | 6.01E-53 |
| FRMD3 | 0.670362 | 2.65E-46 |
| EFNB2 | 0.670203 | 1.15E-33 |
| SYNPO | 0.668928 | 6.69E-48 |

| | | |
|---|---|---|
| BEX4 | 0.666729 | 1.73E-51 |
| MGAT4B | 0.666108 | 1.55E-53 |
| GSTA4 | 0.66444 | 8.29E-52 |
| CHI3L1 | 0.664265 | 3.16E-40 |
| GABARAP | 0.66296 | 1.76E-55 |
| GMDS | 0.661575 | 2.36E-38 |
| TXNIP | 0.659235 | 2.37E-33 |
| APLP2 | 0.6557 | 9.1E-61 |
| UBL7 | 0.65525 | 5.07E-42 |
| TRIOBP | 0.655012 | 1.85E-41 |
| AQP7 | 0.652488 | 1.1E-45 |
| APOPT1 | 0.649724 | 1.26E-46 |
| NEAT1 | 0.647802 | 2.11E-35 |
| TMEM150( | 0.647485 | 2.44E-46 |
| TMEM106I | 0.647385 | 9.17E-38 |
| AGRN | 0.640925 | 5.24E-34 |
| SPTAN1 | 0.633681 | 2.87E-40 |
| NEBL | 0.632215 | 3.45E-46 |
| IMPDH1 | 0.630137 | 8.22E-54 |
| SH3BP5 | 0.628796 | 2.73E-49 |
| SGK1 | 0.624463 | 7.04E-40 |
| MMP23B | 0.623931 | 7.62E-36 |
| FGF1 | 0.623081 | 6.61E-38 |
| RBPMS | 0.619657 | 4.18E-56 |
| HES4 | 0.617026 | 1.32E-27 |
| SYTL1 | 0.61673 | 2.99E-44 |
| NPTN | 0.611425 | 8.03E-40 |
| CD46 | 0.611139 | 8.38E-48 |
| TMEM59 | 0.602348 | 3.36E-44 |
| ARMCX3 | 0.600813 | 1.43E-42 |
| ATP6V0B | 0.599902 | 3.96E-37 |
| DYNLT3 | 0.598518 | 2.36E-36 |
| WASHC1 | 0.595621 | 5.48E-39 |
| DDX17 | 0.594993 | 2.29E-25 |
| SARAF | 0.594778 | 9.48E-39 |
| CTSL | 0.594105 | 5.83E-41 |
| TMSB4X | 0.592833 | 3.73E-75 |
| HACD2 | 0.590899 | 9.23E-37 |
| NTRK2 | 0.58977 | 3.58E-37 |
| MYH9 | 0.588974 | 1.66E-38 |
| PCMTD2 | 0.586788 | 6.97E-40 |
| PSMD6 | 0.581535 | 7.09E-38 |
| TSPAN8 | 0.579422 | 1.01E-16 |
| P4HTM | 0.577874 | 2.55E-48 |
| IL18 | 0.577161 | 3.18E-18 |
| SPATS2L | 0.574214 | 1.53E-29 |
| VWA1 | 0.573621 | 8.48E-34 |
| CTSD | 0.571658 | 4.07E-38 |
| CMTM3 | 0.569379 | 1.24E-30 |
| MVB12A | 0.568484 | 3.04E-42 |
| CRYAB | 0.567577 | 2.42E-39 |
| BMP7 | 0.56718 | 4.97E-37 |
| LAMA5 | 0.566555 | 6.18E-39 |
| ARHGEF1: | 0.565982 | 2.43E-31 |
| HLA-A | 0.565972 | 4.9E-19 |
| SQSTM1 | 0.565187 | 6.74E-35 |

| | | |
|---|---|---|
| RSRP1 | 0.563923 | 4.64E-36 |
| LAPTM4B | 0.559626 | 9.31E-33 |
| GPX3 | 0.557799 | 6.42E-17 |
| TMBIM4 | 0.555786 | 5.31E-35 |
| TRIB2 | 0.552014 | 1.77E-27 |
| ATP6AP1 | 0.550951 | 1.01E-35 |
| UBXN4 | 0.550948 | 7.49E-40 |
| LRRFIP1 | 0.547363 | 1.74E-38 |
| SLC16A1 | 0.54516 | 2.69E-16 |
| ZNF503 | 0.545122 | 1.06E-32 |
| MSI2 | 0.544429 | 7.53E-30 |
| TST | 0.541452 | 2.94E-35 |
| PRMT6 | 0.540036 | 7.18E-33 |
| GRN | 0.53926 | 1.51E-33 |
| GLUL | 0.536884 | 9.06E-34 |
| SKP1 | 0.536765 | 1.92E-37 |
| HSPA5 | 0.536609 | 7.88E-18 |
| CLTB | 0.536275 | 2.85E-38 |
| EVA1B | 0.535928 | 6.65E-29 |
| LMX1B | 0.530489 | 3.58E-37 |
| NDFIP1 | 0.529823 | 3.83E-33 |
| ALCAM | 0.529487 | 5.75E-26 |
| CORO2B | 0.527527 | 1.96E-39 |
| NMRK1 | 0.526862 | 2.92E-35 |
| NLRP1 | 0.525896 | 1.08E-24 |
| TBXAS1 | 0.525522 | 1.33E-36 |
| GPHN | 0.523886 | 3.82E-32 |
| VAPA | 0.522783 | 6.5E-33 |
| OCIAD1 | 0.52243 | 6.85E-34 |
| DSTN | 0.520913 | 1.43E-36 |
| OAZ1 | 0.520874 | 1.12E-36 |
| TULP4 | 0.51962 | 1.63E-27 |
| SELENOT | 0.519589 | 1.63E-39 |
| MGAT5 | 0.519452 | 5.4E-33 |
| ANAPC16 | 0.519345 | 6.79E-27 |
| CXADR | 0.518598 | 4.89E-35 |
| PCMT1 | 0.514562 | 6.18E-35 |
| TMCC3 | 0.511745 | 3.23E-35 |
| CITED2 | 0.509468 | 5.22E-33 |
| WASF2 | 0.509457 | 7.98E-31 |
| LTBP4 | 0.50759 | 1.95E-23 |
| YBX3 | 0.50719 | 5.47E-28 |
| FKBP2 | 0.506149 | 5.45E-31 |
| PARD3B | 0.504913 | 3.14E-36 |
| TSPAN6 | 0.504041 | 8.39E-28 |
| BGN | 0.503419 | 2.95E-19 |
| DAPL1 | 0.502071 | 1.04E-06 |
| AC009336. | 0.500924 | 1.63E-28 |
| ID4 | 0.500054 | 5.13E-20 |
| TMED4 | 0.499327 | 3.2E-31 |
| MXRA7 | 0.498357 | 6.26E-30 |
| IL1R1 | 0.497537 | 2.17E-21 |
| IGFBP2 | 0.496974 | 1.4E-32 |
| SH3BGRL | 0.495901 | 5.65E-21 |
| ATP5IF1 | 0.495168 | 5.73E-56 |
| FAM98A | 0.492702 | 8.68E-36 |

| | | |
|---|---|---|
| NECAB1 | 0.492382 | 1.74E-36 |
| ARGLU1 | 0.490849 | 6.32E-17 |
| LAMB2 | 0.490454 | 1.5E-28 |
| MAGED2 | 0.490234 | 4.76E-24 |
| NDRG2 | 0.48383 | 2.1E-21 |
| PARD6G | 0.48334 | 3.15E-38 |
| CPEB4 | 0.482716 | 8.15E-33 |
| PSAP | 0.481597 | 4.85E-30 |
| TSPAN5 | 0.481495 | 2.22E-31 |
| CAPZA2 | 0.479919 | 9.3E-30 |
| SBDS | 0.479762 | 5.45E-25 |
| CST3 | 0.479315 | 2.8E-35 |
| NCSTN | 0.478984 | 3.51E-33 |
| LAMP1 | 0.478133 | 1.62E-25 |
| ADI1 | 0.477857 | 8.82E-27 |
| PCBD1 | 0.4776 | 1.42E-40 |
| C5orf15 | 0.476889 | 4.58E-30 |
| NME3 | 0.476312 | 5.22E-25 |
| TMEM259 | 0.475429 | 6.26E-21 |
| CBLB | 0.475398 | 2.37E-19 |
| LUC7L3 | 0.475245 | 3.52E-22 |
| MYLK | 0.472164 | 1.7E-25 |
| HIST1H2A | 0.47202 | 2.82E-22 |
| TMED9 | 0.471629 | 1.86E-24 |
| CCBE1 | 0.471355 | 4.11E-36 |
| TMEM50A | 0.469425 | 3.52E-29 |
| NR2F1 | 0.46854 | 3.93E-25 |
| ID3 | 0.468132 | 2.4E-08 |
| GINM1 | 0.467811 | 8.43E-28 |
| ELOB | 0.467706 | 3.6E-20 |
| FOLR1 | 0.467241 | 5.37E-24 |
| YPEL3 | 0.466447 | 5.9E-23 |
| IFNAR1 | 0.465998 | 1.54E-34 |
| RPTOR | 0.46436 | 2.1E-23 |
| LINC00472 | 0.464008 | 3.5E-33 |
| PDIA3 | 0.463584 | 4.14E-19 |
| PRUNE2 | 0.463529 | 1.23E-31 |
| TNS2 | 0.462272 | 2.57E-23 |
| CCNDBP1 | 0.460273 | 2.47E-35 |
| TAX1BP1 | 0.460139 | 2.83E-29 |
| CDIPT | 0.459641 | 2.85E-27 |
| CFAP45 | 0.458913 | 1.4E-28 |
| PAQR7 | 0.458198 | 3.59E-19 |
| HSPA1A | 0.456326 | 1.49E-26 |
| PALLD | 0.455974 | 1.95E-18 |
| UNC5B-AS | 0.454549 | 1.98E-24 |
| DCAF8 | 0.453448 | 2.17E-28 |
| EZR | 0.452462 | 1.16E-35 |
| S100A13 | 0.452295 | 9.7E-21 |
| EMC7 | 0.450966 | 1.23E-25 |
| ADD3 | 0.44997 | 5.51E-23 |
| PTPRQ | 0.448904 | 9.49E-23 |
| GPC6 | 0.446967 | 2.77E-26 |
| SERPINB9 | 0.446188 | 4.74E-30 |
| GFRA3 | 0.44579 | 1.44E-22 |
| ITGB1 | 0.445712 | 1.62E-20 |

| | | |
|---|---|---|
| DOC2B | 0.444604 | 1.91E-23 |
| TRIM17 | 0.443649 | 1.33E-29 |
| NDUFAF3 | 0.442909 | 4.18E-24 |
| CBX6 | 0.442889 | 5.37E-22 |
| CMYA5 | 0.441353 | 5.72E-23 |
| ATP6V1G1 | 0.439648 | 1.98E-26 |
| NR2F2 | 0.43919 | 2.9E-23 |
| TMEM40 | 0.43876 | 2.36E-28 |
| FIS1 | 0.438616 | 7.59E-20 |
| PAWR | 0.437941 | 2.48E-32 |
| TMEM80 | 0.437759 | 3E-27 |
| HERPUD1 | 0.437111 | 2.47E-24 |
| FERMT2 | 0.43699 | 2.87E-27 |
| UQCR10 | 0.434668 | 8.54E-25 |
| PTPRD | 0.434607 | 1.22E-20 |
| DNAJB9 | 0.434545 | 1.27E-26 |
| RFLNB | 0.434463 | 2.81E-10 |
| RBM38 | 0.433978 | 3.26E-20 |
| FAM171A2 | 0.432508 | 9.14E-25 |
| TUBB4B | 0.431822 | 2.55E-40 |
| PLEKHB1 | 0.43177 | 3.32E-31 |
| PTBP3 | 0.431568 | 1.46E-15 |
| CXXC5 | 0.431156 | 7.77E-31 |
| DNAJA1 | 0.430141 | 1.15E-16 |
| MYL12B | 0.428898 | 9.3E-20 |
| RDX | 0.427196 | 1.21E-24 |
| NPR1 | 0.426641 | 2.28E-30 |
| MGST1 | 0.426197 | 8.95E-16 |
| B9D1 | 0.4254 | 7.07E-28 |
| ATP6V0D1 | 0.425043 | 5.88E-26 |
| TMEM179B | 0.424422 | 2.84E-23 |
| MALAT1 | 0.423835 | 0.004504 |
| TMCO3 | 0.423404 | 1.99E-24 |
| SEL1L | 0.423402 | 5.46E-20 |
| PLOD2 | 0.422659 | 1.13E-20 |
| ECHDC2 | 0.422225 | 4.8E-29 |
| FAM89A | 0.421279 | 5.43E-23 |
| ADAMTS9 | 0.420901 | 1.72E-17 |
| RAB3IL1 | 0.420689 | 1.09E-29 |
| GOLIM4 | 0.419856 | 5.55E-16 |
| C9orf3 | 0.419694 | 2.05E-20 |
| EPB41L5 | 0.418929 | 4.49E-22 |
| AC080038. | 0.418437 | 2.99E-25 |
| LRFN4 | 0.418367 | 1.8E-25 |
| AASDH | 0.417629 | 1.17E-24 |
| CFL2 | 0.416754 | 5.82E-21 |
| MTX2 | 0.415163 | 8.47E-22 |
| SGCE | 0.414915 | 1.4E-16 |
| ZNRF3 | 0.414897 | 4.22E-22 |
| FAT1 | 0.414414 | 4.79E-16 |
| ATP2C1 | 0.413886 | 1.22E-21 |
| CMBL | 0.413822 | 5.32E-14 |
| NDUFB2 | 0.413041 | 1.24E-20 |
| HOXA7 | 0.412917 | 3.59E-21 |
| SHISA2 | 0.412813 | 4.12E-18 |
| HOXA9 | 0.412723 | 2.77E-15 |

| | | |
|---|---|---|
| TCTN1 | 0.412504 | 9.57E-20 |
| KTN1 | 0.412432 | 8.28E-21 |
| PLXNB2 | 0.412021 | 8.04E-21 |
| CYHR1 | 0.410641 | 3.98E-22 |
| DAG1 | 0.410155 | 5.93E-20 |
| UBE2M | 0.410132 | 8.16E-18 |
| BAIAP2 | 0.40921 | 4.65E-18 |
| QPRT | 0.408689 | 1.19E-20 |
| BNIP3L | 0.408475 | 3.91E-15 |
| NDUFC1 | 0.407709 | 3.12E-18 |
| LINC01197 | 0.407217 | 1.61E-28 |
| ATP1A1 | 0.405481 | 4.01E-23 |
| DRD4 | 0.404706 | 4.32E-27 |
| LZTS2 | 0.404022 | 2.4E-21 |
| LRP10 | 0.403423 | 3.06E-24 |
| ARL3 | 0.403065 | 3.06E-18 |
| TGFB1 | 0.402843 | 1.03E-23 |
| SMCO4 | 0.402647 | 1.85E-23 |
| TAGLN2 | 0.402432 | 6.49E-16 |
| CARD10 | 0.402277 | 3.92E-25 |
| B3GAT3 | 0.402037 | 6.86E-22 |
| LENG8 | 0.401386 | 3.93E-16 |
| SLC25A4 | 0.400259 | 9.91E-26 |
| SON | 0.399323 | 2.24E-14 |
| SEPT2 | 0.399169 | 1.39E-18 |
| LITAF | 0.398547 | 5E-20 |
| LGALS3BP | 0.39797 | 5.7E-22 |
| PTGER4 | 0.397655 | 2.63E-25 |
| TMEM205 | 0.39722 | 5.38E-28 |
| MAZ | 0.397152 | 1.48E-17 |
| CTSZ | 0.396116 | 3.16E-25 |
| TRIM54 | 0.395122 | 3E-24 |
| SUCO | 0.395078 | 9.47E-20 |
| VCP | 0.394443 | 7.46E-15 |
| RNASEH2( | 0.39444 | 9.55E-22 |
| MARCH3 | 0.39442 | 4.95E-22 |
| FRS2 | 0.394251 | 4.33E-24 |
| COMMD7 | 0.391833 | 3.97E-20 |
| CSRP1 | 0.39176 | 4.91E-17 |
| TCF25 | 0.391686 | 1.86E-15 |
| ITFG1 | 0.391541 | 4.22E-24 |
| JTB | 0.391032 | 9.52E-18 |
| RPN2 | 0.390566 | 1.31E-17 |
| UNC50 | 0.390368 | 1.02E-21 |
| AMIGO2 | 0.390236 | 8.36E-24 |
| SERPIND1 | 0.38986 | 1.16E-26 |
| RAB7A | 0.389624 | 6.37E-17 |
| LRRN2 | 0.388543 | 3.81E-30 |
| HSP90B1 | 0.388314 | 1.08E-09 |
| SYNGR2 | 0.388224 | 2.46E-25 |
| CAPNS1 | 0.387466 | 2.16E-13 |
| SNX6 | 0.382785 | 2.47E-17 |
| PDLIM1 | 0.381255 | 7.85E-14 |
| AC245297. | 0.381078 | 6.59E-22 |
| SH3RF1 | 0.38076 | 2.13E-23 |
| CALR | 0.380515 | 2.25E-10 |

| | | |
|---|---|---|
| MBD2 | 0.380449 | 2.62E-20 |
| PTOV1 | 0.380083 | 1.63E-15 |
| PSMD8 | 0.37897 | 1.22E-14 |
| DST | 0.378919 | 6.99E-06 |
| SERINC1 | 0.378872 | 2.23E-16 |
| KIRREL2 | 0.378853 | 1.68E-27 |
| CLIP3 | 0.377704 | 2.12E-16 |
| ATP6AP2 | 0.377235 | 3.19E-20 |
| BCAR3 | 0.376236 | 9.63E-18 |
| UBB | 0.376033 | 2.75E-14 |
| PDLIM7 | 0.375382 | 5.97E-15 |
| TM7SF2 | 0.37495 | 7.44E-28 |
| LRRC41 | 0.374869 | 8.98E-18 |
| MRC2 | 0.374161 | 2.51E-15 |
| PDE6B | 0.373714 | 6.29E-19 |
| TFG | 0.373626 | 7.65E-13 |
| FBXO21 | 0.373285 | 1.51E-13 |
| COX17 | 0.37059 | 3.52E-08 |
| SEC14L1 | 0.370403 | 2.1E-16 |
| HSBP1L1 | 0.369595 | 2.45E-24 |
| KLRB1 | 0.368802 | 6.58E-20 |
| SAT1 | 0.368345 | 6.98E-23 |
| AKT2 | 0.368228 | 9.45E-21 |
| COX14 | 0.367935 | 2.08E-14 |
| NLK | 0.367405 | 1.46E-19 |
| DUSP14 | 0.367077 | 2.26E-24 |
| SPPL2A | 0.366883 | 4.42E-18 |
| SIL1 | 0.366775 | 1.49E-18 |
| GPAA1 | 0.366772 | 2.64E-19 |
| GUCY1B1 | 0.365845 | 9.66E-17 |
| C11orf71 | 0.365624 | 2.5E-21 |
| CRIM1 | 0.365293 | 2.43E-15 |
| RASSF7 | 0.365072 | 2.96E-15 |
| CA11 | 0.363994 | 1.22E-19 |
| GRHPR | 0.363741 | 7.13E-13 |
| ZBTB10 | 0.36287 | 5.64E-16 |
| BISPR | 0.362658 | 2.78E-24 |
| HSP90AA1 | 0.362261 | 9.11E-16 |
| RHEB | 0.361721 | 1.51E-13 |
| SEC62 | 0.361663 | 1.74E-13 |
| BBX | 0.360425 | 7.97E-11 |
| PC | 0.360376 | 7.99E-19 |
| KYAT1 | 0.360054 | 2.24E-26 |
| TADA3 | 0.359829 | 1.18E-17 |
| LAMP2 | 0.359505 | 1.02E-16 |
| CD200 | 0.359162 | 1.7E-21 |
| RNH1 | 0.358597 | 1.53E-13 |
| DKK3 | 0.357372 | 7.05E-25 |
| EID1 | 0.357333 | 8.47E-16 |
| TEX264 | 0.357115 | 4.09E-19 |
| CD164 | 0.356758 | 2.21E-14 |
| SLC39A10 | 0.356178 | 4.74E-13 |
| TFF3 | 0.355448 | 8.8E-14 |
| ITGA6 | 0.355336 | 1.36E-17 |
| BTG3 | 0.354587 | 9.89E-15 |
| TRAM2 | 0.35457 | 2.69E-20 |

| | | |
|---|---|---|
| PLCG2 | 0.354289 | 6.48E-07 |
| APRT | 0.353897 | 2.17E-14 |
| EAPP | 0.353266 | 7E-14 |
| TIMM8B | 0.352986 | 5.15E-13 |
| PSMC5 | 0.351978 | 6.34E-11 |
| PDLIM5 | 0.351751 | 5.09E-17 |
| ACSL3 | 0.351733 | 1.17E-13 |
| LINC0108 | 0.351696 | 5.44E-20 |
| PGRMC1 | 0.351436 | 2.67E-09 |
| PRNP | 0.351365 | 1.77E-17 |
| TMEM18 | 0.351307 | 2.59E-13 |
| AP2M1 | 0.350948 | 4.7E-09 |
| PSMD3 | 0.350797 | 4.86E-14 |
| SNRPN | 0.349232 | 7.14E-12 |
| MRPL57 | 0.348871 | 2.42E-14 |
| INF2 | 0.348823 | 2.53E-10 |
| CCPG1 | 0.348757 | 7.13E-20 |
| NDUFV3 | 0.348353 | 1.58E-14 |
| SSB | 0.347456 | 2.42E-11 |
| HNRNPH2 | 0.347357 | 2.77E-16 |
| MAPK3 | 0.34735 | 1.7E-19 |
| TNFRSF1 | 0.346627 | 5.87E-16 |
| MXD4 | 0.346544 | 1.3E-17 |
| CMTM6 | 0.345478 | 5.88E-13 |
| MLF2 | 0.344926 | 1.22E-12 |
| IFITM3 | 0.344657 | 5.46E-13 |
| TRAM1 | 0.344591 | 1.57E-12 |
| SRSF5 | 0.344202 | 5.2E-17 |
| ZDHHC12 | 0.343892 | 2.04E-15 |
| PHACTR4 | 0.34352 | 2.35E-11 |
| HAGH | 0.342971 | 5.14E-13 |
| IL13RA1 | 0.342544 | 6.01E-18 |
| SH3GLB2 | 0.342316 | 3.5E-15 |
| RTN3 | 0.342214 | 5.79E-16 |
| ANKRD12 | 0.339828 | 1.77E-08 |
| MSMO1 | 0.339716 | 2.42E-06 |
| CDKN1B | 0.33935 | 8.67E-19 |
| CTBP1 | 0.339236 | 1.22E-09 |
| ORC4 | 0.339092 | 2.11E-10 |
| SLC38A10 | 0.339034 | 1.21E-18 |
| HSPA12A | 0.339033 | 6.01E-20 |
| SLC6A6 | 0.338933 | 1.67E-13 |
| CSNK1A1 | 0.338282 | 2.96E-12 |
| C1GALT1 | 0.337603 | 1.33E-21 |
| DYNLT1 | 0.337388 | 1.67E-10 |
| ZBTB7C | 0.33667 | 3.3E-24 |
| DCTD | 0.33585 | 3.98E-14 |
| NFE2L1 | 0.335802 | 1.01E-15 |
| CCNL2 | 0.33563 | 2.35E-06 |
| ZYX | 0.335337 | 8.27E-15 |
| MYO1C | 0.334545 | 1.27E-14 |
| MYOZ1 | 0.334302 | 1.33E-15 |
| TSPYL1 | 0.333962 | 2.75E-14 |
| TMEM9B | 0.333369 | 3.4E-16 |
| CANX | 0.33227 | 1.66E-12 |
| TMBIM6 | 0.33153 | 1.75E-26 |

| | | |
|---|---|---|
| TMEM115 | 0.331332 | 4.87E-16 |
| HM13 | 0.331213 | 9.31E-09 |
| IGFBP4 | 0.331051 | 8.8E-05 |
| BSG | 0.330835 | 1.88E-27 |
| SLC16A10 | 0.329424 | 5.46E-18 |
| DCBLD2 | 0.329002 | 1.61E-16 |
| CCDC127 | 0.328779 | 5.06E-18 |
| COL4A5 | 0.327642 | 1.17E-09 |
| LRRC8A | 0.327407 | 3.78E-11 |
| MLXIP | 0.327232 | 3.22E-08 |
| ANXA7 | 0.326994 | 1.07E-16 |
| MED4 | 0.326851 | 6.3E-09 |
| AKTIP | 0.326683 | 2.03E-13 |
| PCYOX1 | 0.326678 | 1.67E-11 |
| C2orf74 | 0.326567 | 3.35E-17 |
| VDAC2 | 0.326299 | 1.19E-09 |
| ACTR1B | 0.325518 | 2.81E-11 |
| POLA2 | 0.325491 | 1.46E-18 |
| TSG101 | 0.325299 | 1.57E-13 |
| PEA15 | 0.324912 | 2.13E-16 |
| SFXN2 | 0.324858 | 5.44E-21 |
| ORMDL1 | 0.32468 | 6.1E-12 |
| MUM1 | 0.324462 | 1.34E-10 |
| NDUFA13 | 0.324357 | 1.86E-07 |
| HINT2 | 0.324328 | 1.36E-11 |
| GTF3C1 | 0.324107 | 6.07E-12 |
| SEMA4G | 0.323404 | 1.87E-14 |
| SCAND1 | 0.323015 | 7.43E-09 |
| EFEMP2 | 0.322901 | 1.12E-09 |
| NFE2L3 | 0.322161 | 1.05E-19 |
| ZSCAN16- | 0.321963 | 1.9E-12 |
| PIGP | 0.320567 | 5.8E-12 |
| DYNLRB1 | 0.320511 | 7.44E-08 |
| KRCC1 | 0.320204 | 7.5E-12 |
| LIMK2 | 0.320157 | 3.53E-17 |
| DNAJB12 | 0.320056 | 3.25E-15 |
| SMIM19 | 0.319427 | 7.83E-10 |
| FUCA2 | 0.318941 | 9.78E-19 |
| HEXA | 0.318796 | 9.62E-17 |
| LBH | 0.318641 | 1.02E-14 |
| PGAM1 | 0.318464 | 5.52E-09 |
| ALDH9A1 | 0.317726 | 1.42E-13 |
| C11orf58 | 0.317492 | 7.36E-10 |
| C9orf16 | 0.317065 | 7.44E-06 |
| PGK1 | 0.316974 | 7.09E-08 |
| EMILIN2 | 0.316944 | 5.92E-19 |
| XKR4 | 0.316785 | 1.34E-13 |
| SERINC3 | 0.316416 | 2.31E-17 |
| FNDC3B | 0.316406 | 8.64E-12 |
| RAB2A | 0.316338 | 1.06E-09 |
| RBM7 | 0.315584 | 2.63E-08 |
| PSMD1 | 0.315137 | 4.12E-13 |
| SLC37A4 | 0.315051 | 4.79E-15 |
| TEAD3 | 0.314704 | 5.66E-11 |
| CAPN2 | 0.314615 | 2.17E-10 |
| SLC7A8 | 0.313901 | 1.23E-13 |

| | | |
|---|---|---|
| LMNA | 0.313728 | 6.32E-07 |
| STON1 | 0.313696 | 5.83E-19 |
| ARPC5 | 0.313529 | 9.26E-11 |
| RAB18 | 0.31319 | 1.52E-12 |
| UNC5D | 0.313076 | 2.81E-16 |
| CTDSP1 | 0.312862 | 1.57E-05 |
| GSTT2B | 0.31282 | 1.08E-16 |
| DNAJC3 | 0.312504 | 1E-11 |
| AC137767. | 0.312264 | 1.49E-19 |
| RBM42 | 0.312196 | 2.42E-08 |
| SLC3A2 | 0.312037 | 1.11E-07 |
| ST3GAL1 | 0.311056 | 3.13E-19 |
| PCMTD1 | 0.310255 | 1.18E-11 |
| FAM50A | 0.310036 | 1.25E-05 |
| DSG2 | 0.309932 | 8.02E-19 |
| YWHAE | 0.309831 | 5.75E-09 |
| SYPL1 | 0.309518 | 4.29E-13 |
| RAB14 | 0.309338 | 1.03E-10 |
| PPP1CB | 0.308947 | 0.000185 |
| GHITM | 0.308461 | 7.51E-13 |
| PAK1 | 0.308128 | 1.11E-07 |
| PRDX5 | 0.307784 | 1.8E-06 |
| PSMC2 | 0.307765 | 3.28E-09 |
| CERS2 | 0.306829 | 2.78E-10 |
| HNRNPA2 | 0.306413 | 1.67E-12 |
| YPEL5 | 0.306394 | 9.28E-13 |
| SDHA | 0.306234 | 8.73E-09 |
| MTDH | 0.306027 | 4.35E-08 |
| ZDHHC6 | 0.30587 | 1.39E-12 |
| YIPF4 | 0.305853 | 7.36E-15 |
| TIMP1 | 0.305479 | 7.5E-15 |
| CFLAR | 0.305113 | 8E-09 |
| NDN | 0.304991 | 2.1E-09 |
| PDPN | 0.304667 | 1.41E-19 |
| SMDT1 | 0.304372 | 1.69E-09 |
| PSMB1 | 0.30374 | 5.65E-09 |
| ECM1 | 0.303318 | 3.6E-18 |
| CRELD1 | 0.303156 | 4.88E-16 |
| TMEM132/ | 0.303065 | 1.78E-07 |
| SEC11C | 0.302214 | 9.87E-15 |
| STXBP2 | 0.302209 | 8.99E-20 |
| NOL3 | 0.302068 | 2.32E-16 |
| YWHAZ | 0.301769 | 2.78E-07 |
| UQCRFS1 | 0.301057 | 2.46E-08 |
| AC008264. | 0.300561 | 3.25E-19 |
| SOD1 | 0.300311 | 1.13E-05 |
| ATOX1 | 0.300176 | 1.85E-05 |
| VSIG8 | 0.299923 | 7.06E-17 |
| TRAPPC4 | 0.299731 | 4.5E-11 |
| ASS1 | 0.299597 | 1.53E-12 |
| DDRGK1 | 0.298795 | 2.53E-12 |
| PPP1CA | 0.298642 | 7.45E-09 |
| NAA20 | 0.298423 | 1.29E-05 |
| KAZN | 0.298418 | 2.21E-12 |
| SELENOF | 0.298413 | 2.72E-07 |
| ZNF608 | 0.298235 | 9.9E-09 |

| | | |
|---|---|---|
| IL6R | 0.298116 | 4.44E-17 |
| NAPA | 0.298087 | 9.91E-11 |
| IGF2BP2 | 0.297965 | 1.99E-05 |
| COX5B | 0.297523 | 3.6E-06 |
| PDHB | 0.2972 | 5.95E-10 |
| CIB1 | 0.297053 | 2.48E-12 |
| GPR108 | 0.296656 | 1.08E-11 |
| MKRN2 | 0.296279 | 4.92E-12 |
| MYO1E | 0.295832 | 3.5E-13 |
| DMAP1 | 0.295526 | 3.85E-11 |
| PPP1R9A | 0.295492 | 2.27E-08 |
| H2AFJ | 0.295424 | 2.07E-06 |
| RAB11B | 0.295416 | 0.001478 |
| TMCO1 | 0.295301 | 3.96E-08 |
| EBAG9 | 0.294829 | 1.36E-13 |
| SDF4 | 0.294582 | 6.78E-08 |
| PNKD | 0.294259 | 5.49E-12 |
| MYCN | 0.292967 | 1.1E-21 |
| NIPSNAP3 | 0.292825 | 1.07E-08 |
| MYL6 | 0.292689 | 1.72E-19 |
| MIR4458H | 0.292629 | 1.1E-07 |
| EMC10 | 0.292561 | 4.46E-11 |
| UBXN6 | 0.290771 | 1.23E-15 |
| SMTNL2 | 0.29043 | 4.43E-07 |
| PRODH2 | 0.290118 | 2.55E-15 |
| RNF13 | 0.290037 | 9.65E-12 |
| CHD7 | 0.289762 | 1.34E-17 |
| WASL | 0.289698 | 3.7E-12 |
| ADIPOR1 | 0.289519 | 7.36E-08 |
| CDC42EP | 0.289254 | 6.53E-13 |
| RAB11FIP | 0.288986 | 2.4E-10 |
| OBSL1 | 0.28803 | 1.71E-05 |
| LRRC4C | 0.287755 | 1.02E-05 |
| GALNT10 | 0.287653 | 1.49E-12 |
| NRP2 | 0.287638 | 1.76E-06 |
| CETN3 | 0.287584 | 2.31E-11 |
| RAB6B | 0.287313 | 1.28E-18 |
| ELF2 | 0.286855 | 1.82E-10 |
| TM9SF2 | 0.286578 | 1.42E-07 |
| TMEM30A | 0.285917 | 6.41E-16 |
| BAD | 0.285878 | 7.59E-06 |
| RCN2 | 0.285714 | 2.87E-05 |
| RFC1 | 0.285629 | 9.88E-06 |
| NDUFA2 | 0.285058 | 4.86E-06 |
| NDRG1 | 0.285056 | 6.78E-10 |
| RBM39 | 0.285023 | 0.000273 |
| PRKAR1A | 0.285015 | 2.72E-06 |
| NSMCE1 | 0.285011 | 1.53E-08 |
| SPNS1 | 0.284812 | 3.64E-10 |
| PARVA | 0.284242 | 9.73E-10 |
| GAS1 | 0.284189 | 1.88E-09 |
| FBXW5 | 0.28391 | 7.24E-07 |
| BEX2 | 0.283812 | 7.56E-05 |
| ASMTL | 0.283807 | 2.53E-12 |
| APLP1 | 0.283242 | 1.22E-19 |
| NDUFA1 | 0.282522 | 2.6E-05 |

| | | |
|---|---|---|
| MYO5C | 0.282336 | 8.53E-19 |
| BAG1 | 0.282111 | 1.84E-07 |
| MAT2A | 0.281577 | 6.75E-10 |
| RTL8A | 0.281175 | 9.84E-06 |
| SNRPG | 0.281136 | 5.48E-06 |
| TSPAN4 | 0.280878 | 1.38E-07 |
| LGALS8 | 0.28084 | 1.29E-10 |
| CALCOCO | 0.280779 | 2.04E-12 |
| OSTF1 | 0.280516 | 1.01E-12 |
| UFD1 | 0.280339 | 3.01E-06 |
| DNAJB6 | 0.280205 | 2.44E-07 |
| ATXN10 | 0.279797 | 3.58E-06 |
| NRGN | 0.279572 | 4.42E-11 |
| RPL36AL | 0.279016 | 3.88E-06 |
| PARP3 | 0.278835 | 2.36E-19 |
| JKAMP | 0.27857 | 1.12E-12 |
| CTTN | 0.277578 | 1.22E-06 |
| CLDN1 | 0.277556 | 6.28E-09 |
| TBCA | 0.277049 | 0.001676 |
| NUCB1 | 0.276785 | 8.8E-10 |
| SNX17 | 0.276543 | 3.45E-10 |
| COA4 | 0.276537 | 6.03E-07 |
| YIPF6 | 0.276218 | 1.78E-09 |
| NDUFV2 | 0.2761 | 1.8E-09 |
| TSTD1 | 0.275869 | 3.4E-09 |
| RAD23A | 0.27582 | 7.81E-08 |
| SYTL4 | 0.275802 | 4.11E-20 |
| TIMM17B | 0.275376 | 8.19E-12 |
| TAPBP | 0.275341 | 1.93E-12 |
| GTF2B | 0.275184 | 1.68E-05 |
| EI24 | 0.274182 | 7.55E-06 |
| YY1 | 0.274154 | 0.000248 |
| NAGLU | 0.274124 | 1.07E-12 |
| RGN | 0.27355 | 6.99E-13 |
| PLEKHA5 | 0.273463 | 1.55E-09 |
| ABHD14B | 0.273421 | 2.16E-09 |
| RABGAP1I | 0.27339 | 1.11E-06 |
| FRAS1 | 0.27335 | 1.25E-07 |
| PBX2 | 0.272975 | 1.37E-07 |
| STX7 | 0.272878 | 2.57E-07 |
| VGLL4 | 0.272463 | 1.71E-05 |
| ARHGAP5 | 0.271922 | 2.17E-07 |
| CRIP2 | 0.271758 | 2.32E-14 |
| COPZ1 | 0.271381 | 1.96E-06 |
| PNMA1 | 0.27113 | 1.96E-06 |
| CTDSPL | 0.270848 | 1.27E-05 |
| LINC01031 | 0.270664 | 1.06E-11 |
| SPON2 | 0.27064 | 5.57E-13 |
| DLC1 | 0.270559 | 4.56E-05 |
| JUP | 0.270368 | 2.85E-10 |
| RBFOX1 | 0.270239 | 3.59E-16 |
| VPS26A | 0.269506 | 6.31E-08 |
| TTC14 | 0.269297 | 1.98E-08 |
| PTS | 0.269078 | 4.52E-07 |
| MRPS7 | 0.268741 | 3.18E-09 |
| WDR13 | 0.268561 | 2.41E-08 |

| | | |
|---|---|---|
| CAPRIN2 | 0.268012 | 2.8E-12 |
| LSR | 0.267885 | 1.27E-13 |
| EFEMP1 | 0.26778 | 1.92E-06 |
| TAF3 | 0.267276 | 5.87E-11 |
| GUSB | 0.267194 | 1.4E-10 |
| C8orf58 | 0.266993 | 8.85E-12 |
| RASL11A | 0.266764 | 1.25E-17 |
| MCC | 0.266366 | 2.87E-06 |
| NPAS3 | 0.266353 | 7.1E-17 |
| ANKRD65 | 0.266332 | 2.53E-14 |
| UQCR11 | 0.266115 | 4.96E-05 |
| ILK | 0.266083 | 4.1E-07 |
| IAH1 | 0.265285 | 0.000763 |
| GALNT2 | 0.264557 | 8.44E-10 |
| ADRM1 | 0.26436 | 6.52E-05 |
| TANC1 | 0.264307 | 1.07E-16 |
| HOXA-AS2 | 0.263876 | 6.7E-13 |
| VAMP3 | 0.26365 | 1.88E-06 |
| PSME1 | 0.263119 | 2.79E-07 |
| PAXX | 0.26283 | 8.42E-08 |
| SMAD6 | 0.262406 | 3.47E-14 |
| VPS37A | 0.262114 | 1.86E-07 |
| SKIL | 0.261677 | 2.67E-05 |
| TUSC1 | 0.261652 | 2.9E-06 |
| ARL2BP | 0.261502 | 2.29E-08 |
| PHF10 | 0.261339 | 6.24E-10 |
| ATP5F1C | 0.26129 | 9.58E-05 |
| RHBDD2 | 0.261165 | 1.75E-08 |
| HDAC1 | 0.261148 | 2.1E-09 |
| PGLS | 0.261089 | 0.003866 |
| PSMA4 | 0.261049 | 1.2E-05 |
| ATP5F1B | 0.260964 | 1.71E-05 |
| UBL5 | 0.260611 | 7.97E-05 |
| PKM | 0.2605 | 2.28E-09 |
| ENSA | 0.260471 | 0.000262 |
| PSMC4 | 0.260321 | 1.69E-08 |
| RAPGEF3 | 0.260217 | 1.82E-10 |
| PSMG4 | 0.259656 | 1.22E-11 |
| FAM3A | 0.259595 | 6.28E-08 |
| UROS | 0.259153 | 2.9E-05 |
| PPP1R16A | 0.25907 | 2.23E-11 |
| NT5C | 0.259004 | 0.000749 |
| CDK9 | 0.258734 | 1.08E-09 |
| KIAA0232 | 0.258238 | 7.91E-07 |
| PSMB6 | 0.257759 | 0.000181 |
| ZBTB20 | 0.257584 | 0.000159 |
| YIPF3 | 0.257456 | 1.13E-05 |
| NDUFC2 | 0.257208 | 0.001642 |
| BABAM2 | 0.257025 | 7.08E-10 |
| VPS28 | 0.257014 | 1.4E-05 |
| ILF3-DT | 0.256925 | 0.000281 |
| SRSF8 | 0.256553 | 1.02E-09 |
| FRY | 0.256533 | 3.8E-13 |
| RPH3AL | 0.256471 | 1.21E-12 |
| DAZAP2 | 0.25574 | 7.55E-08 |
| DNAJB2 | 0.255545 | 3.6E-05 |

| | | |
|---|---|---|
| TM2D1 | 0.255415 | 6.78E-07 |
| FSIP2 | 0.255243 | 3.61E-06 |
| ZNF524 | 0.255047 | 5.82E-14 |
| LY6E | 0.2548 | 0.00304 |
| JMJD8 | 0.254586 | 9.89E-10 |
| FDXR | 0.25441 | 4.79E-13 |
| TSPAN31 | 0.254272 | 4.41E-07 |
| YWHAB | 0.253932 | 0.000597 |
| PIGT | 0.253918 | 7.69E-06 |
| PPCS | 0.253912 | 1.56E-06 |
| ARF1 | 0.253586 | 0.001177 |
| MFGE8 | 0.253525 | 1.02E-07 |
| SDHD | 0.253149 | 4.17E-08 |
| SLC35C2 | 0.252663 | 5.02E-10 |
| BAZ2B | 0.252406 | 0.002255 |
| RTN4 | 0.252396 | 6.13E-05 |
| UACA | 0.252235 | 0.00083 |
| ITGB8 | 0.252221 | 5.12E-10 |
| SERF2 | 0.252155 | 2.79E-05 |
| GABPB1-A | 0.251891 | 0.003482 |
| TXNRD2 | 0.251661 | 1.21E-11 |
| ARHGAP2 | 0.251219 | 0.000945 |
| ACYP2 | 0.250672 | 9.11E-14 |
| PAXIP1-AS | 0.250288 | 2.65E-08 |
| RAB21 | 0.250215 | 0.000161 |
| RAD23B | 0.250054 | 1.41E-07 |

| Cluster 13 | | |
|---|---|---|
| Gene | LogFC | pVal |
| APOA1 | 1.947967 | 4.5E-118 |
| IGFBP7 | 1.744684 | 0 |
| BCAM | 1.498494 | 0 |
| MT1X | 1.480444 | 1.72E-97 |
| MT1G | 1.395739 | 6.52E-94 |
| CLEC18A | 1.383335 | 2.6E-276 |
| MT1E | 1.369777 | 1.5E-101 |
| MT2A | 1.369651 | 4.23E-86 |
| MT1H | 1.276562 | 6.02E-76 |
| SLC6A13 | 1.251572 | 1E-224 |
| SERPINA1 | 1.246958 | 1.41E-84 |
| PCSK1N | 1.218454 | 3.4E-195 |
| GPC3 | 1.171185 | 7.5E-202 |
| CLU | 1.135339 | 1.6E-238 |
| ATP5IF1 | 1.102626 | 0 |
| CD9 | 1.068898 | 9.5E-239 |
| MIR503HG | 1.065638 | 3.2E-154 |
| CLEC18B | 1.063441 | 8.8E-210 |
| MT1F | 1.062562 | 8.1E-111 |
| TSPAN12 | 1.057543 | 2E-190 |
| PAX8 | 1.051085 | 7.2E-177 |
| CLDN3 | 1.00068 | 1.5E-222 |
| CFAP126 | 1.000562 | 5.08E-90 |
| ITM2B | 0.983493 | 0 |
| LGALS2 | 0.978476 | 1.4E-170 |
| CDH6 | 0.976773 | 5.3E-143 |
| QPRT | 0.954542 | 7.7E-225 |
| AKR1C3 | 0.916122 | 3E-132 |
| EMID1 | 0.899935 | 8.7E-182 |
| EMX2 | 0.881567 | 6.9E-155 |
| ID4 | 0.87411 | 8.7E-115 |
| BSG | 0.87371 | 0 |
| CYB5A | 0.868318 | 5E-269 |
| ACAA2 | 0.868277 | 3E-203 |
| ETFB | 0.862176 | 3.6E-188 |
| SMIM24 | 0.847739 | 6.5E-212 |
| WT1 | 0.844276 | 2.1E-125 |
| CAMK2N1 | 0.835158 | 9.2E-249 |
| RIDA | 0.823777 | 1.4E-184 |
| PDZK1 | 0.818935 | 1.5E-186 |
| APOE | 0.811024 | 8.5E-151 |
| SNCA | 0.8104 | 1.8E-128 |
| C11orf54 | 0.808464 | 9.3E-150 |
| RBPMS | 0.803576 | 6.3E-208 |
| KRT18 | 0.800281 | 9E-204 |
| VAMP8 | 0.792494 | 1.6E-188 |
| GLS | 0.776952 | 8.9E-153 |
| C12orf75 | 0.775193 | 4E-172 |
| AIG1 | 0.768435 | 6.1E-207 |
| MLLT1 | 0.767862 | 8.21E-97 |
| C5orf49 | 0.753902 | 5E-99 |
| ATP1B1 | 0.753119 | 3.6E-248 |
| CEBPD | 0.748101 | 5E-135 |
| ASS1 | 0.736577 | 3.1E-132 |

| | | |
|---|---|---|
| CLEC18C | 0.730742 | 2.3E-128 |
| SLC39A4 | 0.730242 | 2.7E-101 |
| TM7SF2 | 0.728995 | 3.8E-164 |
| HPN | 0.725882 | 2.4E-190 |
| GPX3 | 0.725735 | 1.12E-65 |
| ALDH1A1 | 0.722626 | 7.4E-109 |
| CETN2 | 0.721967 | 6.3E-125 |
| EPCAM | 0.71897 | 1.4E-245 |
| GOLGA7 | 0.717014 | 3.5E-155 |
| KRT19 | 0.71273 | 1.18E-77 |
| BIN1 | 0.703105 | 2.7E-155 |
| LRRTM1 | 0.701343 | 4.8E-117 |
| KRT8 | 0.699293 | 7.1E-161 |
| S100A1 | 0.695553 | 1.68E-94 |
| SMIM1 | 0.694281 | 1.2E-103 |
| AMN | 0.693097 | 2.5E-122 |
| GATM | 0.693019 | 2.83E-75 |
| CXXC5 | 0.689639 | 4.4E-213 |
| CYBA | 0.679664 | 5.3E-175 |
| RNF181 | 0.673223 | 2.2E-181 |
| CYSTM1 | 0.66991 | 2.9E-188 |
| BHMT | 0.665446 | 1.05E-69 |
| TUBB4B | 0.66312 | 5.3E-192 |
| DNPH1 | 0.650937 | 1.2E-165 |
| HLA-DMB | 0.650667 | 4.03E-53 |
| AQP6 | 0.650098 | 3.8E-114 |
| FOXJ1 | 0.647845 | 1.12E-89 |
| FAM81B | 0.646105 | 1.98E-69 |
| MNS1 | 0.632665 | 1.67E-90 |
| ANXA4 | 0.631194 | 4.4E-196 |
| KIF9 | 0.630559 | 8.9E-110 |
| CAPSL | 0.620953 | 2.37E-76 |
| MLF1 | 0.619689 | 1.06E-91 |
| C1orf194 | 0.610312 | 1.45E-49 |
| MAPK15 | 0.610082 | 1.25E-85 |
| HEPN1 | 0.60563 | 3.3E-76 |
| SLC9A3R1 | 0.605607 | 3.8E-141 |
| KIF12 | 0.605505 | 3.2E-150 |
| PIFO | 0.604809 | 1.62E-86 |
| ARL4C | 0.600281 | 1.54E-59 |
| TMEM150/ | 0.599699 | 9E-122 |
| TMEM178/ | 0.590673 | 2.82E-74 |
| DCDC2 | 0.58434 | 6.87E-98 |
| MPC2 | 0.581873 | 1.3E-227 |
| RSPH1 | 0.577391 | 3.77E-80 |
| AKAP12 | 0.575627 | 6.26E-51 |
| NR2F1 | 0.575395 | 3.82E-94 |
| S100A16 | 0.571637 | 1E-153 |
| PGAM2 | 0.571392 | 4.15E-44 |
| SPINT2 | 0.569468 | 7.1E-221 |
| PLTP | 0.568599 | 6.8E-99 |
| C1QTNF12 | 0.568546 | 1.74E-87 |
| CCDC198 | 0.563778 | 1.3E-109 |
| S100A10 | 0.562114 | 9.3E-170 |
| AGTRAP | 0.560937 | 4.2E-105 |
| SLC51B | 0.556179 | 1.85E-64 |

| | | |
|---|---|---|
| LDHB | 0.5548 | 6.2E-151 |
| SLC39A5 | 0.554397 | 2.06E-81 |
| UGT2B7 | 0.550831 | 3.9E-121 |
| GMDS | 0.549746 | 4.11E-46 |
| PCBD1 | 0.548831 | 6.8E-195 |
| HNF1B | 0.548429 | 2.6E-96 |
| GNG11 | 0.544482 | 6.11E-80 |
| IL32 | 0.543693 | 2.97E-34 |
| PSAT1 | 0.539851 | 4.44E-76 |
| YPEL2 | 0.538837 | 3.7E-93 |
| ERI3 | 0.538643 | 3.3E-121 |
| EPS8L2 | 0.538373 | 1.5E-106 |
| ATOX1 | 0.538116 | 3.4E-139 |
| MT-ND3 | 0.534894 | 7.1E-83 |
| PPP1R16A | 0.532296 | 3.3E-103 |
| UGT2A3 | 0.529806 | 2.1E-75 |
| FAM107B | 0.529572 | 6.2E-103 |
| GK | 0.528626 | 1.89E-85 |
| TMBIM6 | 0.527824 | 4.6E-222 |
| DPY30 | 0.52751 | 4.3E-119 |
| PTTG1IP | 0.527341 | 1.8E-109 |
| IMPA2 | 0.527292 | 2.2E-119 |
| CITED2 | 0.526689 | 3.3E-138 |
| PLAT | 0.523103 | 8E-55 |
| SLC7A7 | 0.522561 | 5.77E-69 |
| MDH1 | 0.52192 | 1.5E-141 |
| RGS9 | 0.519846 | 1.06E-97 |
| BCAS3 | 0.517363 | 2.23E-82 |
| CARHSP1 | 0.516964 | 1.4E-132 |
| SLC37A4 | 0.512982 | 2.2E-117 |
| CUBN | 0.512302 | 1.11E-73 |
| CNDP2 | 0.509112 | 4.31E-98 |
| COX20 | 0.508764 | 1.9E-101 |
| DYNC1LI1 | 0.507064 | 5.43E-94 |
| MT-CO2 | 0.503712 | 2.5E-110 |
| PBLD | 0.503599 | 1.41E-85 |
| RIPPLY1 | 0.502378 | 8.83E-23 |
| EFHC1 | 0.500949 | 1.59E-72 |
| MT-ND4 | 0.498811 | 5.7E-104 |
| CRIP2 | 0.498747 | 2.69E-75 |
| PRR13 | 0.498057 | 2E-115 |
| ARHGAP2 | 0.497479 | 2.5E-75 |
| DSC2 | 0.495999 | 2.59E-73 |
| C1orf210 | 0.492728 | 4.24E-93 |
| FLRT3 | 0.490257 | 6.2E-128 |
| EMX2OS | 0.488949 | 1.64E-72 |
| BORCS7 | 0.486031 | 1.1E-116 |
| WDR54 | 0.480969 | 1.75E-75 |
| DRC1 | 0.478985 | 4.74E-60 |
| DNALI1 | 0.476456 | 1.34E-82 |
| CLDN2 | 0.474287 | 1.24E-52 |
| CD151 | 0.472734 | 4.1E-147 |
| YBX3 | 0.471516 | 6.4E-135 |
| MRPS36 | 0.469788 | 4.5E-126 |
| CRYL1 | 0.469405 | 3.4E-109 |
| MSRB1 | 0.465931 | 1.46E-83 |

| | | |
|---|---|---|
| AIFM1 | 0.464464 | 5.46E-95 |
| IER3IP1 | 0.456718 | 1.55E-92 |
| TMEM176I | 0.45611 | 1.42E-79 |
| CAV2 | 0.455713 | 1.29E-57 |
| MT-ATP6 | 0.455024 | 2.67E-70 |
| MT-ND2 | 0.454111 | 1.79E-78 |
| C1QL4 | 0.452916 | 1.97E-51 |
| CISD1 | 0.452048 | 2.8E-103 |
| DSEL | 0.451559 | 1.45E-47 |
| CIB1 | 0.450126 | 1.2E-108 |
| PRDX6 | 0.45 | 6.1E-155 |
| SUSD3 | 0.448761 | 4.11E-77 |
| HOOK1 | 0.44716 | 5.51E-88 |
| TSTD1 | 0.444945 | 3.43E-53 |
| BNIP3 | 0.444673 | 6.91E-59 |
| C9orf116 | 0.44455 | 4E-69 |
| CLIC1 | 0.441434 | 6.3E-178 |
| SYAP1 | 0.440814 | 9.72E-89 |
| RRAD | 0.440554 | 1.27E-28 |
| STARD10 | 0.438515 | 1.2E-100 |
| PARD6B | 0.43769 | 2.57E-82 |
| UGCG | 0.436985 | 2.69E-71 |
| STRADB | 0.436706 | 1.09E-90 |
| SLC5A8 | 0.436261 | 1.87E-56 |
| PHGDH | 0.435328 | 3.05E-54 |
| MT-CYB | 0.432728 | 5.28E-59 |
| PPIL6 | 0.432528 | 8.91E-66 |
| HTRA1 | 0.429485 | 5.06E-81 |
| BUD31 | 0.428778 | 4.14E-69 |
| DAB2 | 0.427523 | 2.2E-99 |
| FMO1 | 0.426926 | 2.24E-40 |
| CTSB | 0.425902 | 1.46E-86 |
| KCNJ16 | 0.424681 | 4.45E-80 |
| CDH16 | 0.42457 | 5.78E-74 |
| SLC16A10 | 0.424341 | 4.62E-67 |
| SCRN2 | 0.422184 | 1.09E-73 |
| NDUFA5 | 0.421913 | 1.8E-133 |
| AIF1L | 0.420311 | 2.94E-71 |
| ACSM3 | 0.4199 | 3.87E-75 |
| GCHFR | 0.419346 | 9.08E-54 |
| TMEM141 | 0.417447 | 6.23E-96 |
| MT1M | 0.415149 | 1.18E-09 |
| AQP7 | 0.414726 | 4.25E-71 |
| TST | 0.414017 | 3.39E-74 |
| AC022613. | 0.413426 | 1.33E-66 |
| SPATS2L | 0.413136 | 2.67E-71 |
| LRRIQ1 | 0.412692 | 8.96E-55 |
| TECR | 0.410177 | 9.82E-96 |
| ARMC3 | 0.40983 | 5.29E-56 |
| EZR | 0.408646 | 5.5E-103 |
| ERICH5 | 0.407462 | 3.37E-75 |
| PGRMC1 | 0.406697 | 3.7E-109 |
| HEPACAM | 0.406405 | 8.25E-61 |
| PLIN3 | 0.406309 | 1.18E-71 |
| PLAC1 | 0.404456 | 6.21E-64 |
| FABP1 | 0.404047 | 1E-18 |

| | | |
|---|---|---|
| PCK1 | 0.403443 | 3.52E-30 |
| ELSPBP1 | 0.403245 | 2.25E-36 |
| TMEM176/ | 0.402321 | 1.11E-70 |
| MT-CO3 | 0.400816 | 3.48E-72 |
| APRT | 0.400222 | 4.9E-111 |
| CCPG1 | 0.399977 | 7.34E-60 |
| ZNF770 | 0.398052 | 1.66E-70 |
| TCEA3 | 0.396593 | 3.31E-68 |
| MT-CO1 | 0.396409 | 1.79E-98 |
| TSPAN33 | 0.39215 | 7.6E-68 |
| TXNIP | 0.391819 | 3.49E-70 |
| C17orf97 | 0.391494 | 4.02E-61 |
| ARHGAP1 | 0.391219 | 9.64E-74 |
| MT-ND4L | 0.390807 | 1.32E-76 |
| GSTK1 | 0.390257 | 5.21E-80 |
| FTCD | 0.388731 | 9.18E-66 |
| SEPHS2 | 0.387893 | 1.01E-94 |
| MET | 0.387274 | 3.06E-62 |
| CFAP298 | 0.386457 | 1.97E-67 |
| SLC16A4 | 0.384694 | 7.54E-78 |
| ECHDC2 | 0.384622 | 5.31E-71 |
| SLC44A4 | 0.383787 | 5.03E-81 |
| TSPAN1 | 0.38331 | 4.93E-25 |
| EBP | 0.383051 | 3.06E-72 |
| STAT1 | 0.38273 | 1.21E-44 |
| DERL1 | 0.382627 | 4.12E-63 |
| ZMYND10 | 0.382367 | 2.14E-46 |
| ETHE1 | 0.382355 | 3.28E-61 |
| MAOB | 0.382147 | 4.65E-55 |
| EIF3M | 0.381199 | 1.05E-59 |
| GRB14 | 0.380871 | 2.81E-56 |
| LRPAP1 | 0.380743 | 5.4E-119 |
| UCP2 | 0.380727 | 1.39E-55 |
| ARMT1 | 0.38025 | 2.4E-81 |
| TUBG1 | 0.380074 | 1.53E-60 |
| LINC01781 | 0.378274 | 1.98E-72 |
| LYPD6B | 0.37798 | 2.02E-59 |
| RAB25 | 0.377246 | 7.15E-68 |
| CLPTM1L | 0.376075 | 2.43E-75 |
| DDC | 0.375282 | 1.83E-61 |
| ABHD11 | 0.37466 | 4.88E-75 |
| ASRGL1 | 0.374407 | 3.51E-60 |
| CAT | 0.373336 | 1.14E-63 |
| KHK | 0.372261 | 8.99E-48 |
| TMEM205 | 0.372015 | 4.58E-85 |
| CTXN1 | 0.371612 | 1.93E-58 |
| TYRO3 | 0.371498 | 2.17E-64 |
| CLTA | 0.371411 | 2.35E-90 |
| NPTN | 0.371209 | 5.14E-66 |
| SPON1 | 0.371069 | 4.61E-52 |
| ACO2 | 0.36927 | 9.77E-71 |
| MST1 | 0.368169 | 1.94E-63 |
| MSMO1 | 0.364737 | 2.13E-41 |
| TTYH1 | 0.364379 | 8.88E-36 |
| TEX9 | 0.363312 | 6.56E-61 |
| LAPTM4B | 0.362881 | 1.37E-85 |

| | | |
|---|---|---|
| COL18A1 | 0.362809 | 9.88E-74 |
| CKB | 0.362801 | 2.58E-96 |
| NR2F6 | 0.362313 | 3.3E-99 |
| CD46 | 0.36229 | 4.14E-88 |
| JUNB | 0.361378 | 1.81E-48 |
| DNAJB13 | 0.360711 | 3.18E-50 |
| CHCHD10 | 0.360693 | 1.82E-50 |
| HMGN3 | 0.360394 | 5.1E-105 |
| ECHS1 | 0.35989 | 4.56E-89 |
| NDUFB2 | 0.359004 | 1.1E-92 |
| TCTN1 | 0.358461 | 2.41E-58 |
| SAT1 | 0.357498 | 2.34E-64 |
| ARHGAP2 | 0.357339 | 5.74E-74 |
| TMEM59 | 0.357302 | 2.89E-92 |
| SSTR2 | 0.356918 | 1.37E-35 |
| ADGRG1 | 0.355341 | 1.19E-57 |
| MYL9 | 0.354875 | 5.49E-60 |
| HSPE1 | 0.354654 | 4.1E-107 |
| FXYD2 | 0.354633 | 1.4E-77 |
| C9orf24 | 0.354565 | 4.2E-21 |
| ARSE | 0.352781 | 8.4E-67 |
| PALLD | 0.352492 | 3.83E-38 |
| EPHX2 | 0.35205 | 4.61E-63 |
| HMGCS1 | 0.351756 | 1.82E-37 |
| A4GALT | 0.351509 | 3.02E-63 |
| ACMSD | 0.351418 | 2.69E-37 |
| AHCY | 0.350921 | 3.67E-66 |
| NEU1 | 0.350387 | 2.66E-75 |
| GPD1 | 0.350246 | 1.2E-53 |
| FAM183A | 0.34959 | 1.01E-35 |
| CALM3 | 0.349178 | 5.9E-104 |
| TEKT2 | 0.348244 | 5.66E-56 |
| NDUFB10 | 0.347008 | 9.3E-120 |
| ARL1 | 0.346141 | 9.08E-89 |
| P4HTM | 0.345886 | 3.72E-62 |
| METRNL | 0.344399 | 1.08E-63 |
| CCDC74A | 0.343515 | 5.49E-51 |
| CLRN3 | 0.343152 | 4.13E-64 |
| SLC3A1 | 0.342739 | 7.74E-72 |
| TBC1D1 | 0.342382 | 1.54E-51 |
| AKR7A2 | 0.340616 | 1.82E-79 |
| C1QTNF7 | 0.33846 | 9.08E-50 |
| VTN | 0.337711 | 1.82E-23 |
| NIPSNAP1 | 0.334504 | 3.71E-72 |
| CLDN1 | 0.334341 | 3.64E-52 |
| COBLL1 | 0.333663 | 7.8E-46 |
| DBI | 0.332834 | 4.9E-104 |
| HCFC1R1 | 0.332825 | 7.45E-60 |
| HSPA1A | 0.332669 | 2.49E-62 |
| SPINT1 | 0.330272 | 3.12E-59 |
| PPP1R14C | 0.330062 | 2.18E-51 |
| CIDEB | 0.326938 | 2.46E-56 |
| ITGB2 | 0.326737 | 2.14E-41 |
| CALM1 | 0.32652 | 3.5E-102 |
| PDLIM4 | 0.324608 | 3.48E-55 |
| STXBP2 | 0.324301 | 9.6E-64 |

| | | |
|---|---|---|
| ADAMTS9- | 0.324198 | 2.81E-52 |
| CTSC | 0.323802 | 2.87E-54 |
| GAS6 | 0.323431 | 1.29E-40 |
| RPS27L | 0.323055 | 7.77E-87 |
| ODC1 | 0.322557 | 8.67E-57 |
| MYL12B | 0.321834 | 3.33E-81 |
| HLA-A | 0.321452 | 3.21E-43 |
| ORAI3 | 0.32144 | 2.52E-62 |
| DYNC2LI1 | 0.320957 | 8.4E-60 |
| BICC1 | 0.320902 | 1.74E-40 |
| LPCAT3 | 0.320516 | 3.84E-59 |
| DNM1 | 0.320157 | 3.11E-54 |
| AGPAT3 | 0.319102 | 1.39E-48 |
| ATP6V1D | 0.318354 | 1.07E-72 |
| SLC25A4 | 0.318282 | 1.9E-59 |
| BMP7 | 0.317836 | 7.05E-49 |
| MPST | 0.317405 | 2.42E-62 |
| RBM47 | 0.316805 | 9.18E-59 |
| TPT1 | 0.31562 | 1.2E-145 |
| BCAP31 | 0.314748 | 1.35E-68 |
| NDUFC1 | 0.314398 | 1.2E-106 |
| DYNLT1 | 0.313749 | 2.62E-90 |
| SNX5 | 0.3137 | 8.12E-67 |
| RDH11 | 0.313211 | 2.77E-58 |
| GAMT | 0.313054 | 2.07E-43 |
| FUOM | 0.312178 | 5.51E-53 |
| CYB5R3 | 0.311544 | 5.01E-77 |
| PROS1 | 0.310599 | 2.67E-47 |
| B4GALT4 | 0.309932 | 1.97E-50 |
| TEX264 | 0.309762 | 8.03E-65 |
| ARPC3 | 0.30975 | 1.6E-111 |
| PLEKHA1 | 0.309615 | 3.12E-73 |
| CERS2 | 0.308794 | 7E-64 |
| TMED4 | 0.307165 | 3.79E-65 |
| ABO | 0.306902 | 9.16E-57 |
| ATP5MC3 | 0.306199 | 1.04E-83 |
| C19orf33 | 0.306059 | 5.05E-27 |
| SPATA17 | 0.305945 | 1.9E-46 |
| SPEF2 | 0.305841 | 1.86E-48 |
| MGAT4B | 0.305634 | 1.85E-67 |
| PLOD2 | 0.305034 | 1.18E-55 |
| SMIM19 | 0.304683 | 3.3E-63 |
| EPS8L1 | 0.304649 | 3.95E-51 |
| MRPL57 | 0.304596 | 4.9E-64 |
| RSPO3 | 0.303237 | 1.28E-29 |
| BEX2 | 0.30248 | 2.74E-55 |
| SNX4 | 0.300985 | 2.04E-57 |
| ALKBH7 | 0.300668 | 4.43E-79 |
| EPB41L5 | 0.300216 | 2.4E-43 |
| TP53TG1 | 0.2996 | 2.7E-56 |
| MRPL54 | 0.299476 | 3.55E-60 |
| MXRA7 | 0.299344 | 1.11E-55 |
| NECTIN2 | 0.299166 | 3.01E-61 |
| TMEM107 | 0.297127 | 1.88E-51 |
| SESN2 | 0.297096 | 2.7E-51 |
| CRYZ | 0.296782 | 1.02E-57 |

| | | |
|---|---|---|
| CTSZ | 0.296648 | 3E-43 |
| FLYWCH2 | 0.296056 | 2.08E-58 |
| SPA17 | 0.295892 | 2.29E-42 |
| TPM1 | 0.295589 | 7.8E-42 |
| FKBP2 | 0.295086 | 2.67E-72 |
| MORN2 | 0.295038 | 8.11E-52 |
| ANK3 | 0.294916 | 1.42E-55 |
| DDR1 | 0.294501 | 2.6E-56 |
| PPP1R1A | 0.294392 | 9.73E-61 |
| FDPS | 0.293623 | 6.12E-64 |
| ODF3B | 0.293163 | 1.84E-33 |
| GUCY1B1 | 0.292533 | 4.24E-46 |
| NRGN | 0.292364 | 9.23E-26 |
| SKAP2 | 0.292276 | 3.86E-52 |
| MISP3 | 0.291765 | 8.93E-49 |
| DEPTOR | 0.291346 | 1.38E-43 |
| PAWR | 0.291118 | 6.49E-54 |
| CLDN7 | 0.290407 | 4.97E-57 |
| OAZ1 | 0.289468 | 8.22E-97 |
| STAP2 | 0.289465 | 1.27E-45 |
| RASSF10 | 0.289344 | 8.77E-42 |
| KREMEN2 | 0.289343 | 2.51E-44 |
| EFCAB1 | 0.288855 | 9.81E-34 |
| RBM38 | 0.286717 | 2.16E-44 |
| ATRAID | 0.285953 | 3.58E-80 |
| MAPK10 | 0.2859 | 9.13E-46 |
| ATP6V1G1 | 0.285439 | 3E-113 |
| VPS28 | 0.285339 | 5.26E-71 |
| DPCD | 0.285103 | 8.29E-46 |
| F10 | 0.285005 | 1.75E-28 |
| AK4 | 0.284515 | 8.81E-45 |
| ATP6V1F | 0.284406 | 9.42E-78 |
| ESAM | 0.283207 | 3.49E-39 |
| DPP4 | 0.283075 | 9.41E-54 |
| VAV3 | 0.283072 | 4.09E-41 |
| ATP2B1 | 0.282841 | 5.97E-43 |
| USH1C | 0.282064 | 8.72E-62 |
| DNAJB9 | 0.281911 | 4.56E-49 |
| HOXB7 | 0.281837 | 6.94E-54 |
| TUBA4B | 0.281523 | 8.45E-42 |
| PAX2 | 0.281359 | 1.83E-40 |
| FTH1 | 0.281273 | 2.7E-124 |
| FAH | 0.280518 | 9.66E-51 |
| POLD2 | 0.280458 | 2.9E-53 |
| ADH5 | 0.279831 | 1.99E-84 |
| GALM | 0.278346 | 3.83E-48 |
| TRAPPC6/ | 0.278196 | 1.18E-63 |
| CD164 | 0.2778 | 9.63E-70 |
| GIPC2 | 0.277673 | 2.02E-36 |
| LGALS3BF | 0.277348 | 2.67E-42 |
| UPK1B | 0.277059 | 2.02E-37 |
| CST3 | 0.277027 | 2.78E-86 |
| C15orf65 | 0.276913 | 9.66E-44 |
| TOLLIP | 0.276704 | 1.41E-51 |
| B4GAT1 | 0.276153 | 2.06E-52 |
| PDK2 | 0.275727 | 2.16E-43 |

| | | |
|---|---|---|
| CPPED1 | 0.275508 | 2.25E-46 |
| HLA-C | 0.275397 | 2.6E-39 |
| PTGDS | 0.275116 | 0.030106 |
| CLDN10 | 0.27491 | 4.71E-47 |
| ISOC2 | 0.274238 | 5.86E-55 |
| HPCAL1 | 0.273965 | 6.5E-47 |
| AATF | 0.273639 | 5.04E-38 |
| KAZN | 0.273173 | 1.76E-40 |
| KLF13 | 0.273121 | 2E-44 |
| SRI | 0.272514 | 7.59E-55 |
| PACRG | 0.272441 | 4.41E-41 |
| HMGCL | 0.27213 | 4.91E-50 |
| NME5 | 0.27163 | 3.55E-44 |
| WIPI1 | 0.271483 | 5.59E-38 |
| XBP1 | 0.27145 | 8.22E-36 |
| TRAP1 | 0.271394 | 1.02E-49 |
| TM7SF3 | 0.271029 | 4.62E-46 |
| SPAG16 | 0.270989 | 1.48E-47 |
| TMEM256 | 0.270345 | 7.28E-53 |
| ATP5F1C | 0.270091 | 1.64E-89 |
| KLK6 | 0.26985 | 9.61E-31 |
| HSDL2 | 0.26909 | 1.12E-46 |
| MSLN | 0.268919 | 3.59E-35 |
| PSMD8 | 0.268807 | 1.15E-70 |
| CFAP36 | 0.268793 | 1.12E-53 |
| UQCR10 | 0.268288 | 1.6E-102 |
| LINC0048[illegible] | 0.268074 | 1.13E-27 |
| KIF21A | 0.267863 | 6.88E-42 |
| ACLY | 0.267833 | 6.96E-40 |
| MAF | 0.267793 | 6.06E-40 |
| LSR | 0.267757 | 9.18E-51 |
| NPR1 | 0.267756 | 8.15E-47 |
| HINT2 | 0.267704 | 1.39E-46 |
| DCXR | 0.267643 | 3.79E-39 |
| CRAT | 0.266704 | 6.59E-46 |
| PKP2 | 0.266353 | 2.49E-45 |
| FBXO17 | 0.265758 | 3.57E-43 |
| SLC39A11 | 0.265115 | 3.12E-45 |
| SMS | 0.263432 | 1.03E-57 |
| MSRB2 | 0.262574 | 3.76E-49 |
| COQ4 | 0.261686 | 4.99E-51 |
| NEK11 | 0.261427 | 7.12E-42 |
| PDXK | 0.260808 | 5.49E-48 |
| FBXW9 | 0.260665 | 7.41E-42 |
| PRR5 | 0.260049 | 1.07E-42 |
| NDUFAF3 | 0.25949 | 6.64E-48 |
| RASL12 | 0.259139 | 4.27E-41 |
| CHST13 | 0.258863 | 2.04E-37 |
| FIS1 | 0.258483 | 1.5E-53 |
| MT-ND1 | 0.258402 | 1.34E-53 |
| PKN1 | 0.257896 | 3.24E-46 |
| PMM1 | 0.257685 | 4.16E-50 |
| TMEM80 | 0.257553 | 8.96E-37 |
| NUDC | 0.257017 | 9.96E-45 |
| APOM | 0.256704 | 4.07E-27 |
| AP001528. | 0.25655 | 1.12E-42 |

| | | |
|---|---|---|
| COMMD8 | 0.256536 | 8.56E-37 |
| LAMTOR2 | 0.255875 | 4.5E-49 |
| UBAC2 | 0.25571 | 1.31E-51 |
| NUDT5 | 0.254873 | 4.65E-37 |
| ASAH1 | 0.254591 | 4.75E-54 |
| MYO6 | 0.254556 | 8.84E-49 |
| TCTEX1D2 | 0.254128 | 4.81E-44 |
| EFNB2 | 0.253842 | 8.06E-45 |
| FTL | 0.253829 | 9.13E-67 |
| CTSD | 0.253533 | 7.02E-50 |
| CCDC160 | 0.253498 | 4.13E-36 |
| NDFIP2 | 0.253174 | 9.04E-49 |
| MGST3 | 0.253091 | 1.7E-88 |
| TMED5 | 0.2525 | 2.82E-45 |
| IFT27 | 0.251997 | 2.38E-41 |
| FGFR4 | 0.251201 | 1.4E-46 |
| COTL1 | 0.251052 | 3.51E-32 |
| MYLIP | 0.250998 | 2.03E-36 |
| MGMT | 0.250176 | 1.27E-43 |

| Cluster 14 | | |
|---|---|---|
| Gene | LogFC | pVal |
| XIST | 1.561477 | 3.9E-06 |
| MTRNR2L | 1.131585 | 3.94E-46 |
| KCNQ1OT | 1.129057 | 2.55E-10 |
| PMEL | 1.051396 | 0.000141 |
| NEAT1 | 0.920675 | 5.46E-09 |
| MT-ND1 | 0.91368 | 6.11E-27 |
| DDX17 | 0.90506 | 9.4E-09 |
| NKTR | 0.901484 | 6.35E-11 |
| ARID1B | 0.877782 | 1.1E-13 |
| MT-ND4 | 0.875439 | 1.65E-35 |
| MT-ATP6 | 0.864416 | 4.45E-15 |
| MT-CO3 | 0.855271 | 2.58E-17 |
| MT-CO1 | 0.853391 | 2.81E-21 |
| MT-ND5 | 0.829587 | 1.48E-16 |
| FTX | 0.826875 | 3.79E-13 |
| MT-CO2 | 0.823969 | 8.71E-22 |
| GABPB1-A | 0.818186 | 6.63E-09 |
| MLANA | 0.80146 | 4.15E-05 |
| WSB1 | 0.786463 | 1.74E-07 |
| MT-CYB | 0.743382 | 2.39E-10 |
| DST | 0.743223 | 9.53E-08 |
| MT-ND6 | 0.73588 | 9.7E-06 |
| MEG8 | 0.722596 | 0.000419 |
| BCAN | 0.718214 | 0.000113 |
| POLR2J3.1 | 0.713225 | 6.98E-10 |
| COL27A1 | 0.713135 | 1.42E-06 |
| MT-ND2 | 0.705885 | 5.51E-26 |
| MT-ND3 | 0.700616 | 1.96E-11 |
| ANKRD36( | 0.62839 | 0.001325 |
| TIA1 | 0.626839 | 0.001942 |
| PHIP | 0.624173 | 1.98E-05 |
| SYNE2 | 0.623583 | 2.97E-05 |
| MYO9A | 0.621456 | 2.19E-09 |
| CCDC14 | 0.62019 | 5.87E-07 |
| GOLGA4 | 0.612571 | 1.38E-06 |
| MDM4 | 0.604606 | 2.61E-06 |
| RBM5 | 0.600513 | 3.04E-05 |
| MT-ND4L | 0.59992 | 3.82E-09 |
| ASH1L | 0.595117 | 6.41E-06 |
| KMT2C | 0.591827 | 4.14E-06 |
| PCNX4 | 0.587753 | 0.004383 |
| LENG8 | 0.586044 | 3.33E-05 |
| ZNF292 | 0.578912 | 8.58E-05 |
| SMG1 | 0.561207 | 0.000414 |
| KMT2A | 0.551163 | 0.001106 |
| ZNF638 | 0.544061 | 0.00601 |
| NIPBL | 0.537586 | 0.000123 |
| TNRC6A | 0.53378 | 0.013089 |
| EGR1 | 0.532629 | 0.033434 |
| MACF1 | 0.510148 | 0.010244 |
| ZMYM2 | 0.509616 | 0.002221 |
| AC245060. | 0.509141 | 0.00011 |
| PIAS2 | 0.509013 | 0.002304 |
| LPP | 0.507431 | 0.000298 |

| | | |
|---|---|---|
| CDK13 | 0.502215 | 0.001788 |
| OGA | 0.50002 | 0.01325 |
| KANSL1 | 0.491576 | 0.012312 |
| BAZ2B | 0.488859 | 0.00038 |
| SETD5 | 0.486017 | 5.68E-06 |
| AGO3 | 0.484306 | 0.011197 |
| AC092683. | 0.483472 | 0.003462 |
| ATM | 0.482251 | 0.004957 |
| SEMA6A | 0.481524 | 0.000967 |
| FLNA | 0.478691 | 0.033147 |
| ZNF37A | 0.477973 | 0.000864 |
| FZD4 | 0.47672 | 0.0046 |
| AL021368. | 0.472862 | 0.00256 |
| NSG1 | 0.470611 | 0.000136 |
| NF1 | 0.464165 | 0.00142 |
| OGT | 0.464063 | 0.035618 |
| PCM1 | 0.463878 | 0.01348 |
| LAMB1 | 0.459503 | 0.000597 |
| ANKRD36 | 0.456449 | 0.014143 |
| MON2 | 0.454847 | 0.020348 |
| CEP350 | 0.451426 | 0.034841 |
| DGKH | 0.44956 | 0.005275 |
| ERBB3 | 0.44206 | 2.9E-07 |
| CASC15 | 0.434738 | 0.022967 |
| BRAF | 0.434107 | 0.00359 |
| FSIP2 | 0.430901 | 0.011716 |
| CELF1 | 0.430757 | 0.00134 |
| SMAD2 | 0.420608 | 0.012274 |
| ZNF704 | 0.419923 | 0.001624 |
| CCDC198 | 0.413902 | 0.017852 |
| SH3PXD2/ | 0.413533 | 0.00483 |
| GIGYF1 | 0.405641 | 0.000826 |
| PTPRF | 0.40184 | 3.1E-05 |
| MARCH6 | 0.401672 | 0.002522 |
| ERBB4 | 0.400472 | 0.00965 |
| HIPK2 | 0.394498 | 0.000469 |
| AC092069. | 0.392638 | 0.000786 |
| LARP1 | 0.387716 | 0.041966 |
| MLLT6 | 0.384563 | 0.004047 |
| OBSCN | 0.381229 | 0.004351 |
| FRYL | 0.376909 | 0.032916 |
| PHC3 | 0.372579 | 0.006281 |
| JAG1 | 0.370207 | 0.021993 |
| YLPM1 | 0.362988 | 0.017469 |
| EDNRB | 0.348209 | 0.008409 |
| MAP3K2 | 0.34397 | 0.033176 |
| INSR | 0.335355 | 0.034519 |
| PWAR6 | 0.326855 | 0.005057 |
| ZNF117 | 0.322041 | 0.042479 |
| MOB1B | 0.321183 | 0.016184 |
| RREB1 | 0.317055 | 0.013854 |
| TRIM56 | 0.314207 | 0.004779 |
| PHACTR1 | 0.30829 | 0.002207 |

| **Cluster 15** | | |
|---|---|---|
| Gene | LogFC | pVal |
| MT1G | 1.469258 | 2E-220 |
| MT1H | 1.46091 | 9.3E-197 |
| MT2A | 1.405322 | 2.5E-161 |
| MT1X | 1.355577 | 4.9E-148 |
| MT1E | 1.34189 | 3E-189 |
| MT1F | 1.236528 | 2E-174 |
| FTL | 0.839618 | 5.7E-106 |
| MT-ND4 | 0.822184 | 5.24E-28 |
| MT-ND2 | 0.742201 | 1.13E-31 |
| LINC01781 | 0.731429 | 2.88E-70 |
| GPX3 | 0.708799 | 1.89E-35 |
| RHOB | 0.707592 | 4.59E-69 |
| HPN | 0.678289 | 6.35E-87 |
| S100A1 | 0.675203 | 6.35E-43 |
| MT-ND3 | 0.66537 | 1.66E-23 |
| ATP1B1 | 0.657935 | 6.66E-85 |
| PCSK1N | 0.656748 | 7.24E-75 |
| BHMT | 0.652515 | 2.33E-45 |
| MT-ND1 | 0.640161 | 3.4E-15 |
| TMEM176/ | 0.625389 | 1.83E-97 |
| MT-CO2 | 0.617332 | 1.13E-17 |
| LHX1 | 0.616577 | 2.36E-35 |
| CUBN | 0.613636 | 8.18E-43 |
| KLF6 | 0.605272 | 7.03E-31 |
| MTRNR2L | 0.600168 | 1.8E-10 |
| CDH6 | 0.599393 | 1.41E-22 |
| PDGFA | 0.596209 | 6.76E-38 |
| SMIM24 | 0.595547 | 4.5E-68 |
| DAB2 | 0.594659 | 2.39E-56 |
| ERBB3 | 0.586139 | 1.11E-34 |
| MT-CO3 | 0.585768 | 3.85E-22 |
| FMO1 | 0.585735 | 8.99E-61 |
| MT-CYB | 0.578226 | 7.14E-18 |
| CAMK2N1 | 0.564646 | 7.11E-82 |
| PPP1R1A | 0.55906 | 8.08E-52 |
| APOE | 0.544601 | 5.3E-101 |
| ASS1 | 0.541917 | 1.75E-41 |
| SDC4 | 0.534364 | 3.12E-32 |
| MPC2 | 0.531091 | 1.54E-76 |
| MT-ND4L | 0.526104 | 1.03E-28 |
| C1QTNF12 | 0.52252 | 7.14E-45 |
| TSPAN1 | 0.513587 | 2.84E-35 |
| GLYATL1 | 0.510459 | 6.48E-28 |
| MT-ATP6 | 0.501086 | 2.25E-13 |
| CYS1 | 0.501003 | 2.86E-44 |
| FLRT3 | 0.499109 | 7.16E-55 |
| FTH1 | 0.49377 | 1.38E-63 |
| SLC9A3R1 | 0.490558 | 8.17E-44 |
| LRP2 | 0.484783 | 8.54E-19 |
| CHCHD10 | 0.483202 | 8.89E-45 |
| MT-CO1 | 0.478161 | 2.32E-09 |
| SLC16A4 | 0.469812 | 2.31E-33 |
| C19orf33 | 0.467294 | 6.84E-29 |
| ACSM2A | 0.456374 | 4.06E-22 |

| | | |
|---|---|---|
| AMN | 0.45588 | 1.31E-48 |
| KIF12 | 0.447059 | 1.49E-40 |
| SERPINA1 | 0.445531 | 2.15E-22 |
| SMIM1 | 0.444132 | 1.28E-23 |
| RIDA | 0.442799 | 1.82E-37 |
| HNF4A | 0.439495 | 6.59E-20 |
| ATP6V1F | 0.438361 | 7.86E-51 |
| TMEM176I | 0.436834 | 6.29E-53 |
| PDZK1 | 0.435291 | 3.06E-45 |
| PPP1R16A | 0.431084 | 5.21E-33 |
| RAB3IP | 0.430322 | 8.35E-23 |
| PBLD | 0.429036 | 8.6E-38 |
| EMX2 | 0.421737 | 1.29E-32 |
| TMEM256 | 0.421502 | 2.3E-38 |
| ALDH1A1 | 0.42057 | 7.15E-33 |
| CCDC198 | 0.417868 | 5.69E-27 |
| B4GALT1 | 0.417807 | 9.32E-16 |
| ANK3 | 0.415201 | 3.82E-25 |
| C11orf54 | 0.41495 | 5.23E-34 |
| STRADB | 0.414131 | 1.16E-28 |
| TM7SF2 | 0.402456 | 2.08E-36 |
| CRYL1 | 0.401258 | 1.92E-37 |
| BIN1 | 0.401092 | 9.41E-36 |
| IMPA2 | 0.400297 | 4.17E-33 |
| UGT2B7 | 0.399467 | 3.23E-37 |
| PRR13 | 0.395453 | 1.48E-28 |
| LAMTOR5 | 0.395212 | 3.39E-22 |
| RGS14 | 0.394907 | 3.59E-27 |
| RBPMS | 0.391521 | 2.95E-39 |
| LGALS2 | 0.391398 | 3.01E-32 |
| GLS | 0.38732 | 1.71E-17 |
| GATM | 0.386647 | 2.42E-14 |
| MT-ND5 | 0.38632 | 2.53E-10 |
| SLC39A5 | 0.385356 | 2.56E-31 |
| KCNJ15 | 0.381885 | 1.77E-27 |
| TMEM150A | 0.380468 | 8.46E-28 |
| UGCG | 0.377636 | 3.6E-25 |
| GAL3ST1 | 0.377294 | 2.54E-28 |
| NIT2 | 0.376785 | 6.12E-24 |
| ANPEP | 0.375913 | 7.49E-14 |
| CALM3 | 0.374304 | 6.24E-31 |
| RBM47 | 0.374134 | 1.83E-16 |
| CLU | 0.373062 | 3.49E-44 |
| GCHFR | 0.372661 | 1.3E-27 |
| CYB5A | 0.371868 | 2.33E-43 |
| MSRB1 | 0.370043 | 3.34E-23 |
| RDH10 | 0.368665 | 4.97E-13 |
| SLC39A4 | 0.367944 | 1.06E-23 |
| DPP4 | 0.365102 | 3.67E-29 |
| EGR1 | 0.362844 | 3.54E-15 |
| CTSB | 0.361136 | 4.52E-25 |
| ABCC6 | 0.360491 | 1.63E-18 |
| SLC3A1 | 0.360413 | 1.09E-25 |
| CHPT1 | 0.356805 | 6.33E-22 |
| SPP1 | 0.356312 | 1.02E-20 |
| RAB29 | 0.351804 | 2.17E-21 |

| | | |
|---|---|---|
| KIF21A | 0.349448 | 3.27E-15 |
| SLC51B | 0.348956 | 2.97E-19 |
| PPP1R14C | 0.347698 | 7.85E-19 |
| HNF1B | 0.347613 | 4.93E-20 |
| SEMA5A | 0.346375 | 4.25E-09 |
| DPYS | 0.345608 | 1.53E-14 |
| SSTR2 | 0.34458 | 1.18E-17 |
| VAMP8 | 0.344061 | 8.86E-35 |
| PCBD1 | 0.343053 | 3.87E-43 |
| S100A14 | 0.34097 | 9.89E-29 |
| CTNNB1 | 0.336276 | 3.89E-10 |
| NR2F6 | 0.335502 | 4.11E-27 |
| SLC37A4 | 0.335338 | 1.43E-23 |
| EZR | 0.334501 | 1.06E-34 |
| GIPC2 | 0.334265 | 1.78E-19 |
| PPFIBP1 | 0.333075 | 4.29E-13 |
| EIF4EBP2 | 0.332737 | 1.97E-13 |
| ZFP36 | 0.332724 | 1.38E-13 |
| CDK2AP2 | 0.3325 | 3.69E-17 |
| ARID3A | 0.330488 | 1.39E-14 |
| HES4 | 0.329462 | 2.32E-20 |
| PARD6B | 0.328821 | 2.34E-17 |
| ADH6 | 0.328706 | 9.93E-17 |
| ALDH6A1 | 0.327754 | 1.81E-19 |
| GPD1 | 0.32523 | 5.9E-13 |
| MGAT4B | 0.324051 | 4.75E-17 |
| B4GALT5 | 0.323867 | 2E-13 |
| NPC2 | 0.322803 | 8.32E-33 |
| PLEKHA1 | 0.322615 | 2.8E-18 |
| KRT18 | 0.322256 | 6.42E-29 |
| LGMN | 0.321774 | 1.53E-21 |
| HDHD3 | 0.321079 | 1.06E-25 |
| TSTD1 | 0.320613 | 1.04E-16 |
| SORBS2 | 0.320397 | 1.09E-20 |
| ZBTB20 | 0.319481 | 4.52E-11 |
| ZNF704 | 0.317929 | 7.77E-16 |
| EPS8L2 | 0.316392 | 5.61E-25 |
| CEBPD | 0.316342 | 2.74E-23 |
| FGFR3 | 0.313318 | 1.43E-13 |
| CITED2 | 0.312849 | 6.1E-31 |
| DCDC2 | 0.312447 | 3.22E-16 |
| JAG1 | 0.31213 | 2.17E-09 |
| FXYD2 | 0.31085 | 1.77E-25 |
| CLEC18A | 0.31047 | 2.02E-23 |
| CDHR5 | 0.309808 | 8.75E-23 |
| STX3 | 0.30808 | 2.88E-15 |
| TSPAN12 | 0.307584 | 1.83E-14 |
| ANXA4 | 0.30483 | 6.2E-36 |
| CIDEB | 0.301811 | 2.82E-23 |
| C12orf75 | 0.300941 | 3.07E-13 |
| AIG1 | 0.299507 | 9.39E-29 |
| KRT8 | 0.299138 | 4.69E-29 |
| CD68 | 0.298995 | 1.47E-21 |
| CD151 | 0.295546 | 9.45E-34 |
| ODC1 | 0.294626 | 6.03E-27 |
| DBI | 0.294433 | 5.6E-24 |

| | | |
|---|---|---|
| MYO6 | 0.293001 | 3.38E-17 |
| AK4 | 0.2929 | 2.42E-19 |
| SCRN2 | 0.290643 | 2.72E-13 |
| TMED4 | 0.290295 | 4.44E-17 |
| GALM | 0.290122 | 2.56E-18 |
| FAM20C | 0.289793 | 2.52E-14 |
| PRELID1 | 0.289188 | 2.34E-11 |
| CYSTM1 | 0.289011 | 1.4E-25 |
| SLC44A4 | 0.288785 | 2.89E-23 |
| ACAA2 | 0.288644 | 1.93E-23 |
| MITF | 0.2883 | 6.72E-15 |
| KCNJ16 | 0.288174 | 1.94E-13 |
| MDH1 | 0.287843 | 1.07E-21 |
| DMTN | 0.287808 | 8.09E-14 |
| AGT | 0.28747 | 6.92E-16 |
| QPRT | 0.287231 | 4.32E-21 |
| EPPK1 | 0.287074 | 7.08E-12 |
| SLC25A1 | 0.285608 | 3.35E-13 |
| SLC25A5 | 0.285521 | 6.67E-15 |
| CARMIL1 | 0.280653 | 2.08E-06 |
| BRI3 | 0.280629 | 4.91E-21 |
| PHKG1 | 0.2799 | 8.67E-09 |
| VIL1 | 0.279365 | 5.02E-14 |
| MRPS36 | 0.277206 | 8.14E-23 |
| AFP | 0.277042 | 8.08E-13 |
| PTGR1 | 0.276817 | 1.44E-13 |
| DSEL | 0.275495 | 1.14E-10 |
| A4GALT | 0.275083 | 1.96E-12 |
| CXXC5 | 0.274697 | 6.47E-22 |
| IL32 | 0.274043 | 6.64E-13 |
| INSR | 0.27372 | 1.42E-10 |
| NINJ1 | 0.273603 | 2.54E-16 |
| HHLA2 | 0.273264 | 1.29E-14 |
| GPX4 | 0.273263 | 9.69E-33 |
| SLC5A8 | 0.272232 | 2.28E-09 |
| SLC2A4RC | 0.27151 | 1.95E-10 |
| TNFSF12 | 0.271015 | 3.3E-18 |
| APOA1 | 0.269946 | 1.58E-11 |
| DBNDD1 | 0.268114 | 6.93E-10 |
| CLEC18B | 0.268098 | 1.89E-20 |
| SLC40A1 | 0.268079 | 9.65E-14 |
| ERI3 | 0.268052 | 2.3E-12 |
| RAB25 | 0.266992 | 3.86E-15 |
| PANCR | 0.266758 | 9.26E-10 |
| OCIAD2 | 0.265209 | 1.3E-18 |
| SORBS3 | 0.263898 | 6.16E-11 |
| CCNG1 | 0.262609 | 3.59E-12 |
| F10 | 0.261178 | 6.51E-17 |
| NEAT1 | 0.260772 | 1.51E-13 |
| SMS | 0.259027 | 3.98E-14 |
| ENTPD5 | 0.258679 | 6.16E-10 |
| AGPAT2 | 0.258055 | 1.23E-23 |
| DHCR24 | 0.253765 | 2.5E-08 |
| MKNK2 | 0.252599 | 7.4E-10 |
| PAX8 | 0.252297 | 3.25E-18 |
| BICC1 | 0.252002 | 4.2E-05 |

| | | |
|---|---|---|
| FAM84B | 0.251744 | 2.62E-07 |
| TSPAN33 | 0.250996 | 9.27E-08 |
| JUP | 0.250561 | 1.59E-07 |

| Cluster 16 | | |
|---|---|---|
| Gene | LogFC | pVal |
| IGFBP3 | 2.762163 | 1.15E-26 |
| TMSB4X | 2.608727 | 2.85E-34 |
| CLDN5 | 2.518664 | 6.96E-28 |
| COL4A1 | 2.518148 | 2.62E-18 |
| MMP1 | 2.508396 | 5.21E-20 |
| GNG11 | 2.506681 | 8.74E-50 |
| FABP5 | 2.36543 | 1.47E-35 |
| ESM1 | 2.255194 | 3.02E-19 |
| CALM1 | 2.231064 | 2.38E-52 |
| STC1 | 2.192934 | 1.73E-17 |
| CXCR4 | 2.172834 | 9.5E-19 |
| ESAM | 2.104274 | 1.22E-25 |
| CAV1 | 2.0845 | 3.06E-29 |
| RNASE1 | 2.063641 | 2.52E-26 |
| PLVAP | 2.052671 | 6.24E-26 |
| GJA4 | 1.988922 | 2.28E-15 |
| ACKR3 | 1.963416 | 8.83E-17 |
| S100A6 | 1.915798 | 1.09E-27 |
| EGFL7 | 1.897438 | 2.12E-23 |
| CD93 | 1.877466 | 6.39E-20 |
| SOX18 | 1.877155 | 8.21E-23 |
| TM4SF18 | 1.866273 | 1.95E-23 |
| PECAM1 | 1.852122 | 2.64E-21 |
| PGF | 1.813861 | 4.08E-26 |
| ARHGDIB | 1.757135 | 7.91E-24 |
| CCDC85B | 1.753603 | 1.43E-33 |
| CHST1 | 1.735815 | 3.65E-27 |
| PRND | 1.729556 | 1.65E-12 |
| FSCN1 | 1.715905 | 6.46E-27 |
| PRSS23 | 1.709982 | 2.78E-20 |
| ECSCR | 1.708367 | 5.41E-23 |
| TP53I11 | 1.69354 | 4.89E-23 |
| TCF4 | 1.691706 | 6.72E-19 |
| PLXND1 | 1.673293 | 1.19E-22 |
| LAMA4 | 1.612747 | 6.43E-18 |
| FLT1 | 1.610245 | 1.79E-19 |
| B2M | 1.578918 | 2.84E-16 |
| COL4A2 | 1.552366 | 1.71E-11 |
| ADGRL4 | 1.53562 | 1.86E-19 |
| COL15A1 | 1.525308 | 4.44E-18 |
| MAP4K4 | 1.514961 | 8.32E-21 |
| KDR | 1.511781 | 8.47E-20 |
| EFCAB14 | 1.511656 | 1.03E-17 |
| CDH5 | 1.510455 | 1.08E-18 |
| HSPG2 | 1.493948 | 3.59E-13 |
| RAMP2 | 1.491931 | 3.96E-26 |
| APLN | 1.485534 | 4.64E-10 |
| GYPC | 1.48057 | 7.06E-20 |
| NES | 1.475834 | 2.22E-16 |
| FKBP1A | 1.43814 | 7.15E-20 |
| MCAM | 1.432149 | 2.25E-15 |
| ANGPT2 | 1.428211 | 4.61E-13 |
| ARHGAP2 | 1.420068 | 2.2E-21 |
| PFN1 | 1.419091 | 3.68E-18 |

| | | |
|---|---|---|
| ICAM2 | 1.40831 | 1.7E-16 |
| GJA1 | 1.385386 | 5.89E-15 |
| TUBB6 | 1.383719 | 4.93E-20 |
| MYL12A | 1.368416 | 1.98E-38 |
| S100A11 | 1.365586 | 7.84E-32 |
| UNC5B | 1.363672 | 7.48E-14 |
| CRIP2 | 1.354239 | 1.32E-24 |
| GSN | 1.354143 | 1.72E-21 |
| CD34 | 1.353732 | 1.34E-20 |
| ACTB | 1.345837 | 3.87E-33 |
| CAVIN3 | 1.338691 | 1.11E-19 |
| PTP4A3 | 1.304673 | 2.99E-18 |
| TNFSF10 | 1.301006 | 1.42E-15 |
| LMO2 | 1.285917 | 1.96E-17 |
| PHLDA2 | 1.283003 | 1.44E-12 |
| MAP1B | 1.270002 | 9.47E-16 |
| GNAI2 | 1.243784 | 8.99E-20 |
| ARPC1B | 1.240764 | 1.96E-18 |
| SPTBN1 | 1.237578 | 1.04E-10 |
| IGFBP2 | 1.224139 | 2.1E-25 |
| LXN | 1.223953 | 1.45E-11 |
| HOMER3 | 1.222372 | 2.51E-16 |
| LAMC1 | 1.214631 | 3.5E-10 |
| HLA-E | 1.208339 | 5.63E-16 |
| HEY1 | 1.208169 | 4.83E-15 |
| MYL6 | 1.191965 | 5.55E-29 |
| FAM107B | 1.191213 | 2.36E-20 |
| TPM4 | 1.176027 | 4.54E-24 |
| IPO11 | 1.173021 | 2.65E-19 |
| UACA | 1.167939 | 4.68E-14 |
| TUBA1A | 1.159396 | 6.04E-15 |
| CYTOR | 1.158533 | 3.74E-17 |
| SH3BP5 | 1.155349 | 2.28E-17 |
| MGST2 | 1.155258 | 2.46E-13 |
| CD99 | 1.150633 | 1.01E-18 |
| PXDN | 1.149687 | 4.57E-09 |
| SERPINB4 | 1.146318 | 0.007802 |
| AFAP1L1 | 1.140109 | 9.09E-20 |
| TMSB10 | 1.138325 | 1.24E-33 |
| ITGA5 | 1.13081 | 5.51E-12 |
| RNF19A | 1.129361 | 2.53E-12 |
| ARPC2 | 1.124318 | 6.51E-26 |
| TIE1 | 1.123348 | 3.22E-13 |
| ANXA2 | 1.121881 | 7.67E-37 |
| MMRN2 | 1.121211 | 2.3E-13 |
| RPS6KA2 | 1.118028 | 3.66E-16 |
| PEA15 | 1.115213 | 1.52E-14 |
| ADGRF5 | 1.112598 | 3.78E-15 |
| GRN | 1.110402 | 7.18E-11 |
| MYH9 | 1.108243 | 2.32E-12 |
| A2M | 1.10418 | 4.03E-10 |
| ARHGAP1 | 1.104097 | 3.82E-17 |
| RASIP1 | 1.09635 | 4.47E-17 |
| PCDH12 | 1.096261 | 1.01E-11 |
| HPGD | 1.092552 | 6.39E-13 |
| SOX17 | 1.091148 | 8.11E-13 |

| | | |
|---|---|---|
| CD40 | 1.090365 | 4.25E-16 |
| RPS27L | 1.090179 | 4.35E-11 |
| PHLDA1 | 1.082685 | 6.47E-15 |
| NRP1 | 1.080779 | 7.38E-12 |
| RASGRP3 | 1.080662 | 1.02E-13 |
| HLA-B | 1.07921 | 2.78E-06 |
| DEPP1 | 1.078662 | 5.2E-07 |
| ETS1 | 1.07492 | 2.6E-13 |
| SHANK3 | 1.0736 | 5.99E-11 |
| EMP1 | 1.069209 | 1.29E-11 |
| ECM1 | 1.0654 | 2.78E-13 |
| SELENOW | 1.059443 | 5.2E-12 |
| TLNRD1 | 1.055046 | 5.79E-12 |
| ADAM15 | 1.053727 | 5.54E-15 |
| TACC1 | 1.043378 | 5.61E-10 |
| APLP2 | 1.043124 | 2.2E-10 |
| CSTB | 1.042339 | 1.1E-17 |
| CFL1 | 1.0423 | 4.43E-20 |
| SNX3 | 1.042213 | 1.94E-09 |
| GMFG | 1.033532 | 3.41E-12 |
| THY1 | 1.02971 | 1.14E-10 |
| GPRIN3 | 1.024123 | 3.76E-10 |
| IFI16 | 1.022466 | 7.98E-14 |
| TFPI | 1.018589 | 1.26E-11 |
| ARL4C | 1.018283 | 2.81E-10 |
| HYAL2 | 1.018134 | 1.66E-06 |
| NPDC1 | 1.010701 | 5.6E-16 |
| NID1 | 1.007347 | 2.35E-09 |
| AP1S2 | 1.006571 | 3.33E-10 |
| IFITM3 | 1.004593 | 3.26E-16 |
| SEC14L1 | 1.002545 | 3.01E-16 |
| EFNB2 | 1.002372 | 1.76E-07 |
| LAMB1 | 0.99915 | 2.74E-08 |
| CAVIN1 | 0.998996 | 1.23E-15 |
| SH3BGRL3 | 0.99464 | 4.93E-15 |
| SPARC | 0.992062 | 9.91E-10 |
| FILIP1 | 0.987662 | 8.89E-13 |
| SMS | 0.985009 | 6.71E-22 |
| PNP | 0.984248 | 1.21E-14 |
| LIMS1 | 0.981946 | 1.13E-12 |
| GIMAP4 | 0.981881 | 8.26E-13 |
| NOP10 | 0.980104 | 4.75E-23 |
| CXorf36 | 0.977198 | 2.18E-14 |
| POMP | 0.976944 | 5.97E-10 |
| ELOVL5 | 0.973075 | 3.02E-11 |
| CYBA | 0.970794 | 2.97E-16 |
| ABHD17A | 0.958829 | 1.48E-11 |
| CALCRL | 0.951734 | 9.37E-13 |
| HLA-C | 0.951643 | 1.53E-08 |
| VASH1 | 0.951453 | 3.86E-10 |
| YBX3 | 0.950654 | 5.75E-10 |
| TNFAIP8L1 | 0.950205 | 2.32E-13 |
| BRI3 | 0.94544 | 1.56E-13 |
| HMOX2 | 0.938853 | 2.16E-11 |
| ITGB1 | 0.934537 | 8.02E-09 |
| SMAGP | 0.930394 | 2.42E-11 |

| | | |
|---|---|---|
| WDR1 | 0.926469 | 2.83E-13 |
| BCL6B | 0.926348 | 6.37E-13 |
| WWTR1 | 0.924555 | 4.25E-15 |
| MYO1B | 0.920063 | 2.84E-11 |
| TNFRSF4 | 0.917963 | 3.78E-07 |
| MEF2C | 0.914502 | 8.12E-11 |
| VIM | 0.913387 | 2.77E-30 |
| PLAU | 0.912632 | 9.23E-10 |
| ARPC3 | 0.910176 | 1.9E-12 |
| NOTCH4 | 0.908991 | 1.7E-07 |
| CRIM1 | 0.908203 | 7.38E-09 |
| HOPX | 0.907107 | 1.73E-13 |
| TGFB1 | 0.90518 | 1.26E-14 |
| TPM3 | 0.903595 | 6.9E-11 |
| DGKH | 0.898526 | 1.88E-07 |
| BAX | 0.896028 | 4.59E-09 |
| MIR4435-2 | 0.89564 | 2.02E-12 |
| FLNA | 0.894518 | 1.03E-06 |
| ROBO4 | 0.892119 | 2.96E-07 |
| APOD | 0.8905 | 0.015513 |
| C1orf54 | 0.882602 | 4.38E-09 |
| RGS16 | 0.882108 | 0.001462 |
| PROCR | 0.880501 | 9.4E-11 |
| PREX1 | 0.879596 | 8.13E-11 |
| PKIG | 0.874539 | 6.84E-12 |
| SMTN | 0.871051 | 1.08E-08 |
| EPAS1 | 0.870531 | 4.66E-10 |
| DOCK6 | 0.87004 | 5.39E-09 |
| TAGLN2 | 0.869091 | 5.53E-12 |
| RHOC | 0.868774 | 1.12E-13 |
| GUK1 | 0.867071 | 1.74E-11 |
| IL3RA | 0.866057 | 1.13E-09 |
| LBH | 0.86575 | 2.74E-10 |
| CSRP2 | 0.863042 | 5.77E-09 |
| TSPAN13 | 0.860617 | 6.48E-11 |
| RAB13 | 0.860434 | 0.000404 |
| C2CD4B | 0.859538 | 4.47E-10 |
| EIF4EBP1 | 0.85901 | 1.72E-12 |
| RALA | 0.858307 | 6.51E-11 |
| CLEC14A | 0.854087 | 1.13E-05 |
| ABRACL | 0.850358 | 1.61E-13 |
| CD276 | 0.848656 | 2.46E-09 |
| MYCT1 | 0.846327 | 3.75E-10 |
| GFM1 | 0.842271 | 5.82E-06 |
| ARPC5L | 0.842193 | 1.75E-12 |
| FAM69B | 0.841693 | 4.02E-10 |
| TSPAN18 | 0.834663 | 7.03E-08 |
| UBALD2 | 0.829439 | 4.71E-09 |
| PLK2 | 0.829295 | 1.57E-13 |
| TMEM255I | 0.829156 | 5.27E-14 |
| GSTO1 | 0.827308 | 1.42E-10 |
| YES1 | 0.819724 | 3.93E-09 |
| FAM43A | 0.819443 | 5.84E-10 |
| IQGAP1 | 0.81577 | 1E-07 |
| RAPGEF5 | 0.814879 | 6.55E-08 |
| CD63 | 0.813757 | 6.85E-10 |

| | | |
|---|---|---|
| SLC12A2 | 0.805632 | 2.91E-09 |
| CPNE2 | 0.803246 | 4.08E-08 |
| CAV2 | 0.803216 | 5.83E-10 |
| PRDM1 | 0.802688 | 2.22E-08 |
| THSD1 | 0.802443 | 1.14E-06 |
| TGFBR2 | 0.801454 | 7.33E-11 |
| CREM | 0.799257 | 3.56E-08 |
| GAP43 | 0.798655 | 3.04E-06 |
| SLC16A3 | 0.795974 | 2.03E-05 |
| TIMP3 | 0.795515 | 4.6E-09 |
| YWHAH | 0.792914 | 5.84E-12 |
| UBE2J1 | 0.791483 | 6.39E-11 |
| ACTR3 | 0.783966 | 3.89E-10 |
| MSN | 0.783352 | 2.19E-07 |
| OST4 | 0.782696 | 1.67E-11 |
| IL32 | 0.781953 | 3.45E-06 |
| TMEM50A | 0.781636 | 2.47E-08 |
| SOX7 | 0.781129 | 1.35E-07 |
| FDPS | 0.776433 | 7.9E-09 |
| CARD8 | 0.773731 | 8.53E-11 |
| DYSF | 0.771595 | 3.33E-08 |
| DOCK4 | 0.769742 | 2.23E-09 |
| ARPC5 | 0.76574 | 6.86E-08 |
| AP2S1 | 0.763607 | 6.57E-08 |
| BAALC | 0.763177 | 2.78E-11 |
| HLX | 0.759399 | 3.73E-08 |
| DLL4 | 0.755419 | 4.09E-10 |
| DEGS1 | 0.755312 | 6.29E-07 |
| PLS3 | 0.75463 | 5.75E-10 |
| S100A16 | 0.752624 | 2.05E-06 |
| VASP | 0.751748 | 1.67E-10 |
| CEMIP2 | 0.751054 | 1.03E-07 |
| CAP1 | 0.750935 | 1.35E-09 |
| PTPRM | 0.748432 | 3.03E-10 |
| SEMA6B | 0.747926 | 3.11E-08 |
| MAP2K1 | 0.747587 | 3.57E-05 |
| CSRP1 | 0.746556 | 1.38E-11 |
| SIPA1L2 | 0.741824 | 4.15E-06 |
| CCND1 | 0.741019 | 0.000305 |
| HLA-A | 0.740098 | 1.45E-07 |
| PPP1R18 | 0.738598 | 1.8E-11 |
| MRPL17 | 0.737399 | 7.7E-07 |
| TUBA4A | 0.735298 | 0.000145 |
| ARHGDIA | 0.734694 | 3.57E-09 |
| MYO10 | 0.734422 | 5.72E-08 |
| ROBO1 | 0.733992 | 4.95E-07 |
| SVIP | 0.732214 | 3.07E-08 |
| ZEB1 | 0.730319 | 1.14E-07 |
| POLR2L | 0.729369 | 7.28E-09 |
| CITED4 | 0.728944 | 1.01E-05 |
| LINC0148( | 0.728944 | 1.73E-07 |
| FAM89A | 0.727989 | 9.88E-07 |
| FCGRT | 0.725643 | 9.61E-10 |
| DIAPH2 | 0.720306 | 2.83E-07 |
| SH2D3C | 0.719838 | 4.12E-07 |
| REEP5 | 0.715147 | 3.86E-06 |

| | | |
|---|---|---|
| RASSF2 | 0.713112 | 2.22E-10 |
| SLC9A3R2 | 0.712135 | 4.88E-08 |
| PTPN12 | 0.712051 | 1.57E-05 |
| QPCT | 0.709629 | 2.91E-07 |
| TUBA1C | 0.709321 | 1.68E-06 |
| IGFBP4 | 0.708132 | 9.04E-08 |
| NETO2 | 0.706906 | 3.48E-08 |
| CDKN1A | 0.704976 | 1.98E-05 |
| NTAN1 | 0.70282 | 1.5E-09 |
| OSTF1 | 0.702055 | 4.61E-09 |
| MPG | 0.699577 | 2.29E-06 |
| PSMA7 | 0.698375 | 5.52E-14 |
| GALNT18 | 0.697591 | 7.81E-07 |
| CHST15 | 0.695695 | 7.58E-09 |
| ENG | 0.695555 | 1.68E-06 |
| SLC27A3 | 0.694661 | 4.32E-06 |
| EIF4G2 | 0.692884 | 1.1E-10 |
| SPRY4 | 0.692484 | 2.16E-06 |
| EHD4 | 0.690381 | 5.55E-10 |
| UBTD1 | 0.690192 | 2.77E-09 |
| TMEM219 | 0.689981 | 2.15E-06 |
| PARVB | 0.68925 | 3.54E-09 |
| ACTR2 | 0.688978 | 4.16E-09 |
| KLF6 | 0.687891 | 2.5E-11 |
| DOCK9 | 0.687848 | 1.63E-08 |
| GAPDH | 0.686975 | 6.59E-06 |
| GNG2 | 0.685509 | 1.07E-09 |
| FYN | 0.685467 | 7.55E-08 |
| ADAMTS7 | 0.679361 | 4.36E-06 |
| SH3GLB1 | 0.677122 | 5.33E-07 |
| HRAS | 0.676181 | 4.03E-07 |
| KITLG | 0.673227 | 3.68E-07 |
| ATP5F1E | 0.672875 | 7.35E-09 |
| EMCN | 0.672365 | 6.58E-09 |
| MARCKSL | 0.672058 | 5.3E-12 |
| CTSB | 0.67141 | 1.21E-06 |
| CARHSP1 | 0.669869 | 3.36E-08 |
| NOV | 0.669072 | 0.019026 |
| MAST4 | 0.667908 | 6.58E-08 |
| CBLN1 | 0.666139 | 1.79E-07 |
| IFITM2 | 0.66595 | 1.29E-07 |
| ENTPD1 | 0.665526 | 3.31E-06 |
| TMC6 | 0.66406 | 5E-08 |
| CYYR1 | 0.659453 | 6.54E-09 |
| ABL2 | 0.656577 | 3.19E-05 |
| EXOC3L1 | 0.655556 | 1.29E-10 |
| PDGFB | 0.653516 | 1.8E-09 |
| SLC38A1 | 0.650301 | 1.68E-06 |
| CLIC1 | 0.649367 | 1.17E-09 |
| RGCC | 0.648064 | 4.31E-05 |
| BCAP31 | 0.64576 | 0.000909 |
| PRCP | 0.644574 | 1.99E-06 |
| IKBIP | 0.644128 | 1.73E-09 |
| KANK3 | 0.642115 | 2.7E-06 |
| ARPC1A | 0.641024 | 1.79E-07 |
| NOTCH1 | 0.640782 | 4.25E-07 |

| | | |
|---|---|---|
| CDC37 | 0.637433 | 2.55E-08 |
| SHC1 | 0.636586 | 2.03E-07 |
| CAPN2 | 0.636383 | 1.24E-07 |
| FAM198B | 0.636166 | 5.67E-07 |
| PCDH1 | 0.634686 | 4.76E-08 |
| GAPLINC | 0.629777 | 1.33E-06 |
| MPRIP | 0.628944 | 2.29E-07 |
| SNCG | 0.628221 | 1.74E-08 |
| CDC42EP2 | 0.628198 | 1.46E-07 |
| TSPAN15 | 0.626092 | 2.38E-09 |
| N4BP3 | 0.621345 | 3.7E-07 |
| JAM3 | 0.62063 | 3.19E-07 |
| MMP14 | 0.62062 | 2.79E-07 |
| LAPTM5 | 0.620557 | 5.74E-07 |
| SLC2A3 | 0.62022 | 8.25E-07 |
| GPR4 | 0.619932 | 0.004195 |
| ITGA2 | 0.619688 | 8.62E-08 |
| AHNAK | 0.619596 | 4.77E-05 |
| KCNE3 | 0.618186 | 4.01E-06 |
| EVA1B | 0.616423 | 1.82E-08 |
| ARHGEF7 | 0.613768 | 8.88E-08 |
| YWHAB | 0.613506 | 7.46E-06 |
| FNDC3B | 0.613203 | 0.034175 |
| PODXL | 0.611264 | 2.43E-08 |
| ACVRL1 | 0.60787 | 1.52E-08 |
| RGS3 | 0.605665 | 4.46E-06 |
| MGST3 | 0.605201 | 1.9E-09 |
| TAX1BP3 | 0.602557 | 0.002411 |
| CCNY | 0.60146 | 0.000492 |
| MYL12B | 0.599289 | 8.15E-10 |
| FMNL3 | 0.598021 | 7.92E-06 |
| TMEM173 | 0.59789 | 0.000299 |
| SERF2 | 0.59778 | 0.000248 |
| KIF5B | 0.596234 | 0.000357 |
| PSAP | 0.594611 | 0.022844 |
| ELK3 | 0.594149 | 4.01E-06 |
| SCARF1 | 0.592422 | 2.53E-09 |
| PPM1F | 0.592219 | 3.31E-06 |
| SMAD1 | 0.591475 | 1.3E-06 |
| ETS2 | 0.591226 | 5.36E-05 |
| SULF2 | 0.59122 | 0.000198 |
| APP | 0.59072 | 0.000394 |
| TNFRSF10 | 0.588109 | 3.74E-05 |
| GNB1 | 0.586992 | 3.71E-07 |
| MAP4K2 | 0.586682 | 0.000144 |
| RGS5 | 0.586626 | 0.000173 |
| CLIC4 | 0.585291 | 3.59E-05 |
| MAP3K11 | 0.584586 | 2.62E-08 |
| ST3GAL6 | 0.58251 | 0.001029 |
| PTTG1IP | 0.581723 | 0.002023 |
| ARHGEF15 | 0.580082 | 0.000126 |
| ITPKB | 0.579383 | 7.39E-07 |
| DUSP4 | 0.570898 | 9.78E-09 |
| ZYX | 0.570366 | 2.04E-06 |
| GRK5 | 0.566792 | 3.48E-09 |
| EFNA1 | 0.566383 | 0.001352 |

| | | |
|---|---|---|
| RALB | 0.56467 | 7.53E-08 |
| PCAT19 | 0.564119 | 0.0089 |
| HBEGF | 0.563298 | 0.002847 |
| RRAGA | 0.563124 | 1.07E-05 |
| GNB2 | 0.562235 | 0.00016 |
| PPP1CA | 0.561346 | 6.36E-08 |
| ADAM19 | 0.561279 | 1.23E-08 |
| MACF1 | 0.559909 | 0.000337 |
| ERG | 0.559577 | 4.79E-05 |
| DPYSL2 | 0.559403 | 4.18E-05 |
| PGM2L1 | 0.559317 | 5.22E-05 |
| LAMTOR2 | 0.558541 | 3.28E-05 |
| LRP10 | 0.558348 | 0.001151 |
| KCNJ2 | 0.558331 | 0.003132 |
| PALD1 | 0.557058 | 2.83E-09 |
| ADM | 0.555268 | 0.018951 |
| OAZ2 | 0.555124 | 6.42E-06 |
| TCEAL9 | 0.554964 | 0.002799 |
| SERPINH1 | 0.554731 | 7.25E-05 |
| MYL9 | 0.554464 | 0.02191 |
| SH3PXD2/ | 0.553706 | 0.000221 |
| NOVA2 | 0.553564 | 2.6E-09 |
| MRPL33 | 0.551832 | 0.000492 |
| ANXA1 | 0.54821 | 0.005779 |
| CLEC2B | 0.547917 | 1.48E-06 |
| PRKD2 | 0.5469 | 6.23E-07 |
| IDH2 | 0.546586 | 1.44E-10 |
| ARPC4 | 0.545339 | 3.83E-06 |
| ITGA6 | 0.545154 | 5.98E-08 |
| MAPKAPK | 0.542108 | 5.13E-06 |
| KRT18 | 0.541023 | 1.44E-07 |
| ANXA5 | 0.541014 | 0.000832 |
| PPP1R14E | 0.540064 | 0.00315 |
| YWHAZ | 0.539876 | 3.33E-05 |
| NAV1 | 0.539424 | 0.000114 |
| DUSP6 | 0.539287 | 1.63E-06 |
| TUBA1B | 0.53792 | 0.00906 |
| SPTAN1 | 0.537108 | 0.000285 |
| TMEM8A | 0.535215 | 8.78E-09 |
| VPS29 | 0.534625 | 0.000588 |
| SYNJ2 | 0.531602 | 0.004398 |
| HAGLROS | 0.531141 | 8.32E-07 |
| MAP1LC3E | 0.52958 | 0.001338 |
| CTTN | 0.529425 | 0.000126 |
| RAC1 | 0.528456 | 4.59E-05 |
| AKAP13 | 0.524815 | 0.001938 |
| ARHGEF1: | 0.524249 | 7.93E-06 |
| ISG15 | 0.523965 | 0.029246 |
| NID2 | 0.523404 | 3.84E-06 |
| HECW2 | 0.522945 | 6.43E-07 |
| RSU1 | 0.521663 | 2.32E-06 |
| MPZL1 | 0.521444 | 0.002261 |
| LRRFIP1 | 0.519611 | 0.000135 |
| BCAT1 | 0.519577 | 2.05E-06 |
| F2R | 0.517403 | 0.000218 |
| NT5DC2 | 0.516458 | 0.000942 |

| | | |
|---|---|---|
| HDAC7 | 0.515699 | 0.005137 |
| CA2 | 0.515603 | 0.007353 |
| MSL3 | 0.514896 | 0.000431 |
| PLEC | 0.512449 | 6.16E-05 |
| RPP25 | 0.512014 | 0.001544 |
| CMTM3 | 0.511393 | 0.002406 |
| TRIB3 | 0.511199 | 8.98E-06 |
| ERGIC1 | 0.51085 | 0.038742 |
| TM2D2 | 0.51004 | 3.38E-05 |
| LYN | 0.510028 | 7.3E-07 |
| A1BG | 0.507032 | 3.02E-06 |
| PKN1 | 0.505653 | 0.003982 |
| SLC43A3 | 0.50546 | 2.73E-05 |
| EIF2S2 | 0.504278 | 7.77E-05 |
| SERPINB6 | 0.503751 | 0.00028 |
| PCDH17 | 0.503517 | 0.005209 |
| ECE1 | 0.501639 | 5.68E-07 |
| CD200 | 0.499732 | 3.8E-05 |
| RAB1B | 0.499142 | 0.000806 |
| VAT1 | 0.498788 | 0.000478 |
| BMP2K | 0.498546 | 0.000306 |
| RAPGEF2 | 0.49412 | 0.000138 |
| FAM129B | 0.492025 | 2.13E-06 |
| ITGA9 | 0.491068 | 2.15E-07 |
| PLEKHG1 | 0.491016 | 3.2E-08 |
| GRAMD1A | 0.49092 | 4.59E-05 |
| MMP24OS | 0.48713 | 0.012564 |
| FAM126A | 0.48639 | 2.8E-05 |
| TTYH3 | 0.486187 | 0.003467 |
| HIF1A | 0.482924 | 3E-05 |
| FBLIM1 | 0.482248 | 0.002349 |
| COMT | 0.481271 | 0.00892 |
| JCAD | 0.47803 | 4.53E-07 |
| FAM241A | 0.477888 | 8.64E-08 |
| TIMP1 | 0.477288 | 3.22E-05 |
| JAK1 | 0.476869 | 0.000152 |
| AFDN | 0.476699 | 9.47E-06 |
| TRIM47 | 0.475718 | 1.84E-06 |
| ZDHHC14 | 0.47569 | 0.002051 |
| SLC1A4 | 0.474189 | 2.14E-05 |
| TMEM204 | 0.473834 | 0.011732 |
| REEP3 | 0.473118 | 4.18E-06 |
| CLTC | 0.471905 | 6.94E-05 |
| ST8SIA4 | 0.470811 | 3.81E-06 |
| FERMT2 | 0.466986 | 0.000369 |
| CCDC50 | 0.465619 | 0.000279 |
| CCDC167 | 0.464943 | 0.000298 |
| OPTN | 0.464283 | 0.000179 |
| FAM124B | 0.463927 | 0.004265 |
| NRARP | 0.462832 | 7E-07 |
| GPRC5B | 0.461098 | 6.05E-05 |
| RIPOR1 | 0.461015 | 0.000105 |
| CISD3 | 0.460889 | 0.000197 |
| BNIP2 | 0.459239 | 0.00496 |
| TMEM189 | 0.459232 | 9.68E-05 |
| BID | 0.458962 | 0.000262 |

| | | |
|---|---|---|
| ERO1A | 0.458739 | 0.00987 |
| KBTBD11 | 0.457483 | 0.000952 |
| AES | 0.456325 | 0.005311 |
| NOS3 | 0.45523 | 6.44E-06 |
| EPOR | 0.45501 | 0.017335 |
| ABCA9-AS | 0.453401 | 0.002719 |
| SLC4A7 | 0.450346 | 0.000848 |
| ROCK2 | 0.44966 | 0.000532 |
| RAP1B | 0.449201 | 0.012887 |
| COX4I1 | 0.445522 | 3.38E-06 |
| RFLNB | 0.443097 | 0.012222 |
| UTRN | 0.442697 | 0.020048 |
| MYADM | 0.442646 | 0.00312 |
| CORO1B | 0.442319 | 0.016026 |
| RASSF3 | 0.442189 | 0.037733 |
| NTHL1 | 0.442014 | 0.003797 |
| RABAC1 | 0.440951 | 0.007153 |
| IGF2R | 0.440705 | 0.000248 |
| ARSA | 0.439338 | 0.000336 |
| RBMS1 | 0.43851 | 0.003007 |
| COPS9 | 0.4385 | 0.002377 |
| FXYD5 | 0.437821 | 0.001986 |
| CRMP1 | 0.437416 | 0.000401 |
| TWF2 | 0.436791 | 0.004708 |
| CDS2 | 0.435358 | 1.05E-05 |
| VSIR | 0.434491 | 8.06E-05 |
| TBCB | 0.432945 | 8.97E-05 |
| BOK | 0.431346 | 6.15E-05 |
| TGFB1I1 | 0.428177 | 0.001415 |
| NDUFAF3 | 0.426196 | 0.002487 |
| TFPT | 0.425618 | 0.000233 |
| CAPNS1 | 0.424828 | 0.00509 |
| CDC42BPA | 0.422249 | 0.01413 |
| CD47 | 0.42063 | 0.00255 |
| TCIM | 0.420498 | 0.005512 |
| PON2 | 0.420391 | 0.018206 |
| EIF4G1 | 0.420119 | 0.007038 |
| ZNF532 | 0.418783 | 0.0412 |
| GRB10 | 0.418207 | 0.00014 |
| RB1 | 0.417394 | 0.001105 |
| SMYD2 | 0.417175 | 0.002198 |
| SSH2 | 0.416484 | 0.012306 |
| TNFAIP2 | 0.414562 | 0.017005 |
| CD55 | 0.413188 | 0.000199 |
| ADRM1 | 0.411379 | 0.000224 |
| S100A10 | 0.410584 | 0.039071 |
| NRAS | 0.410031 | 0.009378 |
| SH2B3 | 0.409909 | 5.21E-07 |
| ST3GAL4 | 0.409748 | 0.00342 |
| TRAF4 | 0.409387 | 0.001963 |
| WSB2 | 0.408983 | 0.015672 |
| FAM219A | 0.407409 | 8.2E-06 |
| TXNDC17 | 0.407122 | 0.028764 |
| LYPLA1 | 0.406932 | 0.000991 |
| LTBP1 | 0.405966 | 0.011969 |
| SFXN3 | 0.405698 | 0.00935 |

| | | |
|---|---|---|
| FURIN | 0.404849 | 0.000497 |
| CD109 | 0.40364 | 0.000616 |
| CDC42EP3 | 0.403513 | 0.000477 |
| RAB6A | 0.403408 | 0.001507 |
| ATOX1 | 0.402965 | 9.01E-06 |
| RAI14 | 0.401711 | 0.002888 |
| OSBPL10 | 0.401414 | 0.005392 |
| CAPZB | 0.400696 | 0.026722 |
| SPCS3 | 0.400611 | 0.009598 |
| INF2 | 0.398966 | 0.008193 |
| SNN | 0.398919 | 0.000205 |
| AC087627. | 0.39877 | 0.003471 |
| PIK3R6 | 0.397295 | 0.00029 |
| FEZ1 | 0.395323 | 0.003494 |
| TJP1 | 0.394851 | 0.006015 |
| SHE | 0.394077 | 0.001364 |
| BZW1 | 0.393317 | 0.025172 |
| CHD7 | 0.392496 | 3.6E-05 |
| PRKAR2B | 0.391529 | 0.000355 |
| YWHAG | 0.391503 | 0.000696 |
| ZFAND5 | 0.390227 | 0.011104 |
| STAB1 | 0.389998 | 0.000524 |
| PLXNA2 | 0.389119 | 0.000133 |
| MRPL41 | 0.388696 | 0.030709 |
| ITPRIPL2 | 0.388225 | 0.000652 |
| ADA | 0.387194 | 0.049088 |
| C4orf48 | 0.385638 | 0.020005 |
| AGAP1 | 0.385622 | 0.003727 |
| LPAR6 | 0.384869 | 6.69E-05 |
| RNF144A | 0.384436 | 0.000457 |
| NECAP2 | 0.3809 | 0.018985 |
| DBI | 0.38066 | 0.032537 |
| RGL2 | 0.37998 | 0.000436 |
| MTSS1 | 0.378398 | 5.04E-05 |
| GDPD5 | 0.376793 | 2.43E-05 |
| MYLK | 0.376679 | 0.007443 |
| AIDA | 0.376365 | 0.000131 |
| NCKAP1 | 0.373402 | 0.00156 |
| AC007998. | 0.372927 | 0.002205 |
| PPP2R2A | 0.371953 | 0.002785 |
| GAS2L3 | 0.371834 | 0.000777 |
| PSME4 | 0.371459 | 5.23E-05 |
| GRPEL2 | 0.369975 | 0.019756 |
| SH3BP2 | 0.368502 | 0.022744 |
| ANKRD9 | 0.368451 | 8.66E-05 |
| FES | 0.368134 | 0.048803 |
| LIMK2 | 0.368116 | 0.004883 |
| LASP1 | 0.367657 | 0.031932 |
| CREG2 | 0.366426 | 7.89E-05 |
| ACOT9 | 0.365627 | 3.36E-05 |
| PSMD7 | 0.36496 | 0.037216 |
| CORO1C | 0.364348 | 0.000694 |
| RASSF1 | 0.363621 | 0.044163 |
| MCRIP1 | 0.363147 | 0.035924 |
| JAG2 | 0.361802 | 0.000677 |
| TMEM65 | 0.361755 | 0.001245 |

| | | |
|---|---|---|
| CTNNBIP1 | 0.361393 | 0.009327 |
| USP31 | 0.360888 | 0.006279 |
| SCRN1 | 0.358152 | 0.000995 |
| CDC42SE1 | 0.35775 | 0.001906 |
| LPCAT2 | 0.355092 | 0.004653 |
| PRKAR1A | 0.354488 | 0.035964 |
| CBL | 0.354377 | 0.018189 |
| KLHL5 | 0.353441 | 0.000282 |
| SELENON | 0.352949 | 0.002719 |
| TBC1D9 | 0.35281 | 0.036601 |
| MMP15 | 0.351096 | 0.000116 |
| PTK2 | 0.348539 | 0.030056 |
| RPN1 | 0.348355 | 0.023797 |
| LIPA | 0.347612 | 0.014412 |
| ATP11C | 0.346849 | 0.000179 |
| TMEM181 | 0.346004 | 0.001574 |
| TRIM16 | 0.344325 | 0.036972 |
| MAPK11 | 0.344156 | 0.000334 |
| CDK17 | 0.343791 | 0.000295 |
| STK10 | 0.343411 | 0.020538 |
| B3GNT2 | 0.343005 | 0.000401 |
| GFOD1 | 0.342886 | 0.000557 |
| PXDC1 | 0.342777 | 0.010323 |
| RHOJ | 0.342041 | 0.01782 |
| CFLAR | 0.341735 | 0.038378 |
| ICAM3 | 0.337986 | 0.000648 |
| ASAP2 | 0.335644 | 0.002632 |
| TBXA2R | 0.331582 | 0.002178 |
| ANTXR2 | 0.33071 | 0.021447 |
| PGK1 | 0.330239 | 0.004081 |
| UPP1 | 0.330014 | 0.000186 |
| RCAN2 | 0.329647 | 0.030443 |
| HHEX | 0.329635 | 0.032095 |
| S1PR1 | 0.326938 | 0.000236 |
| NLK | 0.3257 | 0.005394 |
| RAPGEF1 | 0.32486 | 0.011675 |
| C20orf204 | 0.321217 | 0.009858 |
| TMEM246 | 0.320484 | 0.008702 |
| ARHGAP2 | 0.320238 | 0.015731 |
| WIPF1 | 0.319932 | 0.033302 |
| MAPRE1 | 0.318585 | 0.019262 |
| GPSM3 | 0.311627 | 0.005394 |
| KIAA1671 | 0.309606 | 0.01084 |
| RGS19 | 0.308398 | 0.016007 |
| MLXIP | 0.303737 | 0.011834 |
| FAM167B | 0.303249 | 0.000726 |
| TSPAN9 | 0.302915 | 0.005389 |
| INKA1 | 0.301956 | 0.035253 |
| RBP1 | 0.300339 | 0.001842 |
| RBPMS | 0.297848 | 0.001645 |
| TCF15 | 0.296461 | 0.00206 |
| EPHX4 | 0.295135 | 0.007859 |
| TTC9 | 0.292883 | 0.014071 |
| CLEC1A | 0.292345 | 0.030012 |
| IL4R | 0.29216 | 0.019453 |
| DYNLT3 | 0.289652 | 0.024113 |

| | | |
|---|---|---|
| AAED1 | 0.288475 | 0.035786 |
| RAPH1 | 0.287872 | 0.000516 |
| AP1B1 | 0.287101 | 0.019078 |
| CCNYL1 | 0.286891 | 0.000501 |
| KCNN3 | 0.286518 | 0.001316 |
| MARCH2 | 0.28631 | 0.000782 |
| ACTG1 | 0.283746 | 6.49E-08 |
| BAG3 | 0.280533 | 0.021301 |
| IL16 | 0.280072 | 0.043156 |
| CSGALNA | 0.278535 | 0.001512 |
| PIEZO1 | 0.278297 | 0.000278 |
| HK1 | 0.27818 | 0.006466 |
| LRRC8C | 0.275373 | 0.046535 |
| APOLD1 | 0.27495 | 0.030279 |
| PTPRB | 0.274236 | 0.005082 |
| BAZ1A | 0.273326 | 0.022926 |
| C15orf39 | 0.268556 | 0.033171 |
| ARL15 | 0.265781 | 0.021869 |
| RGS20 | 0.262174 | 0.004703 |
| FLOT2 | 0.261961 | 0.014739 |
| TCIRG1 | 0.261623 | 0.01037 |
| SLC9A1 | 0.261045 | 0.042045 |
| ST3GAL5 | 0.260959 | 0.026377 |
| CARD8-AS | 0.257894 | 0.005979 |
| PDE8A | 0.256965 | 0.041992 |
| ARHGEF1 | 0.255681 | 0.01156 |
| GIT1 | 0.254208 | 0.029289 |
| HSPA12B | 0.253268 | 0.031869 |
| AK1 | 0.252714 | 0.013528 |
| PITPNC1 | 0.250664 | 0.018576 |

| Cluster 17 | | |
|---|---|---|
| Gene | LogFC | pVal |
| TMEM52B | 2.030704 | 4.19E-24 |
| WFDC2 | 1.998361 | 7.76E-21 |
| KRT19 | 1.754707 | 3.53E-34 |
| TACSTD2 | 1.727391 | 8.15E-18 |
| GATA3 | 1.688349 | 1.32E-23 |
| GSTM3 | 1.65472 | 4.62E-24 |
| S100A4 | 1.498251 | 3.64E-08 |
| EMX2 | 1.416704 | 6.18E-28 |
| CAMK2N1 | 1.316353 | 1.51E-27 |
| ATP1B1 | 1.29427 | 9.7E-22 |
| PAX2 | 1.24906 | 3.16E-17 |
| TMSB4X | 1.226154 | 1.41E-19 |
| SAT1 | 1.224637 | 6.86E-15 |
| GNG11 | 1.183639 | 2.79E-18 |
| MAL | 1.139753 | 1.81E-17 |
| CYB5A | 1.120978 | 8.39E-26 |
| HOXD8 | 1.106617 | 2.82E-18 |
| TFAP2A | 1.089011 | 2.09E-18 |
| ATP1A1 | 1.064783 | 8.21E-15 |
| SCPEP1 | 0.981791 | 1.1E-14 |
| KRT8 | 0.978712 | 2.17E-17 |
| CAPS | 0.968832 | 2.65E-11 |
| CLDN6 | 0.963799 | 6.66E-16 |
| ZNF503 | 0.955059 | 3.93E-18 |
| BTG1 | 0.948584 | 3.41E-13 |
| AOC1 | 0.944983 | 3.68E-10 |
| CD24 | 0.91031 | 2.23E-26 |
| MARCH11 | 0.908975 | 1.05E-10 |
| CD9 | 0.893971 | 3.66E-18 |
| S100A11 | 0.88217 | 2.43E-29 |
| IGFBP7 | 0.874424 | 4.39E-15 |
| KRT18 | 0.849758 | 2.13E-16 |
| EPCAM | 0.847662 | 4.55E-30 |
| TOX3 | 0.839313 | 2.63E-06 |
| PTBP3 | 0.835122 | 7.42E-13 |
| CLDN4 | 0.82599 | 3.19E-25 |
| ANXA11 | 0.825645 | 6.69E-14 |
| GATA3-AS | 0.82068 | 1.67E-13 |
| SCIN | 0.803467 | 2.52E-07 |
| AP1M2 | 0.788841 | 3.93E-16 |
| SH3GLB2 | 0.783273 | 1.78E-10 |
| DMKN | 0.779618 | 7.32E-16 |
| PPDPF | 0.778637 | 2.78E-20 |
| CAPG | 0.762437 | 1.77E-18 |
| ATP6V0B | 0.756425 | 1.39E-10 |
| ANKS1A | 0.74283 | 3.34E-07 |
| LHX1 | 0.736882 | 1.68E-11 |
| SLC14A2 | 0.733347 | 6.4E-08 |
| RAB25 | 0.731229 | 2.42E-17 |
| PKM | 0.728974 | 7.86E-09 |
| CLDN8 | 0.728097 | 3.04E-08 |
| VAV3 | 0.722563 | 5.48E-08 |
| COMT | 0.721353 | 2.01E-11 |
| TMEM213 | 0.715506 | 5.63E-06 |

| | | |
|---|---|---|
| PRR35 | 0.711034 | 7.93E-11 |
| TNFRSF12 | 0.710994 | 0.026729 |
| PAWR | 0.702994 | 2.72E-10 |
| POU3F3 | 0.688721 | 8.54E-12 |
| RAB11FIP | 0.687678 | 9.74E-09 |
| C4orf48 | 0.686363 | 7.99E-11 |
| MECOM | 0.686289 | 1.34E-05 |
| HOXD11 | 0.68595 | 3.47E-11 |
| NUDT4 | 0.685746 | 0.003499 |
| LINC01116 | 0.676446 | 6.32E-10 |
| IER3 | 0.670729 | 0.0013 |
| ZDHHC3 | 0.66764 | 3.99E-10 |
| PFN2 | 0.663429 | 6.77E-10 |
| GPRC5C | 0.659258 | 5.79E-12 |
| NDUFA4 | 0.655844 | 1.76E-19 |
| SDC4 | 0.654736 | 1.59E-07 |
| MTURN | 0.651739 | 8.75E-08 |
| BMP3 | 0.649801 | 1.49E-05 |
| SNX2 | 0.645567 | 5.18E-07 |
| CTSH | 0.640213 | 1.04E-05 |
| QPRT | 0.634385 | 1.13E-15 |
| MLF1 | 0.633759 | 2.01E-11 |
| EMX2OS | 0.629407 | 3.8E-08 |
| CDH1 | 0.628437 | 2.33E-05 |
| RHBG | 0.625082 | 2.46E-05 |
| COA3 | 0.622864 | 1.26E-12 |
| TM7SF2 | 0.618303 | 1.11E-08 |
| AIF1L | 0.618043 | 3.31E-08 |
| REEP5 | 0.606124 | 6.89E-14 |
| CWH43 | 0.60527 | 5.1E-07 |
| PIK3R1 | 0.604237 | 1.94E-07 |
| BLNK | 0.601217 | 9.62E-09 |
| FKBP4 | 0.600812 | 1.14E-11 |
| COBLL1 | 0.599756 | 0.004252 |
| CYCS | 0.599296 | 4.17E-06 |
| DSTN | 0.598983 | 4.78E-13 |
| OCIAD2 | 0.597964 | 6.32E-10 |
| RGL3 | 0.596205 | 6.57E-09 |
| SPINT2 | 0.595855 | 1.83E-17 |
| MAL2 | 0.595449 | 2.66E-09 |
| UCP2 | 0.594082 | 1.48E-05 |
| ATP1B3 | 0.588867 | 0.000254 |
| GIPC1 | 0.5882 | 9.18E-11 |
| TBX3 | 0.587282 | 0.000137 |
| HPRT1 | 0.580781 | 1.64E-06 |
| AP002884. | 0.579378 | 3.8E-05 |
| HDAC1 | 0.576148 | 3.14E-07 |
| IQCK | 0.575663 | 5.61E-11 |
| CSRP1 | 0.575652 | 5.41E-06 |
| HOXA9 | 0.575565 | 2.49E-05 |
| UBAC2 | 0.572123 | 7.51E-08 |
| EDNRB | 0.570748 | 0.003358 |
| GRB14 | 0.570724 | 2.93E-05 |
| TUBA4A | 0.568006 | 1.53E-06 |
| PCBD1 | 0.562745 | 2.91E-10 |
| CEBPD | 0.562245 | 3.96E-11 |

| | | |
|---|---|---|
| COL18A1 | 0.558796 | 7.43E-09 |
| ATP6AP1 | 0.557286 | 9.26E-09 |
| RALBP1 | 0.548732 | 9.01E-05 |
| PGK1 | 0.548142 | 1.77E-05 |
| HIGD1A | 0.546908 | 2.81E-05 |
| CBR1 | 0.546434 | 3.94E-05 |
| DSP | 0.545341 | 7.62E-05 |
| MUC1 | 0.544497 | 6.87E-07 |
| ATP6V0E1 | 0.543887 | 1.32E-15 |
| PNKD | 0.543661 | 2.32E-09 |
| ACADVL | 0.541164 | 6.96E-09 |
| ADGRG1 | 0.539431 | 3.88E-07 |
| BNIP3 | 0.539095 | 1.12E-05 |
| METRNL | 0.538387 | 6.42E-12 |
| CLDN3 | 0.537672 | 1.14E-10 |
| SH3YL1 | 0.535462 | 1.19E-05 |
| BCAM | 0.532442 | 1.14E-17 |
| COX5A | 0.528701 | 1.07E-11 |
| TUBB2B | 0.525415 | 2.8E-05 |
| ELF3 | 0.525368 | 1.19E-06 |
| S100A16 | 0.524856 | 2.64E-08 |
| NDUFA5 | 0.524358 | 4.91E-17 |
| TAGLN2 | 0.523528 | 0.00891 |
| TPD52 | 0.523249 | 2.81E-06 |
| PPP1CB | 0.518793 | 8.04E-05 |
| PERP | 0.518493 | 8.93E-08 |
| CDC42SE1 | 0.517826 | 0.00027 |
| PAX8 | 0.514765 | 2.57E-09 |
| MT-CO1 | 0.514708 | 3.05E-05 |
| GATA2 | 0.512641 | 0.000493 |
| PHGDH | 0.510158 | 3.33E-05 |
| STARD10 | 0.508574 | 1.37E-08 |
| EMX1 | 0.507162 | 4.33E-05 |
| GPRC5B | 0.503074 | 5.8E-05 |
| GABARAP | 0.502418 | 7.68E-05 |
| GLRX | 0.502087 | 2.42E-05 |
| CADPS2 | 0.50135 | 1.55E-07 |
| LHX1-DT | 0.501242 | 1.18E-08 |
| ITM2B | 0.500543 | 4.69E-09 |
| YWHAB | 0.500464 | 3.99E-06 |
| SNRPN | 0.497393 | 7.01E-08 |
| HOOK1 | 0.497378 | 0.0015 |
| ACADL | 0.497273 | 0.000711 |
| COX7B | 0.494277 | 6.27E-13 |
| LRPAP1 | 0.49418 | 4.52E-07 |
| STAP2 | 0.493212 | 7.54E-06 |
| ZFP36L2 | 0.492612 | 0.002634 |
| CDK2AP2 | 0.492287 | 0.016003 |
| YPEL5 | 0.489932 | 2.58E-07 |
| MOCS2 | 0.487688 | 3.92E-05 |
| SPINT1 | 0.487407 | 3.76E-07 |
| PTPRF | 0.486533 | 1.81E-05 |
| NFKBIA | 0.48369 | 0.00892 |
| CXXC5 | 0.482493 | 1.28E-07 |
| CACNB4 | 0.482284 | 0.006669 |
| NR2F6 | 0.481898 | 3.13E-08 |

| | | |
|---|---|---|
| C9orf16 | 0.481213 | 1.81E-07 |
| MT-CO2 | 0.480805 | 5.65E-08 |
| GPR160 | 0.478636 | 8.21E-05 |
| ENO1 | 0.478235 | 7.04E-06 |
| MFHAS1 | 0.478232 | 0.000182 |
| RASSF7 | 0.475916 | 9.46E-07 |
| CAAP1 | 0.475678 | 9.23E-05 |
| PRSS8 | 0.472669 | 0.008813 |
| TMEM176/ | 0.46884 | 2.41E-09 |
| AKIRIN1 | 0.467629 | 0.003359 |
| GGCT | 0.466767 | 1.7E-06 |
| TGFB1 | 0.463822 | 1.27E-05 |
| PRSS22 | 0.462573 | 0.013449 |
| RDH10 | 0.462206 | 0.013763 |
| ACSM3 | 0.461636 | 2.04E-05 |
| UQCR10 | 0.461504 | 1.91E-12 |
| IRF2BPL | 0.458063 | 0.000216 |
| SLC25A39 | 0.457458 | 7.45E-11 |
| EGFL7 | 0.457261 | 0.007315 |
| LLGL2 | 0.457181 | 6.5E-05 |
| ARPC3 | 0.456382 | 1.63E-09 |
| TSTD1 | 0.456031 | 0.000898 |
| PRKAG2 | 0.455481 | 0.000131 |
| KCNJ16 | 0.450413 | 0.000107 |
| HEBP1 | 0.449047 | 2.15E-08 |
| GPI | 0.447461 | 0.015376 |
| SLC25A4 | 0.445812 | 7.02E-06 |
| PNP | 0.445374 | 0.004439 |
| ECHS1 | 0.444406 | 1.09E-09 |
| ANXA3 | 0.443475 | 0.012467 |
| TMEM125 | 0.443125 | 7.98E-05 |
| ABRACL | 0.44308 | 1.53E-06 |
| HACD3 | 0.442044 | 9.1E-06 |
| MARVELD | 0.441856 | 0.003655 |
| ADGRF1 | 0.440537 | 9.55E-08 |
| MCUR1 | 0.440139 | 2.97E-05 |
| SPINT1-AS | 0.437622 | 3.22E-06 |
| MARCKSL | 0.436751 | 9.05E-07 |
| KRAS | 0.435878 | 2.67E-05 |
| NBDY | 0.435286 | 0.001176 |
| NDUFB9 | 0.434566 | 1.2E-07 |
| PTPN13 | 0.432449 | 0.001715 |
| AK2 | 0.431802 | 0.000383 |
| TMEM91 | 0.431086 | 8.08E-05 |
| NECTIN2 | 0.43081 | 5.22E-10 |
| MIF | 0.429419 | 0.010239 |
| MYL12B | 0.429025 | 3.88E-07 |
| LAP3 | 0.427348 | 0.000271 |
| UTRN | 0.427138 | 0.002356 |
| PLEKHA5 | 0.426268 | 0.005799 |
| HMGA1 | 0.425328 | 0.000489 |
| TMEM123 | 0.423885 | 0.000143 |
| PURPL | 0.423089 | 0.005637 |
| MRPS12 | 0.422946 | 0.000108 |
| MANF | 0.422901 | 0.000608 |
| CD55 | 0.42268 | 0.007737 |

| | | |
|---|---|---|
| MRPL33 | 0.421747 | 1.26E-06 |
| CAST | 0.420674 | 1.41E-05 |
| CLDN7 | 0.419518 | 3.57E-09 |
| JUP | 0.419435 | 4.18E-05 |
| APMAP | 0.417474 | 0.020215 |
| EYA3 | 0.41692 | 0.000184 |
| HOMER2 | 0.413165 | 3.52E-05 |
| SRSF5 | 0.412596 | 2.74E-05 |
| SMIM14 | 0.411826 | 0.029889 |
| LYPLA1 | 0.411506 | 6.58E-05 |
| DDR1 | 0.40991 | 0.003166 |
| WSB1 | 0.40936 | 0.0003 |
| TFCP2L1 | 0.409071 | 8.32E-05 |
| HOXD9 | 0.408595 | 3.95E-05 |
| ANAPC16 | 0.407075 | 1.73E-07 |
| HLA-C | 0.406809 | 0.000283 |
| SCUBE3 | 0.404225 | 2.42E-05 |
| SLC35F2 | 0.402541 | 0.000486 |
| SCAND1 | 0.402369 | 6.36E-05 |
| EZR | 0.402248 | 3.37E-05 |
| NAPA | 0.401977 | 5.46E-07 |
| DAZAP2 | 0.401011 | 7.25E-05 |
| COX6A1 | 0.400964 | 0.009341 |
| CRYZ | 0.400043 | 0.045683 |
| CD2AP | 0.398904 | 0.000657 |
| SPTLC2 | 0.398337 | 7.46E-05 |
| NFATC3 | 0.396114 | 0.000997 |
| CLINT1 | 0.394913 | 0.005359 |
| PPIC | 0.393657 | 8.51E-05 |
| KLF5 | 0.393585 | 0.000326 |
| SYNGR2 | 0.391821 | 9.16E-05 |
| S100A13 | 0.391078 | 2.33E-05 |
| SUCLG1 | 0.390964 | 0.002712 |
| NDUFA3 | 0.389168 | 1.44E-06 |
| MRPS36 | 0.388767 | 1.85E-06 |
| NDUFB5 | 0.388047 | 1.78E-06 |
| ARHGAP2 | 0.387065 | 0.005098 |
| CRIP1 | 0.386045 | 0.000218 |
| SMTNL2 | 0.385514 | 0.001265 |
| CRYL1 | 0.385482 | 0.011038 |
| MRPL57 | 0.384378 | 0.010843 |
| MT-ND4 | 0.383908 | 0.010804 |
| DDX5 | 0.382474 | 0.000709 |
| SERTAD3 | 0.382386 | 0.001716 |
| VAPA | 0.381978 | 0.001871 |
| CA12 | 0.381282 | 0.035435 |
| DCXR | 0.379698 | 0.003179 |
| CDV3 | 0.379594 | 0.003106 |
| TMEM72 | 0.378396 | 0.000429 |
| COX5B | 0.378016 | 7.27E-07 |
| YBX3 | 0.377345 | 1.79E-05 |
| CHCHD10 | 0.377169 | 1.16E-05 |
| HOXB7 | 0.376519 | 0.031119 |
| CD46 | 0.376483 | 0.006783 |
| TRAF4 | 0.376319 | 9.34E-07 |
| PSMC4 | 0.374333 | 0.001736 |

| | | |
|---|---|---|
| IVNS1ABP | 0.372378 | 0.028706 |
| ZNF503-AS | 0.37204 | 0.000329 |
| CEBPA | 0.369669 | 0.002187 |
| RABAC1 | 0.369596 | 0.003054 |
| ASAP2 | 0.369126 | 0.010431 |
| IGDCC3 | 0.368647 | 0.003611 |
| PATZ1 | 0.368181 | 0.000676 |
| PKN1 | 0.367655 | 0.003735 |
| TMEM176B | 0.367323 | 4.84E-05 |
| UQCRFS1 | 0.366412 | 0.003688 |
| HMGN3 | 0.365915 | 8.34E-05 |
| CREG1 | 0.364272 | 1.69E-05 |
| ENSA | 0.363347 | 7.11E-06 |
| ELF1 | 0.363326 | 0.022754 |
| KCTD1 | 0.363256 | 0.004175 |
| UBE2H | 0.362507 | 0.002946 |
| PABPN1 | 0.361535 | 0.000604 |
| EEF1E1 | 0.360469 | 0.001505 |
| HSPA1B | 0.358696 | 0.00016 |
| SCP2 | 0.3583 | 0.001594 |
| AK3 | 0.356867 | 0.04586 |
| PLA2G4F | 0.356797 | 0.001922 |
| PEX2 | 0.356127 | 0.001301 |
| ARPC5L | 0.355536 | 3.82E-05 |
| NOP10 | 0.354921 | 2.74E-07 |
| MRPL34 | 0.354814 | 0.000176 |
| NAT14 | 0.354723 | 0.003872 |
| LSM1 | 0.354254 | 0.001944 |
| ARSD | 0.354165 | 0.002434 |
| TMEM9B | 0.353646 | 0.006839 |
| NDUFB2 | 0.352311 | 1.66E-05 |
| MITF | 0.351992 | 0.030263 |
| MCRIP2 | 0.35134 | 0.025592 |
| MT-CO3 | 0.3505 | 0.010085 |
| PLP2 | 0.350137 | 0.002676 |
| PDHA1 | 0.3491 | 0.005679 |
| SDHAF3 | 0.348775 | 2.05E-05 |
| ORMDL2 | 0.348441 | 0.000155 |
| TUBA1C | 0.348201 | 0.018014 |
| COX17 | 0.347565 | 0.004793 |
| TPI1 | 0.345192 | 0.012022 |
| RASSF9 | 0.344855 | 0.006214 |
| AC245297. | 0.344189 | 0.016386 |
| MRPL19 | 0.343294 | 0.004559 |
| SEPHS2 | 0.342292 | 0.016137 |
| SLC38A4 | 0.341584 | 0.009749 |
| TJP3 | 0.340668 | 0.000123 |
| TRAPPC6A | 0.33834 | 0.001355 |
| MPC2 | 0.337286 | 2.43E-09 |
| GSTT2B | 0.337262 | 0.004196 |
| AHCYL1 | 0.336798 | 0.026878 |
| ARHGEF19 | 0.33603 | 0.011455 |
| TMEM106B | 0.33525 | 0.027839 |
| ERP29 | 0.335049 | 4.31E-07 |
| PEX3 | 0.334928 | 0.045087 |
| NDRG1 | 0.334866 | 0.035978 |

| | | |
|---|---|---|
| DYRK4 | 0.332682 | 0.006246 |
| POLR2L | 0.332469 | 3.62E-05 |
| ASAH1 | 0.331846 | 0.008823 |
| TMEM99 | 0.330471 | 0.0002 |
| HLA-E | 0.330456 | 0.026205 |
| RAB38 | 0.330229 | 0.024261 |
| SOX4 | 0.329802 | 0.00126 |
| PFKL | 0.329483 | 0.027593 |
| COX6B1 | 0.327752 | 1.35E-06 |
| PLSCR1 | 0.326849 | 0.049049 |
| TMED4 | 0.326628 | 0.028246 |
| LINC00623 | 0.325737 | 0.008424 |
| DNAJC19 | 0.324731 | 0.041876 |
| NDUFV2 | 0.323459 | 4.71E-08 |
| TOM1L1 | 0.323147 | 0.01072 |
| GCA | 0.322754 | 0.000145 |
| SMDT1 | 0.322503 | 0.000183 |
| HOXD3 | 0.322381 | 0.000337 |
| RTN3 | 0.320057 | 0.000715 |
| NDUFA2 | 0.319865 | 0.026743 |
| MMP24OS | 0.319736 | 0.010195 |
| EMC6 | 0.318825 | 0.000896 |
| CD151 | 0.317542 | 2.23E-08 |
| CALM1 | 0.316014 | 0.003762 |
| ZFYVE21 | 0.316003 | 0.005519 |
| TMEM179B | 0.315753 | 0.001699 |
| BZW2 | 0.315727 | 0.025729 |
| SINHCAF | 0.314991 | 0.00276 |
| SCNN1A | 0.312512 | 0.004541 |
| TMEM205 | 0.310411 | 0.006111 |
| RTN4 | 0.309479 | 1.61E-07 |
| HEBP2 | 0.309133 | 0.00018 |
| CAPN2 | 0.308224 | 0.000416 |
| NDUFA8 | 0.307708 | 0.000779 |
| PPCS | 0.307666 | 0.010913 |
| TMEM134 | 0.306949 | 0.013846 |
| PRDX1 | 0.306628 | 7.22E-05 |
| BAG1 | 0.306447 | 0.027282 |
| HS6ST1 | 0.304973 | 0.037611 |
| NAALADL2 | 0.304556 | 0.006234 |
| SLC35B4 | 0.304334 | 0.005401 |
| VAMP8 | 0.303796 | 4.72E-06 |
| SURF2 | 0.303046 | 0.007954 |
| UQCRQ | 0.302878 | 0.002921 |
| SYTL2 | 0.302312 | 0.006457 |
| NECTIN4 | 0.301951 | 0.001077 |
| MGST3 | 0.301922 | 5.97E-05 |
| C1orf131 | 0.301202 | 0.001101 |
| EIF4A2 | 0.299301 | 0.000879 |
| VAMP2 | 0.298111 | 0.003966 |
| CCNDBP1 | 0.298058 | 0.008046 |
| TMEM8A | 0.297143 | 0.029512 |
| ATP5PD | 0.2969 | 0.000261 |
| UBXN4 | 0.295245 | 0.041282 |
| APLP2 | 0.294853 | 0.009704 |
| AURKAIP1 | 0.294325 | 0.009674 |

| | | |
|---|---|---|
| LSR | 0.293896 | 0.029736 |
| COQ9 | 0.293237 | 0.03938 |
| SEC11C | 0.292822 | 0.028996 |
| ZDHHC12 | 0.29272 | 0.003866 |
| ZFPM1 | 0.292541 | 0.001093 |
| TCIM | 0.29201 | 0.039186 |
| RBX1 | 0.291973 | 0.033988 |
| ACP1 | 0.291083 | 4.99E-05 |
| ZFHX3 | 0.288308 | 0.037311 |
| MYL12A | 0.287795 | 0.011601 |
| ADH5 | 0.287013 | 0.03261 |
| MYL6 | 0.286038 | 2.53E-06 |
| EFCAB14 | 0.285266 | 0.018745 |
| SKP1 | 0.279302 | 1.81E-08 |
| CHCHD1 | 0.278017 | 0.010491 |
| TBC1D8 | 0.277317 | 0.045814 |
| CAPN1 | 0.277217 | 0.012045 |
| COX6C | 0.276189 | 0.000119 |
| TADA3 | 0.275318 | 0.039734 |
| KTN1 | 0.273805 | 0.03956 |
| GSTP1 | 0.273191 | 8.85E-07 |
| PRRG2 | 0.272514 | 0.007898 |
| RHBDD2 | 0.27208 | 0.009803 |
| ARPC2 | 0.271414 | 1.56E-08 |
| TECR | 0.270788 | 0.032731 |
| SLC39A2 | 0.270521 | 0.026588 |
| ATP5IF1 | 0.269697 | 0.01295 |
| UBL5 | 0.269453 | 0.001211 |
| M6PR | 0.26789 | 0.031074 |
| PPP4C | 0.267853 | 0.013803 |
| LAMTOR2 | 0.267266 | 0.01152 |
| ITGA2 | 0.264043 | 0.001265 |
| SMIM5 | 0.261883 | 0.030334 |
| FAM136A | 0.261681 | 0.048834 |
| TMEM167/ | 0.260509 | 0.006165 |
| ATP5MC3 | 0.257375 | 0.001232 |
| TMBIM6 | 0.255503 | 0.000384 |
| ATP5PF | 0.253313 | 0.031853 |
| PSMB3 | 0.252366 | 0.035038 |
| NDUFS5 | 0.251434 | 1.77E-06 |
| HNF1B | 0.251095 | 0.001452 |
