## Supplementary Table 3 for "Enhanced metanephric specification to functional proximal tubule enables toxicity screening and infectious disease modelling in kidney organoids"

| sample | condition | bead_radius | gloms_in_radius | ltl_in_radius | epcam_in_radius | tissue_in_radius |
|---|---|---|---|---|---|---|
| 1 | IWR | 1905017 | 495180 | 41898 | 92217 | 577495 |
| 2 | IWR | 1436790 | 592680 | 27171 | 69770 | 621571 |
| 3 | IWR | 1732366 | 746348 | 40231 | 99303 | 838595 |
| 1 | PBS | 1995803 | 722679 | 115372 | 241557 | 946907 |
| 2 | PBS | 1404122 | 651280 | 78934 | 203536 | 825990 |
| 3 | PBS | 1877655 | 734616 | 72331 | 204901 | 916286 |

| | |
|---|---|
| bead_radius | total number of pixels that are within 200 pixels from a bead edge. |
| gloms_in_radius | total number of pixels defined as NPHS1 positive, within 200 pixels from a bead edge. |
| ltl_in_radius | total number of pixels defined as LTL positive, within 200 pixels from a bead edge. |
| epcam_in_radius | total number of pixels defined as EPCAM positive, within 200 pixels from a bead edge. |
| tissue_in_radius | sum of total number of pixels defined as NPHS1 or EPCAM positive, within 200 pixels from a bead edge. |
